# The anti-tubercular callyaerins target the *Mycobacterium tuberculosis*-specific non-essential membrane protein Rv2113

**DOI:** 10.1101/2023.07.12.548660

**Authors:** David Podlesainski, Emmanuel T. Adeniyi, Yvonne Gröner, Florian Schulz, Violetta Krisilia, Nidja Rehberg, Tim Richter, Daria Sehr, Huzhuyue Xie, Viktor E. Simons, Anna-Lene Kiffe-Delf, Farnusch Kaschani, Thomas R. Ioerger, Markus Kaiser, Rainer Kalscheuer

**Affiliations:** Center of Medical Biotechnology, Chemical Biology, University Duisburg-Essen, Duisburg, Germany; Institute of Pharmaceutical Biology and Biotechnology, Heinrich Heine University, 40225 Düsseldorf, Germany; Department of Computer Science, Texas A&M University, College Station, Texas 77843, United States

## Abstract

Spread of antimicrobial resistances in the pathogen *Mycobacterium tuberculosis* remains a public health challenge. Thus, there is a continuous need for new therapeutic options with modes-of-action differing from current antibiotics. Previously, bioactivity-guided isolation identified the callyaerins, a class of hydrophobic cyclopeptides with an unusual (*Z*)-2,3-diaminoacrylamide unit, as promising antitubercular agents. In this study, we investigated the molecular mechanisms underlying their antimycobacterial properties. Structure-activity relationship studies enabled the identification of the structural determinants relevant for their antibacterial activity. The antitubercular callyaerins are bacteriostatics selectively active against *M. tuberculosis*, including extensively drug-resistant (XDR) strains, with minimal cytotoxicity against human cells and a promising intracellular activity in a macrophage infection model. Via spontaneous resistance mutant screens and various chemical proteomics approaches, we showed that they act by direct targeting of the non-essential, *M. tuberculosis*-specific putative membrane protein Rv2113, thereby triggering a complex stress response in *M. tuberculosis* characterized by global downregulation of lipid biosynthesis, cell division, DNA repair and replication. Our study thus not only identifies Rv2113 as a new *M. tuberculosis*-specific target for antitubercular drugs, which should result in less harm of the microbiome and weaker resistance development in off-target pathogens. It furthermore demonstrates that also non-essential proteins may represent efficacious targets for antimycobacterial drugs.

## INTRODUCTION

Tuberculosis, an ancient infectious disease caused by *Mycobacterium tuberculosis*, remains a global health threat responsible for 1.6 million deaths in 2021 [1]. Despite recent advances in tuberculosis chemotherapy with addition of bedaquiline, pretomanid and delamanid to the antimicrobial arsenal, lack of success in combating tuberculosis lingers [2]. This is partly due to the inability of drugs penetrating compartments harbouring sequestered bacilli, as well as the length and complexity of drug regimens [3]. A standard tuberculosis treatment, for instance, requires a cocktail of antibiotics, usually starting with a 2-month combination therapy of isoniazid, rifampicin, pyrazinamide and ethambutol, followed by a 4-month combination of rifampicin and isoniazid. This prolonged course of treatment often results in poor compliance, which consequently drives emergence of drug resistance [4, 5]. As a matter of fact, in 2021, there was an estimated 3.1% global increase in the number of multidrug-resistant and rifampicin-resistant tuberculosis (MDR/RR-TB) cases compared to the previous year, and about 191,000 deaths associated with MDR/RR-TB [1]. Due to the broad-spectrum activity of some of the antibiotics included in this 6-month tuberculosis regimen, particularly rifampicin exhibiting broad activity against Gram positive bacteria, resistance development can additionally transcend from *M. tuberculosis* to off-target pathogens [6]. Furthermore, long-term use of these broad-spectrum antibiotics causes major distress to the host microbiome, which can significantly impair the host’s immune system as a result of dysbiosis, thereby contributing to several ill health conditions in treated patients [7]. Thus, there remains a persistent need for new resistance-breaking therapeutic options in the ongoing fight against tuberculosis, particularly for antitubercular agents with a weaker resistance development in off-target pathogens and less harm to the microbiome.

Similar to many other clinically used antibiotics, many anti-TB drugs are natural product derivatives [8]. Even today, promising starting points for anti-tubercular drug discovery can therefore be identified by screening microbial organisms for anti-tubercular natural products, particularly if less-investigated habitats and ecological niches such as marine environments are explored, as this lowers the risk for simple rediscovery of known antibiotics [9, 10]. Such an approach led to the discovery of the callyaerins, an anti-tubercular proline-rich cyclopeptide family from *Callyspongia aerizusa*, a marine sponge of Indonesian origin [11–13]. Structurally, these cyclic peptides are composed of a ring system of eight to nine amino acids including an unusual (*Z*)-2,3-diaminoacrylamide unit (DAA, Fig. 1A) that is furthermore C-terminally linked to an exocyclic side chain of two to five amino acids. Among them, particularly callyaerin A and B (CalA, CalB, Fig. 1A) were found as promising anti-tubercular natural products as they possess potent antimycobacterial activity with an encouraging cytotoxic profile [11].

**Figure 1.**
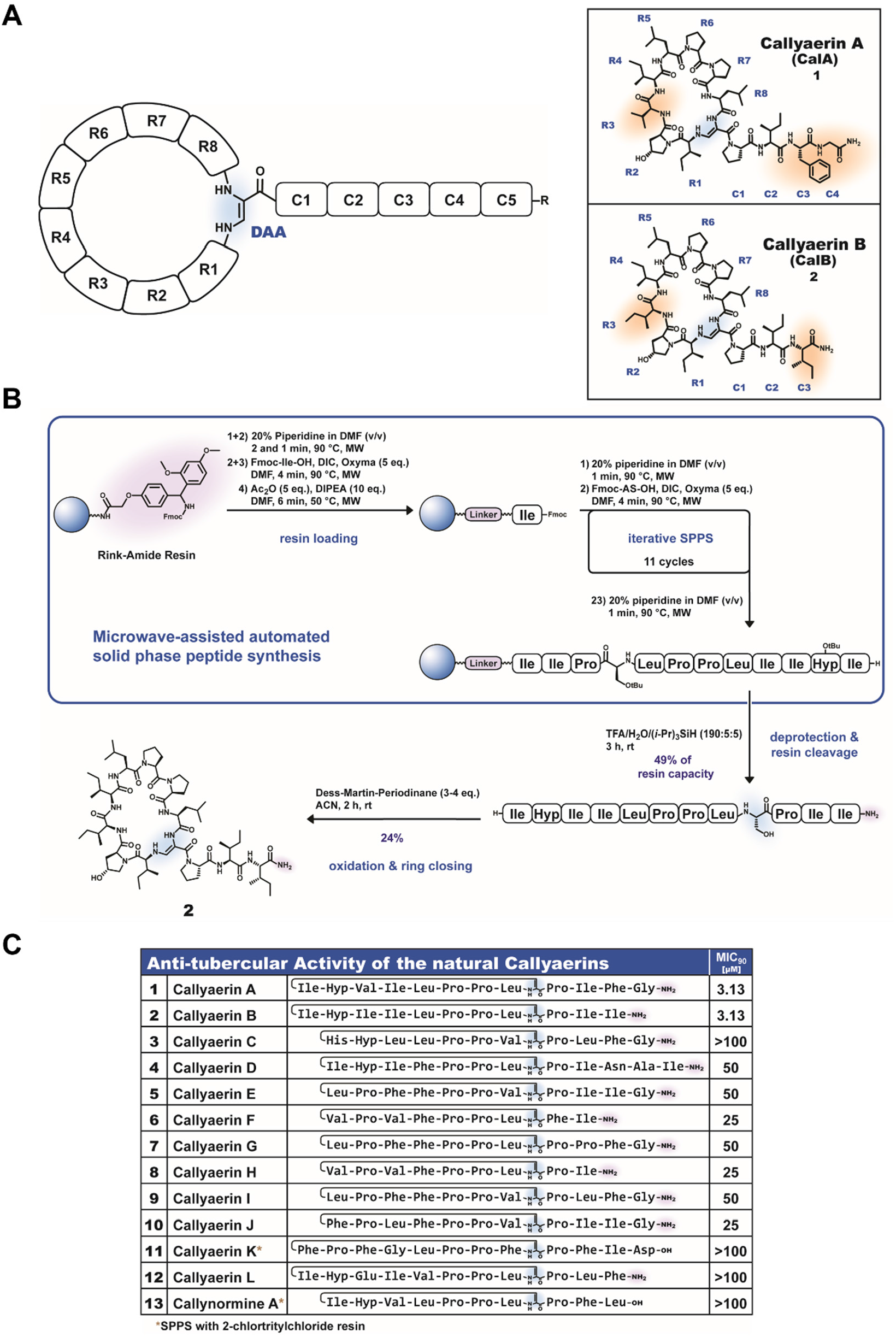
Callyaerin A and B are promising antitubercular agents. **A)** Generic structure scheme for the callyaerin natural product family, (hydroxy)proline rich hydrophobic cyclopeptides with a central (*Z*)-2,3-diaminoacrylic acid (DAA) unit, and chemical structure of callyaerin A (CalA) and B (CalB). Residues that are different between both compounds are indicated in orange, the DAA residue is depicted in blue. **B)** Scheme for the developed solid-phase peptide synthesis for callyaerins, here exemplarily presented for CalB. **C)** Overview on the growth inhibition properties (measured as MIC_90_ values using resazurin dye reduction assay) of chemically synthesized callyaerins vs. *Mycobacterium tuberculosis* H37Rv.

However, an in-depth characterization including structure-activity-relationships (SAR) and mode-of-action (MOA) studies of these anti-tubercular natural products has so far not been pursued. Here, we report the development of a solid phase synthesis protocol based on a modification of a previously published solid phase synthesis of CalA [14] that enabled a straightforward and flexible synthesis of various callyaerin derivatives, some with improved antitubercular activities, for SAR and MOA studies. By a combination of these resources with various genetic and chemical proteomic approaches, we then identified and validated the mycobacterial-specific protein Rv2113 as the direct antimycobacterial target of callyaerins in *M. tuberculosis* H37Rv. Of particular note, Rv2113 is fully dispensable for growth and viability of *M. tuberculosis*, demonstrating that non-essential proteins can represent efficacious targets of antibacterial drugs.

## RESULTS

### Callyaerin A and B display the highest anti-tubercular activity within the callyaerin natural product family

CalA and CalB have previously been identified as promising antitubercular agents (Fig. 1A) [11]. However, only five of the currently fourteen known callyaerins - due to high structural similarities, we count callynormine A [15] as a further callyaerin derivative - have until now been evaluated for their antimycobacterial properties. It therefore remained unclear whether other members of the callyaerin natural product members might represent more promising starting points for further antitubercular drug development.

To address this current limitation and to broaden the evaluation of the antitubercular potential of the callyaerins, we therefore aimed to establish a straightforward and flexible synthesis route to diverse members of the callyaerin natural product family as it may also be later used to generate customized derivatives for structure-activity relationship (SAR) studies. Recently, the first solid phase peptide synthesis of the callyaerin CalA has been reported [14]. This synthesis relied on the incorporation of a suitably protected, specifically synthesized α-formyl glycine building block during solid phase peptide assembly [16], which, after deprotection, allowed the efficient installation of the DAA unit through initial imino linkage and concomitant double-bond migration. Based on these findings, we developed a slightly modified, more straightforward callyaerin synthesis that consists of a solid phase synthesis of a precursor peptide featuring a standard serine residue at the future DAA position, followed by a one-pot selective Dess-Martin oxidation of the hydroxyl residue of serine and concomitant cyclization (Fig. 1B). This protocol allowed a flexible synthesis of diverse callyaerins typically at a 1-5 µmol scale. Notably, this approach exploits the unique chemical properties of the cyclized callyaerins that differ significantly from all other open-chain analogues (oxidized as well as non-oxidized), thereby enabling an efficient and easy purification of the final products by RP-HPLC.

With such a feasible synthesis route at hands, we subsequently synthesized thirteen of the so far fourteen known natural callyaerins (Supplementary Fig. 1, Supporting Information) and tested their *in vitro* antitubercular activity vs. the virulent *M. tuberculosis* strain H37Rv. Synthetic CalA and CalB were the most active callyaerins, showing potent minimum inhibitory concentrations for impairing at least 90% of growth compared to solvent controls (MIC_90_) of 1.56 - 3.13 µM, which are in agreement with previously published data for natural callyaerins (Fig. 1C) [11]. In contrast, all other tested callyaerins were significantly less active or even inactive (i.e., MIC_90_ > 100 µM), indicating that the macrocyclic ring size of eight amino acids plus the DAA residue in CalA and CalB, the overall fold of CalA and CalB with their distinct (hydroxyl)proline pattern as well as the presence of hydrophobic amino acids may be decisive for their anti-tubercular potential (Fig. 1C). We therefore decided to focus our following investigations on the hydrophobic ‘CalA/B-type callyaerins’ as they seem to represent the most promising antitubercular compounds from the callyaerin natural product family.

### CalA/B-type callyaerins are *M. tuberculosis*-selective, intracellularly acting bacteriostatics

We continued our investigations by characterizing the antibacterial and cytotoxicity profile of CalA/B-type callyaerins (Fig. 2A and Supplementary Fig. 2). Interestingly, we found that extensively drug-resistant (XDR) forms of *M. tuberculosis* were also susceptible to CalA and CalB with MIC_90_ ranging from 3.13 - 12.5 µM, indicating that these callyaerins do not share similar targets with clinical drugs to which these XDR forms of *M. tuberculosis* are resistant. The observed slight shift in MIC_90_ might not necessarily be caused by a reduced anti-tubercular activity against these drug-resistant *M. tuberculosis* strains, but due to their decreased fitness in the absence of the antibiotics to which they are resistant [17]. In contrast, only limited growth inhibition was found for other slow- and fast-growing mycobacteria species, including *Mycobacterium bovis* BCG Pasteur, *Mycobacterium smegmatis* and *Mycobacterium marinum*. CalA and CalB were also inactive against other tested Gram-positive (*Bacillus subtilis*, *Staphylococcus aureus*) or Gram-negative (*Escherichia coli*) bacteria, demonstrating a very narrow and specific activity against *M. tuberculosis* strains. Importantly, both natural products displayed a rather low cytotoxicity with IC_50_ values (i.e., concentrations required to inhibit 50% of growth relative to the solvent controls) ≥ 50 µM against a panel of human cell lines resulting in a promising selectivity index of 16-32.

**Figure 2.**
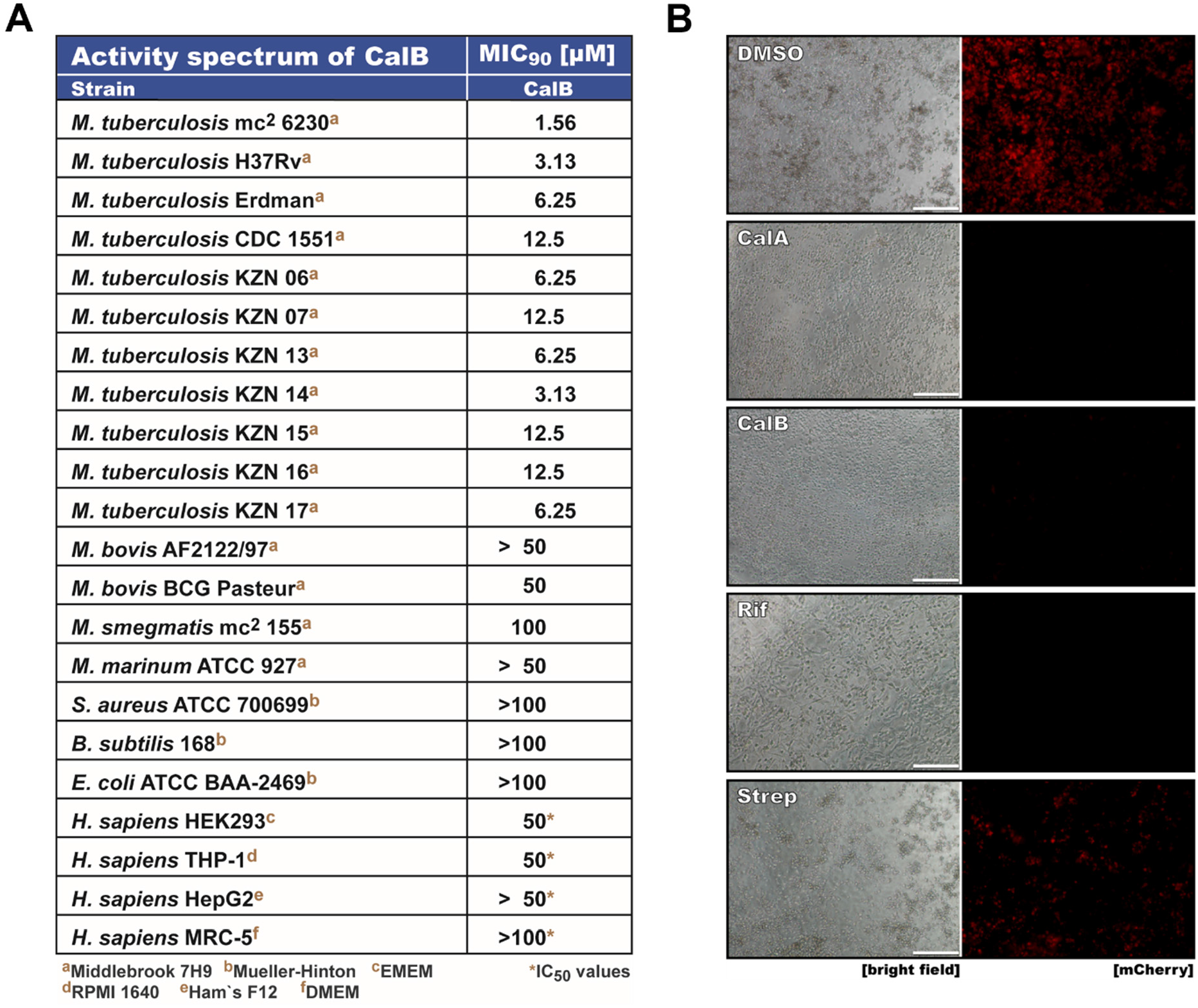
Callyaerins are *M. tuberculosis* selective anti-tubercular agents with intracellular growth inhibition properties. **A)** MIC_90_ values for CalB treatment of various *M. tuberculosis* strains including clinical XDR isolates and further mycobacterial, Gram-positive and Gram-negative strains as well as human cell lines. Media used for cultivation of the respective cells are indicated by brown superscripted letters. **B)** Fluorescence micrographs of intracellular activity of CalA and CalB in a macrophage infection assay. THP-1 cells were differentiated into macrophage-like cells using PMA. Cells were infected with a mCherry expressing reporter strain of H37Rv (multiplicity of infection MOI = 3) for 3 h and subsequently treated as indicated. Intracellular growth was evaluated five days post infection. Scale bar represents 50 µm.

We then performed a killing kinetic study by quantifying viable cell counts following different treatment intervals to determine how CalA/B-type callyaerins exert their growth inhibition effect on *M. tuberculosis* cells. We found that CalB displayed a bacteriostatic effect against cells of *M. tuberculosis* strain mc²6230 that was stable for two weeks (Supplementary Fig. 3).

As current TB treatment relies mainly on combination therapy [3], we also evaluated the *in vitro* efficacy of CalB in combination with other, clinically used anti-tubercular drugs. Although addition of CalB to isoniazid, rifampicin or bedaquiline did not enhance their bactericidal effects, the emergence of resistant mutants observed in monotreatment was effectively prevented (Supplementary Fig. 4). We also determined fractional inhibitory concentration indices (FICI) for the interaction of CalB with either isoniazid, rifampicin, bedaquiline, delamanid or ethambutol in the *M. tuberculosis* strains H37Rv and mc²6230 via a checkerboard assay (Supplementary Fig. 5). The resulting FICI values of 1 - 1.5 for all co-treatments suggested that CalB acts additive to these drugs, which is consistent with the killing kinetic experiment that showed no enhancement of bactericidal effects but suppression of resistance emergence by the bacteriostatic compound.

*M. tuberculosis* is an intracellular pathogen that mostly resides and replicates in phagolysosomes of macrophages [18, 19]. For evaluating the intracellular growth inhibition properties of CalA and CalB, we therefore employed a THP-1 human macrophage infection model system that relies on quantifying cell growth of a mCherry-expressing fluorescent *M. tuberculosis* reporter strain [20]. To this end, macrophages were treated three hours after infection with either DMSO, 15.6 µM CalA, 1.95 µM CalB or with the clinically used anti-TB drugs rifampicin (1 µM) or streptomycin (20 µM). Growth inhibition efficiency was then evaluated five days post infection using fluorescence microscopy (Fig. 2B and Supplementary Fig. 6 for quantification). While macrophages treated with the solvent control DMSO exhibited a heavy intracellular bacterial burden, appeared clumpy and started to detach from the surface, CalA and CalB both substantially inhibited intracellular proliferation of *M. tuberculosis* and resulted in a healthy morphology of the treated macrophages. The effects were comparable to that of the first-line anti-TB drug rifampicin. Remarkably, despite the usage of lower compound concentrations, the intracellular growth inhibition activity of CalA and CalB was superior to streptomycin, which could reduce mycobacterial growth only to about 20% of the DMSO-treated control compared to ca. 1% for CalA and CalB.

In summary, CalA/B-type callyaerins exhibit a bacteriostatic effect *in vitro*, show additive effects in combination with clinical anti-TB drugs, and can penetrate human macrophages for reaching *M. tuberculosis* cells in their phagosomal compartment, resulting in strong inhibition of intracellular growth.

### Structure-antitubercular activity determinants of CalA/B-type callyaerins

CalA and CalB as the founding members of the CalA/B-type callyaerin class are highly hydrophobic peptides that share most of their amino acid composition. Both are characterized by a 9-membered ring system cyclized via a DAA moiety and the conserved site-specific presence of (hydroxyl)proline residues, resulting in a unique and stable peptide fold [14]. They are structurally highly similar, differing only in their R3 and C3 position as well as by the presence of a C4 residue in CalA that is lacking in CalB (Fig. 1A). However, despite these structural differences, both cyclic peptides have a similar potency against *M. tuberculosis* H37Rv (MIC_90_ = 3.13 µM, Fig. 1C), indicating that at least certain positions tolerate structural modifications.

Taking advantage of our flexible chemical synthesis route, we therefore next performed a structure-activity relationship for studying their structural antitubercular determinants. We started with a CalA ‘alanine scan’, i.e., we replaced successively each amino acid of CalA by alanine and determined the resulting antimycobacterial activity for these derivatives (Fig. 3A). For most positions, introduction of an alanine residue resulted in a lower bioactivity, although in some cases (e.g., position R5, R7; see compound **18** and **20**, respectively), these effects were only moderate, while in other positions complete loss of bioactivity was observed (e.g., position R3, C2 and C3; see compound **16**, **23** and **24**, respectively). A notable exception is R4 (compound **17**); here, the substitution of the parent isoleucine residue by alanine was well tolerated and resulted in the same MIC_90_ values as observed for CalA and CalB. Also, an extension of CalA’s exocyclic chain with a further C5 alanine residue (**26**) had no impact on the bioactivity, suggesting that the ‘end region’ of the exocyclic chain may allow attachment of additional chemical tags without significant bioactivity loss.

**Figure 3.**
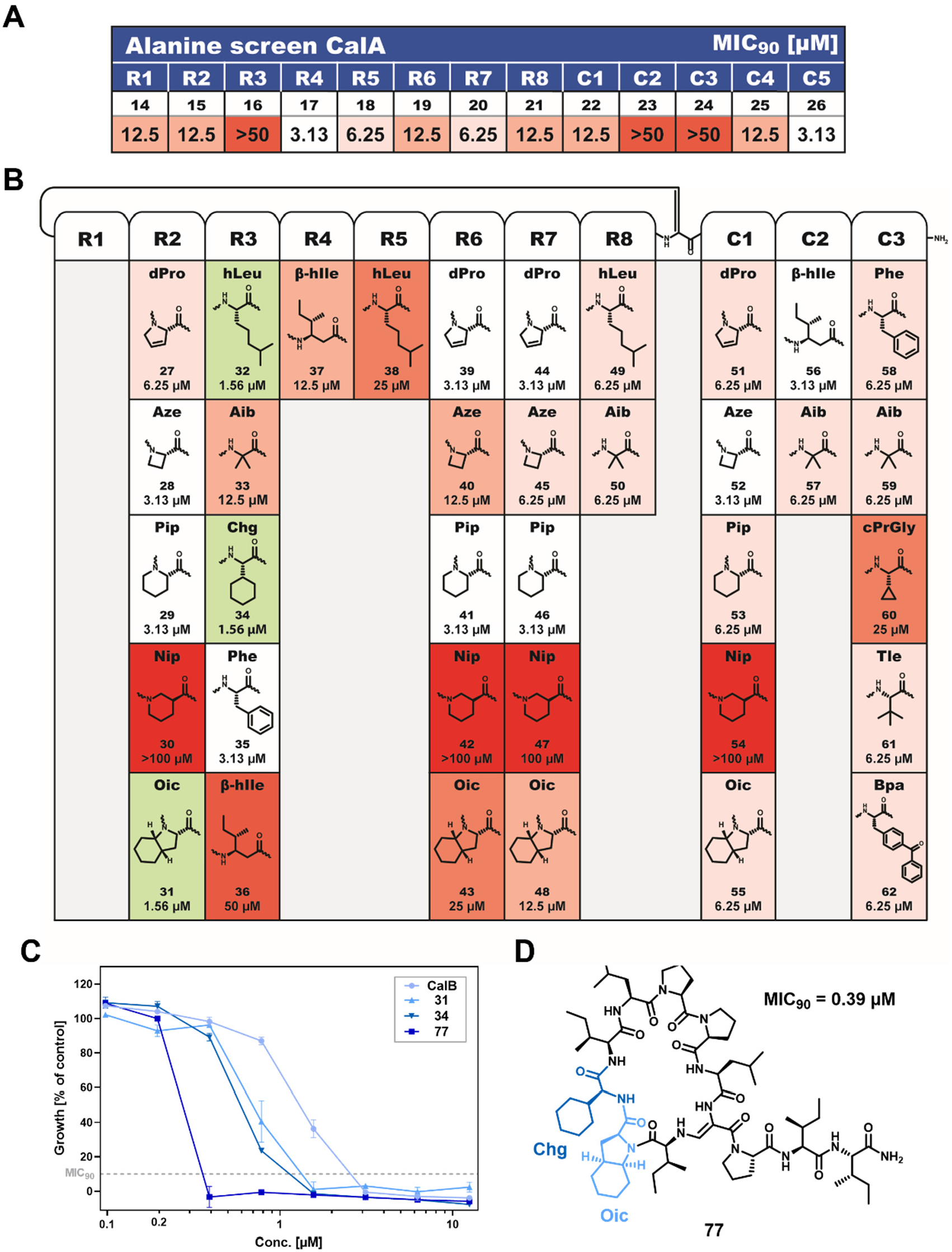
Overview on the structure-activity relationships of CalA/B-type callyaerins. **A)** An alanine scan of the different amino acids of CalA gives a first insight into structural determinants underlying bioactivity. **B)** Overview on single amino acid substitutions performed to study structure-activity-relationships. Further evaluated derivatives are reported in Supplementary Fig. 7. **C)** Concentration-dependent growth inhibition curve for CalB, the slightly more active derivatives **31** and **34** as well as of compound **77** harbouring both structural modifications. Data shown as means of duplicates with SD depicted as error bars. **D)** Chemical structure of compound **77** with a cyclohexylglycine (Chg, dark blue) modification in R3 and an octahydroindole-2-carboxylic acid (Oic, light blue) modification in R2. This rationally designed compound is ten-times more active than the parent compound CalB, displaying a MIC_90_ of 0.39 µM vs. *M. tuberculosis* H37Rv.

We then performed single amino acid substitutions at selected positions within CalB to better test position-dependent structural determinants (Fig. 3B and Supplementary Fig. 7 for further CalA derivatives). We mainly focused on conservative substitutions, i.e., we replaced hydrophobic and/or cyclic amino acids with chemically similar amino acids; these substitutions also included a change of the (hydroxy)proline residues with other cyclic amino acids. These studies revealed that the hydroxyproline residue at R2 is not essential for bioactivity and can also be replaced by a ‘standard’ proline residue (**63**). Indeed, (hydroxyl)proline substitutions by other cyclic amino acids such as dehydroproline or azetidine carboxylic acid were often well tolerated. The incorporation of β-amino acids, in particular nipecotic acid (Nip), however led mostly to fewer active compounds, with the exception of the incorporation of β-hIle at C2 (**56**) that resulted in a derivative with CalA/CalB-similar bioactivities, again highlighting the exocyclic chain end as a region amenable for modification. Most tested amino acid substitutions, however, resulted in derivatives with equal or, more frequently, lower bioactivities compared to the parent compound CalB. Only three substitutions led to moderate, 2-fold more active derivatives: i) a substitution of hydroxyproline at R2 by octahydroindole-2-carboxylic acid (Oic, **31**), and the substitution of isoleucine at R3 by either homoleucine (hLeu, **32**) or cyclohexyl glycine (Chg, **34**, Fig. 3B and 3C). In the case of the amino acid Chg, this effect was also confirmed in the CalA structure (**65**). Importantly, these bioactivity improvements were found to be additive as the designed callyaerin derivative **77** containing the Oic-substitution at R2 and the Chg-substitution at R3 was roughly 10-times more active than the parent compound CalB (Fig. 3C and 3D). Overall, structure-activity-relationships seem particularly driven by hydrophobicity; this notion is further substantiated by the finding that an incorporation of, for example, an aspartate residue at the R5 position (**69**) or a tryptophane residue at the R8 position of CalA (**70**), completely abolishes activity (Supplementary Figure 7A) and, complementarily, Callyaerin K (**11**) and Callyaerin L (**12**), which both belong to the CalA/B-callyaerin family but harbour acidic amino acids in their structure (Fig. 1C), exhibit as well no activity.

Altogether, CalA/B-type callyaerins display distinct structure-activity-relationships, as improper substitutions, even of hydrophobic amino acids with similar chemical properties, strongly reduce bioactivity, thereby indicating that their mode-of-action relies on a defined interaction with one or more distinct target protein(s). The strong preference for hydrophobic amino acids might be a consequence for the need for membrane passage; in addition, it suggests that CalA/B-type callyaerins will allocate to a highly hydrophobic binding site on their target protein.

### *Rv2113* mediates resistance and susceptibility to CalA and CalB

In order to identify the direct target(s) of the antitubercular callyaerins, spontaneous single-step mutants of *M. tuberculosis* H37Rv were isolated after four weeks from 7H10 medium containing a 5-fold MIC of either CalA or CalB, respectively, at a frequency of 10^−6^. In contrast to the wild type, the obtained mutants were highly resistant to CalA or CalB treatment with MIC_90_ values above 50 µM (Supplementary Fig. 8A and B) and also showed cross-resistance to the other callyaerin (Supplementary Fig. 8C), suggesting a common target for both compounds. Subsequent whole-genome sequencing of randomly selected mutants consistently revealed non-synonymous mutations in the gene *Rv2113* (Fig. 4A). In the majority of the analyzed clones, *Rv2113* mutations were accompanied by further mutations; these mutations, however, occurred in diverse genes, strongly suggesting that mutations in *Rv2113* are indeed the molecular cause for callyaerin resistance. The mutations included non-synonymous SNPs resulting in diverse amino acid substitutions (T185A, L28P, L341P, C58R) as well as single nucleotide +c insertions or -c deletions in a 6c homopolymer region 20 nucleotides downstream of the start of the *Rv2113* gene resulting in frame shifts. The bandwidth of identified mutations suggests a loss-of-function-based resistance mechanism.

**Figure 4.**
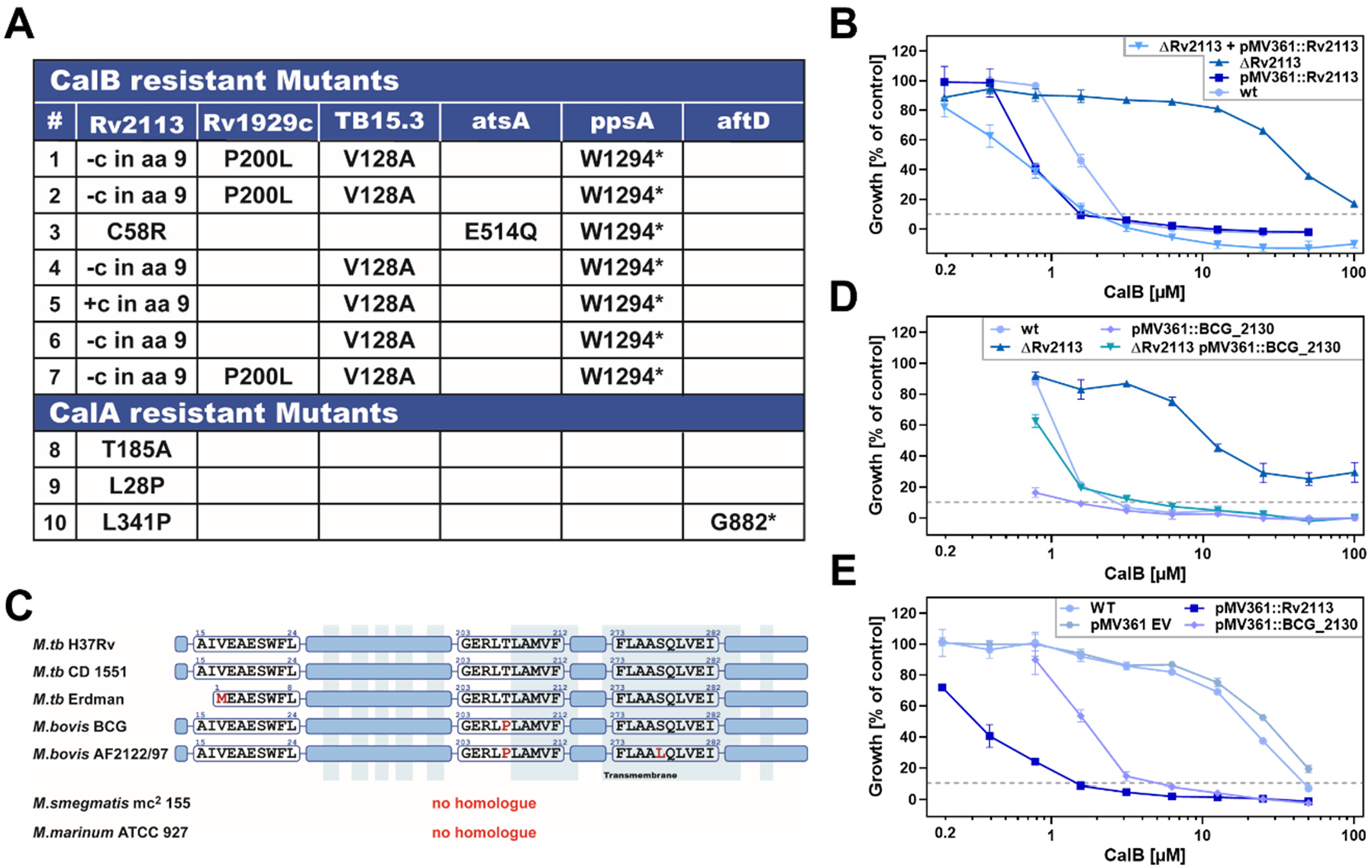
Rv2113 mediates resistance and susceptibility to CalA and CalB. **A)** Overview on the results from screening of spontaneous CalA- and CalB-resistant *M. tuberculosis* H37Rv mutants. Spontaneous single-step resistant mutants emerged at a frequency of 10^−6^ on 7H10 agar containing concentrations corresponding to a 5-fold MIC of CalA or CalB, respectively, after 4-week incubation. The corresponding mutations found in H37Rv were mapped via whole-genome sequencing. **B)** Dose-response curves for CalB concentration-dependent growth inhibition of H37Rv strains. The assay included the wild type, the *ΔRv2113* gene deletion mutant, the complemented strain *ΔRv2113* pMV361::*Rv2113* (treated with CalA) and the overexpressing strain pMV361::*Rv2113*. Data shown as means of duplicates with SD depicted as error bars. **C)** Overview on the sequence variations found in Rv2113 homologs in other mycobacterial strains. **D)** Dose-response curves for CalB concentration-dependent growth inhibition of H37Rv strains. The assay included the wild type, the *ΔRv2113* gene deletion mutant, the pMV361::*BCG_2130* complemented mutant strain *ΔRv2113* and a BCG_2130 overexpressing strain pMV361::*BCG_2130*. Data shown as means of duplicates with SD depicted as error bars. **E)** Dose-response curves for CalB concentration-dependent growth inhibition of *M. bovis* BCG Pasteur strains. The assay included the wild type, the empty vector control pMV361 EV and the overexpressing strains pMV361::*Rv2113* and pMV361::*BCG_2130.* Data shown as means of duplicates with SD depicted as error bars.

To prove the relevance of *Rv2113* for CalA and CalB resistance, we next generated a site-specific *Rv2113* gene deletion mutant in *M. tuberculosis* H37Rv, using specialized transduction (Supplementary Fig. 9). As expected, the generated independent clones of the *M. tuberculosis* Δ*Rv2113* gene deletion mutant were found to be resistant to both CalA (Supplementary Fig. 10) and CalB treatment (Fig. 4B), with MIC_90_ values > 100 µM. Upon complementation of the deletion mutant with a wild-type copy of *Rv2113* constitutively expressed from a single-copy integrative plasmid (Δ*Rv2113* pMV361::*Rv2113*), sensitivity to CalB was restored, thus unambiguously linking the resistance phenotype exclusively to absence of Rv2113. Finally, when the same plasmid pMV361::*Rv2113* was used to generate a merodiploid strain of *M. tuberculosis* H37Rv that overexpresses Rv2113, this led to enhanced susceptibility (2-fold decrease in MIC) to CalB. Although *Rv2113* is not essential for viability, these results suggest that both resistance and susceptibility of *M. tuberculosis* to CalA/B-type callyaerins are intimately linked to Rv2113.

To confirm these findings, we also raised spontaneous CalB-resistant mutants in the generated merodiploid *M. tuberculosis* H37Rv strain harbouring pMV361::*Rv2113*, reasoning that the presence of a second copy of *Rv2113* might force the occurrence of further mutations in other genes. However, we again found relevant non-synonymous SNPs only in *Rv2113* located either in the endogenous or in the merodiploid gene copy, causing A217V and L338P amino acid substitutions, respectively (Supplementary Fig. 11). Importantly, the spontaneous CalB-resistant mutants raised in the *Rv2113*-merodiploid strain exhibited the same MIC as the *M. tuberculosis* wild type when screened against bedaquiline and rifampicin, indicating that the mutations in *Rv2113* do not cause broad-spectrum resistance but specifically only affect susceptibility towards callyaerins (Supplementary Fig. 12). These findings emphasize the crucial role of the non-essential protein Rv2113 in resistance and susceptibility of *M. tuberculosis* to CalA/B-type callyaerins, while no other relevant mutations located in other genes could be identified.

To understand the basis for the selectivity of callyaerins against *M. tuberculosis*, we searched for Rv2113 homologs in the UniProtKB proteome via BLAST [21]. Notably, we found homologues (sequence identity ≥ 95%) of Rv2113 only in a small number of organisms including *M. tuberculosis*, *M. bovis*, *M. orygis*, *M. canettii* and *M. shinjukuense*. Except these mycobacterial strains, the search revealed no further organisms with Rv2113 homologues with more than 70% sequence identity. Rv2113 is annotated as a ‘conserved probable membrane protein’ that, according to sequence analysis and Alphafold prediction [22], harbours eight transmembrane helixes. We previously tested CalA and CalB vs. various (myco)bacterial strains (Fig. 2A). An analysis on the presence of Rv2113 homologs in these strains revealed that apart from the susceptible *M. tuberculosis* strains, Rv2113 homologs are only present in the virulent *M. bovis* AF122/97 (with ‘Mb2137’ corresponding to Rv2113) and non-virulent *M. bovis* BCG Pasteur (with ‘BCG_2130’ corresponding to Rv2113) strains (Fig. 4C). In these strains, however, CalA and CalB were much less active. Although Mb2137 and BCG_2130 show high sequence identities to Rv2113, BCG_2130 harbours one (T207P) and Mb2137 two amino acid substitutions (T207P, S278L). Owing to the fact that the amino acid substitution T207P is common to both *M. bovis* strains, we reasoned that this mutation might account for the low susceptibility (MIC_90_ = 50 µM) of *M. bovis* BCG Pasteur to CalA/B-type callyaerins. We thus heterologously expressed *BCG_2130* in the *M. tuberculosis* Δ*Rv2113* mutant (Δ*Rv2113* pMV361::*BCG_2130*), and observed that this fully restored sensitivity to CalB (Fig. 4D). Further, when *BCG_2130* or *Rv2113* were overexpressed in *M. bovis* BCG Pasteur via a strong constitutive promoter from the single-copy integrative plasmids pMV361::*BCG_2130* and pMV361::*Rv2113*, respectively, this strongly sensitized *M. bovis* BCG Pasteur towards CalB, with a reduction in MIC_90_ from 50 µM to 6.25 µM or 1.56 µM, respectively (Fig. 4E). These findings together suggest that the basis of the poor sensitivity of *M. bovis* BCG Pasteur toward CalA/B-type callyaerins is likely tied to a low expression level of endogenous *BCG_2130*, whereas the single amino acid exchange in BCG_2130 compared to Rv2113 itself does not substantially impair interaction with CalB. However, while the expression level of *Rv2113* or its homologue *BCG_2130* determines sensitivity towards callyaerins in *M. tuberculosis* and *M. bovis* BCG Pasteur, heterologous expression of *BCG_2130* or *Rv2113* did not or only slightly increased susceptibility of *M. smegmatis* towards callyaerins (Supplementary Fig. 13). Similarly, heterologous expression of *Rv2113* in *B. subtilis* did not sensitize this bacterium towards CalB (Supplementary Fig. 14). This suggests that the interaction of CalA/B-type callyaerins with Rv2113 is necessary, but alone not sufficient to establish an antibacterial effect; instead, additional factors that are present in *M. tuberculosis* and *M. bovis* BCG Pasteur, but absent in other bacteria, are required for antitubercular activity.

### CalB directly targets Rv2113

The genetic studies showed that Rv2113 mediates CalB resistance and susceptibility. It is however not clear if this phenotype is caused by a direct interaction with Rv2113 or is a more indirect effect. To prove direct target engagement of CalB in mycobacteria, we therefore devised a chemical proteomics strategy in which suitably modified CalB derivatives were used for direct target identification (Fig. 5A). In this approach, two C4-alkyne tagged CalB derivatives, one with a native leucine residue (**82**, Fig. 5B) and the other with an additional diaziridine-based ‘photoleucine’ residue in R5 (**85**), were used in a (photo)affinity enrichment-LC-MS/MS-based target identification. To this end, a two-step labelling procedure was used in which the alkyne tag was modified with a biotin residue via click chemistry after *in situ* labelling. Accordingly, while **82** could be employed to identify proteins that are targeted by a potential covalent mechanism, e.g., via the DAA unit of the callyaerins by a so far unclear mechanism, **85** could be used to elucidate non-covalent target proteins via an additional photoaffinity labelling step (Fig. 5A). The design of the corresponding probes was deduced from the previous structure-activity-relationship studies (Supplementary Fig. 7C), suggesting the R5 position as a suitable site for the incorporation of the photoleucine residue (Supplementary Fig. 15). Indeed, **82** as well as **85** both efficiently inhibited mycobacterial growth with MIC_90_ values of 6.25 and 12.5 µM, respectively, thus being only slightly less active than the parent compound CalB.

**Figure 5.**
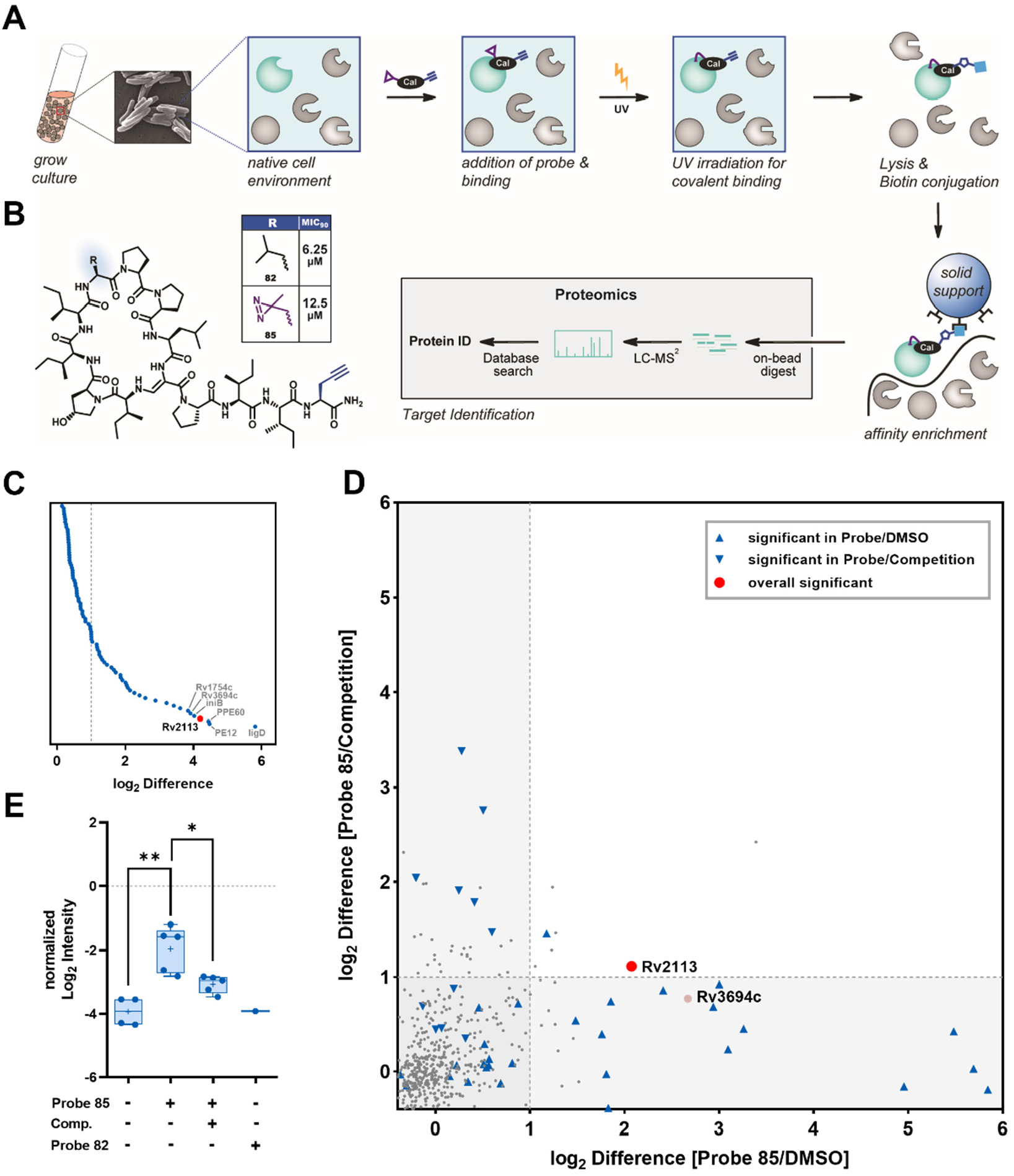
Photoaffinity labelling reveals Rv2113 as a direct target of CalA/B-type callyaerins. **A)** Overview of the workflow for (photoaffinity) labelling of *M. tuberculosis* H37Rv cultures. **B)** Chemical structures of the probes **82** and **85** and their respective MIC_90_ values against *M. tuberculosis* H37Rv. Probe **85** is a photoaffinity probe that besides the alkyne tag for click chemistry also harbours an aziridine-based photoleucine in position R5. **C)** Significant enriched Proteins (P ≤ 0.05 after Student’s t-test using Permutation-based FDR with 250 randomizations and FDR = 0.01) resulting from an affinity-based protein-profiling experiment with 10 µM probe **85** in *M. tuberculosis* H37Rv sorted by their log_2_ FC. **D)** Results from a combined photoaffinity labelling with 12.5 µM **85** (log_2_ FC plotted on x-axis) and competitive photoaffinity labelling approach with application of 30 µM CalB prior to photoaffinity labelling (log_2_ FC plotted on y-axis). Dashed lines indicate the log_2_ FC > 1 threshold. To identifying statistically significant hits from the analysis (marked in blue or red), P ≤ 0.05 (Student’s t-test; Permutation-based FDR with 250 randomizations and FDR = 0.01) was applied. **E)** Boxplot representation of Rv2113 quantification data from the (photoaffinity) labelling experiments. n = 5 technical replicates, asterisks indicate significant differences between indicated samples (** p ≤ 0.01, * p ≤ 0.05, unpaired t-test with Welch’s correction).

To investigate whether CalA/B-type callyaerins exert their bioactivity via a potential covalent interaction mechanism, we first performed a pulldown with **82** but were not able to identify protein targets with this approach (Supplementary Fig. 16). This suggests that CalA and CalB most probably interact via a non-covalent mechanism.

Accordingly, we repeated the experiment with the photoaffinity-tagged probe **85**. Application of 10 µM of the probe to a *M. tuberculosis* H37Rv culture for 3 hours, followed by 20 min UV radiation at 365 nm, cell lysis, click reaction with biotin azide and finally an avidin-based affinity enrichment and LC-MS/MS analysis indeed revealed several significantly enriched targets, with log_2_ fold changes (FC) > 1 as a threshold, among them Rv2113 as one of the top hits (Fig. 5C). For further validation of these initial hits, a second photoaffinity labelling approach was performed at a 12.5 µM probe concentration that was combined with a second, competitive labelling approach that relied on a preincubation with 30 µM CalB (corresponding to a more than 2-fold excess of the competitor) prior to application of the photoaffinity probe **85** (Fig. 5D, Supplementary Fig. 17). Importantly, this combination of both experiments revealed Rv2113 as the only significantly enriched direct target with log_2_ FC > 1 in both experimental setups (Fig. 5D and Fig 5E). Consistent with our previous data, probe **85** showed a significant reduction of activity against our *ΔRv2113* gene deletion mutant (Supplementary Fig. 18A). However, labelling of the ΔRv2113 mutant showed similar enrichment results apart from Rv2113 (Supplementary Fig. 18B). This non-specific labelling might be caused by the highly active carbene species generated during photoactivation of diazirine.

Altogether, these chemical proteomics experiments thus corroborated Rv2113 as a target of CalA/B-type callyaerins. In addition, they prove a direct molecular interaction between this target and the compounds.

### Rv2113 does not mediate intracellular uptake of CalA/B-type callyaerins

Our results so far point towards a central role of Rv2113 in the antibacterial mechanism of CalA/B-type callyaerins. However, whereas most antitubercular antibiotics typically work by inhibiting the function of an essential protein, this cannot straightforwardly explain the molecular mode of action of CalA/B-type callyaerins in *M. tuberculosis* as the Δ*Rv2113* mutant is fully viable and shows no growth defects. We therefore first asked if the membrane protein Rv2113 acts as a molecular transporter of CalA and CalB, thereby mediating intracellular uptake of these compounds, which would allow the compounds to subsequently bind to the ‘real’ targets inside the mycobacterial cell.

To investigate this potential mechanism, we synthesized fluorescent CalB derivatives by conjugating the azide-tagged fluorescent dyes Cy3 or rhodamine via a click reaction to a propargylglycine residue in the C4 position of CalB (compound **82**, Fig. 6A and Supplementary Fig. 19). While the rhodamine derivative **88** was found to be inactive (MIC_90_ > 100 µM), the Cy3 derivative **87** (Fig. 6A) was found to be more than 30-fold more potent than CalB (MIC_90_ = 0.05 µM, Fig. 6B). This improvement in antitubercular activity was a result of the appropriate covalent modification of CalB with Cy3 as neither application of a respective Cy3 control (**89**) alone nor of a 1:1 molar mixture of CalB and Cy3 resulted in comparable MIC_90_ values (Fig. 6B). This improved antitubercular activity furthermore seems to be a specific effect as analogous incorporation of a Cy3 residue at the R3 (**90**) or R4 (**91**) position of CalB again led to less active compounds (Supplementary Fig. 20). Importantly, the Cy3-CalB derivative **87** still inhibited the growth of *M. tuberculosis* in an Rv2113-dependent manner, since the *ΔRv2113* mutant exhibited resistance (Fig. 6C), demonstrating that the dye conjugate most likely relies on the same mechanism as the parent CalB. **87** also maintained its low cytotoxicity to the human cell lines THP-1 and HEK293, resulting in an impressive selectivity index (IC_50_/MIC_90_) of > 125.

**Figure 6.**
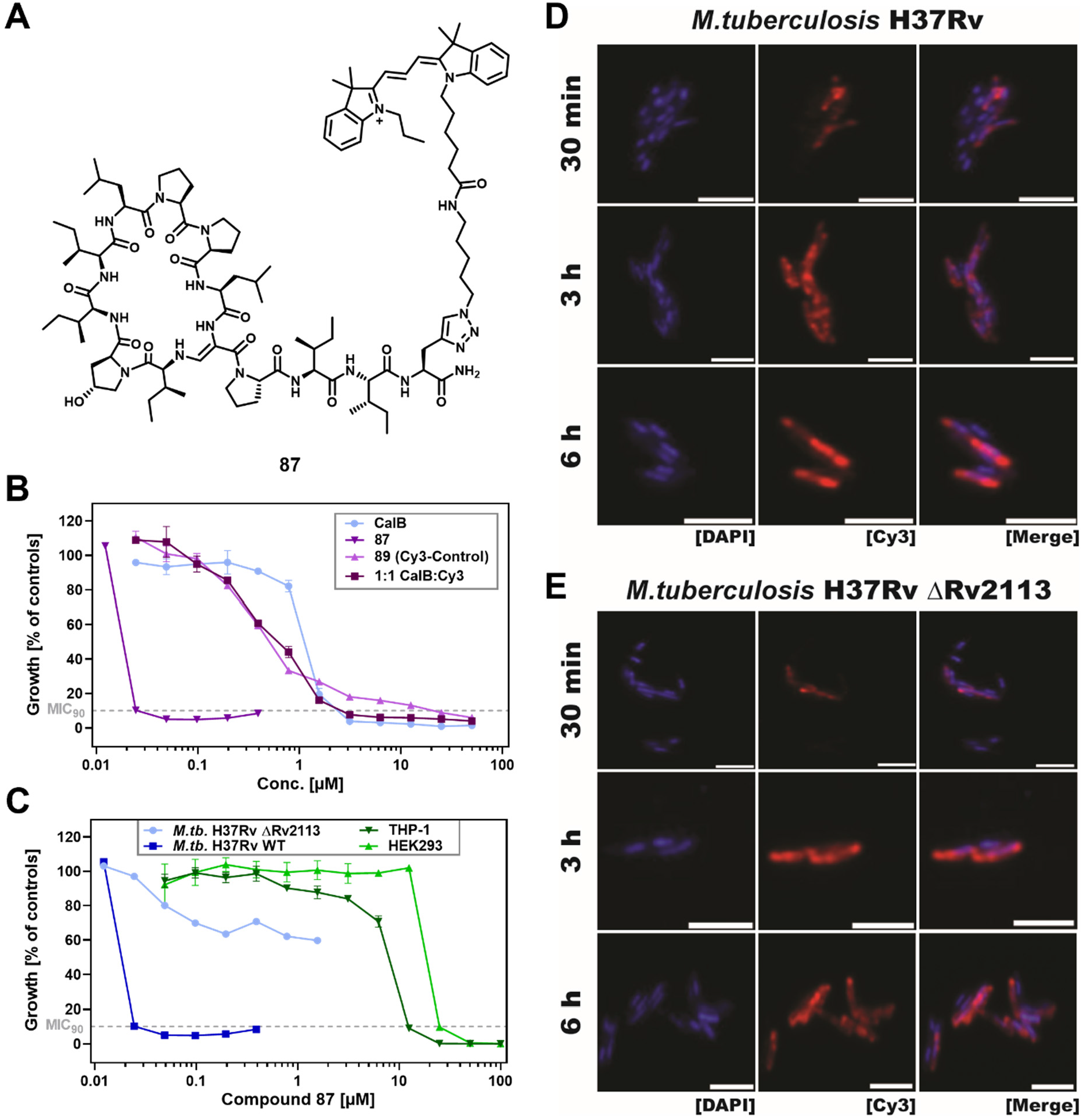
Rv2113 is no molecular transporter for CalB. **A)** Chemical structure of the most active Cy3-CalB conjugate **87**. In this derivative, the Cy3 residue has been conjugated to the C4 position via a Click reaction on a propargyl glycine residue. **B)** Dose-response curves for concentration-dependent growth inhibition of H37Rv strains by the Cy3-CalB derivative **87**, CalB, a Cy3 control (**89**) and a 1:1 mixture of CalB and Cy3. Data shown as means of duplicates with SD depicted as error bars. **C)** Dose-response curves for the **87**-mediated concentration-dependent growth inhibition of cells of *M. tuberculosis* H37Rv wild type and the Δ*Rv2113* mutant as well as of the human cell lines THP-1 and HEK293. **D** and **E)** Fluorescence microscopy images of cells of *M. tuberculosis* H37Rv wild type (**D**) and the Δ*Rv2113* mutant (**E**) treated with 0.2 µM of **87** for the indicated time intervals. After incubation, treated cells were counterstained with DAPI to label all cells. Scale bar represents 3 µm.

With this highly active and fluorescent derivative in hands, we therefore next used fluorescence microscopy to measure compound uptake in *M. tuberculosis* H37Rv cells as well as in the *ΔRv2113* strain (Fig. 6D and Fig. 6E). Intriguingly, **87** was rapidly internalized both by cells of *M. tuberculosis* wild type and the Δ*Rv2113* mutant, with intracellular staining clearly visible as early as after 30 min of incubation. Thus, CalA/B-type callyaerins likely enter *M. tuberculosis* cells independently of Rv2113, ruling out an uptake-dependent mechanism of susceptibility and resistance. As a negative control experiment, we also performed uptake studies with the inactive rhodamine-conjugated derivative (Supplementary Fig. 21). We were not able to detect rhodamine-mediated fluorescence in any of the both strains, thus demonstrating that the observed fluorescence signal from **87** correlates with uptake and not unspecific cell surface binding.

In summary, these experiments demonstrate that Rv2113 does not act as a membrane transporter mediating uptake of CalA or CalB, but as a functional target that is modulated upon binding of these compounds.

### CalB binding to Rv2113 modulates multiple major cellular processes in *M. tuberculosis*

While all experiments so far illustrate the relevance of Rv2113 for the antitubercular activities of CalA/B-type callyaerins, it remains unclear how this interaction translates into a growth inhibitory effect.

Having ruled out an uptake-related role of Rv2113, our first hypothesis was that the interaction of CalA or CalB with Rv2113 might deregulate structure and function of this membrane protein, finally resulting in a loss of membrane integrity; such a mechanism is often observed for ‘membrane active’ antibiotics that trigger an efflux of metabolites and a collapse of the membrane potential. To address this hypothesis, we therefore performed a propidium iodide internalization experiment, which however failed to detect relevant membrane permeabilization in CalB-treated *M. tuberculosis* cells (Supplementary Fig. 22). A lowering of intracellular ATP levels upon CalB treatment as a result from an impairment of energy metabolism of *M. tuberculosis* cells or a reduced ATP production due to a collapse of the membrane potential was also not observed (Supplementary Fig. 23). These findings are corroborated by the fact that compounds affecting bacterial membrane integrity typically exert a strong bactericidal effect [23], while the effect of CalA and CalB on *M. tuberculosis* cells is bacteriostatic. Finally, we also measured the expression of the genes *iniA* and *iniB* from the *iniBAC* gene operon as markers for cell wall stress in mycobacteria after CalB application [24]. While these genes were found to be strongly upregulated by the positive control isoniazid, a known inhibitor of cell wall mycolic acid biosynthesis, CalB treatment led to the downregulation of both marker genes, similar to the negative control rifampicin, an RNA polymerase inhibitor with no direct effect on cell wall structure (Supplementary Fig. 24). In conclusion, we found no evidence that CalA/B-type callyaerins interfere with membrane integrity or cell wall formation in *M. tuberculosis*.

We therefore concluded that CalA/B-type callyaerins act by modulating the molecular function of Rv2113, raising the question as to what is the molecular function of Rv2113 in *M. tuberculosis*. To address this, we therefore performed a comparative full proteome analysis of the *ΔRv2113* gene deletion mutant and the complemented strain *ΔRv2113* pMV361::*Rv2113* (Fig. 7A). Since inadvertent secondary mutations are known to potentially occur during genetic manipulation of *M. tuberculosis*, comparison of these two strains (rather than comparison of the wild type vs. *ΔRv2113* mutant strain) prevents misinterpretations arising from potential secondary mutations or polar effects in the *ΔRv2113* strain, which could result in altered protein abundances unrelated to absence of Rv2113. The analysis revealed only relatively mild overall changes, in accordance with the fact that the *ΔRv2113* knockout strain is fully viable and grows similar to the wild type. The most prominent difference was observed for the Mas protein, a large mycocerosic acid synthase which plays an important role in the synthesis of mycolic acids that constitute the outer membrane of mycobacteria, which was found downregulated in the *ΔRv2113* strain. Further downregulated proteins were Rv1779c, a possible integral membrane protein, and two further proteins involved in lipid biosynthesis, i.e., GrcC2, a dimethylallyltranstransferase, and PrpD, a 2-methylcitrate dehydratase. Rv0073, NadD, PapA5, CydB, MftE and MftF as well as NuoA were found at increased levels in the *ΔRv2113* strain. Of these proteins, PapA5, a phthiocerol/phthiodiolone dimycocerosyl transferase, is also a protein involved in lipid synthesis while the other proteins seem to be involved mainly in diverse redox reactions. Altogether, this analysis seems to suggest that Rv2113 may play a so far unknown non-essential, accessory role in lipid biosynthesis or transport.

**Figure 7.**
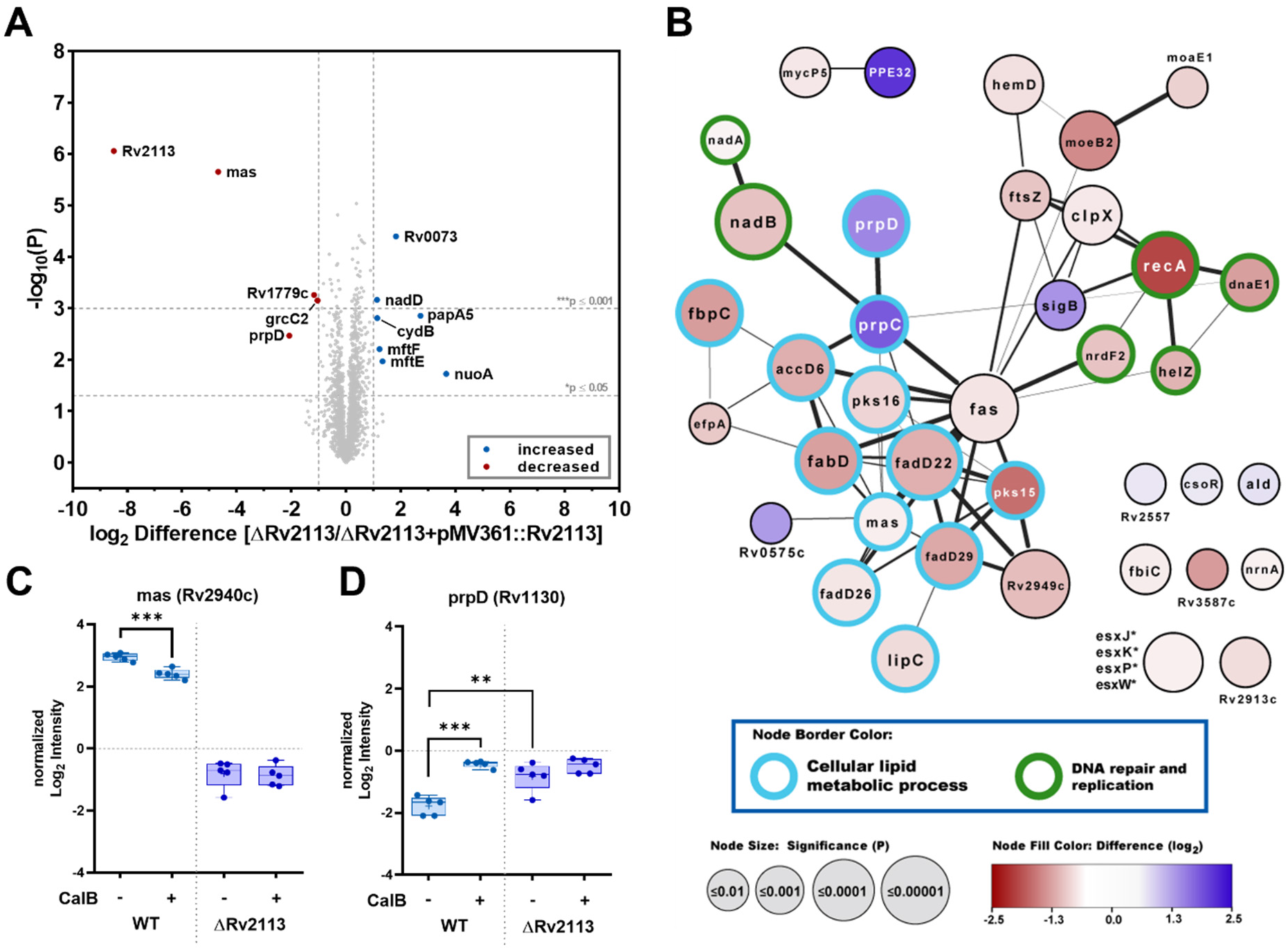
Rv2113-dependent stress response profile suggests that CalB simultaneous impairs multiple pathways. **A)** Full proteome analysis of a complemented strain *M. tuberculosis* H37Rv *ΔRv2113* pMV361::*Rv2113* vs. the corresponding *ΔRv2113* strain. The volcano plot shows the fold-change (log_2_) in protein abundance, plotted against p value (-log_10_). Log_2_ FC values < −1 and > 1 with p < 0.05 (Student’s T-test; Permutation-based FDR with 250 randomizations and FDR = 0.01) are specifically labeled. Proteins more abundant in the *ΔRv2113* strain show positive log_2_ FC values while proteins less abundant have negative values. **B)** Network visualization of proteins which comply the chosen threshold (log_2_ FC ≥ 0.5 or ≤ −0.5, q-value ≤ 0.05; Student’s T-test; Permutation-based FDR with 250 randomizations and FDR = 0.05) from a stress response full proteome profiling of wild type *M. tuberculosis* H37Rv cells treated for 48 h with 31.3 µM CalB (corresponding to a 10-fold MIC_90_) vs. DMSO treatment after STRING analysis. Node color represents the log_2_-FC in protein abundance (increase showed in blue, decrease in red) and node size corresponds to significance (P). Thickness of connection lines indicated the confidence of STRING protein-protein association. Proteins assigned to the GO cluster ‘Cellular lipid metabolic process’ [GO: 0044255] or related to DNA repair and replication are shown with a turquoise or green border, respectively. **C** and **D)** Boxplot presentation of Mas (**C**) and prpD (**D**) quantification data from stress response full proteome profiling of wild type and ΔRv2113 *M. tuberculosis* H37Rv cells treated for 48 h with 31.3 µM CalB (corresponding to a 10-fold MIC_90_) vs. DMSO. n = 5 technical replicates, asterisks indicate significant differences between indicated samples (** p ≤ 0.01, *** p ≤ 0.001, unpaired t-test with Welch’s correction).

To obtain insights into the cellular processes that are specifically affected by CalB treatment in a Rv2113-dependent manner, cells of *M. tuberculosis* H37Rv wild type (Fig. 7B, Supplementary Fig. 25A) and of the CalA and CalB-resistant Δ*Rv2113* mutant (Supplementary Fig. 25B) were challenged with a 10-fold MIC_90_ of CalB for 48 h. Subsequently, whole protein cell lysates were prepared to perform global proteome analysis and comparison of the elicited stress responses. Overall, as expected, treatment of wild type cells with CalB led to more profound stress responses than treatment of the resistant Δ*Rv2113* mutant, which are unambiguously linked to the anti-tubercular nature of CalB, as the inactive compound **36** does not show a similar response at the proteome level (Supplementary Fig. 26). In wild type, the by far most predominant response was downregulation of diverse proteins. A GO analysis revealed that the affected processes were mainly cellular lipid metabolic processes and, to a lesser number, processes associated with DNA repair and replication. As in the comparison of the *ΔRv2113* and the complemented strain *ΔRv2113* pMV361::*Rv2113*, Mas was also found slightly downregulated (Fig. 7C), while PrpD, this time together with PrpC, was found to be one of the rare upregulated proteins (Fig. 7D). Among the downregulated lipid biosynthesis-related proteins were for example Fas (Rv2524c), AccD6 (Rv2247), FadD22 (Rv2948c), FbpC (Rv0129c), FabD (Rv2243), Pks15 (Rv2947c), FadD29 (Rv2950c) and Rv2949c. Rv2525c was previously found to be upregulated in cells of *M. tuberculosis* that were exposed to agents (isoniazid and ethionamide) primarily inhibiting mycolic acid biosynthesis [25]. This suggests that upregulation of Rv2525c might represent an adaptive response to compounds impairing lipid biosynthesis, including callyaerins. Recently, accumulation of methylcitrate cycle intermediates has been linked to tolerance towards first- and second-line TB drugs such as isoniazid and bedaquiline [26]. Therefore, upregulation of PrpC and PrpD might represent a specific stress response aiming at increasing tolerance to CalB. Furthermore, the virulence-associated methyltransferase Rv1405c, among others, has been reported to be responsible for acclimatizing *M. tuberculosis* CDC1551 to acid stress in macrophages [27], while the sigma factor SigB was reported to control a regulon involved in general stress resistance in *M. tuberculosis* [28]. This suggests that upregulation of Rv1405c and SigB may represent additional general stress adaptions in CalB-treated cells.

In summary, CalB treatment elicited a complex stress profile characterized by downregulation of lipid biosynthesis, DNA repair, replication and cell division. On the other hand, the cells responded by induction of several known stress adaptions, including Rv2525c, Rv1405c, PrpC-PrpD, and SigB. This complex stress profile is unique and differs from that of other natural antitubercular cyclic peptides such as teixobactin [29], evybactin [30], lassomycin [31], acyldepsipeptide [32] and cyclomarin [33], implying that callyaerins embark on a distinct mechanism of action that presumably involves simultaneous impairment of several pathways.

## DISCUSSION

We have elucidated structural determinants of CalA/B-type callyaerins, a family of potent antitubercular natural products, and demonstrated a direct targeting of the non-essential membrane protein Rv2113. As Rv2113 is *M. tuberculosis*-specific, this mode-of-action may allow the generation of bacteriostatics with less side effects to the microbiome as well as the spread of off-target pathogen antibacterial resistance. The callyaerins are therefore another promising example of the growing number of cyclo(depsi)peptide-based antitubercular natural products with unique mode-of-actions [34].

The binding of CalA/B-type callyaerins modulate the non-essential function of Rv2113 in a yet-elusive molecular way that elicits pleiotropic effects by simultaneous impairment of several intracellular pathways including downregulation of lipid biosynthesis, DNA repair, replication and cell division. This mode-of-action is thus distinct from other antibiotics that typically either target single critical steps in pathways that are essential for growth or viability of bacterial cells such as DNA replication, cell wall formation or protein, RNA or ATP biosynthesis, among others, or interact with membrane structures, resulting in pore formation and collapse of membrane potential. Our data however confirm that both resistance and susceptibility of *M. tuberculosis* toward CalA/B-type callyaerins are strictly linked to the non-essential membrane protein Rv2113. While a comparative full proteome analysis of the *ΔRv2113* gene deletion mutant and the complemented strain *ΔRv2113* pMV361::*Rv2113* suggests that Rv2113 may play a so far unknown role in lipid biosynthesis or transport, this function is only of accessory nature. Rv2113 is fully dispensable for normal in vitro growth of *M. tuberculosis* under the tested culture conditions and, thus, has not been considered a potential drug target so far. Non-essential proteins, which are connected to the mechanism of antibacterial compounds with phenotypes similar to Rv2113 (i.e., their deletion mediates resistance, while overexpression mediates hypersensitivity), have so far been reported to be associated with only three different processes: (1) Compound uptake, (2) activation of pro-drugs, or (3) deregulation and over-activation of proteases.

An uptake dependent role of non-essential membrane proteins is known, for example, for bacterial multi-solute transporters of the SbmA/BacA family that mediate incorporation of aminoglycoside antibiotics and antimicrobial peptides among other structurally diverse hydrophilic molecules [35]. Inactivation of these transporters confer resistance by preventing uptake of antibacterial compounds. However, our findings with a fluorescent Cy3-CalB derivative showed that CalA/B-type callyaerins enter *M. tuberculosis* cells independently of Rv2113, ruling out an uptake-related role of Rv2113.

Several antibiotics used to treat *M. tuberculosis* such as isoniazid, pyrazinamide and ethionamide are prodrugs that require intracellular activation by enzymes including catalase/peroxidase KatG [36], the pyrazinamidase/nicotinamidase PncA [37], or the Baeyer-Villiger monooxygenases EthA and MymA [38], respectively. Loss of these non-essential enzymes confer resistance by preventing pro-drug activation. However, we have no evidence that CalA/B-type callyaerins are pro-drugs that need direct activation by Rv2113. Rv2113 is a membrane protein comprising 387 amino acids, which harbours eight transmembrane helixes according to Alphafold prediction but does not contain any known catalytic domain.

Acyldepsipeptides (ADEPs) are a class of antibacterial natural compounds that kill Gram-positive bacteria by over-activation of the cytosolic ClpP protease [39]. Similarly, synthetic tripodal peptidyl compounds can inhibit bacterial growth by allosteric over-activation of the non-essential protease DegP [40]. Loss of these non-essential proteases confer resistance by preventing disturbed proteostasis that would otherwise occur by deregulated over-activation. However, a similar mechanism is highly unlikely for CalA/B-type callyaerins for several reasons. First, according to sequence analysis, Rv2113 does not contain any known proteolytic domain nor is it predicted to possess any other catalytic activity. Second, in contrast to other Gram-positive bacteria, the ClpP protease is essential in mycobacteria, and ADEPs were reported to impair growth of mycobacteria by inhibition of the essential ClpP function, not by over-activation [32]. Third, the proteomic stress profile elicited by CalB treatment is distinct and differs from that of cyclic peptidic compounds that cause disturbance of proteostasis in *M. tuberculosis* by either inhibiting or deregulating ClpP function, such as cylcomarins or homoBacPROTACs, respectively [33].

Finally, we also found no evidence that CalA/B-type callyaerins impair structure or function of the mycobacterial membrane or cell wall that could potentially occur if binding of callyaerins to Rv2113 would lead to pore formation.

In summary, we can rule out all known mechanisms of how a non-essential protein could mediate resistance and susceptibility towards an antibacterial compound similar to Rv2113, with deletion conferring resistance and overexpression mediating hypersensitivity. Thus, we conclude that the antitubercular activity of CalA/B-type callyaerins very likely relies on an unprecedented mechanism. This mechanism depends on presence of Rv2113 but also requires additional yet-unknown factors that are unique for *M. tuberculosis* and *M. bovis* BCG, but are obviously either lacking or at least differ substantially in other mycobacteria. One could speculate that this novel mechanism might rely on the unspecific recruitment of essential proteins to Rv2113 following binding of CalA/B-type callyaerins, thereby impairing their function. However, further investigations are necessary to prove or disprove this and other hypotheses concerning the molecular processes that occur following binding of these natural products to Rv2113 in *M. tuberculosis*.

Altogether, all our findings indicate that Rv2113 is a highly interesting protein – it is not only specific for *M. tuberculosis*, its structure is also unique, preventing any function predictions. Our investigation on the role of Rv2113 in the antitubercular mechanism of the CalA/B-type callyaerins strongly suggest that compound binding modulates function of this non-essential protein in a way that derails several metabolic pathways. This mechanism is unprecedented, diametrically differing from typical antibiotics that work by inhibiting essential targets. Further investigations on the functional role of Rv2113 in *M. tuberculosis* will be pursued in the future to shed light on the underlying molecular details of this unique mechanism. Finally, further medicinal chemical optimization of CalA/B-type callyaerins and development of callyaerin conjugates might pave the way for development of urgently needed *M. tuberculosis*-specific anti-TB drugs.

## Supporting information

Callyaerin Supplementary Infos

## ACKNOWLEDGEMENTS

We thank the Deutsche Forschungsgemeinschaft (DFG, German Research Foundation) (KA 2894/7-1 to M.K., KA 2259/5-1 to R.K.) for funding. This work was further partially supported by the DFG – project number 270650915 / GRK 2158 (to RK).

## MATERIALS AND METHODS

### Bacterial strains and culture conditions

*M. tuberculosis* strains including H37Rv, mc²6230 (Δ*panCD* ΔRD1) and several XDR-TB clinical isolates originating from South Africa, *M. bovis* BCG Pasteur as well as *M. smegmatis* mc²155 used in this study were obtained from the laboratory of William R. Jacobs Jr. (Albert Einstein College of Medicine, Bronx, NY). Cells of mycobacteria were grown at 37 °C and 80 rpm in liquid Middlebrook 7H9 medium supplemented with 10% ADS enrichment (5% bovine serum albumin fraction, 2% glucose and 0.85% sodium chloride), 0.5% glycerol and 0.05% tyloxapol, or solid Middlebrook 7H10 agar supplemented with 10% ADS enrichment and 0.5% glycerol. Selective media were supplemented as required with hygromycin (50 µg/mL), apramycin (30 µg/mL) or kanamycin (20 µg/mL). Cultures of non-mycobacterial species such as *Staphylococcus aureus*, *Escherichia coli*, *Bacillus subtilis*, *Enterococcus faecalis*, *E. faecium*, *Pseudomonas aeruginosa* and *Acinetobacter baumannii* were grown at 37 °C and 180 rpm in lysogeny broth (LB) or on solid LB medium.

### Determination of minimal inhibitory concentration (MIC)

The MIC was determined for all bacteria using broth microdilution method. For all mycobacteria species, cultures obtained from exponentially growing cells cultured in supplemented 7H9 medium were diluted and seeded at 1 × 10^5^ CFU/well in 96-well round bottom microplates, in a total volume of 100 µL of supplemented 7H9 medium containing two-fold serially diluted test compounds with a starting concentration of 100 µM. The plates were incubated at 37 °C for 5 days (*M. tuberculosis* strains and *M. bovis* BCG Pasteur) or 24 h (*M. smegmatis*). 10 µL of a 100 µg/mL resazurin solution was subsequently added to each well and further incubated for 16-24 h. To fix the cells for MIC determination, 100 µL of 10% formalin was added to each well (5% final concentration) for at least 30 min. Using a microplate reader (excitation, 540 nm; emission, 590 nm), fluorescence was quantified. Percentage of growth was calculated relative to sterile medium (0% growth) and DMSO solvent control (100% growth).

For nosocomial pathogens and *B. subtilis*, cultures obtained from exponentially growing bacteria in Mueller Hinton broth (MHB) were diluted and seeded at 5 × 10^4^ CFU/well in 96-well round bottom microplates, in a total volume of 100 µL MHB containing two-fold serially diluted test compounds with a starting concentration of 100 µM. Microplates were incubated for 24 h at 37 °C. MIC was determined macroscopically by identifying the lowest concentration that resulted in inhibition of visible bacterial growth.

### Cytotoxicity determination

The cytotoxicity of callyaerins was determined using the human THP-1 monocytic, MRC-5 lung fibroblast, HepG2 liver and HEK293 kidney cell lines. THP-1 monocytes and MRC-5 lung fibroblasts were maintained in Roswell Park Memorial Institute (RPMI) 1640 medium and Dulbeccós modified Eagle medium (DMEM), respectively, both media supplemented with 10% fetal bovine serum (FBS). HEK293 kidney cells were grown in Eagle’s minimum essential medium (EMEM) supplemented with 2 mM L-glutamine, 1 mM sodium pyruvate, 1% non-essential amino acids and 10% FBS, while HepG2 cells were cultivated in Ham’s F12 medium supplemented with 2 mM L-glutamine and 10% FBS. All cells were cultured at 37 °C in a humidified atmosphere of 5% CO_2_. Cells were diluted and seeded at 5 × 10^4^ cells/well in tissue culture-treated 96-well flat bottom microplates, in a total volume of 100 µL of respective media containing two-fold serially diluted test compounds with a starting concentration of 100 µM. 10 µL of a 100 µg/mL resazurin dye dilution was added to each well following 48 h incubation of microplates at 37 °C and 5% CO_2_. After 3 h incubation with resazurin, fluorescence was quantified in a microplate reader (excitation, 540 nm; emission, 590 nm). Percentage of growth was calculated relative to sterile medium (0% growth) and DMSO solvent control (100% growth). Selectivity index of test-compounds was determined as the quotient of the cytotoxic concentration inhibiting 50% growth of human cells and the MIC resulting in 90% inhibition of mycobacterial growth (IC_50_/MIC_90_).

### Mycobacterial killing kinetic

Killing kinetic studies were done by quantifying viable cell counts following different treatment intervals to assess bactericidal or bacteriostatic activity of compounds. For this, a mid-exponentially growing culture of *M. tuberculosis* mc²6230 was adjusted to 2 × 10^6^ CFU/mL and challenged with 5-fold MIC of CalB alone or in combination with clinical anti-TB drugs (isoniazid, rifampicin and bedaquiline) and incubated for 5 weeks at 37 °C and 80 rpm. At various time intervals, 200 µL aliquots were taken and 10-fold serially diluted. Serial dilutions were plated on Middlebrook 7H10 agar plates and incubated at 37 °C for 3 weeks to count colony forming units (CFU).

### Checkerboard assay

The interaction between CalB and clinical anti-TB drugs were quantified by determining the fractional inhibitory concentration indices (FICI) of CalB with these drugs in 96-well round bottom microplates. Compounds tested in combination were two-fold serially diluted in a two-dimensional fashion. Cultures of *M. tuberculosis* H37Rv and mc²6230 were diluted and added to the microplates at a density of 1 × 10^5^ CFU/well in a total volume of 100 µL. After 5-day incubation at 37 °C, 10 µL of a100 µg/mL resazurin solution was added to each well and further incubated for 16-24 h. Mycobacterial growth was quantified as a function of fluorescence intensity measured by a microplate reader (excitation, 540 nm; emission, 590 nm). Subsequently, the FICI was calculated as the sum of the quotients of the MICs of each bioactive agent when tested in combination (A and B) and the MIC when tested alone:

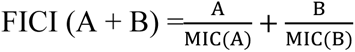

Synergism or antagonism between CalB and clinical anti-TB drugs were represented as FICI ≤ 0.5 or > 2, respectively. Partial synergism was denoted by 0.5 < FICI ≤ 0.75, while 0.75 < FICI ≤ 2 indicates an additive effect [23].

### Intracellular ATP measurement

Exponentially growing culture of *M. tuberculosis* mc²6230 was adjusted to OD (optical density) 0.5 at 600 nm and challenged with 5- or 10-fold MIC of CalB and relevant controls at 37 °C and 80 rpm. At various time intervals, 1 mL culture aliquots were taken, pelleted by centrifugation at 10,000 *g* for 5 min, and supernatants were discarded to remove extracellular ATP. Cell pellets were resuspended in PBS and dispensed in a white opaque 96-well microplate containing BacTiter-Glo reagent (Promega, Madison, WI, USA). The resulting 1:1 mixture was incubated for 5 min and luminescence was measured as a function of ATP content using a microplate reader.

### Propidium iodide internalization assay

A culture of *M. tuberculosis* mc²6230 taken from exponential growth phase was adjusted to OD_600 nm_ = 0.5, harvested by centrifugation at 4,500 *g*, 4 °С for 10 min, washed and resuspended in an equal volume of PBS supplemented with 1% glucose. Cells were then incubated with 15 µg/mL propidium iodide (PI). Cells were dispensed into a black opaque 96-well microplate upon incubation of cells with PI at 37 °C for 30 min to establish a baseline PI fluorescence measurement (excitation, 535 nm; emission, 617 nm) using a microplate reader. Cells were then challenged with 5-fold MIC (final concentration) of CalB or appropriate controls and immediately returned to the microplate reader to quantify uptake of PI by measuring red fluorescence in a kinetic fashion for at least 4 h.

### RNA extraction and real-time quantitative PCR

Cells of an exponentially growing culture of *M. tuberculosis* were adjusted to OD_600 nm_ = 1.5, incubated with 5-fold MIC of compounds and harvested after 24 h by centrifugation at 4,500 *g*, 4 °С for 10 min for RNA extraction. Harvested cells were fixed overnight in 5 mL RNA Protect reagent (Qiagen) after incubation with compounds. Fixed cells were pelleted, resuspended in 1 mL RLT buffer, and then transferred into Precellys® tubes containing a mixture of 0.5 mm and 0.1 mm silica beads. Cells were lysed by bead beating at 50 Hz, and RNA was extracted using the RNeasy Mini kit (Qiagen). Residual genomic DNA was removed by DNase treatment. After RNA integrity was checked using RNA 6000 Nano Chip, 2 µg RNA per sample was reverse transcribed into complementary DNA using the SuperScript™ III First-Strand Synthesis kit (Thermo Fisher). Gene expression was quantified by performing quantitative real-time PCR (qPCR) in a Mx3005P qPCR system (Agilent Technologies) using GoTaq® qPCR Master Mix (Promega). Analyses were performed following the manufactureŕs protocol using gene-specific qPCR primers. Expression values were determined by the ΔΔC_T_ method, normalized to *16SrRNA* using DMSO-treated cells as the control. All qPCR primers (Supplementary Table 1) were designed using the PrimerQuest tool.

### Generation of a targeted *M. tuberculosis* Δ*Rv2113* gene deletion mutant

Site-specific gene deletion in *M. tuberculosis* H37Rv was achieved by specialized phage transduction as described elsewhere [41]. Briefly, the *Rv2113* gene was replaced with a γδ-*sacB*-*hyg*-γδ cassette comprising a hygromycin resistance gene and a *sacB* counterselectable marker flanked by *res*-sites of the γδ-resolvase. For construction of the required allelic exchange substrate, upstream and downstream flanking regions of *Rv2113* were amplified by PCR using primers listed in Supplementary Table 2. Subsequently, the flanking regions were digested with the indicated restriction enzymes and ligated with the *Van91*I-digested p0004S vector. The resulting allelic exchange plasmid was then linearized with *Pac*I, cloned, and packaged into the temperature-sensitive phage ФphAE159, yielding knock-out phages that were propagated in *M. smegmatis* at 30 °C. Allelic exchange in *M. tuberculosis* was achieved by specialized transduction at the non-permissive temperature of 37 °C, using hygromycin for selection, resulting in gene deletion and replacement by γδres-*sacB*-*hyg*-γδres cassette. Obtained hygromycin-resistant transductants were obtained after 3 weeks of incubation and were screened for correct gene disruption by diagnostic PCR.

### Heterologous gene expression in mycobacteria

For genetic complementation and overexpression purposes, the genes *Rv2113* and *BCG_2130* were amplified from genomic DNA of *M. tuberculosis* H37Rv or *M. bovis* BCG Pasteur, respectively, by PCR using the primer pairs listed in Supplementary Table 3, and cloned into the integrative single-copy plasmid pMV361 containing an apramycin resistance marker providing constitutive heterologous gene expression from the HSP60 promoter. The plasmids were transformed into mycobacterial cells via electroporation (1000 Ω, 25 µF, 2500 V). Transformants were selected on Middlebrook 7H10 agar plates containing apramycin (40 mg/l).

### Spontaneous resistant mutant generation

Spontaneous mutants of *M. tuberculosis* H37Rv resistant to callyaerins were generated by plating 500 µL culture aliquots containing ∼ 3 × 10^7^ CFU on 120 mm square agar plates supplemented with 5-fold MIC of CalA or CalB, respectively °C. After 3-4 weeks of incubation at 37 °C, independent colonies were picked for further analyses. Spontaneous mutants raised in the *Rv2113* merodiploid strain of *M. tuberculosis* H37Rv were generated on agar additionally containing 30 µg/mL apramycin to select for presence of the integrative plasmid carrying the merodiploid copy of *Rv2113*.

### Whole-genome sequencing

Genomic DNA of independent resistant mutants was extracted using the CTAB-lysozyme method as described elsewhere [42]. Genomes of resistant mutants were sequenced with an Illumina HiSeq 2500 next generation sequencer (at Texas A&M University, College Station, TX, USA & Biological and Medical Research Center, Heinrich Heine University Düsseldorf, Germany) after preparing sequencing libraries using standard paired-end genomic DNA sample prep kit from Illumina. Genome sequences were compared with that of the parent *M. tuberculosis* H37RvMA (GenBank accession GCA_000751615.1). Paired-end sequence data was collected with a read length of 106 bp. Base-calling was performed with Casava software, v1.8. The reads were assembled using a comparative genome assembly method, with *M. tuberculosis* H37RvMA as a reference sequence [43].

### Intracellular activity in infected macrophages

For differentiation into macrophage-like cells, human THP-1 cells were seeded at a density of 1 × 10^5^ cells per well in a total volume of 100 µL RPMI 1640 medium supplemented with 10% FBS and 50 nM phorbol-12-myristate-13-acetate (PMA) in a 96-well flat-bottom microtiter plate. Cells were incubated overnight in a humidified atmosphere at 37 °C and 5% CO_2_. The next day, cells were washed with PBS twice, and medium was replaced with fresh RPMI 1640 medium supplemented with 10% FBS containing 3 × 10^5^ cells per well of a precultured *M. tuberculosis* H37Rv reporter strain constitutively expressing mCherry resulting in a multiplicity of infection of three (MOI = 3). After three hours of infection, cells were washed with PBS twice, and medium was replaced with fresh RPMI 1640 medium supplemented with 10% FBS containing either 15.6 µM of CalA or 1.95 µM of CalB, respectively. DMSO and standard antibiotics (1 µM rifampicin or 20 µM streptomycin) were used as negative and positive controls, respectively. After five days of cultivation at 37 °C and 5% CO_2_, cells were fixed with a final concentration of 5% formalin and incubated for 30 min at room temperature. Fluorescence was quantified using a Nikon Eclipse TS100 and NIS-Elements (100 x magnification, 500 ms exposure time). Integrated density of red fluorescence was calculated using Fiji (ImageJ).

### Confocal microscopy

Cells of an exponentially growing culture of *M. tuberculosis* H37Rv were adjusted to OD_600 nm_ = 0.4 in Middlebrook 7H9 medium supplemented with 10% ADS enrichment, 0.5% glycerol and 0.05% tyloxapol and were treated with 0.2 µM of either **87** or Cy3 for 0.5, 3 or 6 h. Subsequently, cells were harvested by centrifugation at 5,000 *g* for 5 min and washed once with PBS containing 0.05% tyloxapol. Next, cells were stained with 4′,6-diamidino- 2-phenylindole (DAPI) for 30 min at 4 °C. Upon washing stained cells three times with PBS containing 0.05% tyloxapol, cells were fixed with 10% formaldehyde for 1 h at RT before removal from BSL-3 confinement. Fixed samples were then washed once, resuspended in PBS containing 0.05% tyloxapol and centrifuged at 800 rpm for 10 min to remove cell aggregates. Samples were imaged using Zeiss LSM710 Confocal Laser Scanning Microscope after spreading single-cell suspensions on soft agarose pad. Captured images were subsequently analyzed using ImageJ.

### Photoaffinity enrichment-LC-MS/MS-based target identification

Cells of an exponentially growing culture of *M. tuberculosis* H37Rv were harvested by centrifugation at 4000 *g*, 4 °C for 10 min and washed thrice with PBS supplemented with 0.05% tyloxapol. The cells were then aliquoted into 2-ml centrifugation tubes in a total volume of 1 mL PBS containing 0.05% tyloxapol. For some experiments, samples were then pre-treated with 30 µM CalB as a competitor for 30 minutes, whereas solvent controls contained 0.3% DMSO in total. Next, cells were treated with the respective probes by adding 250 µL of a 5x stock (in PBS; 0.63% DMSO). The samples were incubated for 180 min shaking at 37 °C and afterwards transferred to 6-well plates and exposed to irradiation with UV light (365 nm) for 20 minutes. The cells were transferred back to 2-mL centrifugation tubes, pelleted by centrifugation for 5 min at room temperature and 4500 *g* and washed three times with 1.5 mL of 50 mM Na_2_HPO_4_. Pellets were resuspended in 1,050 µL of 50 mM Na_2_HPO_4_ and cells were lysed by bead-beating using 100 µm silica zirconium beads (50 Hz; 15 min). Subsequently, 350 µL of 4% sodium dodecyl sulfate (SDS; Carl Roth, 2326.2)) in 50 mM Na_2_HPO_4_ were added and samples were incubated at 95 °C for 5 min. After cooling down the samples for 15 min under continuous shaking, lysates were cleared by centrifugation at 13,000 *g* at room temperature for 5 min. Supernatants were collected to be used as protein solutions and were filter-sterilized twice through a bacteria-tight 0.2 µm cellulose acetate syringe filter to remove viable bacteria.

For the use of a two-step enrichment protocol for labeled proteins, a volume equivalent to 250 µg protein (Quantification: 660 nm Protein Assay with ionic detergent compatibility reagent; Thermo Scientific, 22660 & 22663) was transferred to 15 mL centrifugation tubes and filled up with 1% SDS in 50 mM Na_2_HPO_4_ to a total volume of 920 µL. SDS concentration was reduced by addition of 920 µL benzonase-containing buffer (150 u per sample; EMD Millipore, 71206) with subsequent incubation for benzonase treatment at 37 °C for 30 min. In the following, Copper(I)-catalyzed azide-alkyne cycloaddition (CuAAC) was performed by transferring 120 µL reagent-mixture containing 167 µM Biotin-Azide (Jena Bioscience, CLK-AZ104P4), 1.67 mM TBTA (Sigma-Aldrich, 678937) and 33.3 mM TCEP (Sigma-Aldrich, C4706) to the samples and the click reaction was started by addition of 40 µl 100 mM CuSO_4_. The mixture was incubated under continous rotation for 150 min at room temperature. The proteins were precipitated by addition of four times excess of ice-cold methanol (−20 °C, overnight). After sedimentation of the proteins via centrifugation (4 °C, 3220 g, 20 min), the supernatant was removed and the pellet washed with 1 mL ice-cold methanol. After an additional centrifugation step, protein pellets were taken up in 850 µL of 2% SDS in PBS and incubated at 37 °C for several minutes. The total volume was increased by further addition of PBS to a final volume of 3 mL and the remaining pellets were disrupted through sonication. Insoluble particles were sedimented by centrifugation (1400 g, 5 min), the supernatant was transferred to a fresh 15 centrifugation tube and SDS concentration was reduced to 0.2% by filling up the samples with PBS to a total volume of 8.5 mL. For enrichment, avidin agarose (Thermo Scientific, 20225) was equilibrated to 0.2% SDS in PBS and thereafter 100 µL of the slurry suspension was added to each sample (approx. 50 µL bead volume). The suspension was incubated under continuous rotation for 60 min. The beads were sedimented by centrifugation (5 min, 400 g), the supernatant was discarded and the beads were washed four times with 10 mL 1% SDS in H_2_O by using an iterative procedure consisting of incubation (5 min), centrifugation (3 min, 400 g) and supernatant-removal steps. After an additional washing step with 10 mL H_2_O, the beads were transferred to a 1.5 mL centrifugation tube and washed two more times with 1 mL H_2_O (centrifugation 1 min, 3000 g) to remove residual SDS. For on-bead digest, the beads were taken up in 100 µL 0.8 M urea (GE Healthcare, GE17-1319-01) in 50 mM ammonium bicarbonate (ABC; Sigma-Aldrich, 11213) containing 10 mM dithiothreitol (DTT; Sigma-Aldrich, 43815) and incubated for 60 min under continous shaking (1500 rpm). Subsequently, 20 mM iodoacetamide (IAM; Sigma-Aldrich, A3221) was added for protein alkylation (1500 rpm, 75 min). Excess IAM was quenched by increasing the DTT concentration to 25 mM (1500 rpm, 15 min) and the digest was started by the addition of 500 ng trypsin (Promega, V5111) per sample. The digest was incubated under continuous shaking (1250 rpm) for 18 h at 37 °C and stopped by the addition of 5% formic acid (FA; Fisher chemical, A117-50). After sedimentation of the beads (5 min, 600 g), the supernatant was transferred to a fresh centrifugation tube and the beads were washed with 50 µL 1% FA. The supernatants were combined and passed through equilibrated glass microfiber tips (pore size: 1.2 µm, thickness: 0.26 mm, two disks per tip; GE Healthcare, 1822-024) to remove any residual beads. The resulting peptide solution was desalted on home-made C18 StageTips (two discs per tip; 3M, 66883-U) as described before [44] and dry peptides were dissolved in 16 µL 0.1% FA (15 min, 1500 rpm). For analysis, 4 µL of sample was loaded on a self-packed fused silica capillary tube with evacuated-frit tip (75 µm ID x 50 cm, 15 µm orifice; CoAnn Technologies, ICT36007515F-50) filled with either Kinetex XB-C18 (particle size 1.7 µm, Pore Size 100 Å; Phenomenex, 4498) or Reprosil-Pur 120 C18-AQ (particle size 1.9 µm, Pore Size 120 Å; Dr. Maisch, r119.aq.) material. Peptides were separated using a 140 min gradient generated by an EASY-nLC 1200 liquid chromatography (Thermo Fisher Scientific) heated to 50 °C by a PRSO-V1 column oven (Sonation). In case of Kinetex material, a rising proportion of 80% acetonitrile (ACN) in H_2_O with 0.2% FA (solvent B1) in 2% ACN in H_2_O with 0.2% FA (solvent A1) was used for gradient preparation (3-7% B1 in A1 within the first 8 min, 7-27% in the next 84 min, 27-44% in the next 32 min, rise to 100% in the following 6 min, hold for 10 min) at a flow rate of 300 nL/min. In case of Reprosil-Pur material, a rising proportion of 80% acetonitrile (ACN) in H_2_O with 0.1% FA (solvent B2) in H_2_O with 0.1% FA (solvent A2) was used to prepare the gradient (6-32% B2 in A2 within the first 95 min, 32-42% in the next 20 min, rise to 100% in the following 10 min, hold for 15 min) at a flow rate of 250 nL/min. Peptides were ionized using a Nanospray Flex ion source (Thermo Fisher Scientific) with 2300-2500 V spray voltage and MS acquisition was performed in an Orbitrap Fusion Lumos mass spectrometer (Thermo Fisher Scientific). MS1 data acquisition was done in a *m/z* range of 375 to 1800 at a 120000-240000 orbitrap resolution with a maximal injection time of 50 ns. Data dependent MS2 spectra were recorded using a loop cycle of 3 seconds with a dynamic exclusion duration of 20-25 seconds. For precursor isolation a 1.2 *m/z* quadrupole isolation window was used with subsequent stepped HCD fragmentation (20, 30, 40%) and data acquisition at rapid ion trap scan rate with 300% of normalized AGC target. Data processing was done with Proteome Discoverer 2.5 (Thermo Fisher Scientific) using SequestHT protein search and subsequent statistical analysis was done using Perseus 2.0.7.0. [45]

### LC-MS/MS-based whole proteome comparison

Cells of exponentially growing cultures of *M. tuberculosis* H37Rv were adjusted to OD_600 nm_ = 0.75 in a total volume of 20 mL Middlebrook 7H9 medium supplemented with 10% ADS enrichment and 0.05% tyloxapol. Optionally, cells were subjected to treatment with compound or a corresponding volume of DMSO as solvent control for 48 h at 37 °C with shaking at 80 rpm. Subsequently, cells were harvested by centrifugation at 4,500 *g*, 4 °С for 10 min and washed thrice with PBS. Pellets were resuspended in 750 µL of PBS and cells were lysed by bead-beating using 100 µm silica zirconium beads (50 Hz; 15 min). After addition of 250 µL 4× lysis buffer (4% SDS, 40 mM tris(2-carboxyethyl)phosphine (TCEP; Sigma-Aldrich, C4706), 160 mM chloroacetamide (CAM; Sigma-Aldrich, C0267), 200 mM HEPES), samples were incubated for 10 min at 95 °C. Lysates were cleared by centrifugation at 13,000 *g* at 4 °С for 5 min. Supernatants were collected to be used as protein solutions and were filter-sterilized twice through a bacteria-tight 0.2 µm cellulose acetate syringe filter to remove viable bacteria. For the digestion of proteins, a modified SP3 protocol was used. [46] For this purpose, a total volume equivalent to 15 µg protein was transferred from each sample to a fresh vessel (quantification: 660 nm protein assay with ionic detergent compatibility reagent; Thermo Scientific, 22660 & 22663) and the volume was equalized between samples by addition of 1× lysis buffer (1% SDS, 10 mM TCEP, 40 mM CAM, 50 mM HEPES). Benzonase-treatment was started by the addition of 7 u benzonase (37 °C, 1200 rpm, 15-20 min) and subsequent cysteine alkylation was done using 5-10 mM IAM (37 °C, 1200 rpm, 30-60 min). Hydrophilic (Cytiva, #45152105050250) and hydrophobic Sera-Mag SpeedBeads (Cytiva, #65152105050250) were mixed equally and afterwards equilibrated to water. To each sample 150 µg of particles were added and protein binding was induced by adding an equal volume of ethanol to the samples (1250 rpm, 15-30 min). The particles were settled on a magnetic rack and further processed as already described [33]. For analysis, a volume equivalent to 500 ng of peptides was loaded on either a self-packed pulled tip glass capillary column (75 µm ID x 50 cm, 5 µm orifice; ESI Source Solutions, PTC3-75-50-SP; used for comparison of genetic mutants) or a self-packed fused silica capillary tube with evacuated-frit tip (75 µm ID x 50 cm, 15 µm orifice; CoAnn Technologies, ICT36007515F-50; used for stress response analysis) filled Reprosil-Pur 120 C18-AQ (particle size 1.9 µm, Pore Size 120 Å; Dr. Maisch, r119.aq.) material. Peptides were separated using a 140 min gradient generated by an EASY-nLC 1200 liquid chromatography (Thermo Fisher Scientific) heated to 50 °C by a PRSO-V1 column oven (Sonation). For gradient generation, a rising proportion of 80% acetonitrile (ACN) in H_2_O with 0.1% FA (solvent B) in H_2_O with 0.1% FA (solvent A) was used (genetic mutants comparison: 8-35% B in A within the first 100 min, 35-44% in the next 20 min, rise to 100% in the following 10 min, hold for 10 min; stress response analysis: 8-33% B in A within the first 95 min, 33-42% in the next 20 min, rise to 100% in the following 10 min, hold for 15 min) at a flow rate of 300 or 250 nL/min, respectively. Peptides were ionized using a Nanospray Flex ion source (Thermo Fisher Scientific) with 2300 V spray voltage and MS acquisition was performed in an Orbitrap Fusion Lumos mass spectrometer (Thermo Fisher Scientific). MS1 data acquisition was done in a *m/z* range of 375 to 1750 at a 120000-240000 orbitrap resolution with a maximal injection time of 50 ns. Data dependent MS2 spectra were recorded using a loop cycle of 3 seconds with a dynamic exclusion duration of 25-30 seconds. For precursor isolation a 1.2 *m/z* quadrupole isolation window was used with subsequent stepped HCD fragmentation (20, 30, 40%) and data acquisition at rapid ion trap scan rate with 300% of normalized AGC target. Data processing was done with MaxQuant 2.3.0.1 [47] using Andromeda protein search and subsequent statistical analysis was done using Perseus 2.0.7.0. [45]

## Notes

### Competing Interest Statement

The authors have declared no competing interest.

