## Supplementary material for "The anti-tubercular callyaerins target the *Mycobacterium tuberculosis*-specific non-essential membrane protein Rv2113": Callyaerin Supplementary Infos

#### Supplementary Figures

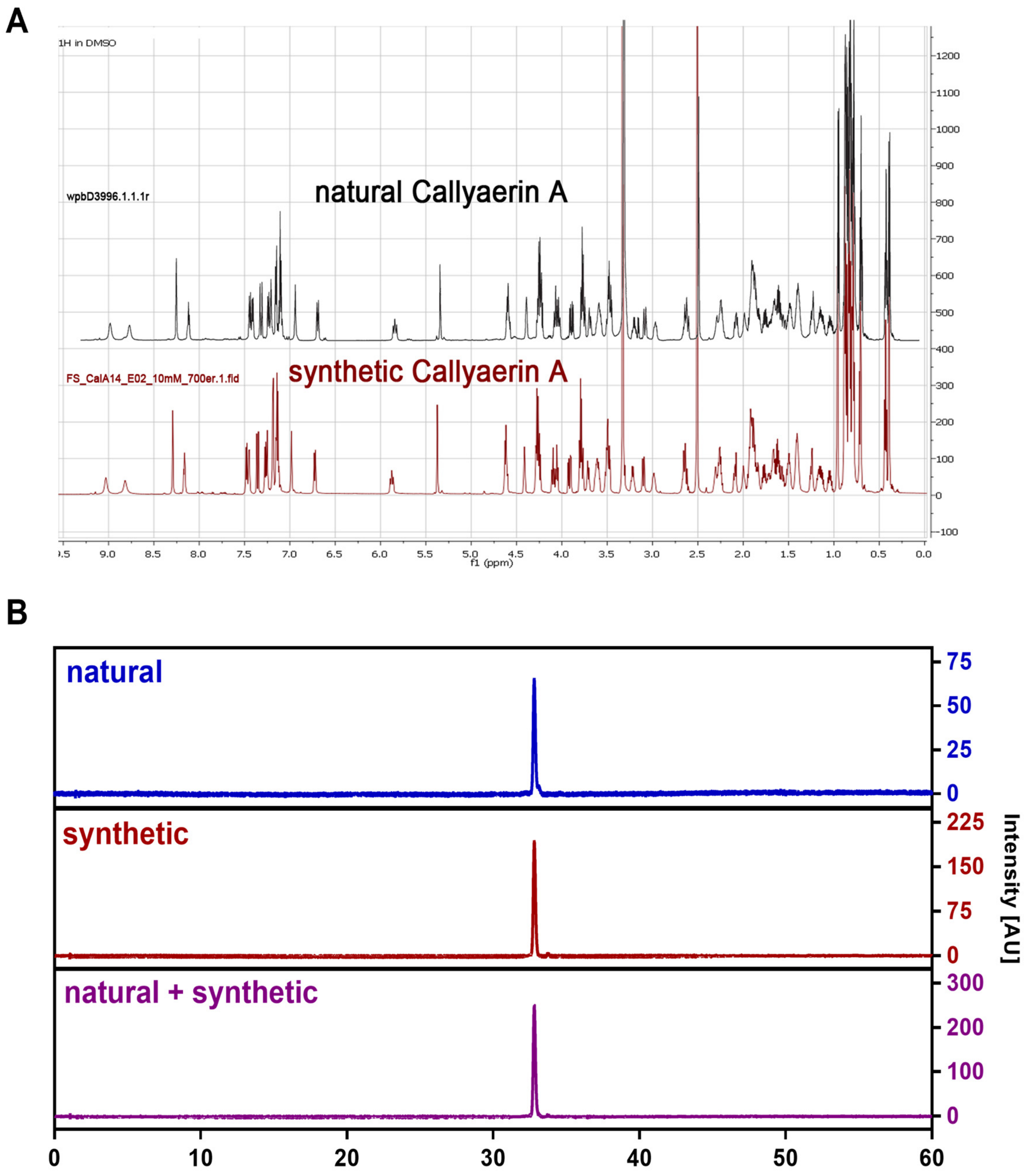

**Supplementary Figure 1.** Superposition of $^{1}$H NMR data (**A**) and LC-MS co-elution (**B**) of the isolated natural Callyaerin A (CalA) and our synthetic variant show structural similarity, thereby confirming the established synthetic approach.

| Activity spectrum of CalA | $MIC_{90}$ [µM] |
|---|---|
| **Strain** | **CalA** |
| *M. tuberculosis* mc[2] 6230[a] | 1.56 |
| *M. tuberculosis* H37Rv[a] | 3.13 |
| *M. tuberculosis* CDC 1551[a] | 3.13 |
| *M. tuberculosis* KZN 06[a] | 12.5 |
| *M. tuberculosis* KZN 13[a] | 12.5 |
| *M. tuberculosis* KZN 14[a] | 12.5 |
| *M. tuberculosis* KZN 16[a] | 6.25 |
| *M. tuberculosis* Beijing-HN878[a] | 25 |
| *M. bovis* AF2122/97[a] | > 50 |
| *M. bovis* BCG Pasteur[a] | > 50 |
| *M. smegmatis* mc[2] 155[a] | >100 |
| *M. marinum* ATCC 927[a] | > 50 |
| *S. aureus* ATCC 700699[b] | >100 |
| *H. sapiens* HEK293[c] | 50* |
| *H. sapiens* THP-1[d] | 50* |
| *H. sapiens* HepG2[e] | 50* |

[a]Middlebrook 7H9 [b]Mueller-Hinton [c]EMEM *$IC_{50}$ values
[d]RPMI 1640 [e]Ham`s F12

**Supplementary Figure 2.** $MIC_{90}$ values of CalA for a variety of fast- and slow-growing mycobacterial strains including clinical isolates and for the nosocomial pathogen *S. aureus* as well as $IC_{50}$ values of human cell lines determined by resazurin assay ($n$ = 2-3). Media used for cultivation of the respective cells are indicated by brown superscripted letters.

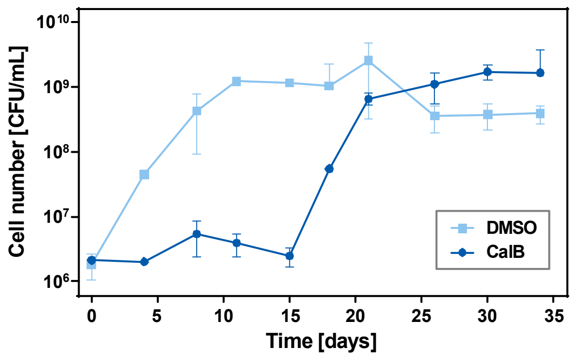

**Supplementary Figure 3.** CalB acts bacteriostatic. Overview on the killing kinetic of *M. tuberculosis* mc²6230 cells in presence of 15.7 µM CalB (5-fold $MIC_{90}$) over a period of 35 days. Viable cell number was determined by plating serially diluted culture aliquots on Middlebrook 7H10 agar plates to quantify colony forming units (CFU) after incubation at 37 °C for 3 weeks ($n = 2$, error bars indicate SD).

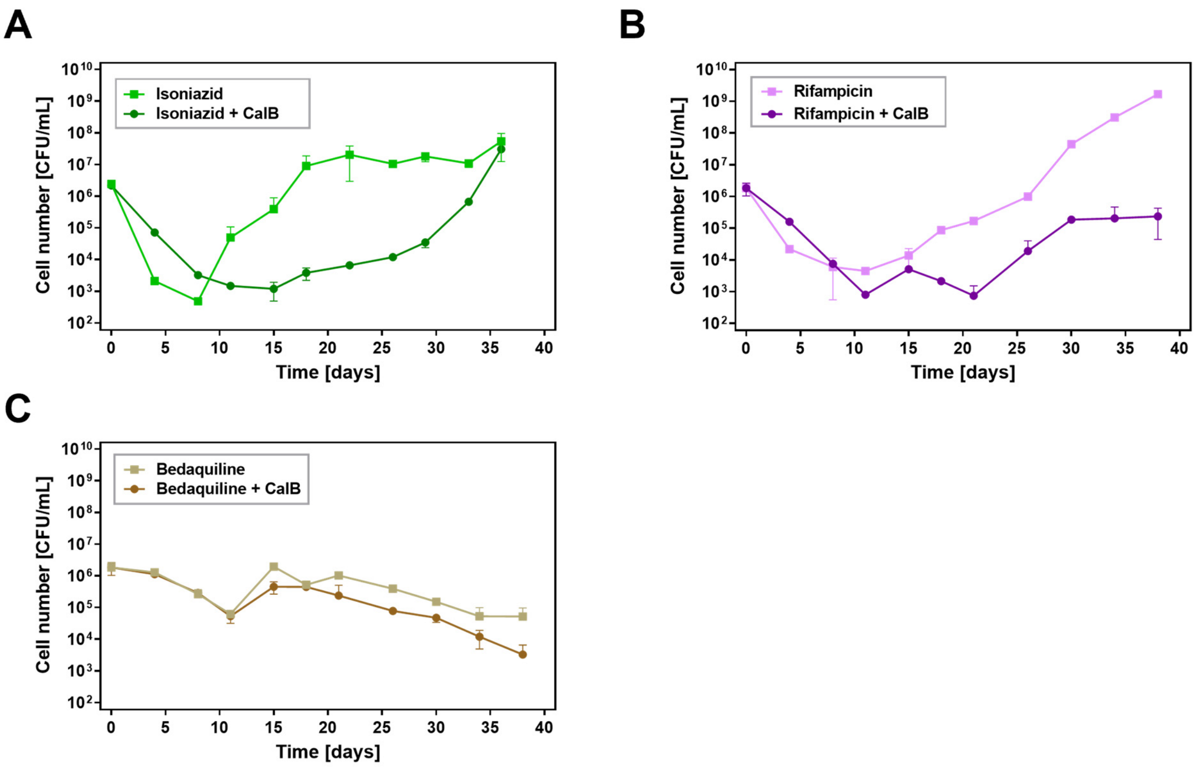

**Supplementary Figure 4.** CalB shows additive effects in combination with clinically used anti-tubercular drugs and slows the emergence of resistant mutants to these agents. Killing kinetic of *M. tuberculosis* mc$^2$6230 treated with **A**) isoniazid (1.95 µM, 5-fold $MIC_{90}$, green), **B**) rifampicin (0.5 µM, 5-fold $MIC_{90}$, purple) or **C**) bedaquiline (1.95 µM, 5-fold $MIC_{90}$, brown) in monotherapy (lighter color) or in combination with CalB (15.65 µM, darker color) over a time course of 36 to 38 days. Viable cell number was determined by plating serially diluted culture aliquots on Middlebrook 7H10 agar plates to quantify colony forming units (CFU) after incubation at 37 °C for 3 weeks ($n$ = 2, error bars indicate SD).

| anti-TB drug | MIC$_{90}$ [µM] Drug | Drug + CalB (A) | CalB + Drug (B) | FIC$_A$ | FIC$_B$ | FICI | Effect |
|---|---|---|---|---|---|---|---|
| **Combinatorial Effects** | | | | | | | ***M. tuberculosis* H37Rv** |
| CalB | 3.13 | | | | | | |
| BDQ | 0.39 | 0.200 | 3.13 | 0.5 | 1.0 | 1.5 | additive |
| RIF | 0.10 | 0.050 | 3.13 | 0.5 | 1.0 | 1.5 | additive |
| INH | 0.39 | 0.200 | 3.13 | 0.5 | 1.0 | 1.5 | additive |
| EMB | 3.13 | 1.560 | 3.13 | 0.5 | 1.0 | 1.5 | additive |
| DEL | 0.05 | 0.025 | 3.13 | 0.5 | 1.0 | 1.5 | additive |
| **Combinatorial Effects** | | | | | | | ***M. tuberculosis* mc$^2$ 6230** |
| CalB | 1.56 | | | | | | |
| BDQ | 0.39 | 0.200 | 0.78 | 0.5 | 0.5 | 1.0 | additive |
| RIF | 0.05 | 0.025 | 1.56 | 0.5 | 1.0 | 1.5 | additive |
| INH | 0.20 | 0.100 | 1.56 | 0.5 | 1.0 | 1.5 | additive |
| EMB | 1.56 | 0.780 | 1.56 | 0.5 | 1.0 | 1.5 | additive |
| DEL | 0.05 | 0.025 | 1.56 | 0.5 | 1.0 | 1.5 | additive |

**Supplementary Figure 5.** Determination of combinatorial effects of CalB with known anti-tubercular drugs (bedaquiline: BDQ, rifampicin: RIF, isoniazid: INH, ethambutol: EMB, delamanid: DEL). Fractional inhibitory concentrations were determined via checkerboard assay ($n$ = 3) through combination of a twofold serial dilution of an anti-tubercular drug with different concentration of CalB (A) or vice versa (B) in *M. tuberculosis* H37Rv or mc$^2$6230. All values are given in µM. Calculation of FIC: FIC = MIC$_{90}$ of drug/combinatorial MIC$_{90}$. Calculation of fractional inhibitory concentration indices (FICI) by addition of the respective FIC$_A$ and FIC$_B$. FICI was defined as follows: synergistic effect when FICI $\leq$ 0.5; partial synergistic effect when FICI > 0.5 $\leq$ 0.75; additive effect when FICI > 0.75 $\leq$ 2; antagonistic effect when FICI > 2.

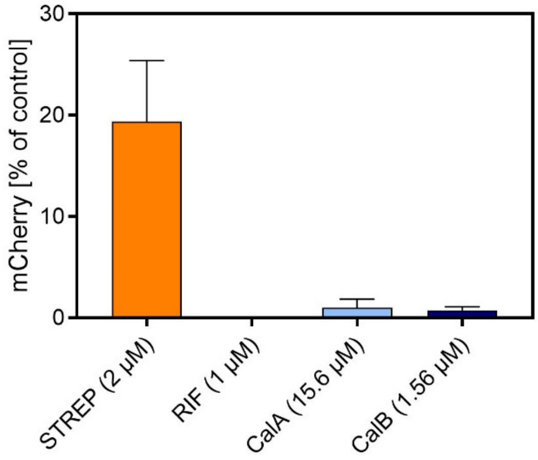

**Supplementary Figure 6.** Quantification of integrated fluorescence intensity from Fig. 2B. The integrated density of red fluorescence (mCherry) from the obtained photomicrographs (shown as mean of three fields per view ± SEM) was normalized to the signal of the DMSO-treated control (corresponding to 100% growth). STREP, streptomycin; RIF, rifampicin.

**A**

| CalA Derivatives | | | |
|---|---|---|---|
| # | Pos. | AS | $MIC_{90}$ [µM] |
| 63 | R2 | Pro | 3.13 |
| 64 | R3 | Abu | 12.5 |
| 65 | R3 | Chg | 1.56 |
| 66 | R3 | Cpg | 3.13 |
| 67 | R3 | Leu | 3.13 |
| 68 | R3 | Phe | 3.13 |
| 69 | R5 | Asp | >100 |
| 70 | R8 | Trp | >100 |
| 71 | R8 | Val | 3.13 |
| 72 | C3 | Leu | 3.13 |
| 73 | C3 | Ile | 1.56 |
| 74 | C3 | -* | 25 |
| 75 | C4 | Aha | 3.13 |
| 76 | C4 | -* | 1.56 |

* Deletion of AS at this and subsequent positions

**B**

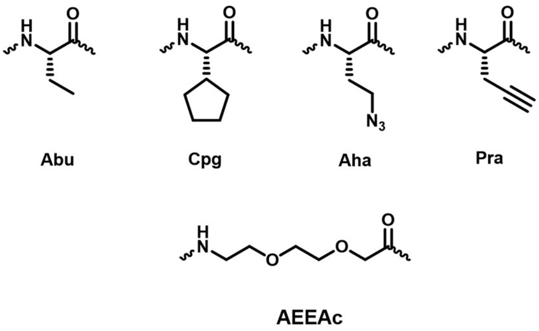

**C**

| Alkyne Probes CalB | | | | | | | | | | | $MIC_{90}$ [µM] | |
|---|---|---|---|---|---|---|---|---|---|---|---|---|
| R1 | R2 | R3 | R4 | R5 | R6 | R7 | R8 | C1 | C2 | C3 | C4 | C5 |
| | | 78 | 79 | | | 80 | | | | 81 | 82 | 83* |
| | | ND | ND | | | 25 | | | | 25 | 6.25 | 12.5 |

* Extention to C5 position was done via the amino acid AEEAc at position C4

**Supplementary Figure 7.** Structure-activity relationships for further CalA and CalB derivatives. **A**) Additionally synthesized CalA derivatives and their corresponding $MIC_{90}$ values against *M. tuberculosis* H37Rv. **B**) Chemical structures corresponding to the abbreviations used in A or B. **C**) Overview on the generated alkyne-modified CalB derivatives and their corresponding $MIC_{90}$ values in *M. tuberculosis* H37Rv. The alkyne modification was incorporated into the compound structure via the amino acid propargyl glycine (Pra).

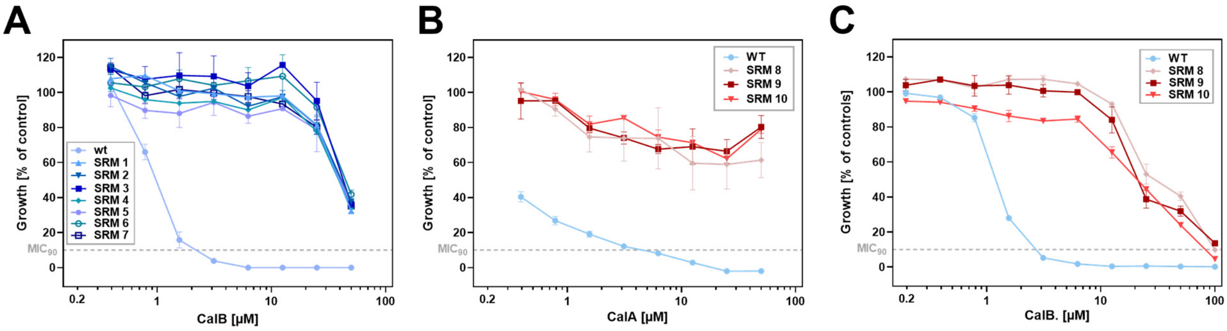

**Supplementary Figure 8.** Dose-response curves for isolated spontaneous resistant mutants (SRM) of *M. tuberculosis* H37Rv. **A**) Dose-response curves reporting CalB-dependent growth inhibition of SRM clones 1-7 isolated from CalB containing media plates. **B**) Dose-response curves reporting CalA-dependent growth inhibition of SRM clones 8-10 isolated from CalA containing media plates. **C**) Dose-response curves reporting CalB-dependent growth inhibition of SRM clones 8-10 isolated from CalA containing media plates. This experiment validates cross-resistance of isolated SRM. Growth of SRM together with that of *M. tuberculosis* H37Rv wild type (WT) was quantified relative to untreated DMSO control (corresponding to 100% growth) and sterile medium control (0% growth) using resazurin dye reduction assay. Data are shown as means of duplicates (**A** & **C**) or triplicates (**B**) with SD depicted as error bars.

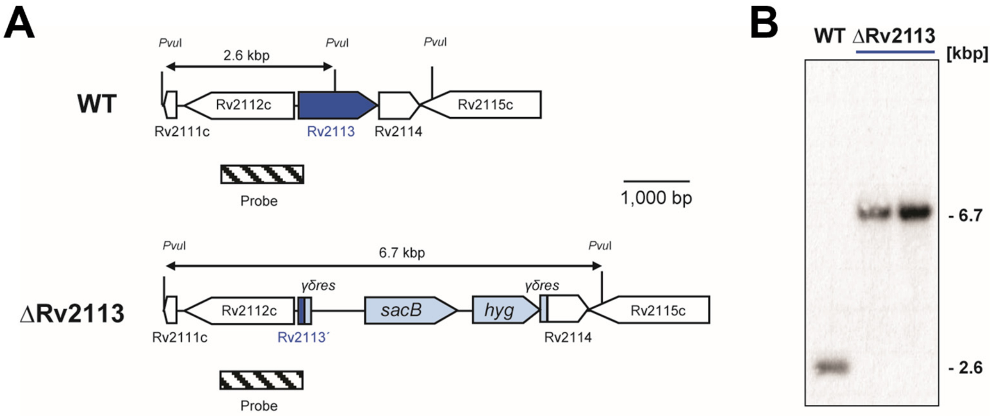

**Supplementary Figure 9.** Overview on the cloning strategy enabling the generation of the site-specific *ΔRv2113 M. tuberculosis* H37Rv gene deletion mutant. **A**) Organization of the *Rv2113* locus in *M. tuberculosis* wild type (WT) as well as in the Δ*Rv2113* gene deletion mutant. The sizes of relevant fragments as well as the location of the probe used for Southern analysis are indicated. γδ*res*: *res*-sites of the γδ-resolvase; *hyg*: hygromycin resistance gene; *sacB*: levansucrase gene from *Bacillus subtilis*. **B**) Southern analyses of *Pvu*I-digested genomic DNA from cells of *M. tuberculosis* wild type as well as the Δ*Rv2113* gene deletion mutant using a probe hybridizing to the position indicated in **A**, showing *Rv2113* gene deletion.

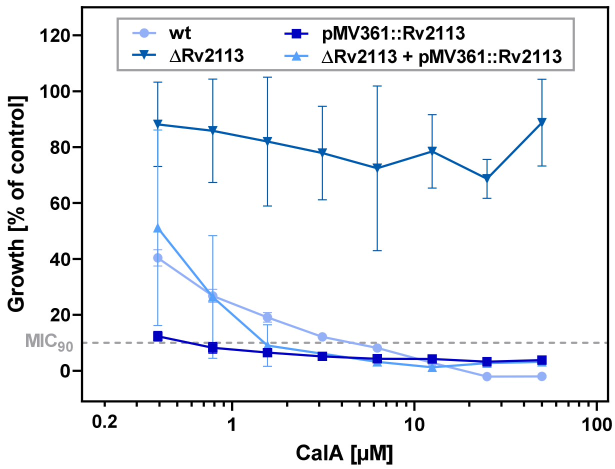

**Supplementary Figure 10**. Dose-response curves for CalA concentration-dependent growth inhibition of H37Rv strains. The assay included the wild type, the Δ*Rv2113* gene deletion mutant, the complemented mutant strain Δ*Rv2113* pMV361::*Rv2113* and the merodiploid strain overexpressing Rv2113 from plasmid pMV361::*Rv2113*. Data shown as means of triplicates with SD depicted as error bars.

**A**

| CalB resistant Rv2113 merodiploid Mutants | | |
|---|---|---|
| # | Rv2113 | Rv3058c |
| 11 | A217V (het) | W205* |
| 12 | L338P (het) | |
| 13 | L338P (het) | |
| 14 | L338P (het) | |
| 15 | L338P (het) | |
| 16 | A217V (het) | W205* |
| 17 | A217V (het) | W205* |
| 18 | A217V (het) | W205* |

**B**

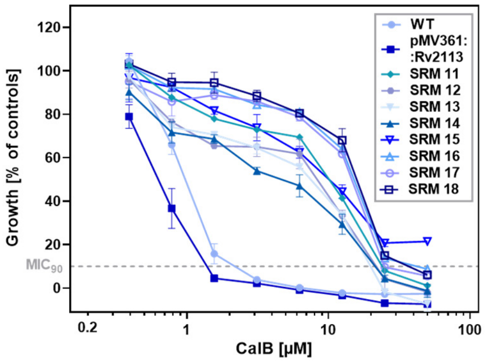

**Supplementary Figure 11.** Overview on the results from the spontaneous CalB-resistant mutant screening raised in the merodiploid strain *M. tuberculosis* H37Rv pMV361::*Rv2113*.harboring two copies of the *Rv2113* gene. **A**) The identified spontaneous resistant mutants (SRMs) emerged at a frequency of ~$10^{-6}$ on 7H10 agar containing 15.7 µM CalB (5-fold $MIC_{90}$) and 30 µg/mL apramycin after 4-week incubation. The corresponding mutations were mapped via whole-genome sequencing. Heterogeneous results (het) were obtained for *Rv2113*, meaning that the relevant SNP was found in ca. 50% of reads. This indicates that the mutation was only manifested either in the endogenous gene or in the merodiploid locus, which could not be distinguished. **B**) Dose-response curves of CalB against *M. tuberculosis* H37Rv wild type (WT), the merodiploid strain *M. tuberculosis* H37Rv pMV361::*Rv2113* and isolated SRMs raised in the merodiploid strain *M. tuberculosis* H37Rv pMV361::*Rv2113* as described in **A**. Growth relative to untreated DMSO control (100%) and sterile medium control (0%) was quantified using resazurin dye reduction assay. Data shown as means of duplicates with SD depicted as error bars.

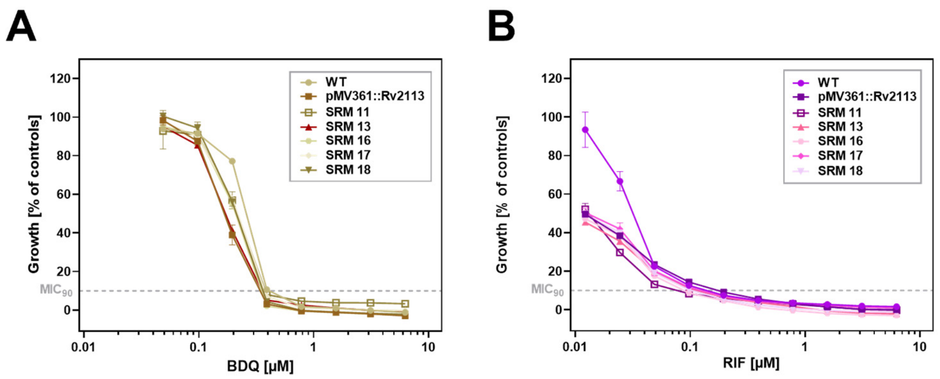

**Supplementary Figure 12.** Susceptibility profiles to bedaquiline and rifampicin treatment of spontaneous resistance mutants (SRMs) raised in the merodiploid strain *M. tuberculosis* H37Rv pMV361::*Rv2113*. **A**) Dose-response curves for **A**) bedaquiline (BDQ) or **B**) rifampicin (RIF) against wild type (WT) *M. tuberculosis* H37Rv, the merodiploid strain *M. tuberculosis* H37Rv pMV361::*Rv2113* and the CalB-resistant SRMs as reported in Supporting Figure 11. Growth relative to untreated DMSO control (corresponding to 100%) and sterile medium control (0%) was quantified using resazurin dye reduction assay. Data shown as means of duplicates with SD depicted as error bars.

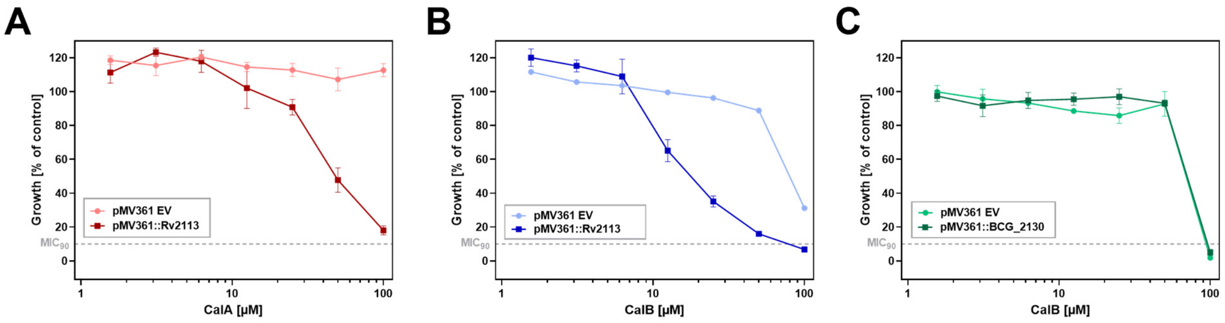

**Supplementary Figure 13.** Dose-response curves of CalA (red, **A**) and CalB (blue, **B**) against *M. smegmatis* mc² 155 pMV361::*Rv2113* cell line (dark color) compared to the pMV361 empty vector control (EV, light color). **C**: Dose-response curves of CalB against *M. smegmatis* mc² 155 pMV361::*BCG_2130* expressing the *M. bovis* BCG Pasteur´s *Rv2113* homologue *BCG_2130* (dark green) compared to the pMV361 empty vector control (EV, light green). Growth relative to untreated DMSO control (corresponding to 100%) and sterile medium control (0%) was quantified using resazurin dye reduction assay. Data shown as means of triplicates with SD depicted as error bars.

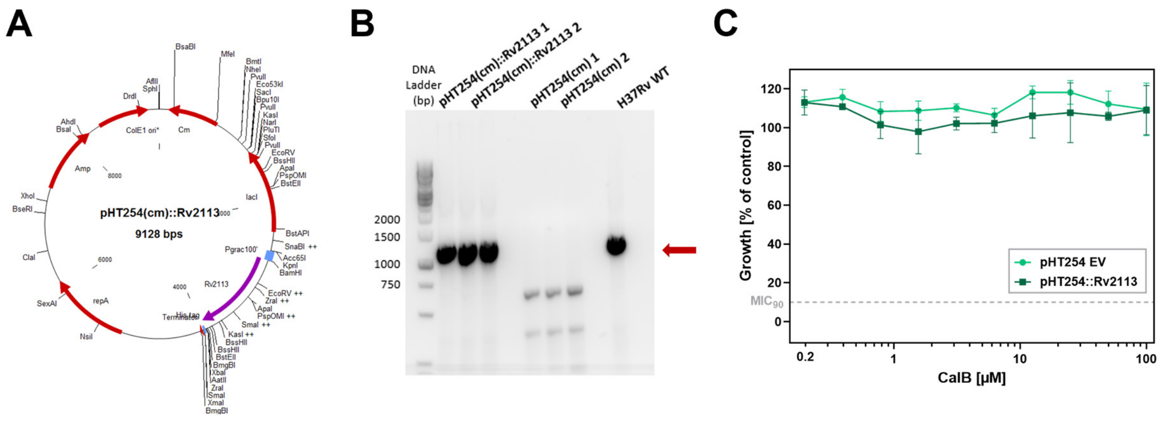

**Supplementary Figure 14**. Generation and MIC determination of *Bacillus subtilis* expressing *Rv2113*. **A**) Plasmid map of *Rv2113* cloned into the *B. subtilis*/*Escherichia coli* shuttle vector pHT254 via *Bam*HI and *Xba*I restriction sites, resulting in *B. subtilis* pHT254(cm)::*Rv2113*. **B**) Diagnostic PCR to prove presence of *Rv2113* in recombinant *B. subtilis* pHT254(cm)::*Rv2113* strains. Genomic DNA obtained from *B. subtilis* harboring the empty vector pHT254(cm) and from *M. tuberculosis* H37Rv were used as negative and positive controls, respectively. Expected band size of *Rv2113* is 1,194 base pairs (red arrow) **C**) Dose-response curves of CalB against *B. subtilis* pHT254::*Rv2113* expressing Rv2113 (dark green) compared to the pHT254 empty vector control (EV, light green). Growth relative to untreated DMSO control (corresponding to 100%) and sterile medium control (0%) was quantified using resazurin dye reduction assay. Data shown as means of duplicates with SD depicted as error bars.

| CalB Diazirine Probes | | | | | | | | | | MIC$_{90}$ [µM] | |
|---|---|---|---|---|---|---|---|---|---|---|---|
| R1 | R2 | R3 | R4 | R5 | R6 | R7 | R8 | C1 | C2 | C3 | C4 |
| | | 84 | | 85 | | | 86 | | | | Pra |
| | | 50 | | 12.5 | | | 50 | | | | |

**Supplementary Figure 15.** Overview on synthesized photoactive CalB probes for the use in an affinity-based protein-profiling approach and corresponding MIC$_{90}$ values in *M. tuberculosis* H37Rv (determined by resazurin assay, $n = 2$). The corresponding photoleucine has been introduced at different positions within the CalB core structure, while an alkyne ligation handle was introduced at C4 position via incorporation of propargylglycine (Pra).

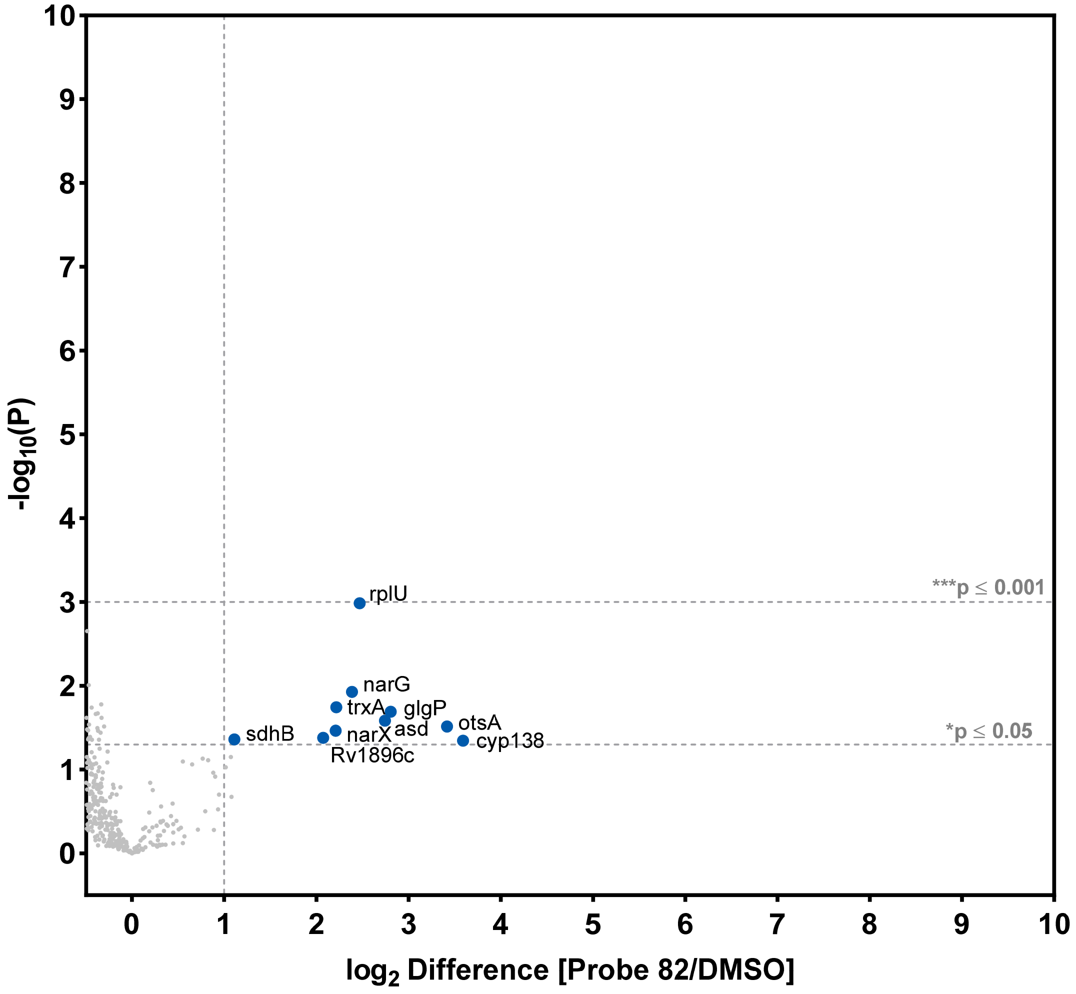

**Supplementary Figure 16.** Activity based protein profiling with a CalB-based alkyne probe (**82**) shows no covalent interaction in *M. tuberculosis* H37Rv. Proteomic data of *M. tuberculosis* H37Rv cells treated with 12.5 µM probe **82** for 3 hours compared to a DMSO control. To identify statistically significant hits from the analysis, $P \leq 0.05$ (Student's T-test; permutation-based FDR with 250 randomizations and FDR = 0.01) was applied. Proteins complying with the chosen threshold of significance and showing a $\log_2$-fold change ≥ 1 are marked in blue. Quantification via label free quantification (LFQ) algorithm of 5 replicates per sample group.

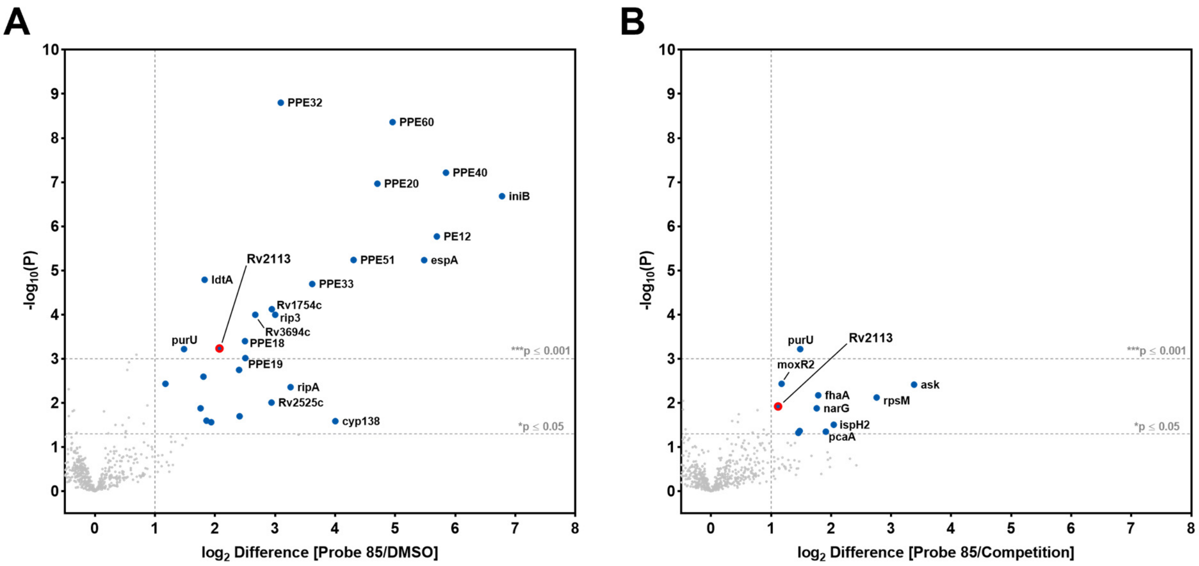

**Supplementary Figure 17.** Affinity-based protein-profiling with a CalB-based photoaffinity-tagged probe (**85**) in combination with competitive labeling in *M. tuberculosis* H37Rv. Proteomic data of *M. tuberculosis* H37Rv cells treated with 12.5 µM probe **85** for 3 hours compared to a DMSO control (**A**) or a competitive labeling approach with preincubation with 30 µM CalB (**B**). To identify statistically significant hits from the analysis, $P \leq 0.05$ (Student's T-test; permutation-based FDR with 250 randomizations and FDR = 0.01) was applied. Proteins complying with the chosen threshold of significance and showing a $\log_2$-fold change $\geq 1$ are marked in blue. Quantification was done via label free quantification (LFQ) algorithm of 5 replicates per sample group.

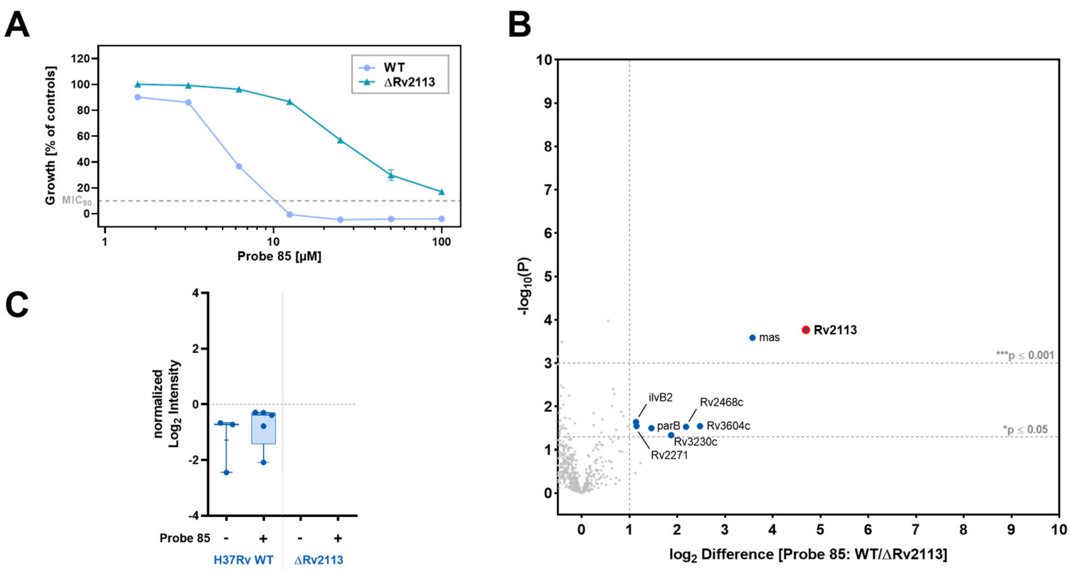

**Supplementary Figure 18.** Photoaffinity labeling of WT in comparison to the Δ*Rv2113* dene deletion mutant. **A**) Dose-response curve for CalB-based photoaffinity-tagged probe (**85**) in *M. tuberculosis* H37 WT or *ΔRv2113* gene deletion mutant (determined by resazurin assay, $n = 2$, SD depicted as error bars). **B**) Comparative affinity-based protein-profiling with **85** in *M. tuberculosis* H37 WT and *ΔRv2113* gene deletion mutant. Proteomic data of cells treated with 12.5 µM probe **85** for 3 hours compared to a DMSO control. To identify statistically significant hits from the analysis, $P \leq 0.05$ (Student's T-test; permutation-based FDR with 250 randomizations and FDR = 0.01) was applied. Proteins complying with the chosen threshold of significance and showing a $\log_2$-fold change ≥ 1 are marked in blue. Quantification was done via label free quantification (LFQ) algorithm of 5 replicates per sample group. **C**) Boxplot representation of mass quantification data from the (photoaffinity) labelling experiments (n = 5 technical replicates).

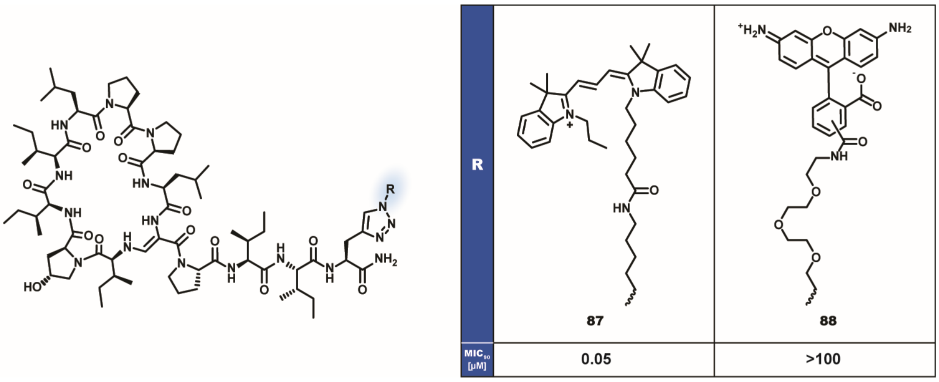

**Supplementary Figure 19.** Chemical structures and $MIC_{90}$ values against *M. tuberculosis* H37Rv (determined by resazurin assay, $n = 2$) of fluorescently-tagged C4-modified CalB derivatives. Compound **87** correspond to a Cy3 tag and **88** represents a rhodamine tag.

| CalB-Cy3 Conjugates | | | | | | | | | | | MIC$_{90}$ [µM] |
|---|---|---|---|---|---|---|---|---|---|---|---|
| R1 | R2 | R3 | R4 | R5 | R6 | R7 | R8 | C1 | C2 | C3 | C4 |
| | | 90 | 91 | | | | | | | | 87 |
| | | >1.56 | >1.56 | | | | | | | | 0.02 |

**Supplementary Figure 20.** Overview on synthesized Cy3-tagged CalB derivatives and corresponding MIC$_{90}$ values in *M. tuberculosis* H37Rv (determined by resazurin assay, $n = 2$). The corresponding Cy3 tag has been introduced at different positions within the CalB core structure. Only incorporation at C4 leads to a highly active derivative.

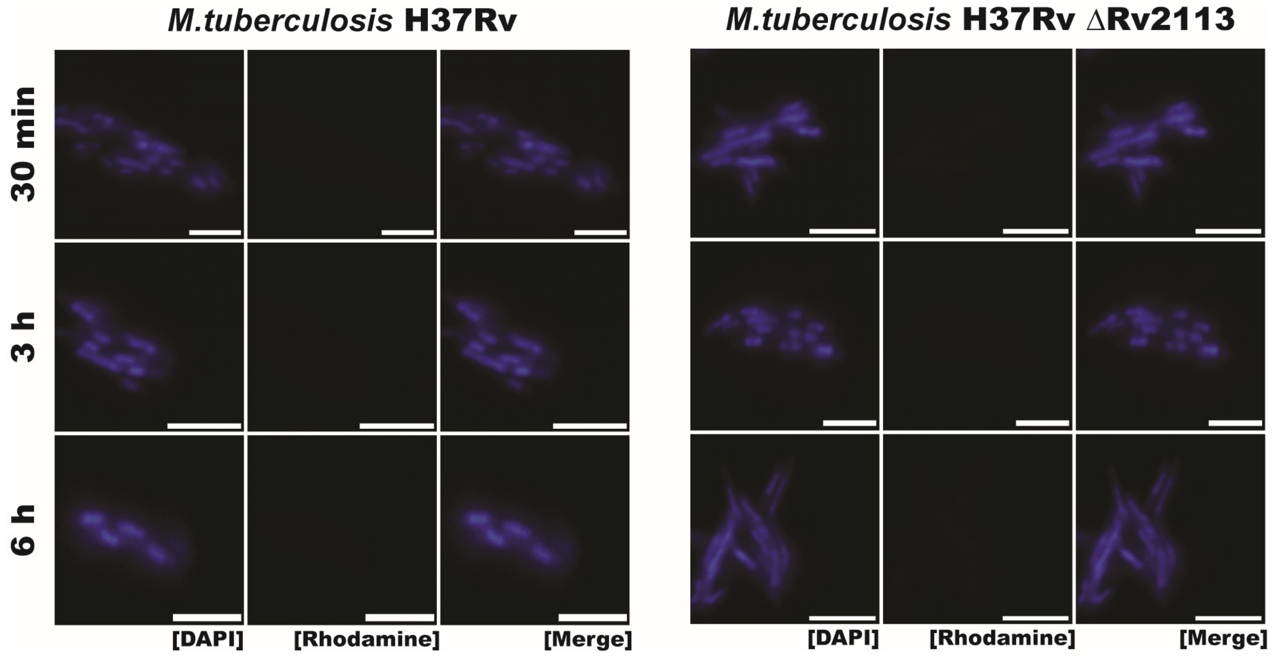

**Supplementary Figure 21.** Fluorescence microscopy images of cells of *M. tuberculosis* H37Rv treated with 0.2 µM of the rhodamine-tagged derivative **88** for the indicated time intervals. After incubation, treated cells were counterstained with DAPI to label all cells. Scale bar represents 3 µm.

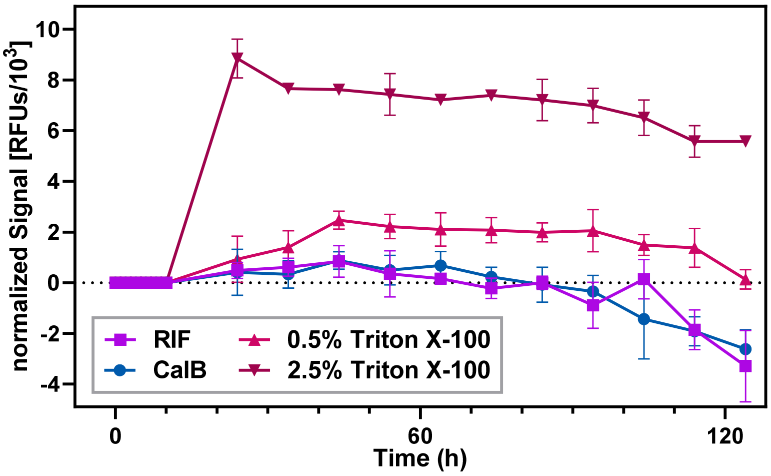

**Supplementary Figure 22.** Effect of CalB treatment on membrane integrity of *M. tuberculosis* as assessed by propidium iodide (PI) uptake assay. Cells of *M. tuberculosis* mc²6230 were preincubated with PI to establish baseline fluorescence and then treated with 15.7 µM CalB (5-fold $MIC_{90}$, blue). Fluorescence intensity (excitation: 535 nm; emission: 617 nm) was measured over time. Addition of 0.5 µM rifampicin (5-fold $MIC_{90}$, purple) and triton X-100 (0.5% as red, 2.5% as brown) served as negative and positive controls, respectively. Data shown as means of triplicates with SD depicted as error bars.

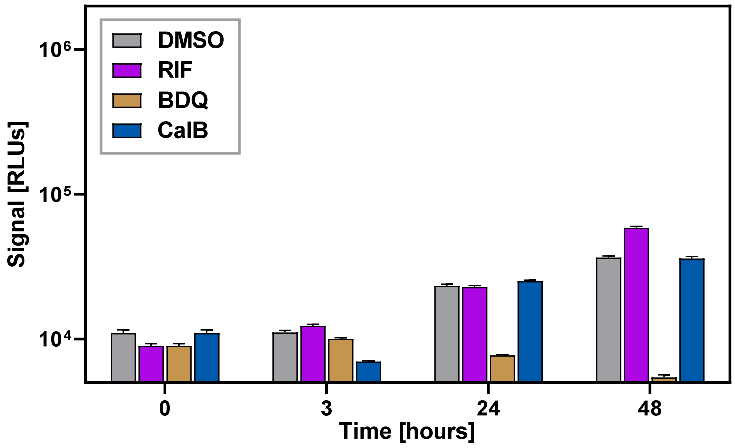

**Supplementary Figure 23.** Effect of CalB on mycobacterial energy metabolism. The amount of intracellular ATP in compound-treated cells was measured as a correlate of interference with energy metabolism. Washed cell pellets of *M. tuberculosis* $mc^2$6230 treated with 31.4 µM CalB (10-fold $MIC_{90}$, blue) were suspended in phosphate-buffered saline and incubated with Bactiter Glo reagent. Luminescence was measured at shown intervals. DMSO (grey) and 1 µM rifampicin (RIF, 10-fold $MIC_{90}$, purple) served as negative controls, while 3.9 µM bedaquiline (BDQ, 10-fold $MIC_{90}$, brown) served as a positive control. Data show means of triplicates with SD depicted as error bars.

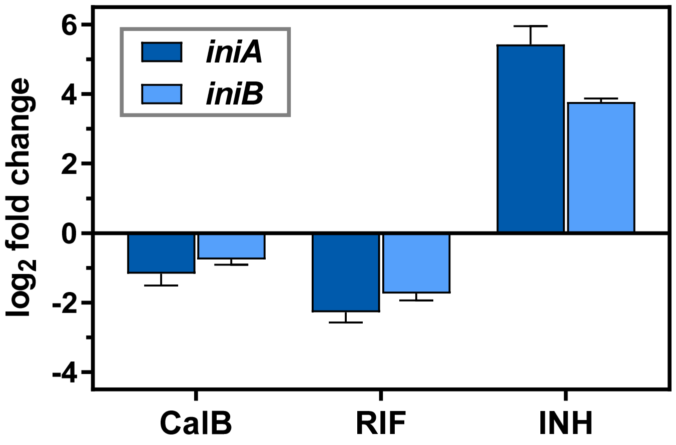

**Supplementary Figure 24.** Expression of *iniA* and *iniB* of the *iniBAC* gene operon in compound-treated cells of *M. tuberculosis*. Quantification of *iniA* (dark blue) and *iniB* (light blue) expression by RT-qPCR in cells of *M. tuberculosis* H37Rv treated with 15.7 µM CalB (5-fold $MIC_{90}$) for 24 h. $Log_2$ fold change was determined by the ΔΔCT method, normalized to 16SrRNA with DMSO-treated cells as control (corresponding to a fold change of 0). Rifampicin (0.5 µM, 5-fold $MIC_{90}$) and isoniazid (1.95 µM, 5-fold $MIC_{90}$) served as negative and positive controls, respectively. Data shown as means of triplicates with SD depicted as error bars.

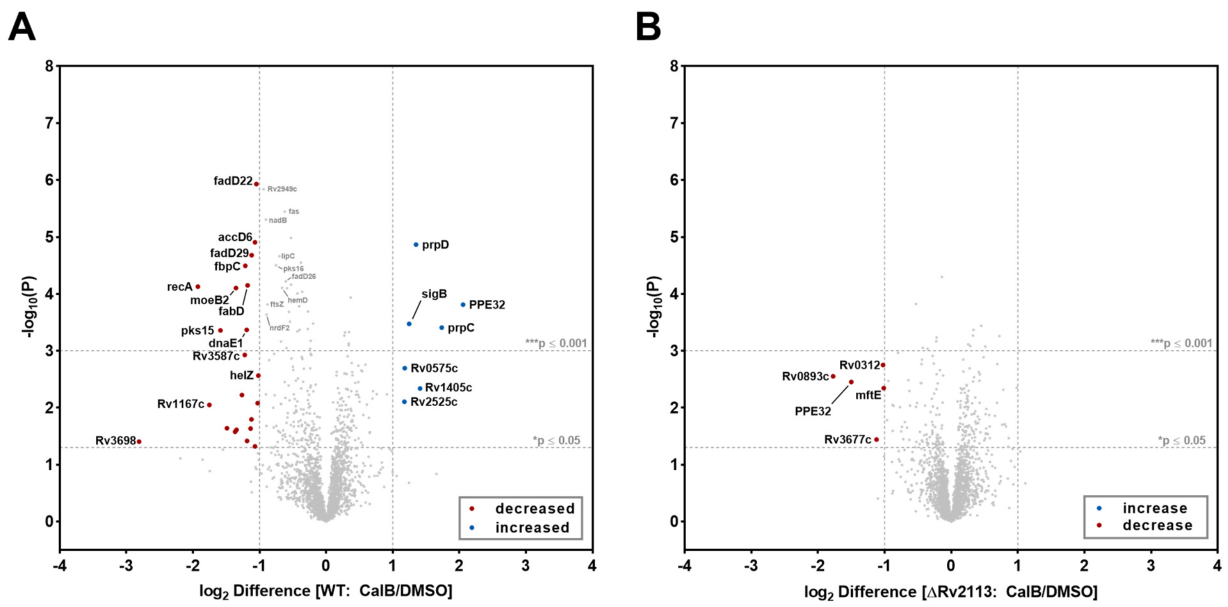

**Supplementary Figure 25.** LC-MS/MS-based proteomic stress profiling of CalB-treated *M. tuberculosis* cells. Whole protein analysis of cells of *M. tuberculosis* H37Rv wild type (WT, **A**) or the Δ*Rv2113* gene deletion mutant (**B**) treated with 31.3 µM CalB (10-fold $MIC_{90}$) for 48 hours compared to the corresponding DMSO control. To identify statistically significant hits from the analysis, $P \leq 0.05$ (Student's T-test; permutation-based FDR with 250 randomizations and FDR = 0.01) was applied. Proteins complying with the chosen threshold of significance and showing a $log_2$-fold change ≥ 1 or ≤ -1 are marked in blue or red, respectively. Quantification was done via label free quantification (LFQ) of 5 replicates per sample group.

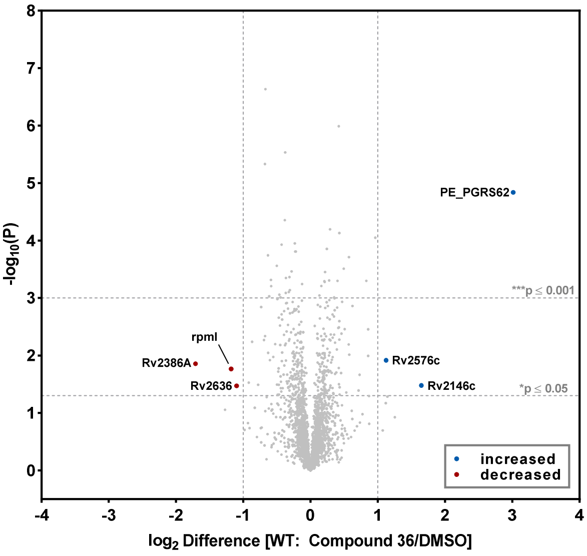

**Supplementary Figure 26.** LC-MS/MS-based proteomic stress profiling of inactive derivative **36** against *M. tuberculosis* shows no distinct response on proteome level. Whole protein analysis of *M. tuberculosis* H37Rv cells treated with 31.3 µM derivative **36** for 48 hours were compared to a DMSO control. To identify statistically significant hits from the analysis, $P \leq 0.05$ (Student's T-test; permutation-based FDR with 250 randomizations and FDR = 0.01) was applied. Proteins complying with the chosen threshold of significance and showing a $\log_2$-fold change $\geq 1$ or $\leq -1$ are marked in blue or red, respectively. Quantification was done via label free quantification (LFQ) algorithm of 5 replicates per sample group.

#### Supplementary Tables

**Table S1.** Oligonucleotides used for RT-qPCR.

| Gene | Forward primer | Reverse primer |
|---|---|---|
| ***iniA*** | CAAAGAGCTC AACGAAGAGT CC | TAGTTCGGAA CCTAGAGACA CC |
| ***iniB*** | GGGCATCGCT AGCCAGATCG | TAACGGGCCC GACAGATGAG G |
| ***16SrRNA*** | GGAATTACTG GGCGTAAAGA GC | CTCCTGATAT CTGCGCATTC C |

**Table S2.** Oligonucleotides employed for the generation of allelic exchange substrates for site-specific gene deletion of *Rv2113*. The restriction sites used for cloning of allelic exchange substrates are underlined.

| | 5´-end primer | 3´-end primer | Restr. site |
|---|---|---|---|
| Upstream flanking sequence | TTTTTCCATA AATTGGACCT GCCGGGATAC CAGAAAGGG | TTTTTCCATT TCTTGGCGAG GGCAGACCAC GCTTCAAGAA C | Van91I |
| Downstream flanking sequence | TTTTTCCATA GATTGGGCCG GTCACGTCGA TGACTAGGTT C | TTTTTCCATC TTTTGGTCGC TGGTAAGCAC GCGGGTATG | Van91I |

**Table S3.** Oligonucleotides used for cloning into mycobacterial expression plasmid pMV361. Relevant restriction sites are underlined.

| Gene | Forward primer | Restr. site | Reverse primer | Restr. site |
|---|---|---|---|---|
| ***Rv2113*** | TTTTTTTAAT TAATGAGCCT TTCCGTCCGT CGC | *Pac*I | TTTTTAAGCT TCTAGTCATC GACGTGACCG GCG | *Hind*III |
| ***BCG_2130*** | GGCCGCTTAA TTAAATGAGC CTTTCCGTCC GTC | *Pac*I | GAGCCGGAAG CTTCTAGTCA TCGACGTGAC C | *Hind*III |

### Chemical synthesis

#### General Information

All chemicals were purchased from *abcr*, *Acros Organics*, *Alpha Aesar*, *Bachem*, *BLD pharm*, *Carbolution*, *Fluorochem*, *Iris Biotech*, *Lumiprobe*, *Merck, TCI Chemicals*, or *Thermo scientific* and used without further purification.

Purification of peptide precursors or compounds via reversed-phase preparative high performance liquid chromatography (prep. HPLC) was performed either on a Prominence UFLC system from *Shimadzu* (System A) with peak detection through an UV detector at λ = 210 nm equipped with a *Phenomenex* C18 column (*Luna* 5 µm C18(2), 100 mm length, 21.2 mm ID) or a Nexera Prep HPLC system from *Shimadzu* (System B) with peak detection through a photodiode array detector at λ = 210 nm equipped with *Macherey-Nagel* C18 columns (*Nucleodur* 5 µm C18 Pyramid, 250 mm length) with either 10 mm or 32 mm ID. In both systems, the mobile phase was generated by passive mixture of solvent B (Acetonitrile (ACN) either with or without 0.1% trifluoracetic acid (TFA)) in solvent A ($H_2O$ either with or without 0.1% TFA) in a linear gradient. Purified peptide precursors or compounds were freeze-dried using an ALPHA 2-4 LD plus lyophilizer from *Christ*.

Liquid chromatography mass spectrometry (LC-MS) analysis was carried out on a *ThermoFisher Scientific* Ultimate 3000 UHPLC system (peak detection at λ = 254 nm) equipped with a *Macherey-Nagel* C18 column (*Nucleodur* 5 µm C18 Pyramid, 250 mm length, 4 mm ID) connected to a *ThermoFisher Scientific* LTQ-XL electrospray ionization mass spectrometer (ESI-MS). The mobile phase was generated by passive mixture of solvent B (ACN with 0.1% formic acid (FA)) in solvent A ($H_2O$ with 0.1% FA) in a linear gradient (as shown). In some cases, the length of the 100% solvent B plateau had to be increased for elution of the compounds. All compounds were analyzed from a 10 mM DMSO stock diluted 1:50 with 50% ACN in $H_2O$ (200 µM analyte, 2% DMSO). This leads to the presence of a DMSO peak in the chromatogram of the compounds (as shown).

| LC-MS Gradient | |
|---|---|
| **min** | **Solvent B [%]** |
| 0.0 | 10 |
| 0.5 | 10 |
| 1.5 | 60 |
| 6.0 | 100 |
| 9.2 | 100 |
| 9.5 | 10 |
| 12.0 | 10 |

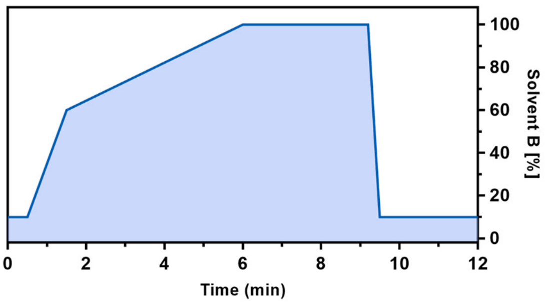

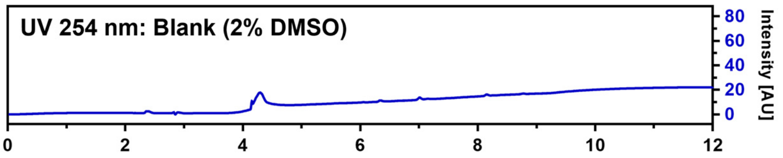

High-resolution mass spectrometry (HRMS) spectra of compounds were recorded on a *ThermoFisher Scientific* Exactive Plus EMR mass spectrometer with a TriVersa NanoMate nanoESI system from *Advion* equipped with 5 µm diameter nozzle spray chips. A total sample volume of 5 µL were picked out of 96-well plates and the ESI spray was generated using 0.8 psi nitrogen backpressure combined with a positive nozzle chip voltage of 1.7 kV. The given *m/z* values represent the average of at least 200 scans.

Nuclear magnetic resonance (NMR) spectra were recorded at 25 °C on a Bruker 400 MHz Avance II NMR spectrometer with a 5 mm PABBI broad band probe (400 MHz for $^1H$ and 101 MHz for $^{13}C$ NMR) or a 700 MHz Avance II NMR spectrometer with a 5 mm TCI cryoprobe (700 MHz for $^1H$ and 176 MHz for $^{13}C$ NMR) spectrometer. $^1H$ NMR spectra were reported in the following manner: chemical shifts (δ) in parts per million (ppm) refer to the residual non-deuterated solvent signals, multiplicities (s: singlet, d: doublet, t: triplet, dd: doublet of doublets, dt: doublet of triplets; td: triplet of doublets, m: multiplet), coupling constants (J) in Hertz (Hz) and number of protons (H).

#### Solid phase peptide synthesis

Peptide precursors for the synthesis of Callyaerin derivatives were synthesized using microwave-assisted automated solid phase peptide synthesis (SPPS) using functionalized polystyrene (PS) resin in a Liberty Blue HT12 Peptide Synthesizer from *CEM*. For this purpose, the respective resin was swollen in dimethylformamide (DMF). The peptide synthesizer was equipped with stock solutions of the corresponding fluorenylmethyloxycarbonyl (Fmoc)-protected amino acids at a concentration of 0.2 M in DMF. Furthermore, stock solutions of the activator reagent *N,N'*-Diisopropylcarbodiimide (DIC, 0.5 M) and the activator base ethyl cyano-hydroxyiminoacetate (Oxyma, 1 M) in DMF as well as a 20% piperidine in DMF solution for Fmoc-deprotection were also generated. If synthesis on Rink amide resin were performed, additional stocks of *N,N*-diisopropylethylamine (DIPEA, 1 M) and acetic anhydrate ($Ac_2O$, 0.5 M) for capping of unreacted binding positions on the resin and optional acetylation of the peptide N-terminus were also added to the device. For the capping of 2-chlorotrityl chloride (2-CTC) resin, a DCM:MeOH:DIPEA (17:2:1) mixture was used instead.

<u>Microwave-assisted SPPS with Rink-amide resin:</u>
Loading of Rink-amide resin with the first amino acid of the sequence was done by starting with an initial deprotection step with 20% piperidine in DMF using microwave heating to 90 °C for 120 seconds. Once the reactor was drained, the deprotection was repeated again for another 65 s. After washing of the resin, the first amino acid was linked to the resin by using 5 equivalents each (compared to initial resin loading capacity) of the respective amino acid, activator and activator base at microwave heating to 90 °C for 240 s. A second double-coupling step was performed, followed by a capping step by using 10 eq. of DIPEA and 5 eq. of $Ac_2OH$ at microwave heating to 50 °C for 360 s to deactivate non-reacted binding positions on the resin. After loading, the next amino acids were coupled by an iterative protocol consisting of piperidine-mediated deprotection (90 °C, 65 s) and double amino acid coupling (5 eq., 90 °C, 240 s) cycles. The synthesis was closed by a final Fmoc deprotection step (90 °C, 65 s), thereby delivering the resin-bound precursor peptide.

<u>Microwave-assisted SPPS with 2-CTC resin:</u>
Loading of 2-CTC resin with the first amino acid of the sequence was performed by using 10 eq. of the initial Fmoc-protected amino acid in combination with 20 eq. of DIPEA at microwave heating to 50 °C for 10 minutes. Once the reactor was drained, the loading step was repeated, followed by capping of unreacted binding positions using DCM:MeOH:DIPEA (17:2:1) at microwave heating to 50 °C for 10 min. After this loading procedure, an iterative protocol of deprotection with 20% piperidine in DMF (50 °C, 5 min) and double amino acid coupling using 5 equivalents each of the respective amino acid, activator and activator base at microwave heating to 50 °C for 10 min was used for elongation of the peptide sequence. The synthesis was closed by a final Fmoc deprotection step (50 °C, 5 min), thereby delivering the resin-bound precursor peptide.

<u>Cleavage of the SPPS-synthesized peptides from the resin:</u>
Peptides were cleaved from the resin by adding a mixture of 2.5% $H_2O$ and 2.5% triisopropyl silane (TIS) in trifluoroacetic acid (TFA) to the peptide-loaded resin for 3 to 5 hours at room temperature. The resin was separated from the cleavage solution which was poured into ice-cold diethyl ether. The occurring peptide precipitation was supported by overnight cooling of this solution at -20 °C. The peptides were separated from the solution via centrifugation at 4 °C and 4200 g for 30 min, the supernatant was discarded and the peptide-pellet was air-dried for further purification via prep. HPLC.

#### Synthetic procedures and characterization of Callyaerin derivatives

##### Callyaerin A (1)

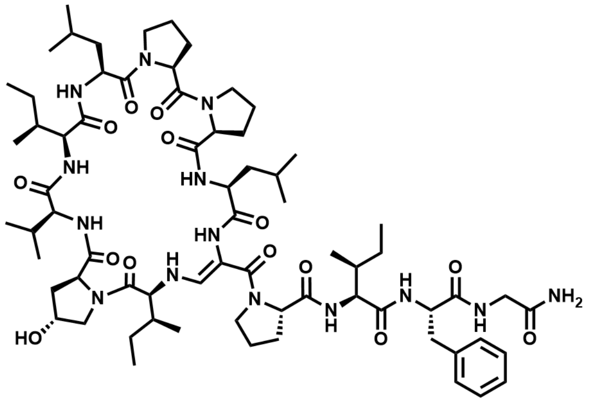

$C_{69}H_{108}N_{14}O_{14}$ M = 1357.71 g/mol

The linear precursor peptide of **1** was prepared via microwave-assisted solid-phase peptide synthesis (SPPS) using Rink amide PS resin at a scale of 0.1 mmol initial resin capacity. After purification by HPLC (solvent A: $H_2O$, acidified with 0.1% TFA; solvent B: ACN, acidified with 0.1% TFA), 38 mg (27 µmol, 27%) of the peptide precursor was obtained as a white powder. For the oxidation of the serine residue and concomitant cyclization, the peptide was dissolved in 10 mL of ACN and 3 equivalents of Dess-Martin-Periodinane (DMP) were added. The resulting suspension was shaken for 2 hours at RT and immediately purified by HPLC (solvent A: $H_2O$; solvent B: ACN) to give 6.1 mg (4.5 µmol, 16.4%) of **1** in a total yield of 4.5% (calculated in relation to initial resin capacity) as a white powder.

$^1$H NMR (700 MHz, DMSO-$d_6$) δ 9.02 (s, 1H), 8.81 (s, 1H), 8.28 (s, 1H), 8.15 (t, J = 6.1 Hz, 1H), 7.47 (d, J = 8.7 Hz, 1H), 7.44 (d, J = 7.0 Hz, 1H), 7.35 (d, J = 13.2 Hz, 1H), 7.26 (d, J = 8.6 Hz, 1H), 7.24 (s, 1H), 7.19-7.11 (m, 5H), 6.97 (s, 1H), 6.72 (d, J = 10.0 Hz, 1H), 5.87 (t, J = 11.7 Hz, 1H), 5.36 (d, J = 3.1 Hz, 1H), 4.61 (m, 2H), 4.40 (m, 1H), 4.30-4.22 (m, 4H), 4.09 (t, J = 10.2 Hz, 1H), 4.05 (t, J = 9.0 Hz, 1H), 3.90 (dd, J = 16.6, 7.1 Hz, 1H), 3.79 (d, J = 8.5 Hz, 2H), 3.77 (d, J = 11.6 Hz, 1H), 3.70 (dd, J = 11.5, 3.6 Hz, 1H), 3.64-3.57 (m, 2H), 3.52-3.49 (m, 2H), 3.48 (dd, J = 16.0, 4.6 Hz, 1H), 3.30 (m, 1H), 3.21 (td, J = 11.1, 5.8 Hz, 1H), 3.09 (m, 1H), 2.98 (m, 1H), 2.65 (m, 1H), 2.61 (m, 1H), 2.30 (m, 1H), 2.26 (m, 1H), 2.23 (m, 1H), 2.08 (m, 1H), 1.99 (m, 1H), 1.95-1.81 (m, 7H), 1.77 (m, 1H), 1.71 (m, 1H), 1.68-1.54 (m, 6H), 1.49 (m, 1H), 1.44-1.36 (m, 3H), 1.24 (m, 1H), 1.19-1.09 (m, 2H), 1.04 (m, 1H), 0.95 (d, J = 6.6 Hz, 3H), 0.88-0.86 (m, 6H), 0.85 (d, J = 6.6 Hz, 3H), 0.83-0.80 (m, 9H), 0.80-0.76 (m, 7H), 0.70 (t, J = 7.4 Hz, 3H), 0.42 (t, J = 7.3 Hz, 3H), 0.38 (d, J = 6.8 :Hz, 3H); $^{13}$C NMR (176 MHz, DMSO) δ 173.37, 172.72, 172.40, 172.33, 172.02, 172.02, 171.53, 171.47, 171.46, 171.24, 171.12, 170.83, 167.77, 143.42, 138.06, 129.02, 129.01, 127.78, 127.76 126.11, 98.17, 68.74, 66.12, 64.41, 64.04, 62.49, 61.97, 59.84, 59.27, 58.64, 56.77, 55.66, 50.06, 49.15, 48.78, 47.04, 46.12, 41.97, 41.17, 40.55, 40.02, 37.90, 37.78, 36.16, 34.96, 29.45, 28.57, 27.18, 26.23, 25.85, 25.55, 25.07, 24.92, 24.86, 24.71, 24.62, 24.07, 23.26, 22.89, 21.94, 21.08, 19.56, 19.23, 15.24, 14.92, 14.25, 11.01, 10.57, 10.26; HRMS (ESI): *m/z* calculated for $[M+H]^+$: 1357.8242, found: 1357.8235 (-0.5 ppm); calculated for $[M+2H]^{2+}$: 679.4158, found: 679.4153 (-0.7 ppm); MS$^2$ HRMS (ESI, HCD): *m/z* calculated for ring fragment $[M_{ring}+H]^+$: 926.5710, found: 926.5695 (-1.6 ppm).

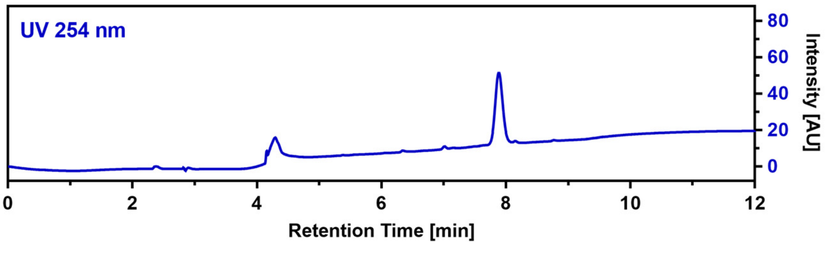

**Callyaerin B (2)**

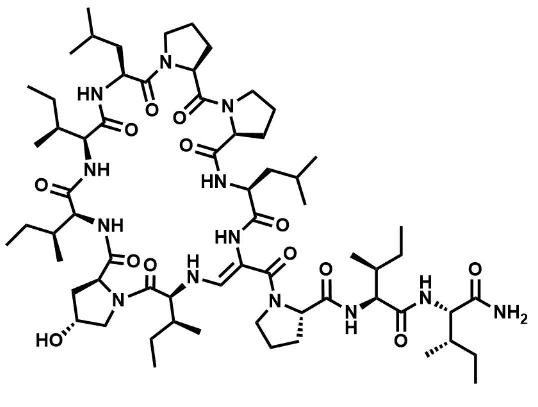

$C_{65}H_{109}N_{13}O_{13}$ M = 1280.67 g/mol

The linear precursor peptide of **2** was prepared via microwave-assisted SPPS using Rink amide PS resin at a scale of 0.1 mmol initial resin capacity. After purification by HPLC (solvent A: $H_2O$, acidified with 0.1% TFA; solvent B: ACN, acidified with 0.1% TFA), 63 mg (49 µmol, 49%) of the peptide precursor was obtained as a white powder. For the oxidation of the serine residue and concomitant cyclization, 72 mg of the peptide was dissolved in 15 mL of ACN and 3 equivalents of DMP were added. The resulting suspension was shaken for 2.5 hours at RT and immediately purified by HPLC (solvent A: $H_2O$; solvent B: ACN) to give 16.7 mg (13 µmol, 24%) of **2** in a total yield of 11.8% (calculated in relation to initial resin capacity) as a white powder.

$^{1}$H NMR (700 MHz, DMSO) δ 9.00 (s, 1H), 8.86 (s, 1H), 8.28 (s, 1H), 7.59 (d, *J* = 10.3 Hz, 1H), 7.37 (d, *J* = 6.9 Hz, 1H), 7.23 (d, *J* = 9.6 Hz, 1H), 7.16 (s, 1H), 7.05 (d, *J* = 13.2 Hz, 1H), 6.97 (s, 1H), 6.64 (d, *J* = 10.0 Hz, 1H), 5.71 (dd, *J* = 13.3, 10.1 Hz, 1H), 5.41 (s, 1H), 4.62 (q, *J* = 6.8 Hz, 1H), 4.55 (td, *J* = 10.4, 3.8 Hz, 1H), 4.41 (s, 1H), 4.26-4.24 (m, 2H), 4.23 (t, *J* = 4.0 Hz, 1H), 4.15 (dd, *J* = 9.6, 5.2 Hz, 1H), 4.10 (dd, *J* = 10.0, 8.6 Hz, 1H), 4.04 (t, *J* = 8.9 Hz, 1H), 3.95 (t, *J* = 10.2 Hz, 1H), 3.82-3.76 (m, 2H), 3.65 (dd, *J* = 11.2, 3.7 Hz, 1H), 3.63-3.56 (m, 2H), 3.52-3.45 (m, 2H), 3.32-3.28 (m, 1H), 3.15 (td, *J* = 9.9, 4.6 Hz, 1H), 3.07 (dd, *J* = 11.1, 6.8 Hz, 1H), 2.55-2.51 (m, 1H), 2.32-2.25 (m, 2H), 2.24-2.20 (m, 1H), 2.10-2.05 (m, 1H), 2.01-1.96 (m, 1H), 1.93-1.87 (m, 5H), 1.87-1.82 (m, 2H), 1.80-1.77 (m, 1H), 1.75 (dd, *J* = 13.6, 7.1 Hz, 1H), 1.72-1.66 (m, 2H), 1.64-1.56 (m, 3H), 1.55-1.50 (m, 1H), 1.49-1.37 (m, 7H), 1.30-1.25 (m, 1H), 1.25-1.21 (m, 1H), 1.18-1.09 (m, 2H), 1.07-1.02 (m, 1H), 1.02-0.95 (m, 2H), 0.90-0.80 (m, 30H), 0.79-0.77 (m, 6H), 0.75 (t, *J* = 7.4 Hz, 3H), 0.72 (t, *J* = 7.4 Hz, 3H); $^{13}$C NMR (176 MHz, DMSO) δ 173.5, 172.8, 172.7, 172.3, 172.1, 171.5, 171.4, 171.4, 171.3, 171.2, 167.2, 142.7, 98.0, 68.8, 64.6, 64.3, 64.0, 62.5, 61.4, 60.0, 59.3, 57.9, 57.5, 56.7, 50.1, 49.2, 48.7, 47.1, 46.2, 41.2, 40.5, 37.9, 37.5, 36.4, 36.2, 36.2, 32.4, 29.7, 28.5, 26.3, 25.8, 25.6, 25.1, 24.9, 24.9, 24.8, 24.6, 23.9, 23.5, 23.1, 22.7, 22.2, 20.9, 15.8, 15.7, 15.3, 15.3, 14.4, 11.5, 11.1, 10.8, 10.3, 9.9; HRMS (ESI): *m/z* calculated for $[M+H]^+$: 1280.8341, found: 1280.8322 (-1.5 ppm); calculated for $[M+2H]^{2+}$: 640.9207, found: 640.9195(-1.9 ppm); MS$^2$ HRMS (ESI, HCD): *m/z* calculated for ring fragment $[M_{ring}+H]^+$: 940.5866, found: 940.5832 (-3.6 ppm).

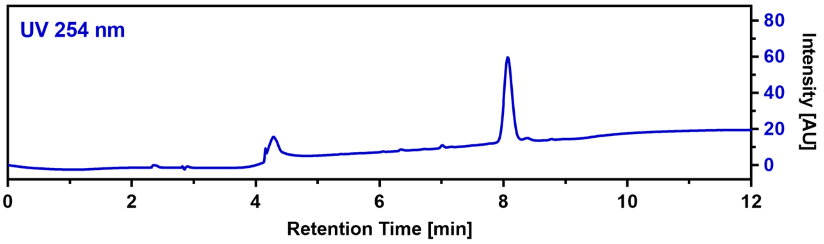

**Callyaerin C (3)**

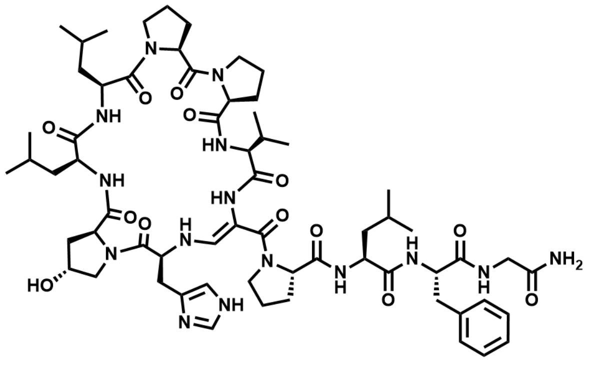

$C_{63}H_{93}N_{15}O_{13}$ M = 1268.53 g/mol

The linear precursor peptide of **3** was prepared via SPPS using Rink amide PS resin at a scale of 0.1 mmol initial resin capacity and purified by HPLC (solvent A: $H_2O$, acidified with 0.1% TFA; solvent B: ACN, acidified with 0.1% TFA). The obtained peptide was then dissolved in 10 mL of ACN and 4 equivalents DMP were added. The resulting suspension was shaken for one hour at RT and immediately purified by HPLC (solvent A: $H_2O$; solvent B: ACN) to give 8.7 mg (6.9 µmol) of **3** in a total yield of 6.9% (calculated in relation to initial resin capacity) as a white powder.

$^1$H NMR (400 MHz, DMSO) δ 12.03 (s, 1H), 9.84 (d, *J* = 7.7 Hz, 1H), 8.57 (s, 1H), 7.90 (d, *J* = 7.2 Hz, 1H), 7.80 (d, *J* = 9.3 Hz, 1H), 7.73 (dd, *J* = 7.1, 5.2 Hz, 1H), 7.65 (d, *J* = 13.8 Hz, 1H), 7.57 (d, *J* = 6.3 Hz, 1H), 7.39 (s, 1H), 7.37-7.34 (m, 2H), 7.32 (d, *J* = 10.4 Hz, 1H), 7.19 (s, 1H), 7.18-7.14 (m, 3H), 7.09 (s, 1H), 6.00 (s, 1H), 5.64 (dd, *J* = 13.8, 9.9 Hz, 1H), 5.09 (d, *J* = 3.7 Hz, 1H), 4.67 (dd, *J* = 10.2, 4.4 Hz, 1H), 4.52-4.45 (m, 1H), 4.45-4.37 (m, 2H), 4.35-4.27 (m, 2H), 4.27-4.20 (m, 1H), 4.19-4.10 (m, 2H), 4.07 (s, 1H), 3.95-3.83 (m, 2H), 3.71-3.62 (m, 1H), 3.60-3.45 (m, 6H), 3.30-3.25 (m, 1H), 3.23-3.14 (m, 1H), 2.99 (t, *J* = 12.6 Hz, 1H), 2.59-2.52 (m, 1H), 2.38-2.26 (m, 3H), 2.25-2.16 (m, 2H), 2.12-2.06 (m, 1H), 2.03-1.98 (m, 1H), 1.97-1.85 (m, 6H), 1.83-1.70 (m, 3H), 1.67-1.56 (m, 6H), 1.54-1.43 (m, 2H), 1.26-1.15 (m, 2H), 1.11 (d, *J* = 6.9 Hz, 3H), 0.97 (d, *J* = 6.9 Hz, 3H), 0.91-0.84 (m, 12H), 0.82 (d, *J* = 6.5 Hz, 3H), 0.75 (d, *J* = 6.4 Hz, 3H); HRMS (ESI): *m/z* calculated for $[M+H]^+$: 1568.7150, found: 1268.7173 (1.8 ppm); calculated for $[M+2H]^{2+}$: 634.8612, found: 634.8616 (0.6 ppm); $MS^2$ HRMS (ESI, HCD): *m/z* calculated for ring fragment $[M_{ring}+H]^+$: 837.4618, found: 837.4627 (1.1 ppm).

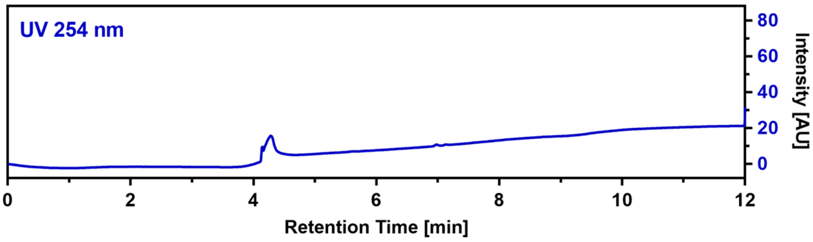

**Callyaerin D (4)**

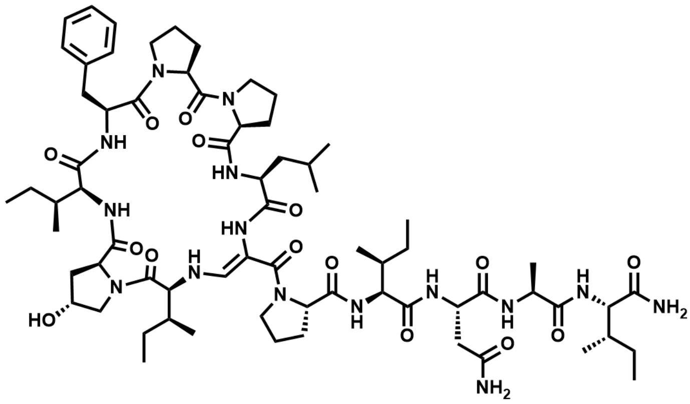

$C_{69}H_{107}N_{15}O_{15}$ M = 1386.71 g/mol

The linear precursor peptide of **4** was prepared via SPPS using Rink amide PS resin at a scale of 0.1 mmol initial resin capacity and purified by HPLC (solvent A: $H_2O$, acidified with 0.1% TFA; solvent B: ACN, acidified with 0.1% TFA). The obtained peptide was then dissolved in 10 mL of ACN and 4 equivalents of DMP were added. The resulting suspension was shaken for one hour at RT and immediately purified by HPLC (solvent A: $H_2O$; solvent B: ACN) to give 5.7 mg (4.1 µmol) of **4** in a total yield of 4.1% (calculated in relation to initial resin capacity) as a white powder.

$^1$H NMR (400 MHz, DMSO) δ 8.42 (s, 1H), 7.88 (d, $J$ = 7.5 Hz, 1H), 7.76 (d, $J$ = 6.8 Hz, 1H), 7.64 (d, $J$ = 5.0 Hz, 1H), 7.56 (d, $J$ = 7.1 Hz, 1H), 7.53 (d, $J$ = 9.3 Hz, 1H), 7.50 (d, $J$ = 6.0 Hz, 1H), 7.38 (d, $J$ = 3.3 Hz, 1H), 7.35 (s, 1H), 7.32-7.22 (m, 6H), 7.10 (s, 1H), 7.02 (s, 1H), 6.92 (s, 1H), 5.29 (dd, $J$ = 13.8, 10.1 Hz, 1H), 5.19 (d, $J$ = 3.8 Hz, 1H), 4.65-4.59 (m, 1H), 4.55 (q, $J$ = 7.0 Hz, 1H), 4.51-4.44 (m, 1H), 4.44-4.38 (m, 2H), 4.36-4.30 (m, 2H), 4.21-4.11 (m, 3H), 4.05 (dd $J$ = 8.9, 6.8 Hz, 1H), 4.00 (dd, $J$ = 5.0, 3.4 Hz, 1H), 3.96-3.90 (m, 2H), 3.68-3.60 (m, 2H), 3.56-3.49 (m, 1H), 3.40-3.35 (m, 1H), 3.28-3.23 (m, 1H), 3.06 (dd, $J$ = 13.1, 9.5 Hz, 1H), 2.80-2.73 (m, 1H), 2.73-2.66 (m, 1H), 2.39-2.30 (m, 1H), 2.27-2.12 (m, 4H), 2.10-2.04 (m, 1H), 2.02-1.88 (m, 5H), 1.86-1.68 (m, 9H), 1.65-1.56 (m, 1H), 1.52-1.30 (m, 8H), 1.27 (d, $J$ = 7.2 Hz, 3H), 1.23 (s, 1H), 1.17-1.06 (m, 1H), 0.98-0.91 (m, 1H), 0.89-0.79 (m, 30H); HRMS (ESI): $m/z$ calculated for $[M+H]^+$: 1386.8144, found: 1386.8153 (0.6 ppm); calculated for $[M+2H]^{2+}$: 693.9109, found: 693.9108 (-0.1 ppm); MS$^2$ HRMS (ESI, HCD): $m/z$ calculated for ring fragment $[M_{ring}+H]^+$: 861.4869, found: 861.4870 (0.1 ppm).

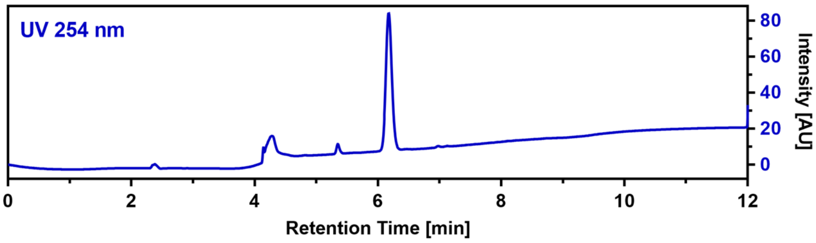

**Callyaerin E (5)**

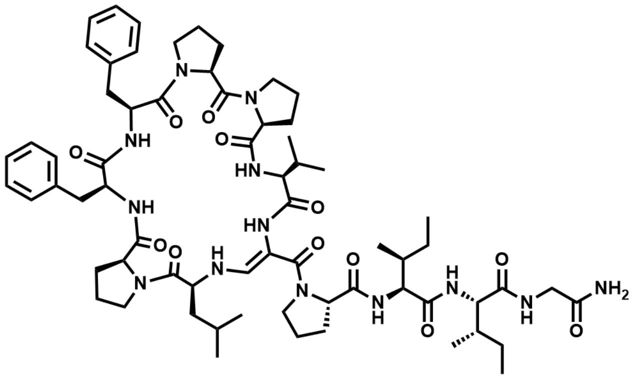

The linear precursor peptide of **5** was prepared via SPPS using Rink amide PS resin at a scale of 0.23 mmol initial resin capacity. After purification by HPLC (solvent A: $H_2O$, acidified with 0.1% TFA; solvent B: ACN, acidified with 0.1% TFA), 130 mg (101 µmol, 44%) of the peptide precursor was obtained as a white powder. For the oxidation of the serine residue and concomitant cyclization, the peptide was dissolved in 10 mL of ACN and 4 equivalents of DMP were added. The resulting suspension was shaken for one hour at RT and immediately purified by HPLC (solvent A: $H_2O$; solvent B: ACN) to give 60.1 mg (47.6 µmol, 46.9%) of **5** in a total yield of 20.7% (calculated in relation to initial resin capacity) as a white powder.

$^1$H NMR (400 MHz, DMSO) δ 8.37 (s, 1H), 7.89 (t, $J$ = 6.0 Hz, 1H), 7.72 (d, $J$ = 6.7 Hz, 1H), 7.63 (d, $J$ = 8.0 Hz, 1H), 7.37 (d, $J$ = 8.2 Hz, 1H), 7.33-7.16 (m, 12H), 7.06-7.01 (m, 2H), 6.99-6.94 (m, 1H), 5.27 (dd, $J$ = 13.5, 9.7 Hz, 1H), 4.68-4.60 (m, 1H), 4.56 (dd, $J$ = 10.2, 3.9 Hz, 1H), 4.45-4.32 (m, 4H), 4.16 (dd, $J$ = 9.9, 7.5 Hz, 1H), 4.09 (t, $J$ = 7.7 Hz, 1H), 4.04-3.96 (m, 2H), 3.96-3.90 (m, 1H), 3.71 (dd, $J$ = 16.6, 6.2 Hz, 1H), 3.63-3.54 (m, 3H), 3.53-3.46 (m, 1H), 3.46-3.37 (m, 1H), 3.33-3.29 (m, 1H), 3.26-3.16 (m, 1H), 3.04-2.95 (m, 3H), 2.71 (dd, $J$ = 13.4, 4.8 Hz, 1H), 2.49-2.43 (m, 1H), 2.37-2.27 (m, 1H), 2.25-2.11 (m, 4H), 2.10-2.01 (m, 1H), 1.98-1.53 (m, 14H), 1.49-1.34 (m, 3H), 1.30-1.16 (m, 3H), 1.05 (d, $J$ = 6.9 Hz, 3H), 0.96 (dd, $J$ = 6.7, 3.3 Hz, 6H), 0.90 (d, $J$ = 6.6 Hz, 3H), 0.88-0.82 (m, 9H), 0.79 (t, $J$ = 7.3 Hz, 3H); $^{13}$C NMR (101 MHz, DMSO) δ 173.5, 172.6, 172.1, 171.7, 171.5, 171.5, 171.3, 171.0, 171.0, 169.1, 167.7, 143.1, 138.1, 137.9, 129.0, 128.4, 128.2, 128.1, 126.5, 126.4, 98.5, 63.5, 63.1, 61.0, 58.4, 57.8, 56.6, 54.0, 53.0, 48.7, 46.6, 46.4, 42.3, 42.2, 36.1, 35.5, 35.4, 35.3, 29.9, 28.8, 28.7, 27.0, 25.7, 25.5, 24.8, 24.4, 24.2, 23.8, 20.8, 18.3, 17.7, 15.7, 15.5, 11.2,

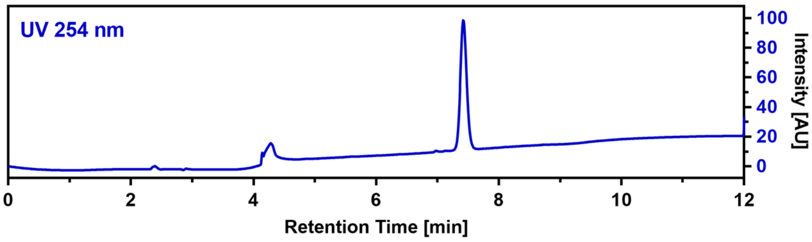

10.6; HRMS (ESI): *m/z* calculated for $[M+H]^+$: 1262.7296, found: 1262.7304 (0.6 ppm); calculated for $[M+2H]^{2+}$: 631.8685, found: 631.8686 (0.2 ppm); $MS^2$ HRMS (ESI, HCD): *m/z* calculated for ring fragment $[M_{ring}+H]^+$: 865.4607, found: 865.4597 (-1.2 ppm).

**Callyaerin F (6)**

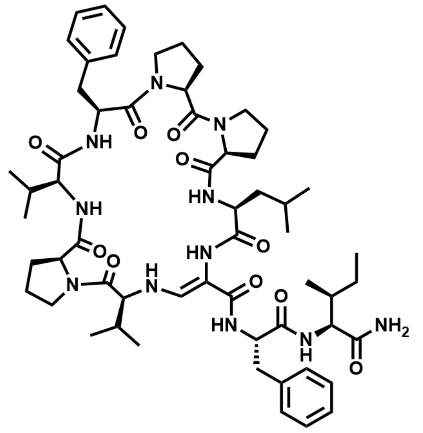

$C_{58}H_{83}N_{11}O_{10}$ M = 1094.37 g/mol

The linear precursor peptide of **6** was prepared via SPPS using Rink amide PS resin at a scale of 0.23 mmol initial resin capacity. After purification by HPLC (solvent A: $H_2O$, acidified with 0.1% TFA; solvent B: ACN, acidified with 0.1% TFA), 116 mg (104 µmol, 45%) of the peptide precursor was obtained as a white powder. For the oxidation of the serine residue and concomitant cyclization, the peptide was dissolved in 10 mL of ACN and 4 equivalents of DMP were added. The resulting suspension was shaken for one hour at RT and immediately purified by HPLC (solvent A: $H_2O$; solvent B: ACN) to give 14.1 mg (12.9 µmol, 12.4%) of **6** in a total yield of 5.6% (calculated in relation to initial resin capacity) as a white powder.

$^1$H NMR (400 MHz, DMSO) δ 8.25 (s, 1H), 7.66 (d, *J* = 9.6 Hz, 1H), 7.57-7.46 (m, 3H), 7.32-7.13 (m, 11H), 7.03 (s, 1H), 6.98 (s, 1H), 6.17 (d, *J* = 7.8 Hz, 1H), 5.14 (dd, *J* = 13.7, 10.0 Hz, 1H), 4.63 (td, *J* = 10.2, 3.9 Hz, 1H), 4.57-4.50 (m, 1H), 4.45-4.39 (m, 1H), 4.37 (d, *J* = 2.0 Hz, 1H), 4.31-4.23 (m, 2H), 4.01-3.92 (m, 2H), 3.89-3.82 (m, 2H), 3.77-3.68 (m, 1H), 3.59 (t, *J* = 8.3 Hz, 1H), 3.39-3.35 (m, 1H), 3.24 (t, *J* = 8.9 Hz, 1H), 3.01 (dd, *J* = 12.9, 9.6 Hz, 1H), 2.96-2.87 (m, 2H), 2.74 (dd, *J* = 12.9, 3.5 Hz, 1H), 2.35-2.26 (m, 2H), 2.22-2.00 (m, 5H), 1.97-1.88 (m, 2H), 1.88-1.77 (m, 4H), 1.77-1.63 (m, 4H), 1.50-1.38 (m, 2H), 1.33-1.21 (m, 1H), 1.16-1.03 (m, 1H), 0.93 (d, *J* = 4.7 Hz, 3H), 0.91 (d, *J* = 4.2 Hz, 3H), 0.88-0.84 (m, 6H), 0.82 (d, *J* = 6.3 Hz, 3H), 0.79-0.74 (m, 6H), 0.65 (d, *J* = 6.6 Hz, 3H); HRMS (ESI): *m/z* calculated for $[M+H]^+$: 1094.6397, found: 1094.6412 (1.4 ppm); calculated for $[M+2H]^{2+}$: 547.8235, found: 547.8242(1.3 ppm); $MS^2$ HRMS (ESI, HCD): *m/z* calculated for ring fragment $[M_{ring}+H]^+$: 817.4607, found: 817.4609 (0.2 ppm).

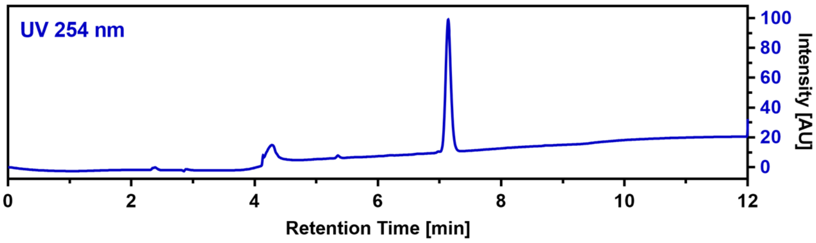

**Callyaerin G (7)**

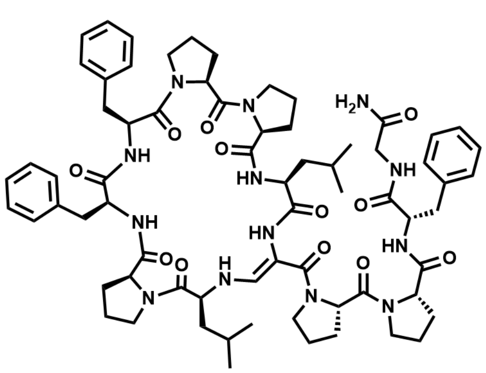

$C_{69}H_{91}N_{13}O_{12}$ M = 1294.57 g/mol

The linear precursor peptide of **7** was prepared via SPPS using Rink amide PS resin at a scale of 0.23 mmol initial resin capacity. After purification by HPLC (solvent A: $H_2O$, acidified with 0.1% TFA; solvent B: ACN, acidified with 0.1% TFA), 83 mg (63 µmol, 27%) of the peptide precursor was obtained as a white powder. For the oxidation of the serine residue and concomitant cyclization, the peptide was dissolved in 10 mL of ACN and 4 equivalents of DMP were added. The resulting suspension was shaken for one hour at RT and immediately purified by HPLC (solvent A: $H_2O$; solvent B: ACN) to give 35 mg (27 µmol, 42.9%) of **7** in a total yield of 11.7% (calculated in relation to initial resin capacity) as a white powder.

$^1$H NMR (400 MHz, DMSO) δ 8.24 (s, 1H), 7.98 (dd, *J* = 7.9, 4.8 Hz, 1H), 7.75-7.66 (m, 2H), 7.52 (d, *J* = 9.7 Hz, 1H), 7.47 (d, *J* = 13.5 Hz, 1H), 7.38-7.34 (m, 2H), 7.30-7.10 (m, 16H), 5.33 (dd, *J* = 13.5, 9.9 Hz, 1H), 4.64 (q, *J* = 7.0 Hz, 1H), 4.53-4.40 (m, 2H), 4.38-4.20 (m, 5H), 4.11 (dd,

$J$ = 8.4, 6.6 Hz, 1H), 3.99-3.81 (m, 5H), 3.54-3.44 (m, 2H), 3.43-3.35 (m, 4H), 3.25 (dd, $J$ = 13.2, 3.7 Hz, 1H), 3.04-2.94 (m, 2H), 2.93-2.84 (m, 2H), 2.79-2.72 (m, 1H), 2.68 (dd, $J$ = 13.2, 6.3 Hz, 1H), 2.34-2.23 (m, 1H), 2.20-2.07 (m, 3H), 2.06-1.93 (m, 3H), 1.93-1.82 (m, 7H), 1.82-1.73 (m, 4H), 1.72-1.58 (m, 5H), 1.55-1.43 (m, 1H), 1.41-1.29 (m, 1H), 1.15-1.01 (m, 2H), 0.88 (d, $J$ = 6.3 Hz, 3H), 0.84 (d, $J$ = 6.2 Hz, 3H), 0.54 (d, $J$ = 6.6 Hz, 3H), 0.38 (d, $J$ = 6.3 Hz, 3H); $^{13}$C NMR (101 MHz, DMSO) δ 173.3, 172.1, 171.9, 171.6, 171.4, 171.0, 171.0, 170.9, 170.7, 169.6, 168.1, 145.6, 138.2, 138.2, 137.7, 129.9, 129.2, 128.4, 128.2, 128.1, 128.0, 126.5, 126.3, 125.9, 98.0, 63.7, 63.0, 62.8, 61.7, 57.6, 54.6, 54.1, 52.5, 50.3, 47.7, 46.7, 46.3, 42.2, 36.9, 36.7, 35.2, 28.7, 27.9, 26.8, 25.6, 25.0, 24.4, 23.3, 22.9, 20.8, 20.2; HRMS (ESI): *m/z* calculated for $[M+H]^+$: 1294.6983, found: 1294.7013 (2.3 ppm); calculated for $[M+2H]^{2+}$: 647.8528, found: 847.8531 (0.5 ppm); MS$^2$ HRMS (ESI, HCD): *m/z* calculated for ring fragment $[M_{ring}+H]^+$: 879.4763, found: 879.4742 (-2.4 ppm).

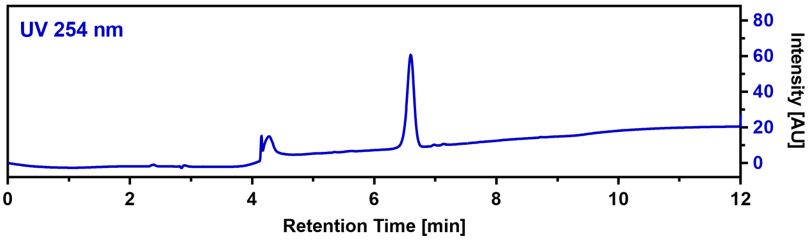

**Callyaerin H (8)**

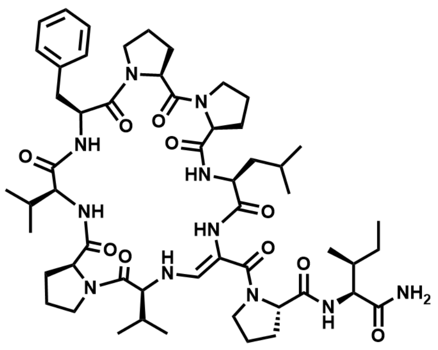

$C_{54}H_{81}N_{11}O_{10}$ M = 1044.31 g/mol

The linear precursor peptide of **8** was prepared via SPPS using Rink amide PS resin at a scale of 0.23 mmol initial resin capacity. After purification by HPLC (solvent A: $H_2O$, acidified with 0.1% TFA; solvent B: ACN, acidified with 0.1% TFA), 98 mg (92 µmol, 40%) of the peptide precursor was obtained as a white powder. For the oxidation of the serine residue and concomitant cyclization, the peptide was dissolved in 10 mL of ACN and 4 equivalents of DMP were added. The resulting suspension was shaken for one hour at RT and immediately purified by HPLC (solvent A: $H_2O$; solvent B: ACN) to give 4.7 mg (4.5 µmol, 4.9%) of **8** in a total yield of 2.0% (calculated in relation to initial resin capacity) as a white powder.

$^1$H NMR (400 MHz, DMSO) δ 8.47 (s, 1H), 7.59 0(d, $J$ = 5.1 Hz, 1H), 7.55 (d, $J$ = 9.6 Hz, 1H), 7.52 (s, 1H), 7.50 (d, $J$ = 5.5 Hz, 1H), 7.31-7.20 (m, 6H), 6.97 (s, 1H), 6.89 (s, 1H), 5.26 (dd, $J$ = 13.9, 9.9 Hz, 1H), 4.61-4.48 (m, 3H), 4.43 (d, $J$ = 1.9 Hz, 1H), 4.37-4.31 (m, 2H), 4.17 (dd, $J$ = 9.6, 7.7 Hz, 1H), 4.05-4.00 (m, 1H), 4.00-3.94 (m, 1H), 3.86 (t, $J$ = 4.7 Hz, 1H), 3.84-3.79 (m, 1H), 3.79-3.71 (m, 1H), 3.65-3.58 (m, 1H), 3.39-3.32 (m, 1H), 3.30-3.22 (m, 2H), 3.04 (dd, $J$ = 13.0, 9.5 Hz, 1H), 2.77 (dd, $J$ = 13.0, 3.6 Hz, 1H), 2.40-2.30 (m, 1H), 2.28-2.15 (m, 4H), 2.14-2.07 (m, 2H), 2.07-2.02 (m, 1H), 1.99-1.89 (m, 2H), 1.89-1.81 (m, 4H), 1.80-1.65 (m, 5H), 1.60-1.36 (m, 4H), 1.24 (s, 1H), 1.22-1.13 (m, 1H), 0.93 (d, $J$ = 6.9 Hz, 3H), 0.90-0.80 (m, 18H), 0.68 (d, $J$ = 6.6 Hz, 3H); HRMS (ESI): *m/z* calculated for $[M+H]^+$: 1044.6241, found: 1044.6243 (0.2 ppm); calculated for $[M+2H]^{2+}$: 522.8157, found: 522.8137 (-3.8 ppm); MS$^2$ HRMS (ESI, HCD): *m/z* calculated for ring fragment $[M_{ring}+H]^+$: 817.4607, found: 817.4602 (-0.6 ppm).

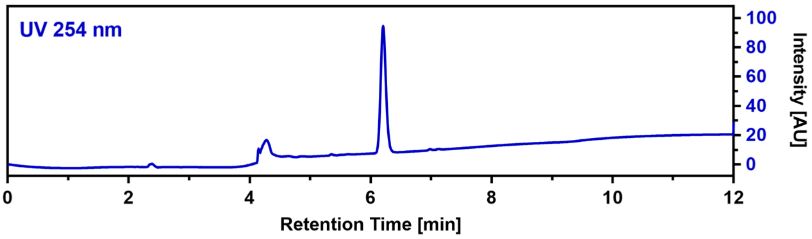

**Callyaerin I (9)**

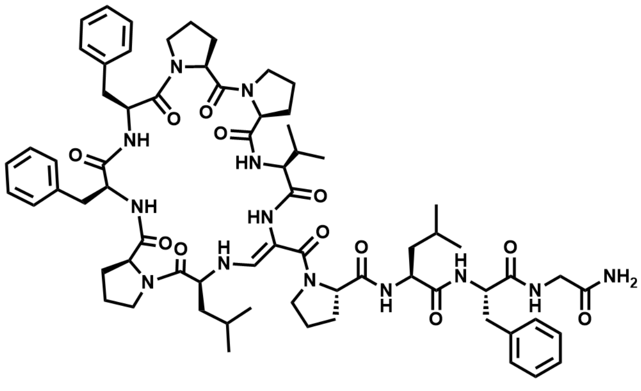

$C_{69}H_{93}N_{13}O_{12}$ M = 1296.58 g/mol

The linear precursor peptide of **9** was prepared via SPPS using Rink amide PS resin at a scale of 0.23 mmol initial resin capacity. After purification by HPLC (solvent A: $H_2O$, 0acidified with 0.1% TFA; solvent B: ACN, acidified with 0.1% TFA), 117 mg (89 µmol, 39%) of the peptide precursor was obtained as a white powder. For the oxidation of the serine residue and concomitant cyclization, the peptide was dissolved in 10 mL of ACN and 4 equivalents of DMP were added. The resulting suspension was shaken for one hour at RT and immediately purified by HPLC (solvent A: $H_2O$; solvent B: ACN) to give 47.7 mg (36.8 µmol, 41.3%) of **9** in a total yield of 16% (calculated in relation to initial resin capacity) as a white powder.

$^1$H NMR (400 MHz, DMSO) δ 8.45 (s, 1H), 7.78 (d, *J* = 6.9 Hz, 1H), 7.68 (d, *J* = 9.7 Hz, 1H), 7.65 (d, *J* = 6.7 Hz, 1H), 7.53 (dd, *J* = 7.3, 5.1 Hz, 1H), 7.44 (d, *J* = 13.5 Hz, 1H), 7.32-7.22 (m, 11H), 7.21-7.18 (m, 1H), 7.16-7.11 (m, 6H), 7.03 (d, *J* = 10.1 Hz, 1H), 5.37 (dd, *J* = 13.6, 9.8 Hz, 1H), 4.68-4.59 (m, 2H), 4.47-4.35 (m, 3H), 4.30 (dd, *J* = 10.7, 7.2 Hz, 1H), 4.27-4.21 (m, 1H), 4.18 (dd, *J* = 10.0, 7.4 Hz, 1H), 4.01-3.94 (m, 2H), 3.89 (dd, *J* = 17.0, 7.4 Hz, 1H), 3.85-3.78 (m, 1H), 3.68-3.60 (m, 1H), 3.56-3.45 (m, 3H), 3.45-3.38 (m, 1H), 3.38-3.30 (m, 1H), 3.29-3.21 (m, 2H), 3.01-2.92 (m, 3H), 2.81-2.72 (m, 1H), 2.68 (dd, *J* = 13.3, 4.7 Hz, 1H), 2.48-2.43 (m, 1H), 2.34-2.27 (m, 1H), 2.26-2.19 (m, 1H), 2.18-2.10 (m, 2H), 2.07-1.99 (m, 1H), 1.96-1.82 (m, 4H), 1.81-1.68 (m, 4H), 1.67-1.49 (m, 6H), 1.48-1.36 (m, 1H), 1.26-1.22 (m, 1H), 1.20 (d, *J* = 6.9 Hz, 3H), 1.15-1.10 (m, 1H), 1.07 (d, *J* = 7.0 Hz, 3H), 1.04-0.98 (m, 1H), 0.87 (d, *J* = 6.5 Hz, 3H), 0.74 (d, *J* = 6.5 Hz, 3H), 0.44 (d, *J* = 6.4 Hz, 3H), 0.32 (d, *J* = 6.6 Hz, 3H).; $^{13}$C NMR (101 MHz, DMSO) δ 174.2, 173.5, 172.1, 171.8, 171.7, 171.0, 170.7, 170.7, 169.1, 168.4, 143.5, 138.3, 138.1, 137.9, 129.8, 129.0, 128.4, 128.2, 128.1, 127.7, 126.5, 126.4, 126.0, 98.1, 63.5, 63.2, 63.0, 62.0, 57.6, 56.9, 53.8, 52.9, 48.8, 46.7, 46.4, 46.1, 42.2, 38.1, 37.7, 36.0, 35.2, 30.2, 29.4, 28.9, 28.7, 27.0, 26.0, 25.5, 24.9, 24.4, 24.2, 23.5, 23.1, 22.7, 20.8, 19.9, 18.2, 17.7; HRMS (ESI): *m/z* calculated for $[M+H]^+$: 1296.7139, found: 1296.7161 (1.7 ppm); calculated for $[M+2H]^{2+}$: 648.8606, found: 648.8610 (0.6 ppm); MS$^2$ HRMS (ESI, HCD): *m/z* calculated for ring fragment $[M_{ring}+H]^+$: 865.4607, found: 865.4597 (-1.2 ppm).

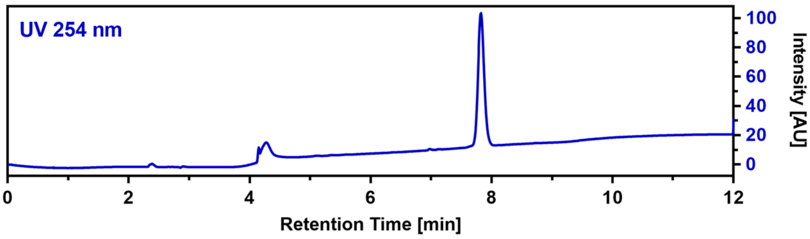

**Callyaerin J (10)**

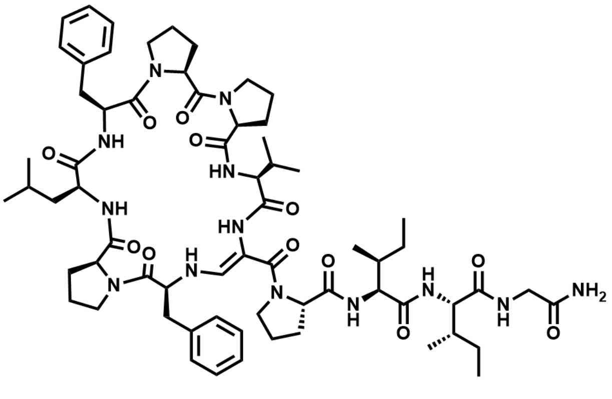

$C_{66}H_{95}N_{13}O_{12}$ M = 1262.57 g/mol

The linear precursor peptide of **10** was prepared via SPPS using Rink amide PS resin at a scale of 0.1 mmol initial resin capacity and purified by HPLC (solvent A: $H_2O$, acidified with 0.1% TFA; solvent B: ACN, acidified with 0.1% TFA). The obtained peptide was then dissolved in 10 mL of ACN and 4 equivalents DMP were added. The resulting suspension was shaken for one hour at RT and immediately purified by HPLC (solvent A: $H_2O$; solvent B: ACN) to give 29 mg (23 µmol) of **10** in a total yield of 23% (calculated in relation to initial resin capacity) as a white powder.

$^1$H NMR (400 MHz, DMSO) δ 8.21 (s, 1H), 7.84 (t, $J$ = 6.0 Hz, 1H), 7.69 (d, $J$ = 7.5 Hz, 1H), 7.44 (d, $J$ = 6.2 Hz, 1H), 7.32 (d, $J$ = 8.4 Hz, 1H), 7.29-7.16 (m, $J$ = 7.2, 6.8 Hz, 10H), 7.14 (d, $J$ = 6.7 Hz, 1H), 7.10 (d, $J$ = 10.1 Hz, 1H), 7.07-7.01 (m, 3H), 5.44 (dd, $J$ = 13.5, 9.6 Hz, 1H), 4.64-4.57 (m, 2H), 4.51 (dd, $J$ = 10.2, 5.1 Hz, 1H), 4.36 (dd, $J$ = 11.1, 7.7 Hz, 1H), 4.30 (t, $J$ = 7.5 Hz, 1H), 4.15 (dd, $J$ = 9.8, 7.5 Hz, 1H), 4.12-4.06 (m, 2H), 4.02 (dd, $J$ = 10.1, 6.9 Hz, 1H), 3.98-3.92 (m, 2H), 3.89-3.80 (m, 1H), 3.70 (dd, $J$ = 16.7, 6.5 Hz, 1H), 3.58-3.45 (m, 3H), 3.40-3.34 (m, 2H), 3.21-3.10 (m, 1H), 3.01 (dd, $J$ = 13.5, 8.8 Hz, 1H), 2.73-2.61 (m, 2H), 2.56-2.52 (m, 1H), 2.33 (h, $J$ = 6.7 Hz, 1H), 2.28-2.16 (m, 3H), 2.16-2.06 (m, 2H), 1.98-1.86 (m, 4H), 1.86-1.71 (m, 6H), 1.68-1.53 (m, 4H), 1.52-1.37 (m, 4H), 1.32-1.17 (m, 2H), 1.12-1.07 (m, 1H), 1.05 (d, $J$ = 6.8 Hz, 3H), 1.01 (d, $J$ = 6.8 Hz, 3H), 0.92 (d, $J$ = 6.5 Hz, 3H), 0.89-0.84 (m, 6H), 0.82 (d, $J$ = 6.5 Hz, 3H), 0.69 (d, $J$ = 6.9 Hz, 3H), 0.65 (t, $J$ = 7.7 Hz, 3H); $^{13}$C NMR (101 MHz, DMSO) δ 172.9, 172.5, 172.1, 172.0, 171.6, 171.5, 171.2, 171.0, 170.9, 169.1, 167.4, 142.3, 138.0, 136.1, 129.2, 129.0, 128.4, 128.1, 126.7, 126.4, 99.3, 63.6, 63.4, 63.0, 61.4, 60.6, 58.6, 57.2, 56.9, 52.7, 52.1, 48.6, 46.6, 42.2, 36.4, 35.5, 35.3, 29.9, 29.2, 27.0, 25.7, 25.5, 25.0, 24.4, 24.2, 22.7, 21.3, 18.9, 18.5, 15.6, 11.3, 10.9; HRMS (ESI): $m/z$ calculated for $[M+H]^+$: 1262.7296, found: 1262.7307 (0.9 ppm); calculated for $[M+2H]^{2+}$: 631.8685, found: 631.8688 (0.5 ppm); MS$^2$ HRMS (ESI, HCD): $m/z$ calculated for ring fragment $[M_{ring}+H]^+$: 865.4607, found: 865.4595 (-1.4 ppm).

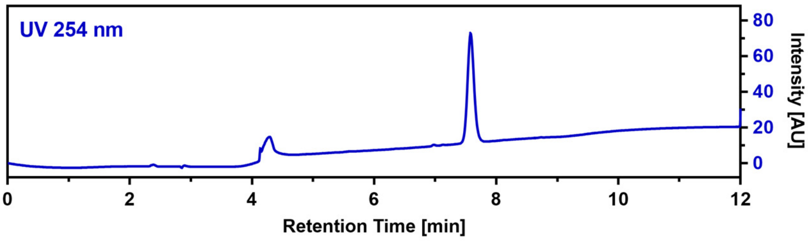

**Callyaerin K (11)**

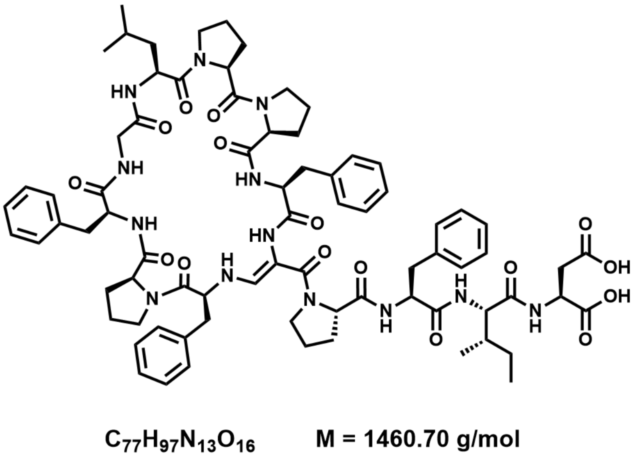

$C_{77}H_{97}N_{13}O_{16}$ M = 1460.70 g/mol

The linear precursor peptide of **11** was prepared via microwave-assisted SPPS using 2-CTC PS resin at a scale of 0.1 mmol initial resin capacity. After purification by HPLC (solvent A: $H_2O$, acidified with 0.1% TFA; solvent B: ACN, acidified with 0.1% TFA), 44 mg (30 µmol, 30%) of the peptide precursor was obtained as a white powder. For the oxidation of the serine residue and concomitant cyclization, the peptide was dissolved in 10 mL of ACN and 3 equivalents of DMP were added. The resulting suspension was shaken for 2.5 hours at RT and immediately purified by HPLC (solvent A: $H_2O$;

solvent B: ACN) to give 7.6 mg (5.2 µmol, 13.5%) of **11** in a total yield of 5.2% (calculated in relation to initial resin capacity) as a white powder.

$^{1}$H NMR (400 MHz, DMSO) δ 9.26 (d, *J* = 8.4 Hz, 1H), 8.94 (s, 1H), 8.33 (d, *J* = 7.5 Hz, 1H), 7.81 (d, *J* = 9.4 Hz, 1H), 7.47 (s, 1H), 7.41-7.34 (m, 4H), 7.30-7.23 (m, 8H), 7.23-7.16 (m, 6H), 7.12-7.08 (m, 2H), 7.07-7.03 (m, 2H), 6.98 (s, 1H), 5.64-5.48 (m, 1H), 5.18-5.07 (m, 1H), 4.61 (s, 1H), 4.53-4.45 (m, 1H), 4.32 (ddd, *J* = 12.7, 9.3, 3.6 Hz, 1H), 4.22-4.12 (m, 2H), 4.10-3.97 (m, 4H), 3.96-3.86 (m, 2H), 3.66 (s, 1H), 3.56-3.44 (m, 3H), 3.29-3.21 (m, 4H), 3.21-3.14 (m, 2H), 3.10-3.04 (m, 1H), 2.99-2.91 (m, 2H), 2.90-2.82 (m, 3H), 2.73-2.63 (m, 2H), 2.58 (dd, *J* = 16.4, 6.1 Hz, 1H), 2.21-2.11 (m, 1H), 2.07-1.99 (m, 1H), 1.97-1.73 (m, 6H), 1.68-1.52 (m, 4H), 1.50-1.27 (m, 8H), 1.23 (s, 1H), 1.14-1.05 (m, 1H), 0.88 (d, *J* = 6.5 Hz, 3H), 0.86-0.81 (m, 6H), 0.70 (t, *J* = 7.3 Hz, 3H), 0.65-0.53 (m, 1H); HRMS (ESI): *m/z* calculated for $[M+H]^{+}$: 1460.7249, found: 1460.7254 (0.3 ppm); calculated for $[M+2H]^{2+}$: 730.8661, found: 730.8662 (0.1 ppm); $MS^{2}$ HRMS (ESI, HCD): *m/z* calculated for ring fragment $[M_{ring}+H]^{+}$: 970.4822, found: 940.4814 (-0.8 ppm).

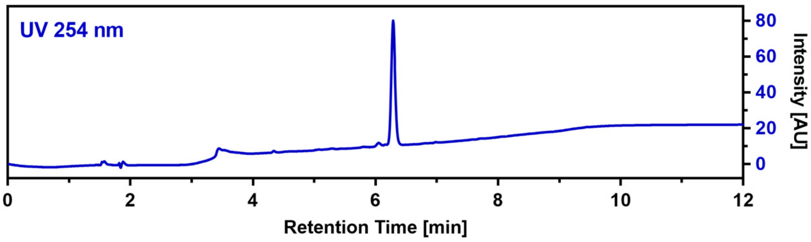

**Callyaerin L (12)**

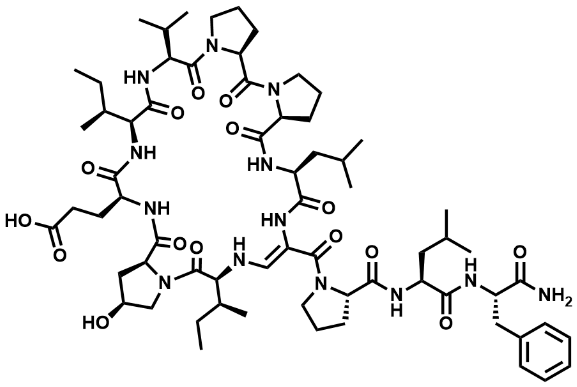

$C_{66}H_{101}N_{13}O_{15}$ M = 1316.61 g/mol

The linear precursor peptide of **12** was prepared via microwave-assisted SPPS using Rink amide PS resin at a scale of 0.1 mmol initial resin capacity. After purification by HPLC (solvent A: $H_2O$, acidified with 0.1% TFA; solvent B: ACN, acidified with 0.1% TFA), 43 mg (32 µmol, 32%) of the peptide precursor was obtained as a white powder. For the oxidation of the serine residue and concomitant cyclization, the peptide was dissolved in 8 mL of ACN and 3 equivalents of DMP were added. The resulting suspension was shaken for 2 hours at RT and immediately purified by HPLC (solvent A: $H_2O$; solvent B: ACN) to give 3.7 mg (2.8 µmol, 8.7%) of **12** in a total yield of 2.8% (calculated in relation to initial resin capacity) as a white powder.

$^{1}$H NMR (400 MHz, DMSO) δ 8.75 (s, 1H), 8.27 (s, 1H), 7.76 (d, *J* = 7.6 Hz, 1H), 7.59-7.51 (m, 2H), 7.37 (d, *J* = 7.4 Hz, 1H), 7.32-7.26 (m, 1H), 7.25-7.18 (m, 3H), 7.14-7.04 (m, 3H), 6.91 (d, *J* = 9.9 Hz, 1H), 6.78 (s, 1H), 5.45-5.36 (m, 1H), 5.36-5.32 (m, 1H), 4.61-4.52 (m, 2H), 4.40 (s, 1H), 4.30-4.23 (m, 2H), 4.22-4.03 (m, 4H), 3.99-3.90 (m, 2H), 3.88-3.77 (m, 2H), 3.75-3.66 (m, 2H), 3.63-3.48 (m, 4H), 3.22-3.10 (m, 2H), 2.74-2.66 (m, 1H), 2.33-2.23 (m, 3H), 2.22-2.17 (m, 2H), 2.15-2.02 (m, 6H), 1.98-1.80 (m, 7H), 1.77-1.57 (m, 5H), 1.56-1.46 (m, 3H), 1.42-1.30 (m, 3H), 1.23- (s, 1H), 1.19-1.11 (m, 1H), 0.98 (d, *J* = 6.9 Hz, 3H), 0.96-0.91 (m, 6H), 0.88-0.79 (m, 9H), 0.76-0.68 (m, 9H), 0.15 (t, *J* = 7.3 Hz, 3H); HRMS (ESI): *m/z* calculated for $[M+H]^{+}$: 1316.7613, found: 1316.7621 (0.6 ppm); calculated for $[M+2H]^{2+}$: 658.8843, found: 658.884 (0.2 ppm); $MS^{2}$ HRMS (ESI, HCD): *m/z* calculated for ring fragment $[M_{ring}+H]^{+}$: 942.5295, found: 942.5301 (0.6 ppm).

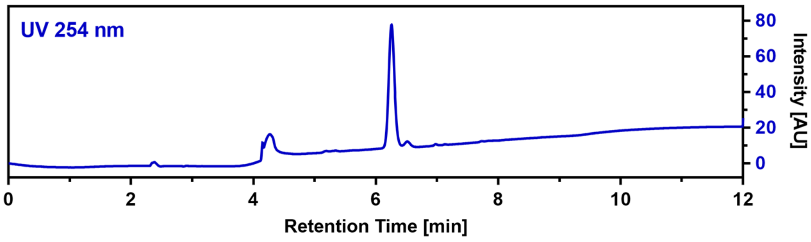

**Callynormine A (13)**

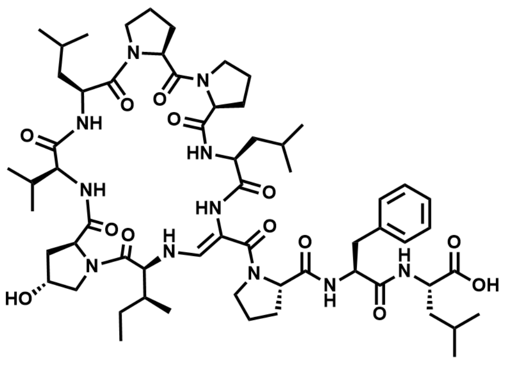

$C_{61}H_{93}N_{11}O_{13}$ M = 1188.48 g/mol

The linear precursor peptide of **13** was prepared via microwave-assisted SPPS using Rink amide PS resin at a scale of 0.1 mmol initial resin capacity. After purification by HPLC (solvent A: $H_2O$, acidified with 0.1% TFA; solvent B: ACN, acidified with 0.1% TFA), 55 mg (46 µmol, 46%) of the peptide precursor was obtained as a white powder. For the oxidation of the serine residue and concomitant cyclization, the peptide was dissolved in 10 mL of ACN and 3 equivalents of DMP were added. The resulting suspension was shaken for 2 hours at RT and immediately purified by HPLC (solvent A: $H_2O$; solvent B: ACN) to give 17.7 mg (14.9 µmol, 32.7%) of **13** in a total yield of 14.9% (calculated in relation to initial resin capacity) as a white powder.

$^1$H NMR (400 MHz, DMSO) δ 8.24 (s, 1H), 7.75 (d, *J* = 9.4 Hz, 1H), 7.56 (d, *J* = 3.8 Hz, 1H), 7.54 (s, 1H), 7.37-7.26 (m, 5H), 7.20 (d, *J* = 5.9 Hz, 1H), 7.19-7.17 (m, 1H), 7.05 (d, *J* = 13.7 Hz, 1H), 5.24 (dd, *J* = 13.7, 9.4 Hz, 1H), 5.17 (d, *J* = 3.7 Hz, 1H), 4.72-4.63 (m, 1H), 4.54 (q, *J* = 6.9 Hz, 1H), 4.44-4.36 (m, 3H), 4.35-4.26 (m, 3H), 4.19-4.09 (m, 2H), 3.98-3.88 (m, 2H), 3.69-3.58 (m, 3H), 3.54-3.43 (m, 3H), 3.29-3.16 (m, 3H), 2.77 (t, *J* = 12.8 Hz, 1H), 2.35-2.26 (m, 1H), 2.23-2.10 (m, 2H), 2.09-1.95 (m, 5H), 1.94-1.79 (m, 6H), 1.74-1.57 (m, 4H), 1.56-1.43 (m, 6H), 1.41-1.33 (m, 1H), 1.19-1.08 (m, 1H), 0.96-0.75 (m, 30H); HRMS (ESI): *m/z* calculated for $[M+H]^+$: 1188.7027, found: 1188.7022 (-0.4 ppm); calculated for $[M+2H]^{2+}$: 594.8550, found: 594.8534 (-2.7 ppm); $MS^2$ HRMS (ESI, HCD): *m/z* calculated for ring fragment $[M_{ring}+H]^+$: 813.4870, found: 813.4849 (-2.6 ppm).

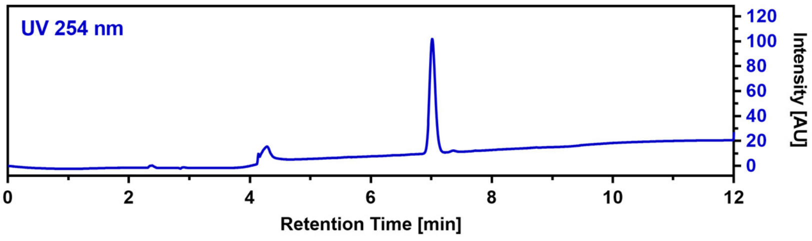

**CalA_R1A (14)**

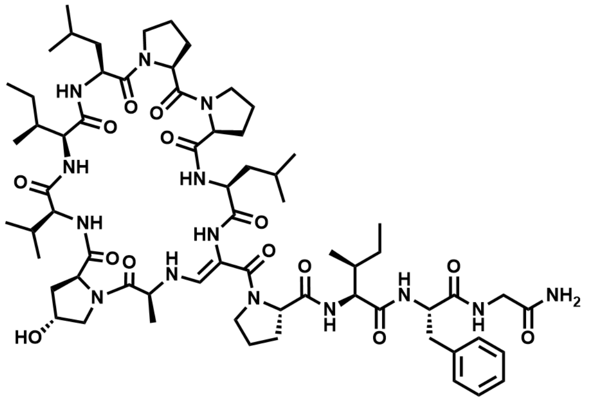

$C_{66}H_{102}N_{14}O_{14}$ M = 1315.63 g/mol

The linear precursor peptide of **14** was prepared via SPPS using Rink amide PS resin at a scale of 0.23 mmol initial resin capacity. After purification by HPLC (solvent A: $H_2O$, acidified with 0.1% TFA; solvent B: ACN, acidified with 0.1% TFA), 102 mg (77 µmol, 33%) of the peptide precursor was obtained as a white powder. For the oxidation of the serine residue and concomitant cyclization, the peptide was dissolved in 10 mL of ACN and 4 equivalents of DMP were added. The resulting suspension was shaken for one hour at RT and immediately purified by HPLC (solvent A: $H_2O$; solvent B: ACN) to give 8.0 mg (6.1 µmol, 7.9%) of **14** in a total yield of 2.7% (calculated in relation to initial resin capacity) as a white powder.

$^1$H NMR (400 MHz, DMSO) δ 8.36 (s, 1H), 8.25 (t, *J* = 6.2 Hz, 1H), 7.74 (d, *J* = 6.6 Hz, 1H), 7.59 (d, *J* = 13.6 Hz, 1H), 7.56 (d, *J* = 9.1 Hz, 1H), 7.35 (d, *J* = 7.8 Hz, 1H), 7.28 (s, 1H), 7.25-7.18 (m, 1H), 7.18-7.11 (m, 6H), 6.96 (s, 1H), 6.87 (d, *J* = 10.0 Hz, 1H), 5.77-5.67 (m, 1H), 5.33 (d, *J* = 3.3 Hz, 1H), 4.72-4.65 (m, 1H), 4.63-4.53 (m, 2H), 4.39 (s, 1H), 4.30-4.23 (m, 3H), 4.12 (dd, *J* = 10.0, 7.2 Hz, 1H), 4.06-3.95 (m, 2H), 3.94-3.84 (m, 3H), 3.67 (d, *J* = 11.3 Hz, 1H), 3.63-3.52 (m, 3H), 3.51-3.39 (m, 3H), 3.29-3.17 (m, 2H), 3.11 (dd, *J* = 14.2, 3.4 Hz, 1H), 2.70-2.61 (m, 1H), 2.30-2.21 (m, 3H), 2.17-2.10 (m, 1H), 2.06-1.91 (m, 4H), 1.90-1.80 (m, 4H), 1.76-1.66 (m, 3H),

1.64-1.42 (m, 8H), 1.39-1.30 (m, 1H), 1.20-1.12 (m, 1H), 1.09-1.01 (m, 2H), 0.99 (d, *J* = 6.5 Hz, 3H), 0.94-0.90 (m, 6H), 0.89-0.87 (m, 3H), 0.86-0.82 (m, 9H), 0.79 (d, *J* = 6.8 Hz, 3H), 0.76 (t, *J* = 7.4 Hz, 3H), 0.69 (t, *J* = 7.4 Hz, 3H), 0.50 (d, *J* = 6.8 Hz, 3H); HRMS (ESI): *m/z* calculated for $[M+H]^+$: 1315.7773, found: 1315.7745 (-2.1 ppm); calculated for $[M+2H]^{2+}$: 658.3923, found: 658.3919 (-0.6 ppm); $MS^2$ HRMS (ESI, HCD): *m/z* calculated for ring fragment $[M_{ring}+H]^+$: 884.5241, found: 884.5244 (0.3 ppm).

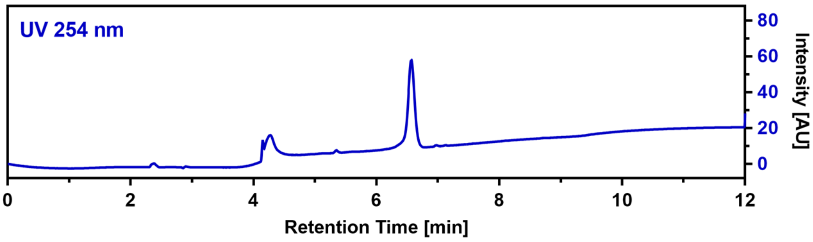

**CalA_R2A (15)**

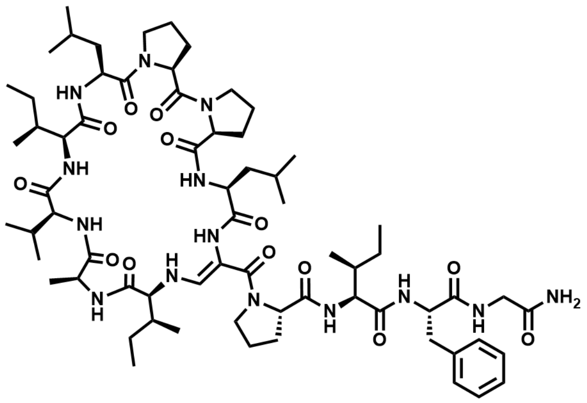

$C_{67}H_{105}N_{14}O_{13}$    M = 1314.67 g/mol

The linear precursor peptide of **15** was prepared via SPPS using Rink amide PS resin at a scale of 0.1 mmol initial resin capacity and purified by HPLC (solvent A: $H_2O$, acidified with 0.1% TFA; solvent B: ACN, acidified with 0.1% TFA). The obtained peptide was then dissolved in 10 mL of ACN and 4 equivalents DMP were added. The resulting suspension was shaken for one hour at RT and immediately purified by HPLC (solvent A: $H_2O$; solvent B: ACN) to give 3.2 mg (2.4 µmol) of **15** in a total yield of 2.4% (calculated in relation to initial resin capacity) as a white powder.

$^1H$ NMR (400 MHz, DMSO) δ 8.77 (s, 1H), 8.47 (d, *J* = 6.7 Hz, 1H), 8.15 (s, 1H), 7.70 (d, *J* = 6.9 Hz, 1H), 7.64 (t, *J* = 6.2 Hz, 1H), 7.57-7.51 (m, 2H), 7.21-7.07 (m, 9H), 6.74 (d, *J* = 9.6 Hz, 1H), 5.82-5.71 (m, 1H), 4.63-4.55 (m, 1H), 4.54-4.47 (m, 1H), 4.31-4.22 (m, 3H), 3.94 (t, *J* = 8.8 Hz, 1H), 3.86-3.77 (m, 3H), 3.74 (t, *J* = 7.3 Hz, 1H), 3.63-3.55 (m, 3H), 3.54-3.45 (m, 1H), 3.43-3.38 (m, 1H), 3.30-3.22 (m, 3H), 3.01 (dd, *J* = 10.6, 6.9 Hz, 1H), 2.69-2.60 (m, 2H), 2.32-2.20 (m, 3H), 2.00-1.94 (m, 1H), 1.93-1.82 (m, 7H), 1.79-1.68 (m, 4H), 1.65-1.52 (m, 4H), 1.47-1.35 (m, 4H), 1.28 (d, *J* = 7.0 Hz, 3H), 1.23 (s, 1H), 1.16-1.12 (m, 1H), 1.11 (s, 2H), 0.95 (d, *J* = 6.4 Hz, 3H), 0.92-0.78 (m, 21H), 0.74-0.69 (m, 6H), 0.45 (d, *J* = 6.8 Hz, 3H), 0.43-0.38 (m, 3H); HRMS (ESI): *m/z* calculated for $[M+H]^+$: 1315.8137, found: 1315.8145 (0.6 ppm); calculated for $[M+2H]^{2+}$: 658.4105, found: 658.4105 (0 ppm); $MS^2$ HRMS (ESI, HCD): *m/z* calculated for ring fragment $[M_{ring}+H]^+$: 884.5605, found: 884.5591 (-1.6 ppm).

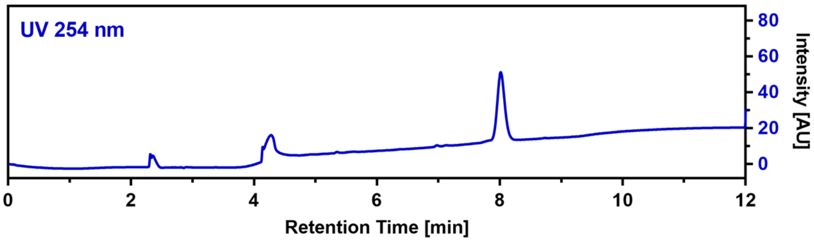

**CalA_R3A (16)**

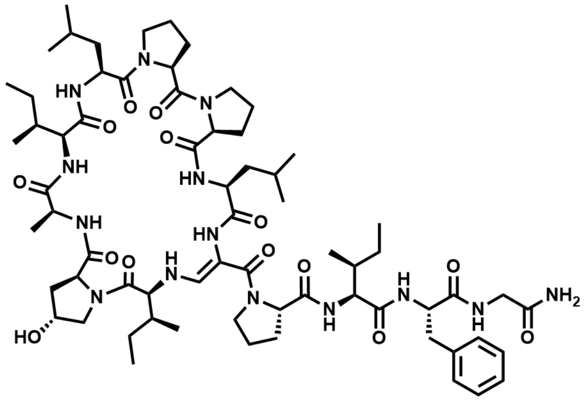

$C_{67}H_{104}N_{14}O_{14}$ M = 1329.65 g/mol

The linear precursor peptide of **16** was prepared via SPPS using Rink amide PS resin at a scale of 0.1 mmol initial resin capacity and purified by HPLC (solvent A: $H_2O$, acidified with 0.1% TFA; solvent B: ACN, acidified with 0.1% TFA). The obtained peptide was then dissolved in 10 mL of ACN and 4 equivalents DMP were added. The resulting suspension was shaken for one hour at RT and immediately purified by HPLC (solvent A: $H_2O$; solvent B: ACN) to give 8.3 mg (6.2 µmol) of **16** in a total yield of 6.2% (calculated in relation to initial resin capacity) as a white powder.

HRMS (ESI): *m/z* calculated for $[M+H]^+$: 1329.7929, found: 1329.7947 (1.4 ppm); calculated for $[M+2H]^{2+}$: 665.4001, found: 665.4004 (0.5 ppm); $MS^2$ HRMS (ESI, HCD): *m/z* calculated for ring fragment $[M_{ring}+H]^+$: 898.5397, found: 898.5389 (-0.9 ppm).

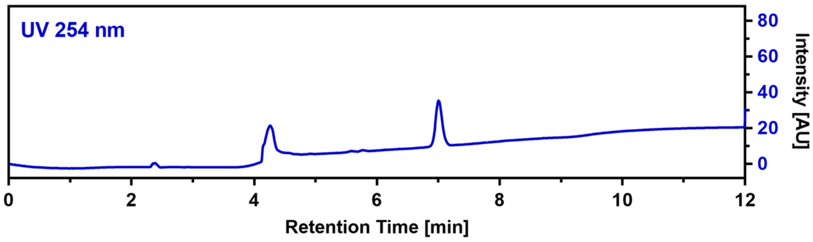

**CalA_R4A (17)**

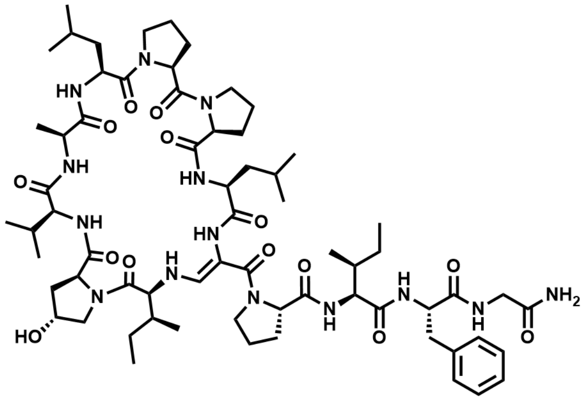

$C_{66}H_{102}N_{14}O_{14}$ M = 1315.63 g/mol

The linear precursor peptide of **17** was prepared via SPPS using Rink amide PS resin at a scale of 0.1 mmol initial resin capacity and purified by HPLC (solvent A: $H_2O$, acidified with 0.1% TFA; solvent B: ACN, acidified with 0.1% TFA). The obtained peptide was then dissolved in 10 mL of ACN and 4 equivalents DMP were added. The resulting suspension was shaken for one hour at RT and immediately purified by HPLC (solvent A: $H_2O$; solvent B: ACN) to give 21.5 mg (16.3 µmol) of **17** in a total yield of 16.3% (calculated in relation to initial resin capacity) as a white powder.

$^1$H NMR (400 MHz, DMSO) δ 8.91 (s, 1H), 8.68 (s, 1H), 8.30 (s, 1H), 8.20-8.12 (m, 1H), 7.50 (d, *J* = 6.9 Hz, 1H), 7.46 (d, *J* = 8.8 Hz, 1H), 7.36 (d, *J* = 13.4 Hz, 1H), 7.28-7.09 (m, 7H), 6.99-6.94 (m, 1H), 6.74 (d, *J* = 10.0 Hz, 1H), 5.84 (dd, *J* = 13.2, 10.2 Hz, 1H), 5.35 (d, *J* = 3.1 Hz, 1H), 4.65-4.57 (m, 1H), 4.57-4.52 (m, 1H), 4.41 (s, 1H), 4.31-4.21 (m, 4H), 4.12-4.02 (m, 2H), 3.97 (dd, *J* = 7.3, 5.8 Hz, 1H), 3.91 (dd, *J* = 16.5, 7.1 Hz, 1H), 3.79 (t, *J* = 8.5 Hz, 1H), 3.76-3.70 (m, 1H), 3.69-3.61 (m, 1H), 3.62-3.55 (m, 1H), 3.54-3.43 (m, 3H), 3.38-3.33 (m, 1H), 3.25-3.17 (m, 1H), 3.10 (dd, *J* = 13.9, 2.9 Hz, 1H), 3.04-2.97 (m, 1H), 2.68-2.55 (m, 2H), 2.33-2.20 (m, 3H), 2.11-2.05 (m, 1H), 2.02-1.95 (m, 1H), 1.94-1.80 (m, 7H), 1.78-1.63 (m, 5H), 1.63-1.51 (m, 4H), 1.50-1.35 (m, 4H), 1.31-1.22 (m, 1H), 1.14 (d, *J* = 7.3 Hz, 3H), 1.08-0.99 (m, 1H), 0.95 (d, *J* = 6.5 Hz, 3H), 0.89-0.77 (m, 18H), 0.70 (t, *J* = 7.4 Hz, 3H), 0.44 (t,

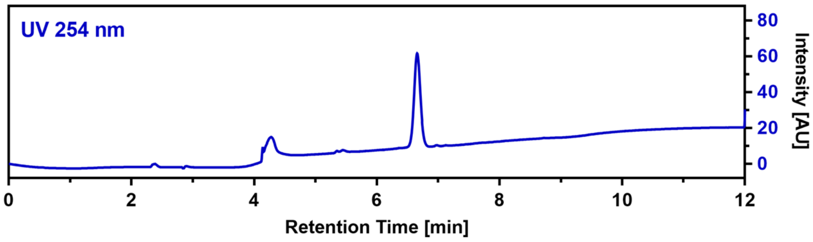

$J$ = 7.4 Hz, 3H), 0.38 (d, $J$ = 6.8 Hz, 3H); HRMS (ESI): $m/z$ calculated for $[M+H]^+$: 1315.7773, found: 1315.7780 (0.5 ppm); calculated for $[M+2H]^{2+}$: 658.3923, found: 658.3925 (0.3 ppm); $MS^2$ HRMS (ESI, HCD): $m/z$ calculated for ring fragment $[M_{ring}+H]^+$: 884.5241, found: 884.5243 (0.2 ppm).

**CalA_R5A (18)**

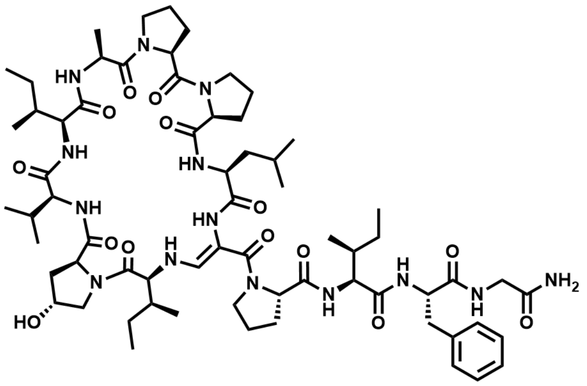

$C_{66}H_{102}N_{14}O_{14}$ M = 1315.63 g/mol

The linear precursor peptide of **18** was prepared via SPPS using Rink amide PS resin at a scale of 0.1 mmol initial resin capacity and purified by HPLC (solvent A: $H_2O$, acidified with 0.1% TFA; solvent B: ACN, acidified with 0.1% TFA). The obtained peptide was then dissolved in 10 mL of ACN and 4 equivalents DMP were added. The resulting suspension was shaken for one hour at RT and immediately purified by HPLC (solvent A: $H_2O$; solvent B: ACN) to give 34.3 mg (26.1 µmol) of **18** in a total yield of 26.1% (calculated in relation to initial resin capacity) as a white powder.

$^1H$ NMR (400 MHz, DMSO) δ 9.02 (d, $J$ = 7.0 Hz, 1H), 8.84 (s, 1H), 8.30 (s, 1H), 8.08 (t, $J$ = 6.2 Hz, 1H), 7.54 (d, $J$ = 8.4 Hz, 1H), 7.49 (d, $J$ = 6.6 Hz, 1H), 7.32 (d, $J$ = 13.4 Hz, 1H), 7.28 (d, $J$ = 8.9 Hz, 1H), 7.25-7.10 (m, 6H), 7.01-6.94 (m, 2H), 5.85 (dd, $J$ = 13.2, 10.2 Hz, 1H), 5.35 (d, $J$ = 3.1 Hz, 1H), 4.69 (p, $J$ = 6.4 Hz, 1H), 4.47-4.38 (m, 2H), 4.30-4.20 (m, 4H), 4.09-4.01 (m, 2H), 3.89 (dd, $J$ = 16.5, 7.0 Hz, 1H), 3.83-3.75 (m, 1H), 3.74-3.62 (m, 4H), 3.57-3.49 (m, 2H), 3.48-3.42 (m, 2H), 3.31-3.21 (m, 1H), 3.11 (dd, $J$ = 13.8, 3.0 Hz, 1H), 2.96 (dd, $J$ = 10.9, 6.9 Hz, 1H), 2.69-2.57 (m, 2H), 2.31-2.16 (m, 3H), 2.13-2.05 (m, 1H), 2.04-1.98 (m, 1H), 1.96-1.81 (m, 6H), 1.80-1.59 (m, 6H), 1.54-1.42 (m, 2H), 1.42-1.30 (m, 3H), 1.28-1.19 (m, 1H), 1.16 (d, $J$ = 6.2 Hz, 3H), 1.13-1.07 (m, 1H), 1.06-0.96 (m, 1H), 0.90 (d, $J$ = 6.2 Hz, 3H), 0.85-0.70 (m, 23H), 0.38 (t, $J$ = 7.3 Hz, 3H), 0.35 (d, $J$ = 6.8 Hz, 3H); $^{13}C$ NMR (101 MHz, DMSO) δ 173.5, 173.0, 172.9, 172.1, 172.0, 171.8, 171.7, 171.5, 171.2, 171.2, 170.9, 167.8, 143.2, 138.1, 129.2, 127.9, 126.3, 98.4, 68.9, 64.1, 62.5, 61.9, 58.9, 56.7, 55.7, 50.7, 48.9, 47.2, 46.1, 42.1, 40.5, 38.4, 37.6, 36.2, 35.0, 29.5, 28.6, 27.2, 25.9, 25.6, 25.1, 24.2, 23.0, 20.7, 19.5, 19.2, 16.2, 15.1, 14.9, 14.3, 11.0, 10.7, 10.4; HRMS (ESI): $m/z$ calculated for $[M+H]^+$: 1315.7773, found: 1315.7791 (1.4 ppm); calculated for $[M+2H]^{2+}$: 658.3923, found: 658.3927 (0.6 ppm); $MS^2$ HRMS (ESI, HCD): $m/z$ calculated for ring fragment $[M_{ring}+H]^+$: 884.5241, found: 884.5246 (0.6 ppm).

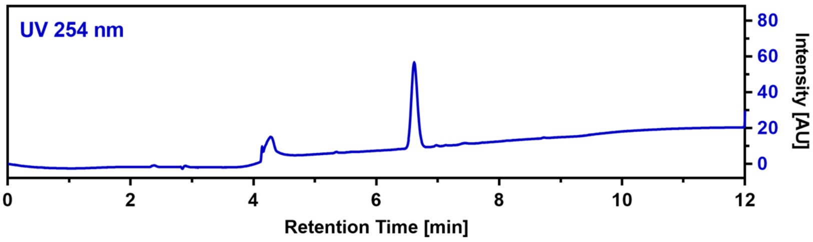

##### CalA_R6A (19)

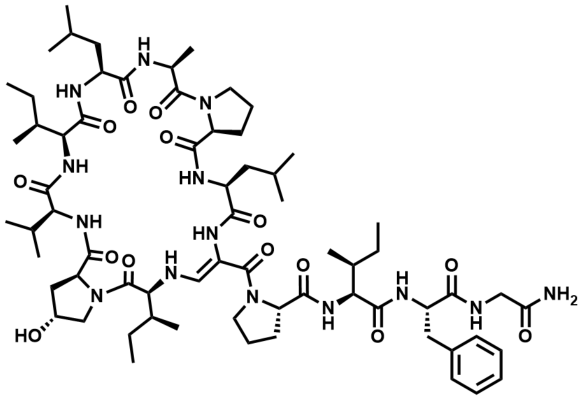

$C_{67}H_{106}N_{14}O_{14}$ M = 1331.67 g/mol

The linear precursor peptide of **19** was prepared via SPPS using Rink amide PS resin at a scale of 0.1 mmol initial resin capacity and purified by HPLC (solvent A: $H_2O$, acidified with 0.1% TFA; solvent B: ACN, acidified with 0.1% TFA). The obtained peptide was then dissolved in 10 mL of ACN and 4 equivalents DMP were added. The resulting suspension was shaken for one hour at RT and immediately purified by HPLC (solvent A: $H_2O$; solvent B: ACN) to give 12.8 mg (9.6 µmol) of **19** in a total yield of 9.6% (calculated in relation to initial resin capacity) as a white powder.
HRMS (ESI): *m/z* calculated for $[M+H]^+$: 1331.8086, found: 1331.8096 (0.8 ppm); calculated for $[M+2H]^{2+}$: 666.4079, found: 666.4066 (-2.0 ppm); MS$^2$ HRMS (ESI, HCD): *m/z* calculated for ring fragment $[M_{ring}+H]^+$: 900.5554, found: 900.5546 (-0.9 ppm).

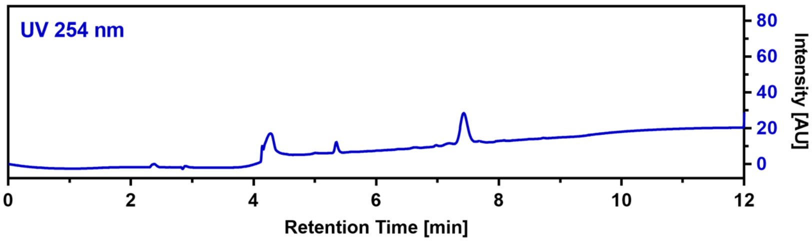

##### CalA_R7A (20)

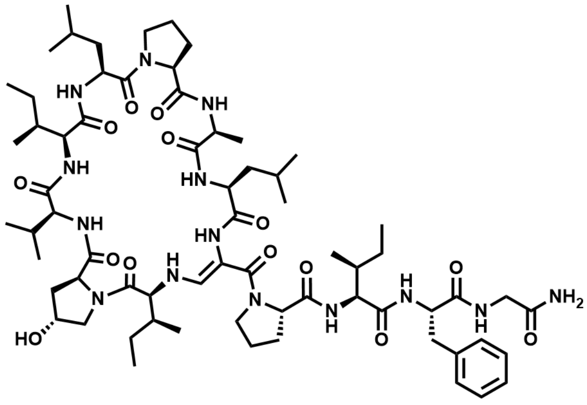

$C_{67}H_{106}N_{14}O_{14}$ M = 1331.67 g/mol

The linear precursor peptide of **20** was prepared via SPPS using Rink amide PS resin at a scale of 0.1 mmol initial resin capacity and purified by HPLC (solvent A: $H_2O$, acidified with 0.1% TFA; solvent B: ACN, acidified with 0.1% TFA). The obtained peptide was then dissolved in 10 mL of ACN and 4 equivalents DMP were added. The resulting suspension was shaken for one hour at RT and immediately purified by HPLC (solvent A: $H_2O$; solvent B: ACN) to give 8.1 mg (6.1 µmol) of **20** in a total yield of 6.1% (calculated in relation to initial resin capacity) as a white powder.
HRMS (ESI): *m/z* calculated for $[M+H]^+$: 1331.8086, found: 1331.8097 (0.8 ppm); calculated for $[M+2H]^{2+}$: 666.4079, found: 666.4081 (0.3 ppm); MS$^2$ HRMS (ESI, HCD): *m/z* calculated for ring fragment $[M_{ring}+H]^+$: 900.5554, found: 900.5558 (0.4 ppm).

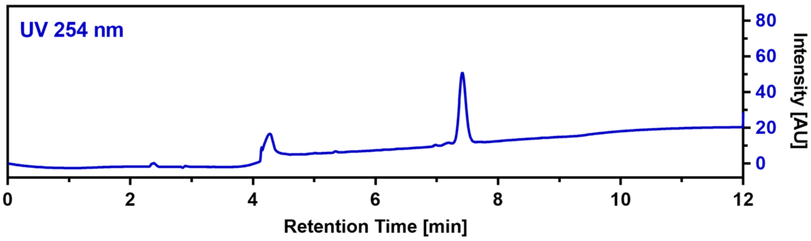

**CalA_R8A (21)**

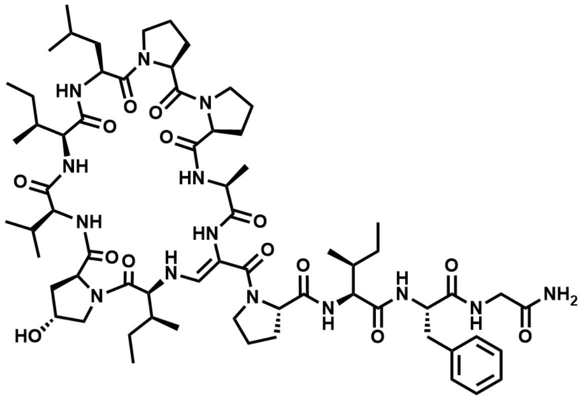

$C_{66}H_{102}N_{14}O_{14}$ M = 1315.63 g/mol

The linear precursor peptide of **21** was prepared via SPPS using Rink amide PS resin at a scale of 0.1 mmol initial resin capacity and purified by HPLC (solvent A: $H_2O$, acidified with 0.1% TFA; solvent B: ACN, acidified with 0.1% TFA). The obtained peptide was then dissolved in 10 mL of ACN and 4 equivalents DMP were added. The resulting suspension was shaken for one hour at RT and immediately purified by HPLC (solvent A: $H_2O$; solvent B: ACN) to give 11.0 mg (8.4 µmol) of **21** in a total yield of 8.4% (calculated in relation to initial resin capacity) as a white powder.

$^1$H NMR (400 MHz, DMSO) δ 8.99 (d, $J$ = 6.6 Hz, 1H), 8.81 (s, 1H), 8.32 (s, 1H), 8.11 (dd, $J$ = 7.0, 5.3 Hz, 1H), 7.48 (d, $J$ = 8.5 Hz, 1H), 7.44 (d, $J$ = 6.4 Hz, 1H), 7.35 (d, $J$ = 13.4 Hz, 1H), 7.27-7.12 (m, 7H), 6.99-6.95 (m, 1H), 6.89 (d, $J$ = 9.5 Hz, 1H), 5.82 (dd, $J$ = 13.3, 10.1 Hz, 1H), 5.36 (s, 1H), 4.55-4.46 (m, 2H), 4.41 (s, 1H), 4.30-4.21 (m, 4H), 4.10-4.02 (m, 2H), 3.90 (dd, $J$ = 16.6, 7.1 Hz, 1H), 3.79-3.68 (m, 4H), 3.65-3.57 (m, 2H), 3.55-3.46 (m, 2H), 3.30-3.21 (m, 2H), 3.10 (dd, $J$ = 13.8, 3.0 Hz, 1H), 2.99 (dd, $J$ = 10.8, 6.9 Hz, 1H), 2.68-2.59 (m, 2H), 2.33-2.17 (m, 3H), 2.12-2.04 (m, 1H), 2.03-1.97 (m, 1H), 1.96-1.79 (m, 7H), 1.75-1.62 (m, 3H), 1.57-1.48 (m, 3H), 1.45 (d, $J$ = 7.4 Hz, 3H), 1.42-1.33 (m, 3H), 1.27-1.10 (m, 4H), 1.09-0.97 (m, 1H), 0.92 (d, $J$ = 6.5 Hz, 3H), 0.89-0.85 (m, 1H), 0.84-0.75 (m, 18H), 0.72 (t, $J$ = 7.4 Hz, 3H), 0.41 (t, $J$ = 7.3 Hz, 3H), 0.36 (d, $J$ = 6.7 Hz, 3H); HRMS (ESI): *m/z* calculated for $[M+H]^+$: 1315.7773, found: 1315.7780 (0.5 ppm); calculated for $[M+2H]^{2+}$: 658.3923, found: 658.3925 (0.3 ppm); MS$^2$ HRMS (ESI, HCD): *m/z* calculated for ring fragment $[M_{ring}+H]^+$: 884.5241, found: 884.5244 (0.3 ppm).

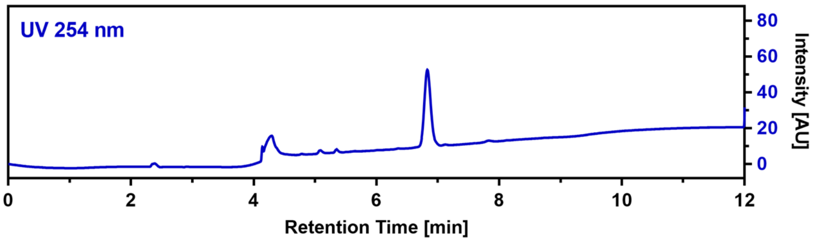

**CalA_C1A (22)**

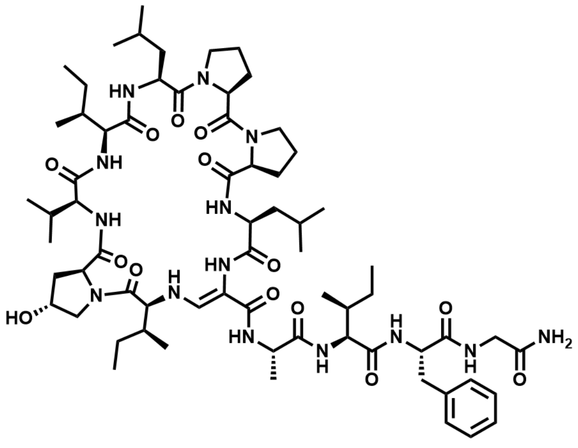

$C_{67}H_{106}N_{14}O_{14}$ M = 1331.67 g/mol

The linear precursor peptide of **22** was prepared via SPPS using Rink amide PS resin at a scale of 0.23 mmol initial resin capacity. After purification by HPLC (solvent A: $H_2O$, acidified with 0.1% TFA; solvent B: ACN, acidified with 0.1% TFA), 93 mg (69 µmol, 30%) of the peptide precursor was obtained as a white powder. For the oxidation of the serine residue and concomitant cyclization, the peptide was dissolved in 10 mL of ACN and 4 equivalents of DMP were added. The resulting suspension was shaken for one hour at RT and immediately purified by HPLC (solvent A: $H_2O$; solvent B: ACN) to give 2.4 mg (1.8 µmol, 2.6%) of **22** in a total yield of 0.8% (calculated in relation to initial resin capacity) as a white powder.

HRMS (ESI): *m/z* calculated for $[M+H]^+$: 1331.8086, found: 1331.8094 (0.6 ppm); calculated for $[M+2H]^{2+}$: 666.4079, found: 666.4071 (-1.2 ppm); $MS^2$ HRMS (ESI, HCD): *m/z* calculated for ring fragment $[M_{ring}+H]^+$: 926.5710, found: 926.5729 (2.1 ppm).

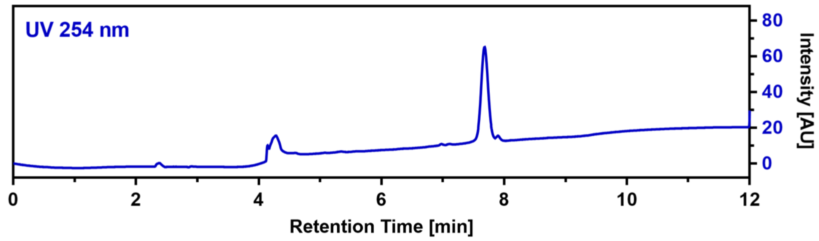

**CalA_C2A (23)**

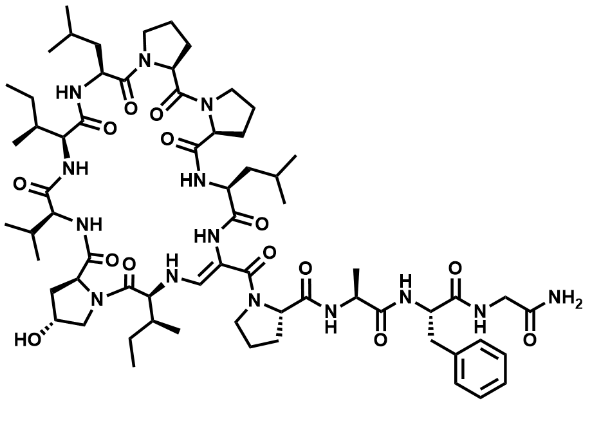

$C_{66}H_{102}N_{14}O_{14}$ M = 1315.63 g/mol

The linear precursor peptide of **23** was prepared via SPPS using Rink amide PS resin at a scale of 0.1 mmol initial resin capacity and purified by HPLC (solvent A: $H_2O$, acidified with 0.1% TFA; solvent B: ACN, acidified with 0.1% TFA). The obtained peptide was then dissolved in 10 mL of ACN and 4 equivalents DMP were added. The resulting suspension was shaken for one hour at RT and immediately purified by HPLC (solvent A: $H_2O$; solvent B: ACN) to give 3.4 mg (2.6 µmol) of **23** in a total yield of 2.6% (calculated in relation to initial resin capacity) as a white powder.

$^1H$ NMR (400 MHz, DMSO) δ 8.99 (s, 1H), 8.77 (s, 1H), 8.17 (s, 1H), 7.87 (dd, *J* = 7.1, 5.2 Hz, 1H), 7.81 (d, *J* = 7.4 Hz, 1H), 7.58 (d, *J* = 9.2 Hz, 1H), 7.36 (d, *J* = 7.1 Hz, 1H), 7.27 (d, *J* = 13.3 Hz, 1H), 7.23-7.19 (m, 3H), 7.11-7.08 (m, 3H), 7.03 (s, 1H), 6.76 (d, *J* = 9.8 Hz, 1H), 5.84-5.73 (m, 1H), 4.63 (q, *J* = 6.6 Hz, 1H), 4.56 (dd, *J* = 10.0, 5.0 Hz, 1H), 4.40 (s, 1H), 4.29-4.21 (m, 4H), 4.08-4.01 (m, 1H), 3.97-3.91 (m, 2H), 3.87 (dd, *J* = 16.8, 7.2 Hz, 1H), 3.83-3.76 (m, 2H), 3.71-3.61 (m, 3H), 3.52-3.45 (m, 2H), 3.36-3.30 (m, 1H), 3.28-3.19 (m, 1H), 3.16-3.09 (m, 1H), 3.03-2.96 (m, 1H), 2.75 (t, *J* = 12.7 Hz, 1H), 2.69-2.59 (m, 1H), 2.34 – 2.18 (m, 3H), 2.13 – 2.05 (m, 1H), 2.02 – 1.96 (m, 1H), 1.95 – 1.81 (m, 8H), 1.78 – 1.67 (m, 3H), 1.66-1.52 (m, 4H), 1.50-1.43 (m, 1H), 1.42-1.31 (m, 2H), 1.30-1.25 (m, 1H), 1.23 (s, 1H), 1.17-1.10 (m, 1H), 1.08 (d, *J* = 7.4 Hz, 3H), 0.95 (d, *J* = 6.5 Hz, 3H), 0.90 (d, *J* = 6.4 Hz, 3H), 0.87 (d, *J* = 2.9 Hz, 3H), 0.86 (d, *J* = 3.0 Hz, 3H), 0.84-0.79 (m, 9H), 0.77 (t, *J* = 7.4 Hz, 3H), 0.74 (d, *J* = 7.2 Hz, 3H), 0.62-0.52 (m, 1H), 0.27 (t, *J* = 7.3 Hz, 3H); HRMS (ESI): *m/z* calculated for $[M+H]^+$: 1315.7773, found: 1315.7785 (0.9 ppm); calculated for $[M+2H]^{2+}$: 658.3923, found: 658.3925 (0.3 ppm); $MS^2$ HRMS (ESI, HCD): *m/z* calculated for ring fragment $[M_{ring}+H]^+$: 926.5710, found: 926.5726 (1.7 ppm).

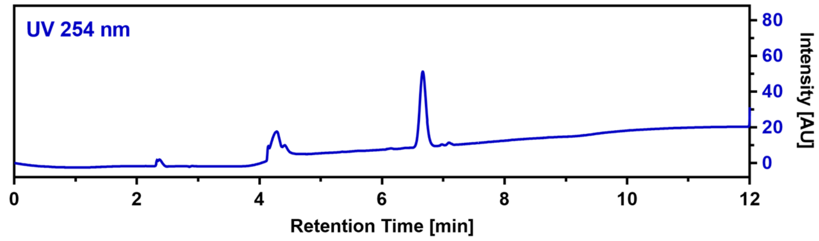

**CalA_C3A (24)**

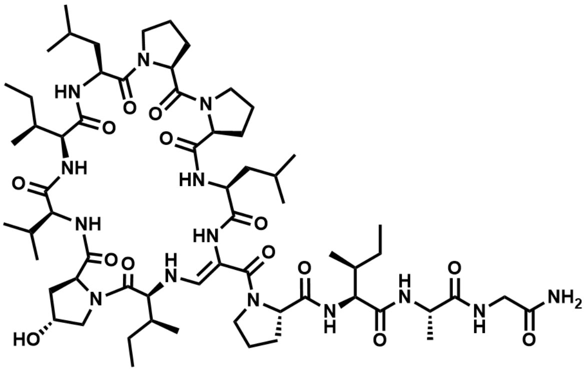

$C_{63}H_{104}N_{14}O_{14}$ M = 1281.61 g/mol

The linear precursor peptide of **24** was prepared via SPPS using Rink amide PS resin at a scale of 0.1 mmol initial resin capacity and purified by HPLC (solvent A: $H_2O$, acidified with 0.1% TFA; solvent B: ACN, acidified with 0.1% TFA). The obtained peptide was then dissolved in 10 mL of ACN and 4 equivalents DMP were added. The resulting suspension was shaken for one hour at RT and immediately purified by HPLC (solvent A: $H_2O$; solvent B: ACN) to give 1.4 mg (1.1 µmol) of **24** in a total yield of 1.1% (calculated in relation to initial resin capacity) as a white powder.

HRMS (ESI): *m/z* calculated for $[M+H]^+$: 1281.7929, found: 1281.7929 (0.0 ppm).

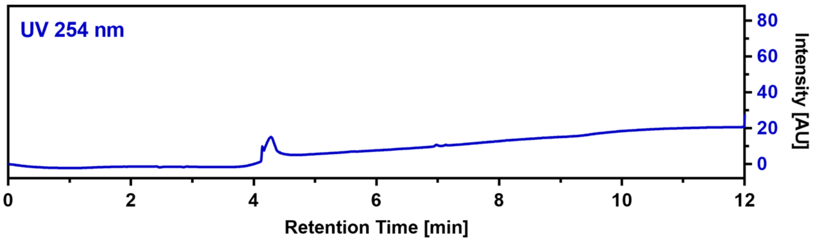

**CalA_C4A (25)**

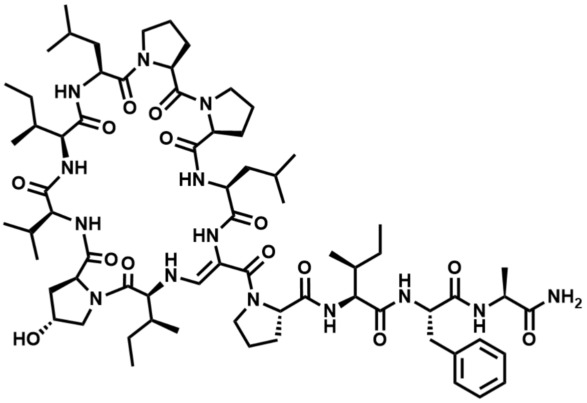

$C_{70}H_{110}N_{14}O_{14}$ M = 1371.73 g/mol

The linear precursor peptide of **25** was prepared via SPPS using Rink amide PS resin at a scale of 0.1 mmol initial resin capacity and purified by HPLC (solvent A: $H_2O$, acidified with 0.1% TFA; solvent B: ACN, acidified with 0.1% TFA). The obtained peptide was then dissolved in 10 mL of ACN and 4 equivalents DMP were added. The resulting suspension was shaken for one hour at RT and immediately purified by HPLC (solvent A: $H_2O$; solvent B: ACN) to give 2.6 mg (3.6 µmol) of **25** in a total yield of 3.6% (calculated in relation to initial resin capacity) as a white powder.

$^1H$ NMR (400 MHz, DMSO) δ 8.98 (s, 1H), 8.77 (s, 1H), 8.13 (s, 1H), 7.72 (d, *J* = 7.3 Hz, 1H), 7.48 (d, *J* = 7.3 Hz, 1H), 7.42 (d, *J* = 8.5 Hz, 1H), 7.36 (d, *J* = 7.1 Hz, 1H), 7.26-7.20 (m, 2H), 7.18 (d, *J* = 13.3 Hz, 1H), 7.10-7.02 (m, 4H), 6.74 (d, *J* = 9.9 Hz, 1H), 6.68 (s, 1H), 5.78-5.66 (m, 1H), 4.64 (q, *J* = 6.8 Hz, 1H), 4.60-4.51 (m, 1H), 4.38 (s, 1H), 4.29-4.19 (m, 3H), 4.18-4.11 (m, 1H), 4.10-4.03 (m, 2H), 3.90-3.83 (m, 2H), 3.66-3.56 (m, 4H), 3.53-3.46 (m, 2H), 3.39-3.31 (m, 1H), 3.27-3.18 (m, 1H), 3.14-3.07 (m, 1H), 2.99 (t, *J* = 8.9 Hz, 1H), 2.81 (t, *J* = 12.7 Hz, 1H), 2.70-2.58 (m, 1H), 2.34-2.19 (m, 3H), 2.12-2.05 (m, 1H), 2.02-1.96 (m, 1H), 1.95-1.80 (m, 8H), 1.78-1.68 (m, 4H), 1.66-1.53 (m, 4H), 1.51-1.43 (m, 1H), 1.42-1.33 (m, 2H), 1.31 (d, *J* = 7.3 Hz, 3H), 1.28-1.25 (m, 1H), 1.23 (s, 1H), 1.20-1.06 (m, 3H), 0.96 (d, *J* = 6.5 Hz, 3H), 0.92 (d, *J* = 6.4 Hz, 3H), 0.88 (d, *J* = 2.2 Hz, 3H), 0.87 (d, *J* = 2.1 Hz, 3H), 0.84-0.79 (m, 9H), 0.77 (t, *J* = 7.5 Hz, 3H), 0.73 (d, *J* = 6.7 Hz, 3H), 0.70 (t, *J* = 7.4 Hz, 3H), 0.60-0.53 (m, 1H), 0.50 (d, *J* = 6.8 Hz, 3H), 0.26 (t, *J* = 7.3 Hz, 3H); HRMS (ESI): *m/z* calculated for $[M+H]^+$: 1371.8399, found: 1371.8411

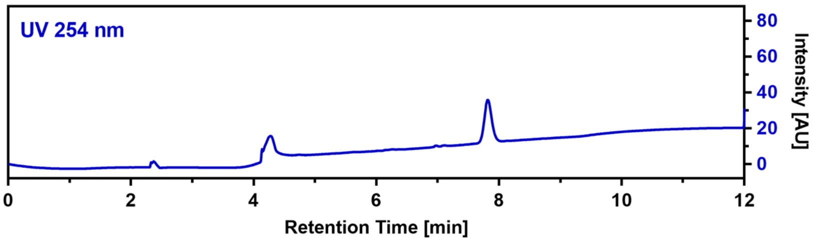

(0.9 ppm); calculated for $[M+2H]^{2+}$: 686.4236, found: 686.4234 (-0.3 ppm); $MS^2$ HRMS (ESI, HCD): *m/z* calculated for ring fragment $[M_{ring}+H]^+$: 926.5710, found: 926.5709 (-0.1 ppm).

**CalA_C5A (26)**

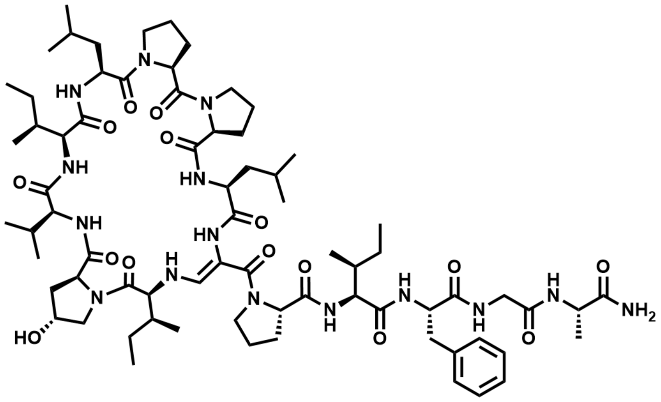

$C_{72}H_{113}N_{15}O_{15}$ M = 1428.79 g/mol

The linear precursor peptide of **26** was prepared via SPPS using Rink amide PS resin at a scale of 0.1 mmol initial resin capacity and purified by HPLC (solvent A: $H_2O$, acidified with 0.1% TFA; solvent B: ACN, acidified with 0.1% TFA). The obtained peptide was then dissolved in 10 mL of ACN and 4 equivalents DMP were added. The resulting suspension was shaken for one hour at RT and immediately purified by HPLC (solvent A: $H_2O$; solvent B: ACN) to give 3.0 mg (2.1 µmol) of **26** in a total yield of 2.1% (calculated in relation to initial resin capacity) as a white powder.

$^1H$ NMR (400 MHz, DMSO) δ 9.04 (d, *J* = 6.9 Hz, 1H), 8.83 (s, 1H), 8.49 (t, *J* = 6.0 Hz, 1H), 8.34 (s, 1H), 8.22 (d, *J* = 7.8 Hz, 1H), 7.54 (d, *J* = 13.3 Hz, 1H), 7.47 (d, *J* = 7.2 Hz, 1H), 7.44 (d, *J* = 9.4 Hz, 1H), 7.27 (d, *J* = 8.5 Hz, 1H), 7.20-7.11 (m, 6H), 7.04 (s, 1H), 6.72 (d, *J* = 10.1 Hz, 1H), 5.96-5.87 (m, 1H), 5.36 (d, *J* = 3.2 Hz, 1H), 4.67-4.58 (m, 2H), 4.39 (s, 1H), 4.34-4.24 (m, 4H), 4.23-4.16 (m, 2H), 4.06 (t, *J* = 9.4 Hz, 1H), 4.00 (dd, *J* = 15.9, 6.5 Hz, 1H), 3.82-3.75 (m, 2H), 3.74-3.69 (m, 1H), 3.66-3.57 (m, 2H), 3.54-3.45 (m, 3H), 3.25-3.17 (m, 1H), 3.08 (d, *J* = 13.5 Hz, 1H), 3.02-2.94 (m, 1H), 2.72-2.62 (m, 1H), 2.61-2.52 (m, 1H), 2.33-2.21 (m, 3H), 2.13-2.05 (m, 1H), 2.02-1.97 (m, 1H), 1.96-1.83 (m, 7H), 1.82-1.72 (m, 3H), 1.62 (s, 6H), 1.52-1.37 (m, 6H), 1.24 (d, *J* = 4.8 Hz, 3H), 1.23-1.20 (m, 1H), 1.18-1.10 (m, 1H), 1.06-0.98 (m, 1H), 0.95 (d, *J* = 6.3 Hz, 3H), 0.88-0.76 (m, 24H), 0.73-0.67 (m, 4H), 0.45 (t, *J* = 7.2 Hz, 3H), 0.35 (d, *J* = 6.7 Hz, 3H); HRMS (ESI): *m/z* calculated for $[M+H]^+$: 1428.8613, found: 1428.8625 (0.8 ppm); calculated for $[M+2H]^{2+}$: 714.9343, found: 714.9346 (0.4 ppm); $MS^2$ HRMS (ESI, HCD): *m/z* calculated for ring fragment $[M_{ring}+H]^+$: 926.5710, found: 926.5725 (1.6 ppm).

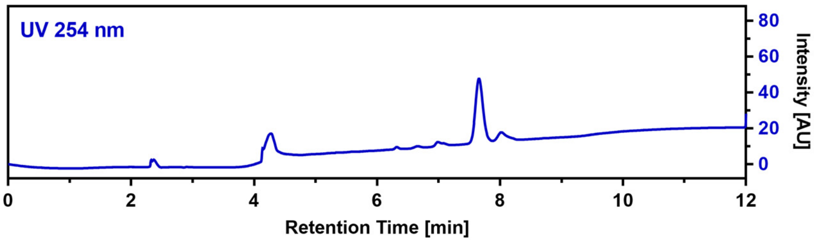

**CalB_R2dPro (27)**

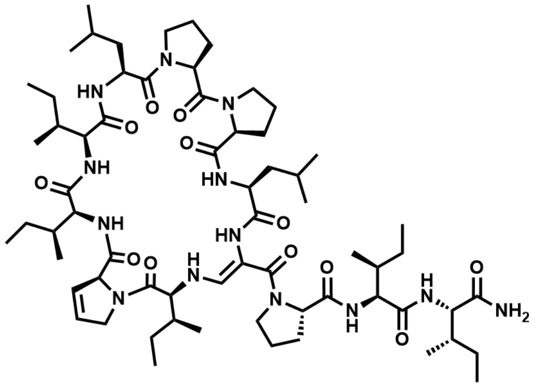

$C_{65}H_{107}N_{13}O_{12}$ M = 1262.65 g/mol

The linear precursor peptide of **27** was prepared via microwave-assisted SPPS using Rink amide PS resin at a scale of 0.1 mmol initial resin capacity. After purification by HPLC (solvent A: $H_2O$, acidified with 0.1% TFA; solvent B: ACN, acidified with 0.1% TFA), 69 mg (54 µmol, 54%) of the peptide precursor was obtained as a white powder. For the oxidation of the serine residue and concomitant cyclization, the peptide was dissolved in 15 mL of ACN and 4 equivalents of DMP were added. The resulting suspension was shaken for 5 hours at RT and immediately purified by HPLC (solvent A: $H_2O$; solvent B: ACN) to give 0.5 mg (0.4 µmol, 0.7%) of **27** in a total yield of 0.4% (calculated in relation to initial resin capacity) as a white powder.

HRMS (ESI): *m/z* calculated for $[M+H]^+$: 1262.8235, found: 1262.8218 (-1.3 ppm); calculated for $[M+2H]^{2+}$: 631.9154, found: 631.9143 (-1.7 ppm); $MS^2$ HRMS (ESI, HCD): *m/z* calculated for ring fragment $[M_{ring}+H]^+$: 922.5761, found: 922.5724 (-4.0 ppm).

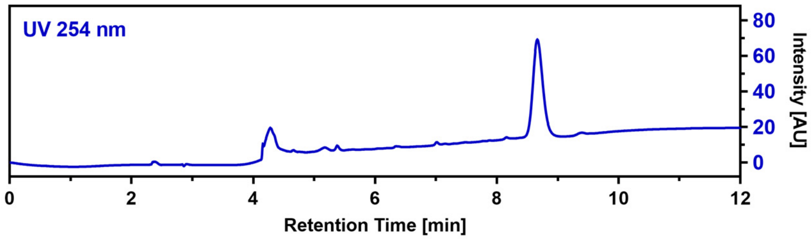

**CalB_R2Aze (28)**

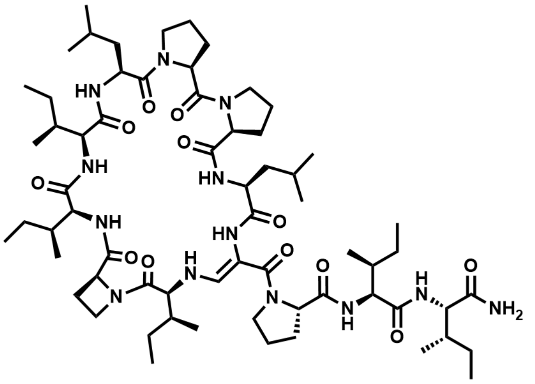

$C_{64}H_{107}N_{13}O_{12}$ M = 1250.6 g/mol

The linear precursor peptide of **28** was prepared via microwave-assisted SPPS using Rink amide PS resin at a scale of 0.1 mmol initial resin capacity. After purification by HPLC (solvent A: $H_2O$, acidified with 0.1% TFA; solvent B: ACN, acidified with 0.1% TFA), 55 mg (43 µmol, 43%) of the peptide precursor was obtained as a white powder. For the oxidation of the serine residue and concomitant cyclization, the peptide was dissolved in 10 mL of ACN and 3 equivalents of DMP were added. The resulting suspension was shaken for 2 hours at RT and immediately purified by HPLC (solvent A: $H_2O$; solvent B: ACN) to give 8.4 mg (6.7 µmol, 15.5%) of **28** in a total yield of 6.7% (calculated in relation to initial resin capacity) as a white powder. $^1H$ NMR (400 MHz, DMSO-d6) δ 9.00 (d, *J* = 7.1 Hz, 1H), 8.50 (d, *J* = 6.6 Hz, 1H), 8.22 (s, 1H), 7.59 (d, *J* = 5.8 Hz, 1H), 7.57 (d, *J* = 3.0 Hz, 1H), 7.20 (d, *J* = 9.7 Hz, 1H), 7.12 (s, 1H), 7.02 (d, *J* = 13.2 Hz, 1H), 6.91 (s, 1H), 6.73 (d, *J* = 9.8 Hz, 1H), 5.55-5.45 (m, 1H), 4.66-4.59 (m, 2H), 4.55-4.46 (m, 1H), 4.39-4.31 (m, 1H), 4.28-4.18 (m, 3H), 4.12-4.04 (m, 2H), 3.99-3.92 (m, 1H), 3.86-3.80 (m, 1H), 3.67-3.61 (m, 1H), 3.60-3.53 (m, 2H), 3.51-3.44 (m, 2H), 3.35-3.26 (m, 1H), 3.23-3.13 (m, 2H), 2.61-2.53 (m, 1H), 2.46-2.38 (m, 1H), 2.30-2.16 (m, 3H), 2.01-1.94 (m, 1H), 1.93-1.76 (m, 8H), 1.74-1.65 (m, 3H), 1.65-1.56 (m, 3H), 1.52-1.36 (m, 8H), 1.35-1.22 (m, 2H), 1.21-1.11 (m, 2H), 1.08-0.96 (m, 3H), 0.92-0.74 (m, 42H); HRMS (ESI): *m/z* calculated for $[M+H]^+$: 1250.8235, found: 1250.8229 (-0.5 ppm); calculated for $[M+2H]^{2+}$: 625.9154, found: 625.9148 (-1.0 ppm); $MS^2$ HRMS (ESI, HCD): *m/z* calculated for ring fragment $[M_{ring}+H]^+$: 910.5761, found: 910.5733 (-3.1 ppm).

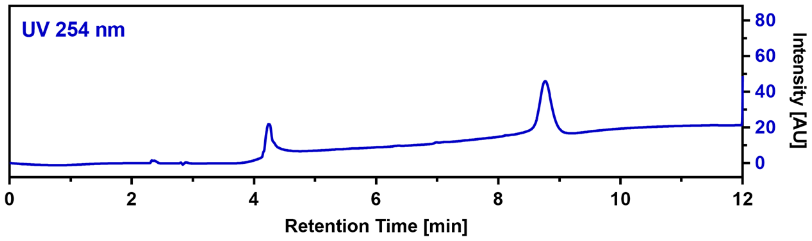

**CalB_R2Pip (29)**

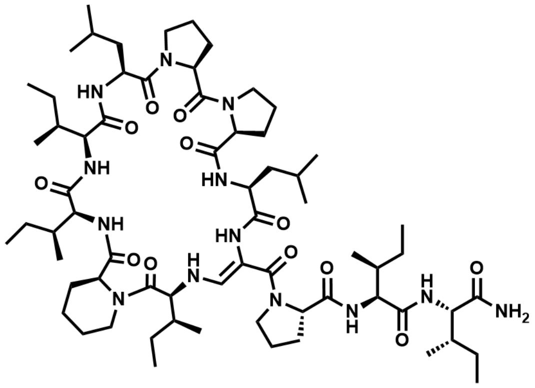

$C_{66}H_{111}N_{13}O_{12}$ M = 1278.7 g/mol

The linear precursor peptide of **29** was prepared via microwave-assisted SPPS using Rink amide PS resin at a scale of 0.1 mmol initial resin capacity. After purification by HPLC (solvent A: $H_2O$, acidified with 0.1% TFA; solvent B: ACN, acidified with 0.1% TFA), 38 mg (29 µmol, 29%) of the peptide precursor was obtained as a white powder. For the oxidation of the serine residue and concomitant cyclization, the peptide was dissolved in 8 mL of ACN and 3 equivalents of DMP were added. The resulting suspension was shaken for 2 hours at RT and immediately purified by HPLC (solvent A: $H_2O$; solvent B: ACN) to give 2.0 mg

(1.6 µmol, 5.3%) of **29** in a total yield of 1.6% (calculated in relation to initial resin capacity) as a white powder.

$^1$H NMR (400 MHz, DMSO) δ 8.85 (d, $J$ = 7.2 Hz, 1H), 8.61 (d, $J$ = 6.8 Hz, 1H), 8.24 (s, 1H), 7.57 (d, $J$ = 10.2 Hz, 1H), 7.22 (d, $J$ = 9.6 Hz, 1H), 7.16 (s, 1H), 7.09-7.00 (m, 2H), 6.90 (s, 1H), 6.67 (d, $J$ = 10.1 Hz, 1H), 5.70 (dd, $J$ = 13.2, 9.9 Hz, 1H), 4.64 (q, $J$ = 6.7 Hz, 1H), 4.55 (td, $J$ = 10.3, 3.6 Hz, 1H), 4.34-4.19 (m, 3H), 4.16-4.03 (m, 4H), 3.98-3.89 (m, 1H), 3.83-3.75 (m, 1H), 3.62-3.42 (m, 5H), 3.20-3.06 (m, 3H), 2.61-2.52 (m, 1H), 2.35-2.17 (m, 4H), 1.96-1.82 (m, 9H), 1.81-1.71 (m, 3H), 1.68-1.55 (m, 6H), 1.53-1.37 (m, 9H), 1.31-1.19 (m, 3H), 1.16-0.97 (m, 4H), 0.91-0.69 (m, 42H); HRMS (ESI): $m/z$ calculated for $[M+H]^+$: 1278.8548, found: 1278.8534 (-1.1 ppm); calculated for $[M+2H]^{2+}$: 639.9311, found: 639.9300 (-1.7 ppm); $MS^2$ HRMS (ESI, HCD): $m/z$ calculated for ring fragment $[M_{ring}+H]^+$: 938.6074, found: 938.6056 (-1.9 ppm).

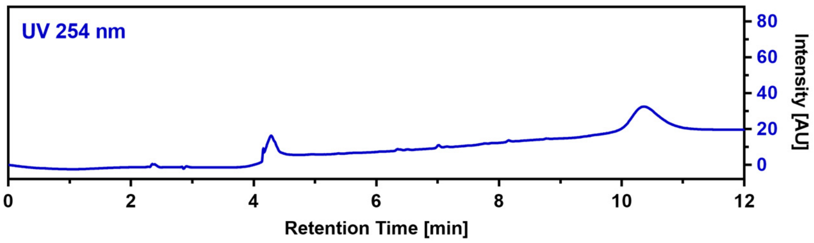

**CalB_R2Nip (30)**

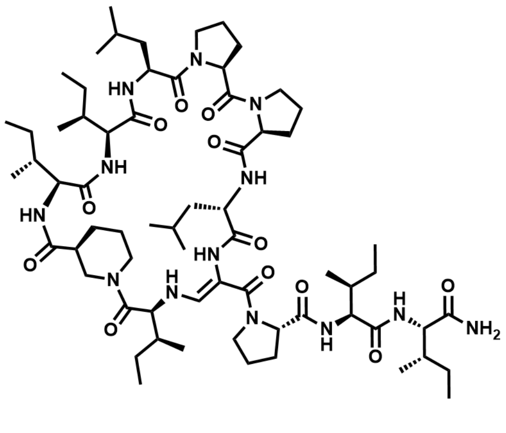

$C_{66}H_{111}N_{13}O_{12}$ M = 1278.7 g/mol

The linear precursor peptide of **30** was prepared via microwave-assisted SPPS using Rink amide PS resin at a scale of 0.1 mmol initial resin capacity. After purification by HPLC (solvent A: $H_2O$, acidified with 0.1% TFA; solvent B: ACN, acidified with 0.1% TFA), 21 mg (16 µmol, 16%) of the peptide precursor was obtained as a white powder. For the oxidation of the serine residue and concomitant cyclization, the peptide was dissolved in 5 mL of ACN and 4 equivalents of DMP were added. The resulting suspension was shaken for 3 hours at RT and immediately purified by HPLC (solvent A: $H_2O$; solvent B: ACN) to give 1.4 mg (1.1 µmol, 6.6%) of **30** in a total yield of 1.1% (calculated in relation to initial resin capacity) as a white powder.

HRMS (ESI): $m/z$ calculated for $[M+H]^+$: 1278.8548, found: 1278.8551 (0.2 ppm); calculated for $[M+2H]^{2+}$: 639.9311, found: 639.9309 (-0.3 ppm); $MS^2$ HRMS (ESI, HCD): $m/z$ calculated for ring fragment $[M_{ring}+H]^+$: 938.6074, found: 938.6057 (-1.8 ppm).

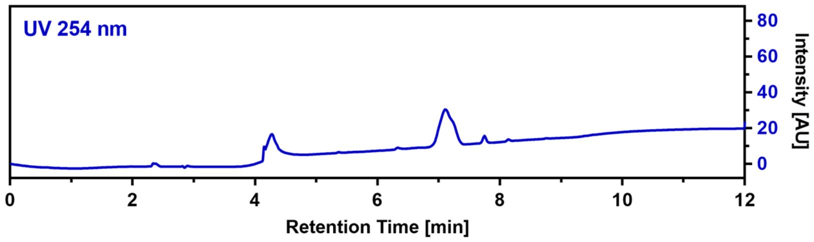

#### CalB_R2Oic (31)

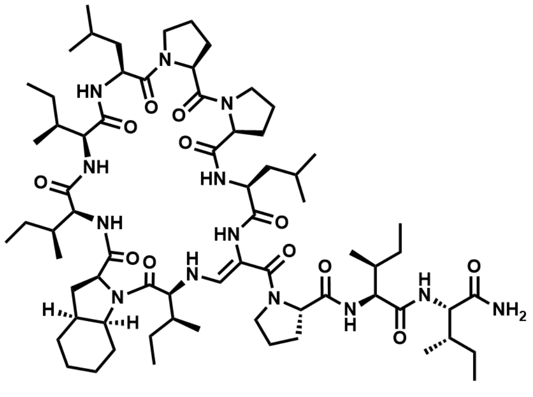

$C_{69}H_{115}N_{13}O_{12}$ M = 1318.8 g/mol

The linear precursor peptide of **31** was prepared via microwave-assisted SPPS using Rink amide PS resin at a scale of 0.1 mmol initial resin capacity. After purification by HPLC (solvent A: $H_2O$, acidified with 0.1% TFA; solvent B: ACN, acidified with 0.1% TFA), 61 mg (45 µmol, 45%) of the peptide precursor was obtained as a white powder. For the oxidation of the serine residue and concomitant cyclization, the peptide was dissolved in 15 mL of ACN and 4 equivalents of DMP were added. The resulting suspension was shaken for 3 hours at RT and immediately purified by HPLC (solvent A: $H_2O$; solvent B: ACN) to give 26 mg (20 µmol, 44%) of **31** in a total yield of 19.8% (calculated in relation to initial resin capacity) as a white powder.

$^1$H NMR (400 MHz, DMSO) δ 8.84 (d, $J$ = 7.0 Hz, 1H), 8.59 (d, $J$ = 6.6 Hz, 1H), 8.27 (s, 1H), 7.60 (d, $J$ = 3.4 Hz, 1H), 7.58 (s, 1H), 7.29 (d, $J$ = 9.6 Hz, 1H), 7.12 (d, $J$ = 13.4 Hz, 1H), 7.04 (s, 1H), 7.02 (s, 1H), 6.67 (d, $J$ = 9.9 Hz, 1H), 5.62-5.53 (m, 1H), 4.65-4.57 (m, 1H), 4.57-4.48 (m, 1H), 4.31-4.20 (m, 2H), 4.20-4.14 (m, 1H), 4.14-4.01 (m, 3H), 3.95 (t, $J$ = 8.7 Hz, 1H), 3.81 (t, $J$ = 10.1 Hz, 1H), 3.77-3.71 (m, 1H), 3.62-3.40 (m, 4H), 3.30-3.24 (m, 1H), 3.23-3.13 (m, 1H), 3.11-3.04 (m, 1H), 2.69-2.57 (m, 1H), 2.37-2.17 (m, 3H), 2.05-1.96 (m, 2H), 1.95-1.82 (m, 8H), 1.81-1.55 (m, 10H), 1.54-1.38 (m, 8H), 1.33-0.98 (m, 10H), 0.92-0.69 (m, 44H); HRMS (ESI): $m/z$ calculated for $[M+H]^+$: 1318.8861, found: 1318.8862 (0.1 ppm); calculated for $[M+2H]^{2+}$: 659.9467, found: 659.9461 (-0.9 ppm); $MS^2$ HRMS (ESI, HCD): $m/z$ calculated for ring fragment $[M_{ring}+H]^+$: 978.6387, found: 978.6381 (-0.6 ppm).

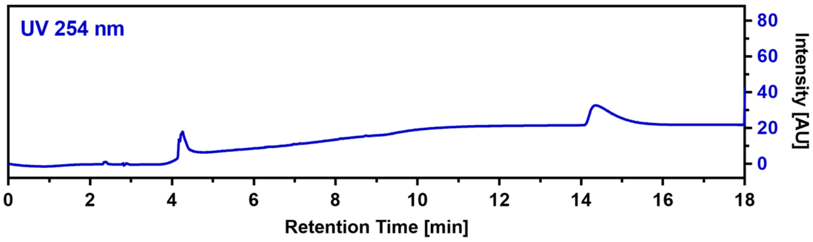

#### CalB_R3hLeu (32)

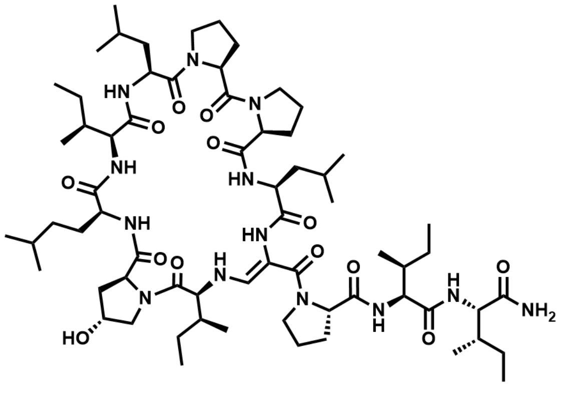

$C_{66}H_{111}N_{13}O_{13}$ M = 1294.7 g/mol

The linear precursor peptide of **32** was prepared via microwave-assisted SPPS using Rink amide PS resin at a scale of 0.1 mmol initial resin capacity. After purification by HPLC (solvent A: $H_2O$, acidified with 0.1% TFA; solvent B: ACN, acidified with 0.1% TFA), 25 mg (19 µmol, 19%) of the peptide precursor was obtained as a white powder. For the oxidation of the serine residue and concomitant cyclization, the peptide was dissolved in 5 mL of ACN and 3 equivalents of DMP were added. The resulting suspension was shaken for 2 hours at RT and immediately purified by HPLC (solvent A: $H_2O$; solvent B: ACN) to give 2.1 mg (1.6 µmol, 8.5%) of **32** in a total yield of 1.6% (calculated in relation to initial resin capacity) as a white powder.

$^{1}$H NMR (400 MHz, DMSO) δ 8.87 (d, *J* = 6.8 Hz, 1H), 8.30 (s, 1H), 8.21 (d, *J* = 8.1 Hz, 1H), 7.60 (d, *J* = 10.0 Hz, 1H), 7.45 (d, *J* = 7.3 Hz, 1H), 7.23 (d, *J* = 9.6 Hz, 1H), 7.15 (s, 1H), 7.06 (d, *J* = 13.2 Hz, 1H), 6.97 (s, 1H), 6.67 (d, *J* = 9.9 Hz, 1H), 5.61 (dd, *J* = 13.2, 10.1 Hz, 1H), 4.66 (q, *J* = 6.8 Hz, 1H), 4.52 (td, *J* = 10.3, 3.6 Hz, 1H), 4.42 (s, 1H), 4.29-4.20 (m, 3H), 4.19-4.13 (m, 1H), 4.13-4.05 (m, 1H), 4.06-3.92 (m, 3H), 3.78-3.68 (m, 2H), 3.62-3.49 (m, 5H), 3.21-3.11 (m, 1H), 2.36-2.17 (m, 3H), 2.13-1.99 (m, 3H), 1.96-1.77 (m, 9H), 1.73-1.55 (m, 6H), 1.54-1.38 (m, 8H), 1.37-1.29 (m, 1H), 1.27-1.20 (m, 2H), 1.19-1.06 (m, 4H), 1.05-0.94 (m, 2H), 0.93-0.69 (m, 42H); HRMS (ESI): *m/z* calculated for $[M+H]^{+}$: 1294.8497, found: 1294.8481 (-1.2 ppm); calculated for $[M+2H]^{2+}$: 647.9285, found: 647.9274 (-1.7 ppm); MS$^2$ HRMS (ESI, HCD): *m/z* calculated for ring fragment $[M_{ring}+H]^{+}$: 954.6023, found: 954.5990 (-3.5 ppm).

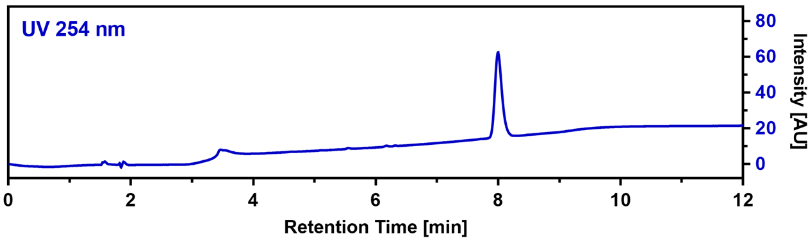

**CalB_R3Aib (33)**

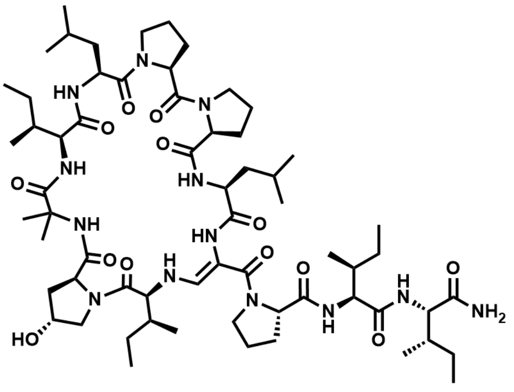

$C_{63}H_{105}N_{13}O_{13}$ M = 1252.6 g/mol

The linear precursor peptide of **33** was prepared via microwave-assisted SPPS using Rink amide PS resin at a scale of 0.1 mmol initial resin capacity. After purification by HPLC (solvent A: $H_2O$, acidified with 0.1% TFA; solvent B: ACN, acidified with 0.1% TFA), 42 mg (33 µmol, 33%) of the peptide precursor was obtained as a white powder. For the oxidation of the serine residue and concomitant cyclization, the peptide was dissolved in 10 mL of ACN and 4 equivalents of DMP were added. The resulting suspension was shaken for 5 hours at RT and immediately purified by HPLC (solvent A: $H_2O$; solvent B: ACN) to give 2.6 mg (2.1 µmol, 6.3%) of **33** in a total yield of 2.1% (calculated in relation to initial resin capacity) as a white powder.

HRMS (ESI): *m/z* calculated for $[M+H]^{+}$: 1252.8028, found: 1252.8015 (-1.0 ppm); calculated for $[M+2H]^{2+}$: 626.9050, found: 626.9044 (-1.0 ppm); MS$^2$ HRMS (ESI, HCD): *m/z* calculated for ring fragment $[M_{ring}+H]^{+}$: 912.5554, found: 912.5514 (-4.4 ppm).

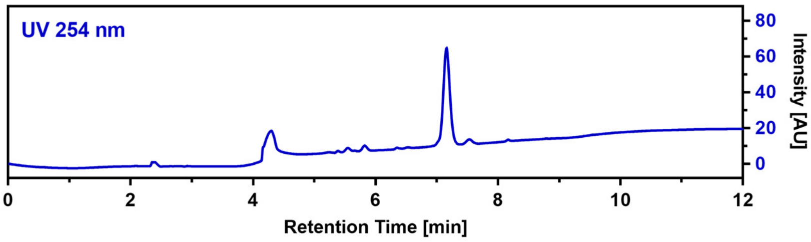

**CalB_R3Chg (34)**

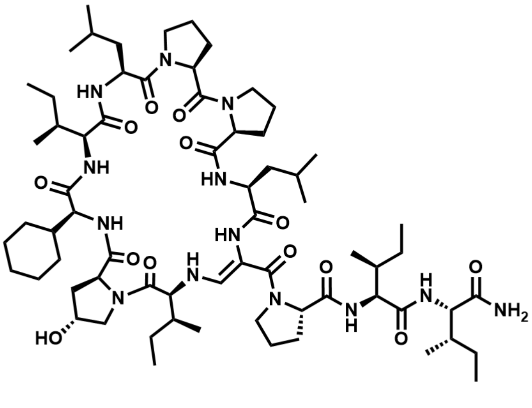

$C_{67}H_{111}N_{13}O_{13}$ M = 1306.7 g/mol

The linear precursor peptide of **34** was prepared via microwave-assisted SPPS using Rink amide PS resin at a scale of 0.1 mmol initial resin capacity. After purification by HPLC (solvent A: $H_2O$, acidified with 0.1% TFA; solvent B: ACN, acidified with 0.1% TFA), 40 mg (31 µmol, 31%) of the peptide precursor was obtained as a white powder. For the oxidation of the serine residue and concomitant cyclization, the peptide was dissolved in 10 mL of ACN and 4 equivalents of DMP were added. The resulting suspension was shaken for 5 hours at RT and immediately purified by HPLC (solvent A: $H_2O$; solvent B: ACN) to give 6.8 mg (5.1 µmol, 16.5%) of **34** in a total yield of 5.1% (calculated in relation to initial resin capacity) as a white powder.

$^1$H NMR (400 MHz, DMSO) δ 9.02 (d, $J$ = 7.0 Hz, 1H), 8.86 (d, $J$ = 6.6 Hz, 1H), 8.29 (s, 1H), 7.59 (d, $J$ = 10.1 Hz, 1H), 7.36 (d, $J$ = 7.0 Hz, 1H), 7.24 (d, $J$ = 9.7 Hz, 1H), 7.17 (s, 1H), 7.05 (d, $J$ = 13.3 Hz, 1H), 6.97 (s, 1H), 6.65 (d, $J$ = 10.0 Hz, 1H), 5.78-5.67 (m, 1H), 5.41 (s, 1H), 4.66-4.59 (m, 1H), 4.58-4.50 (m, 1H), 4.41 (s, 1H), 4.29-4.21 (m, 3H), 4.19-4.08 (m, 3H), 4.08-4.01 (m, 1H), 4.00-3.91 (m, 1H), 3.84-3.76 (m, 2H), 3.70-3.43 (m, 6H), 3.21-3.11 (m, 1H), 3.11-3.03 (m, 1H), 2.44-2.34 (m, 1H), 2.35-2.17 (m, 3H), 2.14-2.03 (m, 1H), 2.02-1.95 (m, 1H), 1.95-1.82 (m, 8H), 1.82-1.55 (m, 10H), 1.52-1.37 (m, 8H), 1.33-1.21 (m, 3H), 1.20-1.08 (m, 4H), 1.05-0.94 (m, 3H), 0.92-0.68 (m, 36H); HRMS (ESI): $m/z$ calculated for $[M+H]^+$: 1306.8497, found: 1306.8494 (-0.2 ppm); calculated for $[M+2H]^{2+}$: 653.9285, found: 653.9279 (-0.9 ppm); $MS^2$ HRMS (ESI, HCD): $m/z$ calculated for ring fragment $[M_{ring}+H]^+$: 966.6023, found: 966.5992 (-3.2 ppm).

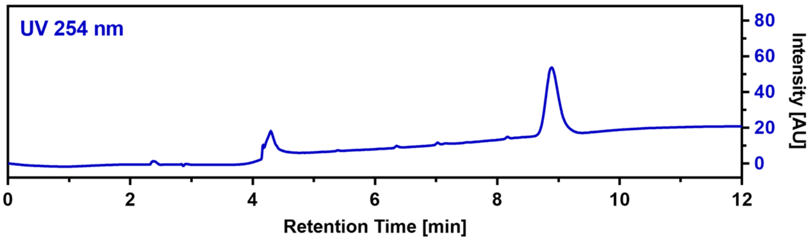

**CalB_R3F (35)**

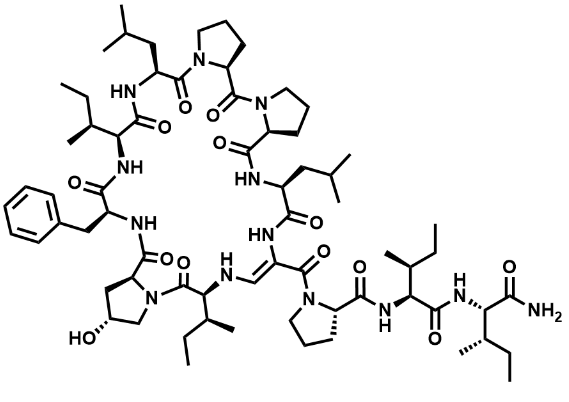

$C_{68}H_{107}N_{13}O_{13}$ M = 1314.7 g/mol

The linear precursor peptide of **35** was prepared via microwave-assisted SPPS using Rink amide PS resin at a scale of 0.1 mmol initial resin capacity. After purification by HPLC (solvent A: $H_2O$, acidified with 0.1% TFA; solvent B: ACN, acidified with 0.1% TFA), 37 mg (28 µmol, 28%) of the peptide precursor was obtained as a white powder. For the oxidation of the serine residue and concomitant cyclization, the peptide was dissolved in 10 mL of ACN and 3 equivalents of DMP were added. The resulting suspension was shaken for 2 hours at RT and immediately purified by HPLC (solvent A: $H_2O$; solvent B: ACN) to give 8.2 mg (6.2 µmol, 22.4%) of **35** in a total yield of 6.2% (calculated in relation to initial resin capacity) as a white powder.

$^{1}$H NMR (400 MHz, DMSO) δ 8.79 (d, $J$ = 6.8 Hz, 1H), 8.27 (s, 1H), 8.24 (d, $J$ = 8.4 Hz, 1H), 7.58 (d, $J$ = 10.1 Hz, 1H), 7.50 (d, $J$ = 7.4 Hz, 1H), 7.36-7.28 (m, 2H), 7.27-7.20 (m, 2H), 7.19-7.16 (m, 1H), 7.15 (s, 1H), 7.04 (d, $J$ = 13.2 Hz, 1H), 6.98 (s, 1H), 6.64 (d, $J$ = 9.9 Hz, 1H), 5.67-5.56 (m, 1H), 5.33 (s, 1H), 4.68 (q, $J$ = 6.8 Hz, 1H), 4.51 (td, $J$ = 10.4, 3.6 Hz, 1H), 4.40 (s, 1H), 4.27-4.20 (m, 2H), 4.19-4.06 (m, 3H), 4.05-3.93 (m, 3H), 3.70-3.64 (m, 1H), 3.60-3.40 (m, 6H), 3.20-3.11 (m, 1H), 3.11-3.04 (m, 1H), 2.32-2.16 (m, 3H), 2.01-1.95 (m, 1H), 1.94-1.82 (m, 8H), 1.79-1.52 (m, 11H), 1.51-1.39 (m, 6H), 1.39-1.31 (m, 1H), 1.29-1.18 (m, 2H), 1.17-1.08 (m, 1H), 1.06-0.96 (m, 2H), 0.94-0.69 (m, 36H); HRMS (ESI): *m/z* calculated for $[M+H]^+$: 1314.8184, found: 1314.8184 (0.0 ppm); calculated for $[M+2H]^{2+}$: 657.9129, found: 657.9125 (-0.6 ppm); $MS^2$ HRMS (ESI, HCD): *m/z* calculated for ring fragment $[M_{ring}+H]^+$: 974.5710, found: 974.5693 (-1.7 ppm).

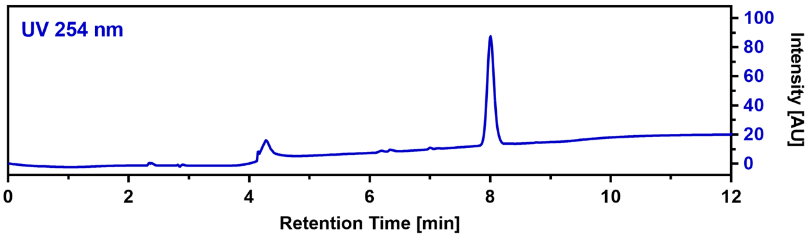

**CalB_R3β-hIle (36)**

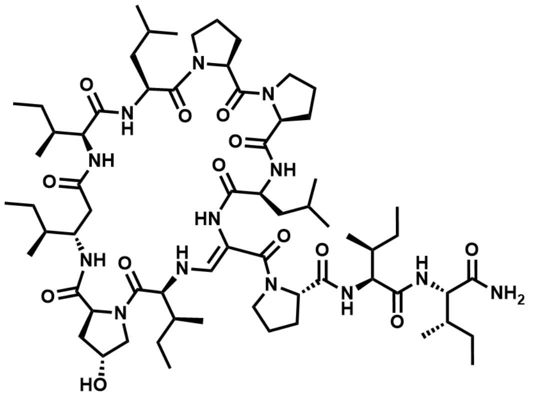

The linear precursor peptide of **36** was prepared via microwave-assisted SPPS using Rink amide PS resin at a scale of 0.1 mmol initial resin capacity. After purification by HPLC (solvent A: $H_2O$, acidified with 0.1% TFA; solvent B: ACN, acidified with 0.1% TFA), 51 mg (39 µmol, 39%) of the peptide precursor was obtained as a white powder. For the oxidation of the serine residue and concomitant cyclization, the peptide was dissolved in 10 mL of ACN and 4 equivalents of DMP were added. The resulting suspension was shaken for 3 hours at RT and immediately purified by HPLC (solvent A: $H_2O$; solvent B: ACN) to give 3.7 mg (2.9 µmol, 7.4%) of **36** in a total yield of 2.9% (calculated in relation to initial resin capacity) as a white powder. HRMS (ESI): *m/z* calculated for $[M+H]^+$: 1294.8497, found: 1294.8488 (-0.7 ppm); calculated for $[M+2H]^{2+}$: 647.9285, found: 647.9277 (-1.2 ppm); $MS^2$ HRMS (ESI, HCD): *m/z* calculated for ring fragment $[M_{ring}+H]^+$: 954.6023, found: 954.5994 (-3.0 ppm).

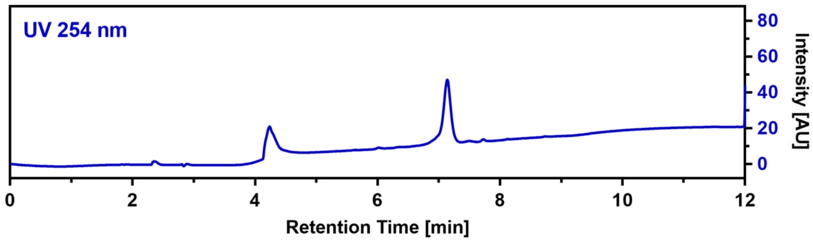

**CalB_R4β-hIle (37)**

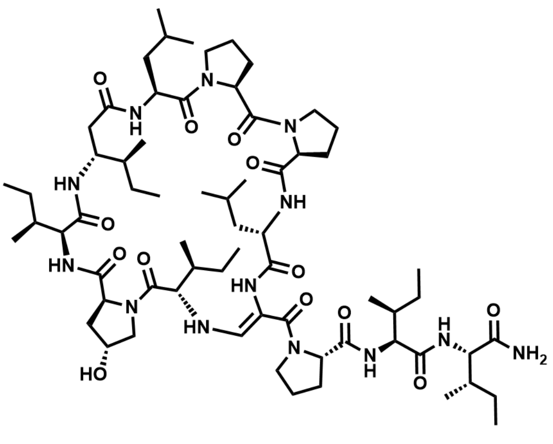

$C_{66}H_{111}N_{13}O_{13}$ M = 1294.7 g/mol

The linear precursor peptide of **37** was prepared via microwave-assisted SPPS using Rink amide PS resin at a scale of 0.1 mmol initial resin capacity. After purification by HPLC (solvent A: $H_2O$, acidified with 0.1% TFA; solvent B: ACN, acidified with 0.1% TFA), 44 mg (33 µmol, 33%) of the peptide precursor was obtained as a white powder. For the oxidation of the serine residue and concomitant cyclization, the peptide was dissolved in 5 mL of ACN and 4 equivalents of DMP were added. The resulting suspension was shaken for 4 hours at RT and immediately purified by HPLC (solvent A: $H_2O$; solvent B: ACN) to give 0.9 mg (0.7 µmol, 2.1%) of **37** in a total yield of 0.7% (calculated in relation to initial resin capacity) as a white powder.

HRMS (ESI): *m/z* calculated for $[M+H]^+$: 1294.8497, found: 1294.8492 (-0.4 ppm); calculated for $[M+2H]^{2+}$: 647.9285, found: 647.9277 (-1.2 ppm); $MS^2$ HRMS (ESI, HCD): *m/z* calculated for ring fragment $[M_{ring}+H]^+$: 954.6023, found: 954.5995 (-2.9 ppm).

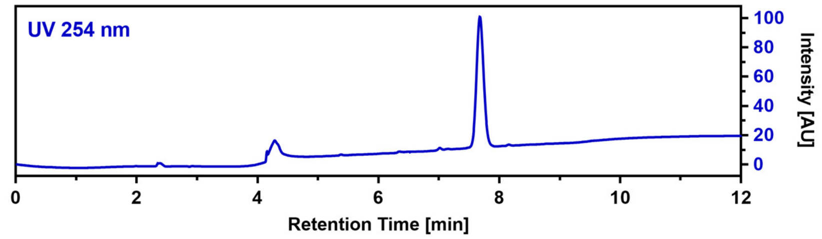

**CalB_R5hLeu (38)**

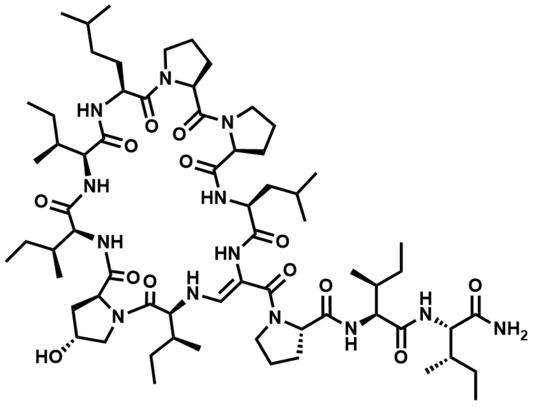

$C_{66}H_{111}N_{13}O_{13}$ M = 1294.7 g/mol

The linear precursor peptide of **38** was prepared via microwave-assisted SPPS using Rink amide PS resin at a scale of 0.1 mmol initial resin capacity. After purification by HPLC (solvent A: $H_2O$, acidified with 0.1% TFA; solvent B: ACN, acidified with 0.1% TFA), 33 mg (25 µmol, 25%) of the peptide precursor was obtained as a white powder. For the oxidation of the serine residue and concomitant cyclization, the peptide was dissolved in 8 mL of ACN and 3 equivalents of DMP were added. The resulting suspension was shaken for 2 hours at RT and immediately purified by HPLC (solvent A: $H_2O$; solvent B: ACN) to give 7.8 mg (6.0 µmol, 23.7%) of **38** in a total yield of 6.0% (calculated in relation to initial resin capacity) as a white powder.

$^1$H NMR (400 MHz, DMSO) δ 8.96 (d, *J* = 6.9 Hz, 1H), 8.82 (s, 1H), 8.29 (s, 1H), 7.59 (d, *J* = 10.0 Hz, 1H), 7.33 (d, *J* = 7.0 Hz, 1H), 7.23 (d, *J* = 9.7 Hz, 1H), 7.14 (s, 1H), 7.05 (d, *J* = 13.3 Hz, 1H), 6.95 (s, 1H), 6.73 (d, *J* = 9.8 Hz, 1H), 5.68 (dd, *J* = 13.2, 10.1 Hz, 1H), 5.39 (d, *J* = 3.0 Hz, 1H), 4.59 (q, *J* = 6.8 Hz, 1H), 4.52 (td, *J* = 10.2, 3.3 Hz, 1H), 4.41 (s, 1H), 4.28-4.19 (m, 3H), 4.15-4.00 (m, 3H), 3.93 (t, *J* = 10.0 Hz, 1H), 3.82-3.75 (m, 2H), 3.69-3.63 (m, 1H), 3.61-3.48 (m, 3H), 3.47-3.38 (m, 2H), 3.21-3.12 (m, 1H), 3.12-3.03 (m, 1H), 2.33-2.17 (m, 3H), 2.11-2.04 (m, 1H), 2.03-1.94 (m, 2H), 1.94-1.75 (m, 9H), 1.73-1.56 (m, 4H), 1.55-1.37 (m, 10H), 1.28-1.10 (m, 5H), 1.06-0.94 (m, 3H), 0.91-0.69 (m, 42H); HRMS (ESI): *m/z* calculated for $[M+H]^+$:

1294.8497, found: 1294.8497 (0.0 ppm); calculated for $[M+2H]^{2+}$: 647.9285, found: 647.9279 (-0.9 ppm); $MS^2$ HRMS (ESI, HCD): *m/z* calculated for ring fragment $[M_{ring}+H]^+$: 954.6023, found: 954.6000 (-2.4 ppm).

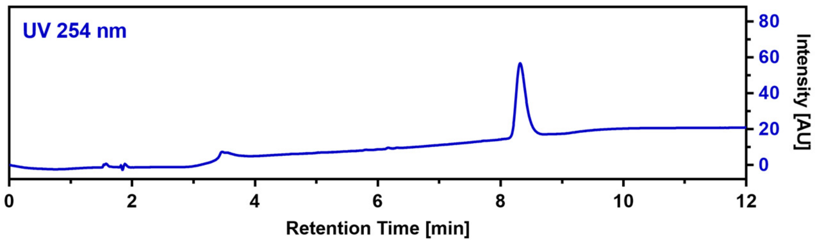

**CalB_R6dPro (39)**

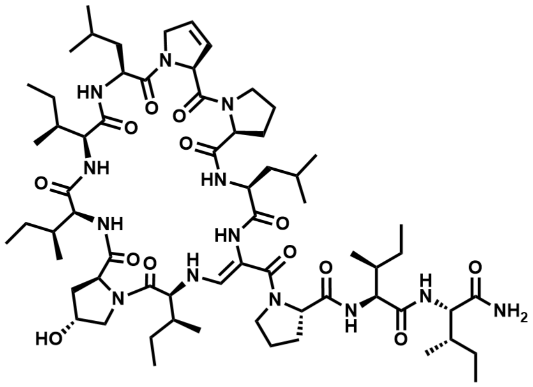

$C_{65}H_{107}N_{13}O_{13}$ M = 1278.65 g/mol

The linear precursor peptide of **39** was prepared via microwave-assisted SPPS using Rink amide PS resin at a scale of 0.1 mmol initial resin capacity. After purification by HPLC (solvent A: $H_2O$, acidified with 0.1% TFA; solvent B: ACN, acidified with 0.1% TFA), 23 mg (18 µmol, 18%) of the peptide precursor was obtained as a white powder. For the oxidation of the serine residue and concomitant cyclization, the peptide was dissolved in 5 mL of ACN and 4 equivalents of DMP were added. The resulting suspension was shaken for 4 hours at RT and immediately purified by HPLC (solvent A: $H_2O$; solvent B: ACN) to give 5.6 mg (4.4 µmol, 25.1%) of **39** in a total yield of 4.4% (calculated in relation to initial resin capacity) as a white powder.

$^1$H NMR (400 MHz, DMSO) δ 8.75 (d, *J* = 7.0 Hz, 1H), 8.59 (d, *J* = 6.8 Hz, 1H), 8.32 (s, 1H), 7.63 (d, *J* = 10.1 Hz, 1H), 7.35 (d, *J* = 7.2 Hz, 1H), 7.25-7.17 (m, 2H), 7.06 (d, *J* = 13.3 Hz, 1H), 6.90 (s, 1H), 6.61 (d, *J* = 9.9 Hz, 1H), 6.14-6.09 (m, 1H), 5.85-5.80 (m, 1H), 5.68 (dd, *J* = 13.2, 10.0 Hz, 1H), 4.82-4.77 (m, 1H), 4.72 (q, *J* = 6.8 Hz, 1H), 4.55 (td, *J* = 10.4, 3.7 Hz, 1H), 4.43-4.35 (m, 2H), 4.35-4.27 (m, 1H), 4.27-4.19 (m, 2H), 4.14-4.06 (m, 3H), 3.95 (t, *J* = 10.0 Hz, 1H), 3.89 (t, *J* = 7.6 Hz, 1H), 3.77-3.72 (m, 1H), 3.69-3.63 (m, 1H), 3.57-3.49 (m, 1H), 3.29-3.21 (m, 2H), 3.20-3.12 (m, 1H), 3.11-3.05 (m, 1H), 2.48-2.41 (m, 1H), 2.31-2.19 (m, 2H), 2.09-2.01 (m, 1H), 1.93-1.81 (m, 5H), 1.80-1.59 (m, 7H), 1.56-1.38 (m, 9H), 1.38-1.30 (m, 1H), 1.23 (s, 1H), 1.19-1.08 (m, 2H), 1.06-0.95 (m, 3H), 0.94-0.70 (m, 42H); HRMS (ESI): *m/z* calculated for $[M+H]^+$: 1278.8184, found: 1278.8171 (-1.0 ppm); calculated for $[M+2H]^{2+}$: 639.9129, found: 639.9119 (-1.6 ppm); $MS^2$ HRMS (ESI, HCD): *m/z* calculated for ring fragment $[M_{ring}+H]^+$: 938.5710, found: 938.5702 (-0.9 ppm).

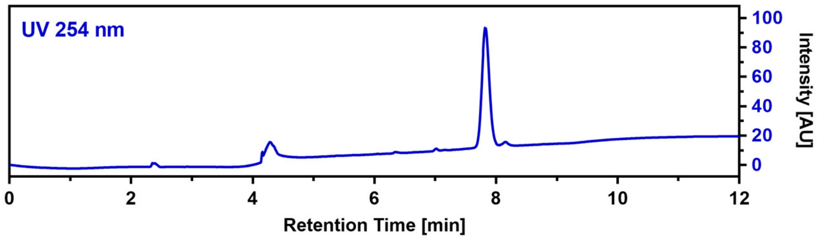

**CalB_R6Aze (40)**

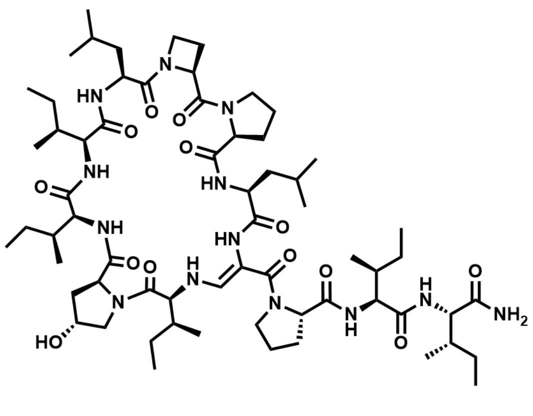

$C_{64}H_{107}N_{13}O_{13}$ M = 1266.64 g/mol

The linear precursor peptide of **40** was prepared via microwave-assisted SPPS using Rink amide PS resin at a scale of 0.1 mmol initial resin capacity. After purification by HPLC (solvent A: $H_2O$, acidified with 0.1% TFA; solvent B: ACN, acidified with 0.1% TFA), 30 mg (23 µmol, 23%) of the peptide precursor was obtained as a white powder. For the oxidation of the serine residue and concomitant cyclization, the peptide was dissolved in 5 mL of ACN and 3 equivalents of DMP were added. The resulting suspension was shaken for 2 hours at RT and immediately purified by HPLC (solvent A: $H_2O$; solvent B: ACN) to give 1.1 mg (0.9 µmol, 3.7%) of **40** in a total yield of 0.9% (calculated in relation to initial resin capacity) as a white powder.

HRMS (ESI): *m/z* calculated for $[M+H]^+$: 1266.8184, found: 1266.8150 (-2.7 ppm); calculated for $[M+2H]^{2+}$: 633.9129, found: 633.9111 (-2.8 ppm); $MS^2$ HRMS (ESI, HCD): *m/z* calculated for ring fragment $[M_{ring}+H]^+$: 926.5710, found: 926.5690 (-2.2 ppm).

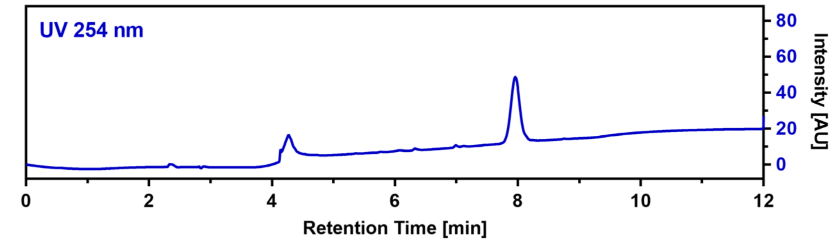

**CalB_R6Pip (41)**

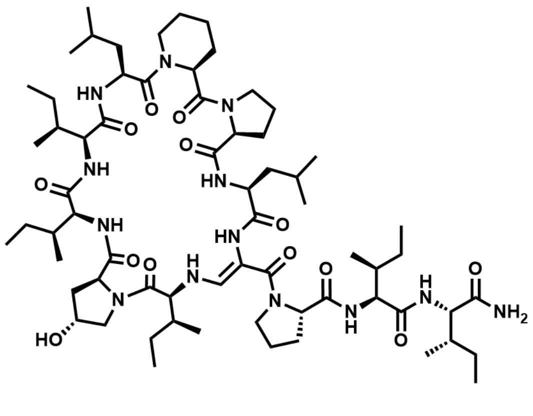

$C_{66}H_{111}N_{13}O_{13}$ M = 1294.69 g/mol

The linear precursor peptide of **41** was prepared via microwave-assisted SPPS using Rink amide PS resin at a scale of 0.1 mmol initial resin capacity. After purification by HPLC (solvent A: $H_2O$, acidified with 0.1% TFA; solvent B: ACN, acidified with 0.1% TFA), 57 mg (44 µmol, 44%) of the peptide precursor was obtained as a white powder. For the oxidation of the serine residue and concomitant cyclization, the peptide was dissolved in 10 mL of ACN and 4 equivalents of DMP were added. The resulting suspension was shaken for 2 hours at RT and immediately purified by HPLC (solvent A: $H_2O$; solvent B: ACN) to give 1.1 mg (0.8 µmol, 2.0%) of **41** in a total yield of 0.8% (calculated in relation to initial resin capacity) as a white powder.

HRMS (ESI): *m/z* calculated for $[M+H]^+$: 1294.8497, found: 1294.8492 (-0.4 ppm); calculated for $[M+2H]^{2+}$: 647.9285, found: 947.9285 (-0.0 ppm); $MS^2$ HRMS (ESI, HCD): *m/z* calculated for ring fragment $[M_{ring}+H]^+$: 954.6023, found: 954.6011 (-1.3 ppm).

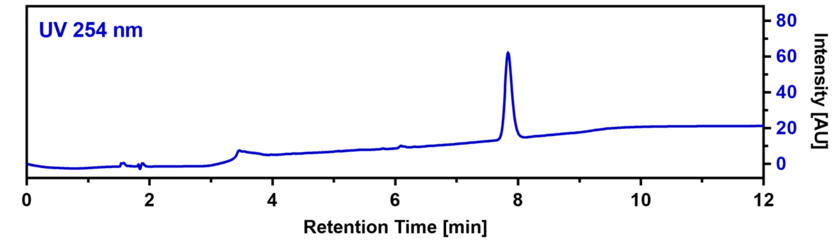

**CalB_R6Nip (42)**

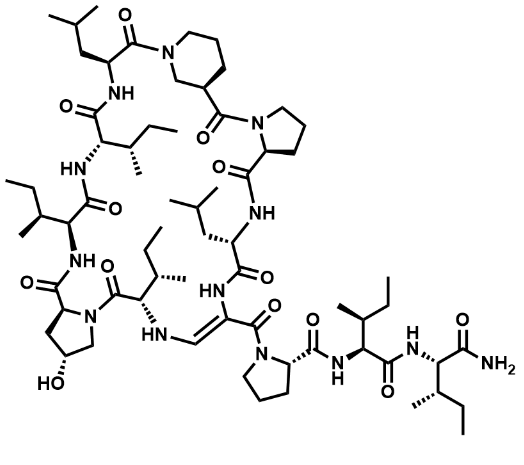

$C_{66}H_{111}N_{13}O_{13}$ M = 1294.69 g/mol

The linear precursor peptide of **42** was prepared via microwave-assisted SPPS using Rink amide PS resin at a scale of 0.1 mmol initial resin capacity. After purification by HPLC (solvent A: $H_2O$, acidified with 0.1% TFA; solvent B: ACN, acidified with 0.1% TFA), 30 mg (23 µmol, 23%) of the peptide precursor was obtained as a white powder. For the oxidation of the serine residue and concomitant cyclization, the peptide was dissolved in 10 mL of ACN and 4 equivalents of DMP were added. The resulting suspension was shaken for 2.5 hours at RT and immediately purified by HPLC (solvent A: $H_2O$; solvent B: ACN) to give 1.1 mg (0.9 µmol, 3.8%) of **42** in a total yield of 0.9% (calculated in relation to initial resin capacity) as a white powder.
HRMS (ESI): *m/z* calculated for $[M+H]^+$: 1294.8497, found: 1294.8494 (-0.2 ppm); calculated for $[M+2H]^{2+}$: 647.9285, found: 647.9282 (-0.5 ppm); $MS^2$ HRMS (ESI, HCD): *m/z* calculated for ring fragment $[M_{ring}+H]^+$: 954.6023, found: 954.6000 (-2.4 ppm).

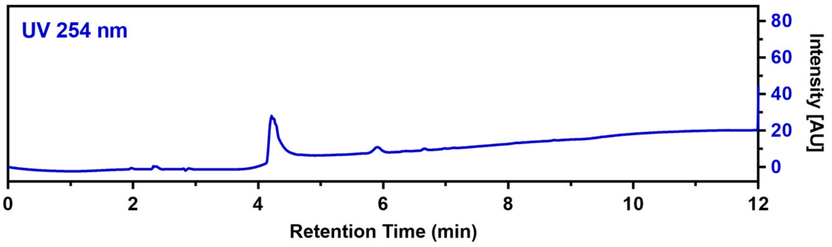

**CalB_R6Oic (43)**

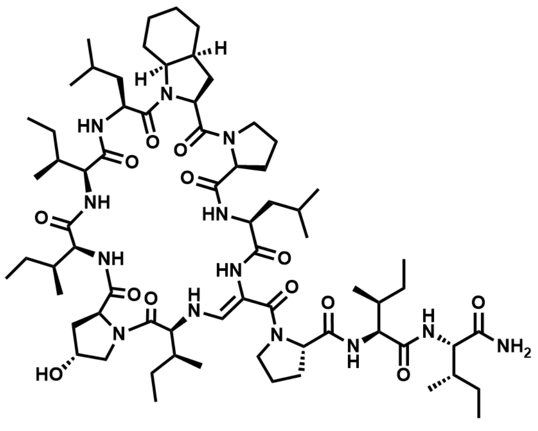

$C_{69}H_{115}N_{13}O_{13}$ M = 1334.76 g/mol

The linear precursor peptide of **43** was prepared via microwave-assisted SPPS using Rink amide PS resin at a scale of 0.1 mmol initial resin capacity. After purification by HPLC (solvent A: $H_2O$, acidified with 0.1% TFA; solvent B: ACN, acidified with 0.1% TFA), 47 mg (35 µmol, 35%) of the peptide precursor was obtained as a white powder. For the oxidation of the serine residue and concomitant cyclization, the peptide was dissolved in 10 mL of ACN and 4 equivalents of DMP were added. The resulting suspension was shaken for 3 hours at RT and immediately purified by HPLC (solvent A: $H_2O$; solvent B: ACN) to give 1.0 mg (0.7 µmol, 2.2%) of **43** in a total yield of 0.7% (calculated in relation to initial resin capacity) as a white powder.
HRMS (ESI): *m/z* calculated for $[M+H]^+$: 1334.8810, found: 1334.8797 (-1.0 ppm); calculated for $[M+2H]^{2+}$: 667.9442, found: 667.9469 (4.0 ppm); $MS^2$ HRMS (ESI, HCD): *m/z* calculated for ring fragment $[M_{ring}+H]^+$: 994.6336, found: 994.6319 (-1.7 ppm).

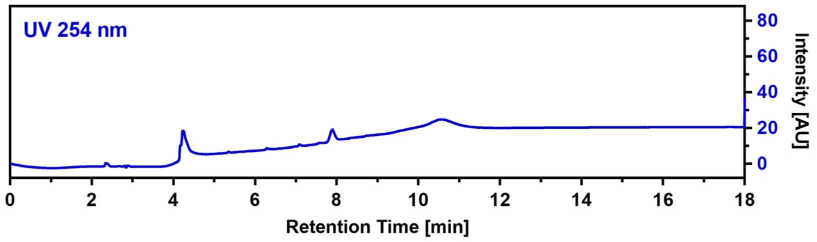

**CalB_R7dPro (44)**

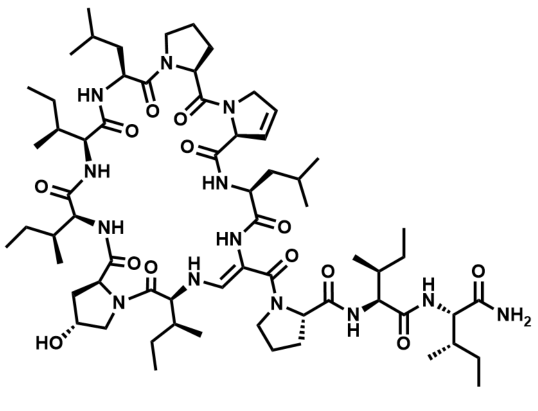

$C_{65}H_{107}N_{13}O_{13}$ M = 1278.65 g/mol

The linear precursor peptide of **44** was prepared via microwave-assisted SPPS using Rink amide PS resin at a scale of 0.1 mmol initial resin capacity. After purification by HPLC (solvent A: $H_2O$, acidified with 0.1% TFA; solvent B: ACN, acidified with 0.1% TFA), 25 mg (19 µmol, 19%) of the peptide precursor was obtained as a white powder. For the oxidation of the serine residue and concomitant cyclization, the peptide was dissolved in 5 mL of ACN and 4 equivalents of DMP were added. The resulting suspension was shaken for 3 hours at RT and immediately purified by HPLC (solvent A: $H_2O$; solvent B: ACN) to give 3.5 mg (2.7 µmol, 14.5%) of **44** in a total yield of 2.7% (calculated in relation to initial resin capacity) as a white powder.

HRMS (ESI): *m/z* calculated for $[M+H]^+$: 1278.8184, found: 1278.8185 (0.1 ppm); calculated for $[M+2H]^{2+}$: 639.9129, found: 639.9124 (-0.8 ppm); $MS^2$ HRMS (ESI, HCD): *m/z* calculated for ring fragment $[M_{ring}+H]^+$: 938.5710, found: 938.5672 (-4.0 ppm).

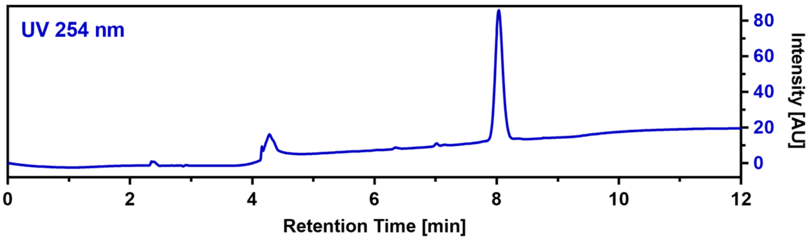

**CalB_R7Aze (45)**

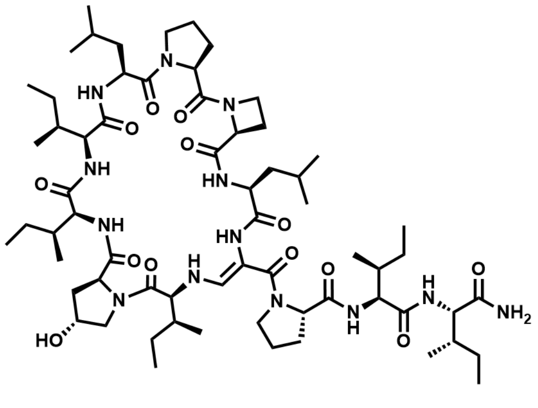

$C_{64}H_{107}N_{13}O_{13}$ M = 1266.64 g/mol

The linear precursor peptide of **45** was prepared via microwave-assisted SPPS using Rink amide PS resin at a scale of 0.1 mmol initial resin capacity. After purification by HPLC (solvent A: $H_2O$, acidified with 0.1% TFA; solvent B: ACN, acidified with 0.1% TFA), 31 mg (24 µmol, 24%) of the peptide precursor was obtained as a white powder. For the oxidation of the serine residue and concomitant cyclization, the peptide was dissolved in 5 mL of ACN and 3 equivalents of DMP were added. The resulting suspension was shaken for 2 hours at RT and immediately purified by HPLC (solvent A: $H_2O$; solvent B: ACN) to give 2.1 mg (1.7 µmol, 7.0%) of **45** in a total yield of 1.7% (calculated in relation to initial resin capacity) as a white powder.

$^1$H NMR (400 MHz, DMSO) δ 8.98 (d, *J* = 7.0 Hz, 1H), 8.84 (d, *J* = 6.6 Hz, 1H), 8.21 (s, 1H), 7.56 (d, *J* = 10.1 Hz, 1H), 7.29 (d, *J* = 7.3 Hz, 1H), 7.20 (d, *J* = 9.8 Hz, 1H), 7.17 (s, 1H), 7.08 (d, *J* = 5.6 Hz, 1H), 7.05 (d, *J* = 8.4 Hz, 1H), 6.95 (s, 1H), 5.86 (dd, *J* = 13.2, 10.0 Hz, 1H), 5.39 (d, *J* = 3.1 Hz, 1H), 4.71-4.63 (m, 2H), 4.59 (dd, *J* = 9.3, 6.1 Hz, 1H), 4.43-4.39 (m, 1H), 4.38-4.32 (m, 1H), 4.26-4.19 (m, 2H), 4.18-4.08 (m, 2H), 4.02-3.92 (m, 2H), 3.87 (t, *J* = 8.4 Hz, 1H), 3.83 (d, *J* = 7.3 Hz, 1H), 3.78 (d, *J* = 11.7 Hz, 1H), 3.64 (dd, *J* = 11.1, 3.6 Hz, 1H), 3.56-3.40 (m, 2H), 3.06 (dd, *J* = 11.1, 6.9 Hz, 1H), 3.02-2.95 (m, 1H), 2.65-2.57 (m, 1H), 2.34-2.23 (m, 2H), 2.11-2.00 (m, 2H), 1.99-1.95 (m, 1H), 1.94-1.82 (m, 6H), 1.82-1.72 (m, 4H), 1.71-1.60 (m, 3H), 1.56-1.37 (m, 8H), 1.35-1.27 (m, 1H), 1.23 (s, 1H), 1.18-1.09 (m, 2H), 1.08-0.96 (m, 3H), 0.94-0.71 (m, 42H); HRMS (ESI): *m/z* calculated for $[M+H]^+$: 1266.8184, found: 1266.8172

(-0.9 ppm); calculated for $[M+2H]^{2+}$: 633.9129, found: 633.9117 (-1.9 ppm); $MS^2$ HRMS (ESI, HCD): *m/z* calculated for ring fragment $[M_{ring}+H]^+$: 926.5710, found: 926.5674 (-3.9 ppm).

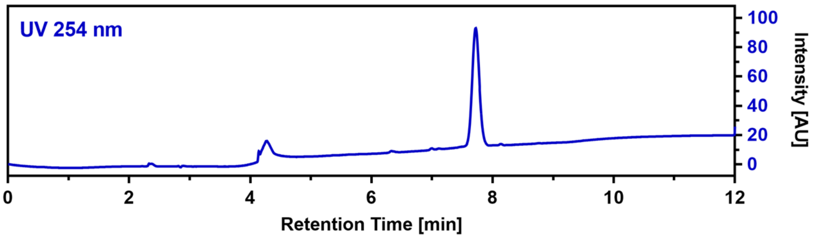

**CalB_R7Pip (46)**

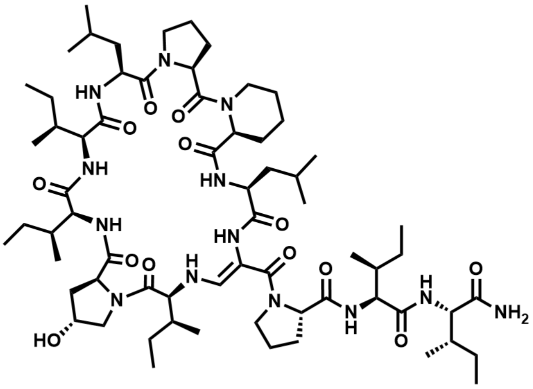

$C_{66}H_{111}N_{13}O_{13}$ M = 1294.69 g/mol

The linear precursor peptide of **46** was prepared via microwave-assisted SPPS using Rink amide PS resin at a scale of 0.1 mmol initial resin capacity. After purification by HPLC (solvent A: $H_2O$, acidified with 0.1% TFA; solvent B: ACN, acidified with 0.1% TFA), 35 mg (27 µmol, 27%) of the peptide precursor was obtained as a white powder. For the oxidation of the serine residue and concomitant cyclization, the peptide was dissolved in 5 mL of ACN and 3 equivalents of DMP were added. The resulting suspension was shaken for 3 hours at RT and immediately purified by HPLC (solvent A: $H_2O$; solvent B: ACN) to give 1.3 mg (1.0 µmol, 3.8%) of **46** in a total yield of 1% (calculated in relation to initial resin capacity) as a white powder.

HRMS (ESI): *m/z* calculated for $[M+H]^+$: 1294.8497, found: 1294.8487 (-0.8 ppm); calculated for $[M+2H]^{2+}$: 647.9285, found: 947.9277 (-1.2 ppm); $MS^2$ HRMS (ESI, HCD): *m/z* calculated for ring fragment $[M_{ring}+H]^+$: 954.6023, found: 954.5999 (-2.5 ppm).

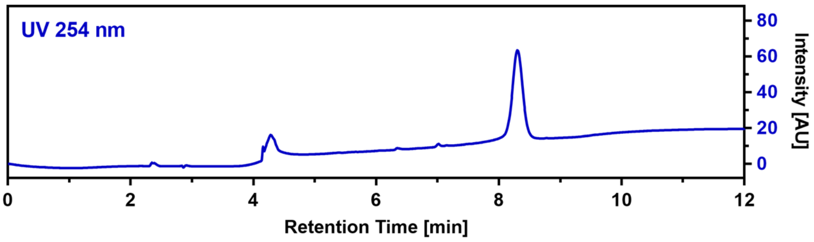

**CalB_R7Nip (47)**

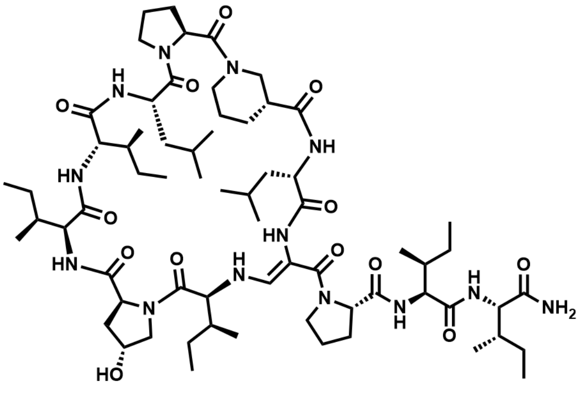

$C_{66}H_{111}N_{13}O_{13}$ M = 1294.69 g/mol

The linear precursor peptide of **47** was prepared via microwave-assisted SPPS using Rink amide PS resin at a scale of 0.1 mmol initial resin capacity. After purification by HPLC (solvent A: $H_2O$, acidified with 0.1% TFA; solvent B: ACN, acidified with 0.1% TFA), 32 mg (24 µmol, 24%) of the peptide precursor was obtained as a white powder. For the oxidation of the serine residue and concomitant cyclization, the peptide was dissolved in 10 mL of ACN and 4 equivalents of DMP were added. The resulting suspension was shaken for 2 hours at RT and immediately purified by HPLC (solvent A: $H_2O$; solvent B: ACN) to give 0.8 mg (0.6 µmol, 2.6%) of **47** in a total yield of 0.6% (calculated in relation to initial resin capacity) as a white powder.

HRMS (ESI): *m/z* calculated for $[M+H]^+$: 1294.8497, found: 1294.8482 (-1.2 ppm); calculated for $[M+2H]^{2+}$: 647.9285, found: 647.9276 (-1.4 ppm); $MS^2$ HRMS (ESI, HCD): *m/z* calculated for ring fragment $[M_{ring}+H]^+$: 954.6023, found: 954.5998 (-2.6 ppm).

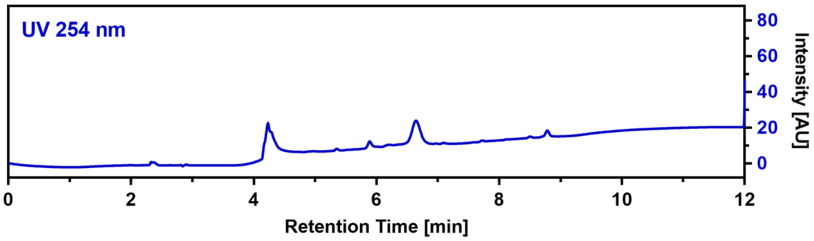

**CalB_R7Oic (48)**

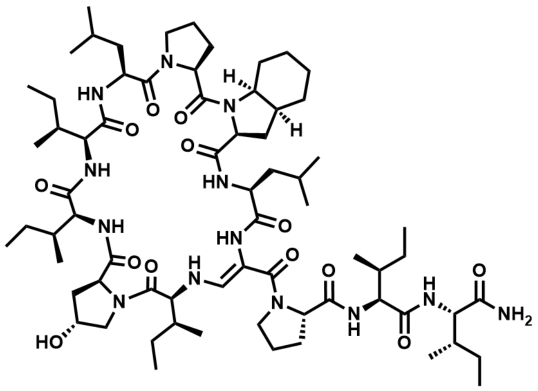

$C_{69}H_{115}N_{13}O_{13}$ M = 1334.76 g/mol

The linear precursor peptide of **48** was prepared via microwave-assisted SPPS using Rink amide PS resin at a scale of 0.1 mmol initial resin capacity. After purification by HPLC (solvent A: $H_2O$, acidified with 0.1% TFA; solvent B: ACN, acidified with 0.1% TFA), 47 mg (34 µmol, 34%) of the peptide precursor was obtained as a white powder. For the oxidation of the serine residue and concomitant cyclization, the peptide was dissolved in 10 mL of ACN and 4 equivalents of DMP were added. The resulting suspension was shaken for 3 hours at RT and immediately purified by HPLC (solvent A: $H_2O$; solvent B: ACN) to give 4.9 mg (3.7 µmol, 10.7%) of **48** in a total yield of 3.7% (calculated in relation to initial resin capacity) as a white powder.

$^1$H NMR (400 MHz, DMSO) δ 9.05 (d, *J* = 7.0 Hz, 1H), 8.95 (d, *J* = 6.3 Hz, 1H), 8.26 (s, 1H), 7.56 (d, *J* = 10.1 Hz, 1H), 7.48 (d, *J* = 6.9 Hz, 1H), 7.25-7.16 (m, 2H), 7.03 (d, *J* = 13.2 Hz, 1H), 6.95 (s, 1H), 6.62 (d, *J* = 10.3 Hz, 1H), 5.92-5.81 (m, 1H), 5.41 (d, *J* = 2.9 Hz, 1H), 4.55 (td, *J* = 10.4, 3.4 Hz, 1H), 4.47-4.37 (m, 3H), 4.29-4.19 (m, 2H), 4.18-4.05 (m, 3H), 3.94 (t, *J* = 10.2 Hz, 1H), 3.89-3.83 (m, 1H), 3.82-3.72 (m, 2H), 3.71-3.60 (m, 2H), 3.49 (t, *J* = 9.1 Hz, 1H), 3.19-3.01 (m, 2H), 2.59-2.52 (m, 1H), 2.40-2.22 (m, 3H), 2.18-2.00 (m, 3H), 1.96-1.82 (m, 6H), 1.80-1.55 (m, 10H), 1.52-1.35 (m, 10H), 1.32-1.10 (m, 7H), 1.07-0.93 (m, 3H), 0.92-0.68 (m, 42H); HRMS (ESI): *m/z* calculated for $[M+H]^+$: 1334.8810, found: 1334.8812 (0.1 ppm); calculated for $[M+2H]^{2+}$: 667.9442, found: 667.9436 (-0.9 ppm); $MS^2$ HRMS (ESI, HCD): *m/z* calculated for ring fragment $[M_{ring}+H]^+$: 994.6336, found: 994.6326 (-1.0 ppm).

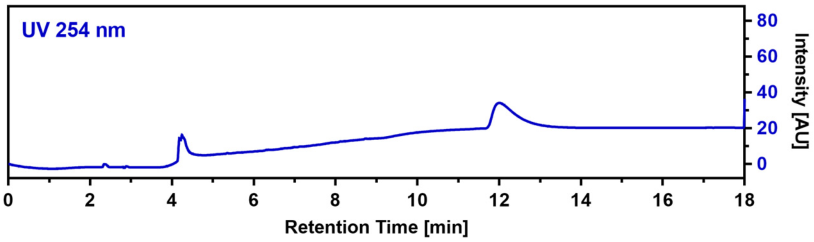

**CalB_R8hLeu (49)**

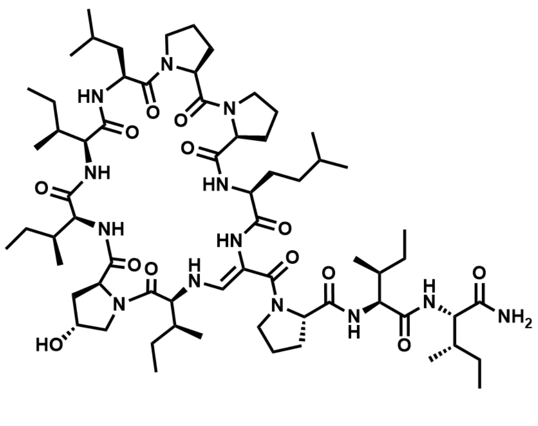

$C_{66}H_{111}N_{13}O_{13}$ M = 1294.69 g/mol

The linear precursor peptide of **49** was prepared via microwave-assisted SPPS using Rink amide PS resin at a scale of 0.1 mmol initial resin capacity. After purification by HPLC (solvent A: $H_2O$, acidified with 0.1% TFA; solvent B: ACN, acidified with 0.1% TFA), 43 mg (33 µmol, 33%) of the peptide precursor was obtained as a white powder. For the oxidation of the serine residue and concomitant cyclization, the peptide was dissolved in 10 mL of ACN and 4 equivalents of DMP were added. The resulting suspension was shaken for 3 hours at RT and immediately purified by HPLC (solvent A: $H_2O$; solvent B: ACN) to give 7.5 mg (5.8 µmol, 17.8%) of **49** in a total yield of 5.8% (calculated in relation to initial resin capacity) as a white powder.

$^1$H NMR (400 MHz, DMSO) δ 9.00 (d, *J* = 6.9 Hz, 1H), 8.87 (d, *J* = 6.3 Hz, 1H), 8.28 (s, 1H), 7.58 (d, *J* = 10.1 Hz, 1H), 7.40 (d, *J* = 6.8 Hz, 1H), 7.24 (d, *J* = 9.7 Hz, 1H), 7.15 (s, 1H), 7.06 (d, *J* = 13.3 Hz, 1H), 6.97 (s, 1H), 6.64 (d, *J* = 9.9 Hz, 1H), 5.70 (dd, *J* = 13.3, 10.1 Hz, 1H), 5.40 (s, 1H), 4.62 – 4.54 (m, 2H), 4.47 – 4.36 (m, 2H), 4.29 – 4.21 (m, 3H), 4.18 – 4.13 (m, 1H), 4.11 – 4.01 (m, 2H), 3.95 (t, *J* = 10.1 Hz, 1H), 3.81-3.72 (m, 2H), 3.69-3.55 (m, 3H), 3.54-3.45 (m, 2H), 3.29-3.22 (m, 1H), 3.20-3.10 (m, 1H), 3.10-3.03 (m, 1H), 2.34-2.19 (m, 3H), 2.12-1.96 (m, 3H), 1.97-1.82 (m, 8H), 1.82-1.55 (m, 5H), 1.53-1.34 (m, 10H), 1.30-1.21 (m, 2H), 1.20-1.10 (m, 3H), 1.07-0.94 (m, 2H), 0.92-0.68 (m, 42H); HRMS (ESI): *m/z* calculated for $[M+H]^+$: 1294.8497, found: 1294.8484 (-1.0 ppm); calculated for $[M+2H]^{2+}$: 647.9285, found: 647.9275 (-1.5 ppm); $MS^2$ HRMS (ESI, HCD): *m/z* calculated for ring fragment $[M_{ring}+H]^+$: 954.6023, found: 954.5991 (-3.4 ppm).

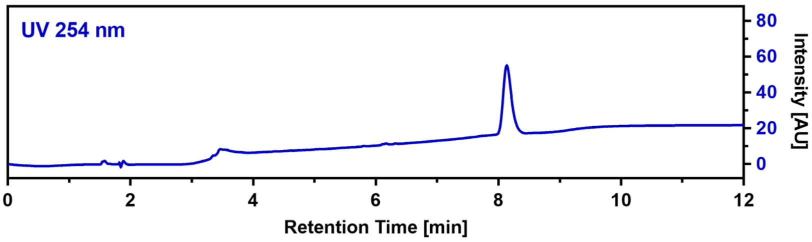

**CalB_R8Aib (50)**

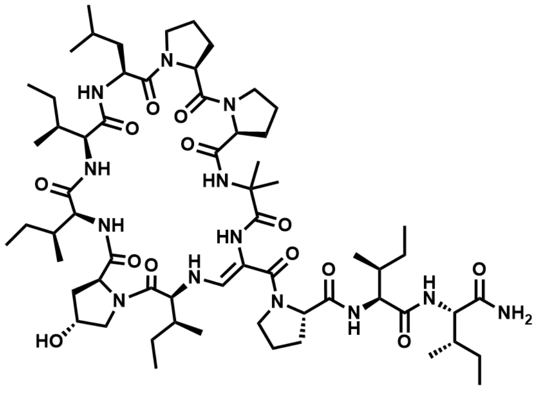

$C_{63}H_{105}N_{13}O_{13}$ M = 1252.61 g/mol

The linear precursor peptide of **50** was prepared via microwave-assisted SPPS using Rink amide PS resin at a scale of 0.1 mmol initial resin capacity. After purification by HPLC (solvent A: $H_2O$, acidified with 0.1% TFA; solvent B: ACN, acidified with 0.1% TFA), 56 mg (44 µmol, 44%) of the peptide precursor was obtained as a white powder. For the oxidation of the serine residue and concomitant cyclization, the peptide was dissolved in 10 mL of ACN and 4 equivalents of DMP were added. The resulting suspension was shaken for 5 hours at RT and immediately purified by HPLC (solvent A: $H_2O$; solvent B: ACN) to give 0.8 mg (0.6 µmol, 1.5%) of **50** in a total yield of 0.6% (calculated in relation to initial resin capacity) as a white powder.

HRMS (ESI): $m/z$ calculated for $[M+H]^+$: 1252.8028, found: 1252.8011 (-1.4 ppm); calculated for $[M+2H]^{2+}$: 626.9050, found: 626.9041 (-1.4 ppm); $MS^2$ HRMS (ESI, HCD): $m/z$ calculated for ring fragment $[M_{ring}+H]^+$: 912.5554, found: 912.5525 (-3.2 ppm).

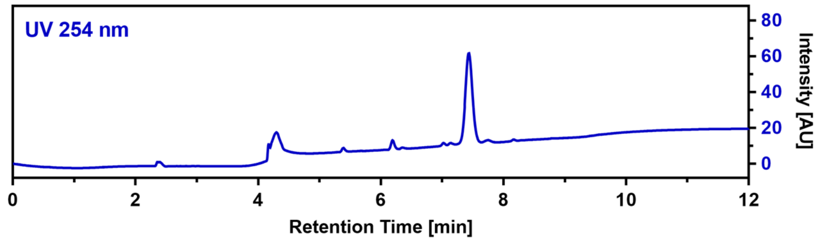

**CalB_C1dPro (51)**

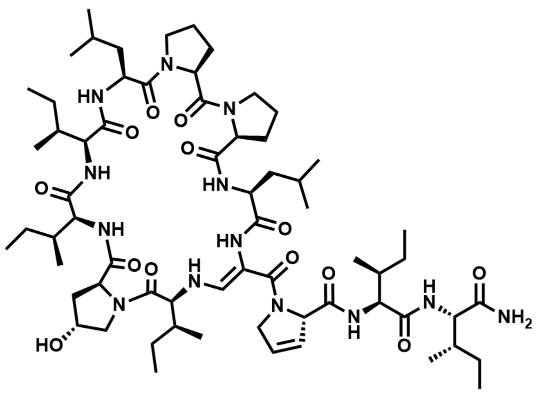

The linear precursor peptide of **51** was prepared via microwave-assisted SPPS using Rink amide PS resin at a scale of 0.1 mmol initial resin capacity. After purification by HPLC (solvent A: $H_2O$, acidified with 0.1% TFA; solvent B: ACN, acidified with 0.1% TFA), 29 mg (22 µmol, 22%) of the peptide precursor was obtained as a white powder. For the oxidation of the serine residue and concomitant cyclization, the peptide was dissolved in 5 mL of ACN and 4 equivalents of DMP were added. The resulting suspension was shaken for 3 hours at RT and immediately purified by HPLC (solvent A: $H_2O$; solvent B: ACN) to give 2.5 mg (2.0 µmol, 8.9%) of **51** in a total yield of 2.0% (calculated in relation to initial resin capacity) as a white powder.

HRMS (ESI): $m/z$ calculated for $[M+H]^+$: 1278.8184, found: 1278.8171 (-1.0 ppm); calculated for $[M+2H]^{2+}$: 639.9129, found: 639.9121 (-1.3 ppm); $MS^2$ HRMS (ESI, HCD): $m/z$ calculated for ring fragment $[M_{ring}+H]^+$: 940.5866, found: 940.5830 (-3.8 ppm).

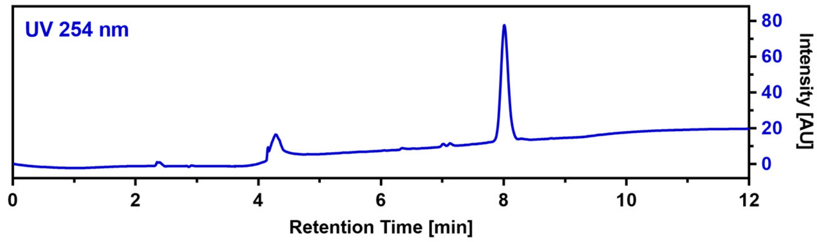

**CalB_C1Aze (52)**

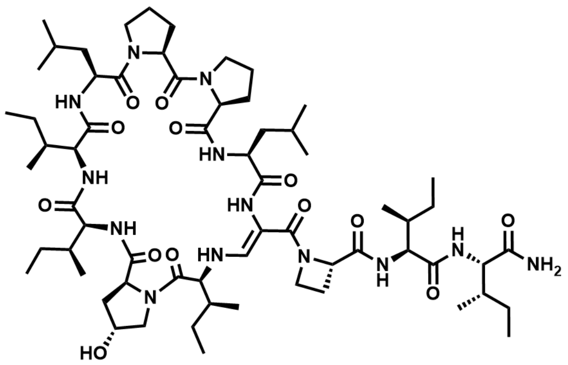

The linear precursor peptide of **52** was prepared via microwave-assisted SPPS using Rink amide PS resin at a scale of 0.1 mmol initial resin capacity. After purification by HPLC (solvent A: $H_2O$, acidified with 0.1% TFA; solvent B: ACN, acidified with 0.1% TFA), 50 mg (39 µmol, 39%) of the peptide precursor was obtained as a white powder. For the oxidation of the serine residue and concomitant cyclization, the peptide was dissolved in 10 mL of ACN and 3 equivalents of DMP were added. The resulting suspension was shaken for 2 hours at RT and immediately purified by HPLC (solvent A: $H_2O$; solvent B: ACN) to give 3.2 mg (2.5 µmol, 6.5%) of **52** in a total yield of 2.5% (calculated in relation to initial resin capacity) as a white powder. $^1H$ NMR (400 MHz, DMSO) δ 8.94 (s, 1H), 8.83 (s, 1H), 7.91 (s, 1H), 7.82 (d, $J$ = 9.3 Hz, 1H), 7.48 (d, $J$ = 9.0 Hz, 1H), 7.38 (d, $J$ = 6.9 Hz, 1H), 7.19 (d, $J$ = 13.3 Hz, 1H), 7.08 (s, 1H), 6.97 (s, 1H), 6.80 (d, $J$ = 10.0 Hz, 1H), 5.66-5.56 (m, 1H), 5.40 (d, $J$ = 3.2 Hz, 1H), 4.65-4.46 (m, 3H), 4.40 (s, 1H), 4.27-4.15 (m, 3H), 4.11-4.05 (m, 1H), 4.05-3.98 (m, 1H), 3.98-3.90 (m, 2H), 3.88-3.73 (m, 3H), 3.72-3.65 (m, 1H), 3.64-3.53 (m, 2H), 3.52-3.42 (m, 1H), 3.27-3.19 (m, 1H), 3.13-3.05 (m, 1H), 2.44-2.32 (m, 1H), 2.31-2.17 (m, 2H), 2.13-1.68 (m, 12H),

1.66-1.33 (m, 11H), 1.32-1.21 (m, 2H), 1.20-0.93 (m, 5H), 0.92-0.71 (m, 42H); HRMS (ESI): *m/z* calculated for $[M+H]^+$: 1266.8184, found: 1266.8185 (0.1 ppm); calculated for $[M+2H]^{2+}$: 633.9129, found: 633.9125 (-0.6 ppm); $MS^2$ HRMS (ESI, HCD): *m/z* calculated for ring fragment $[M_{ring}+H]^+$: 940.5866, found: 940.5855 (-1.2 ppm).

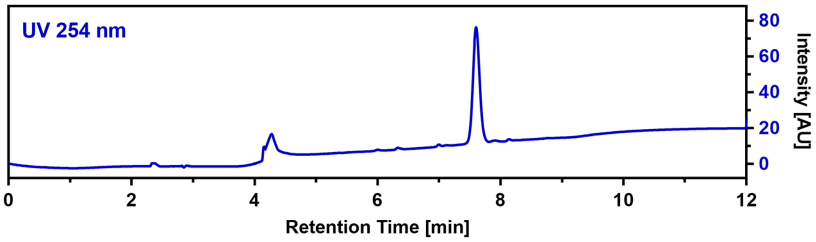

**CalB_C1Pip (53)**

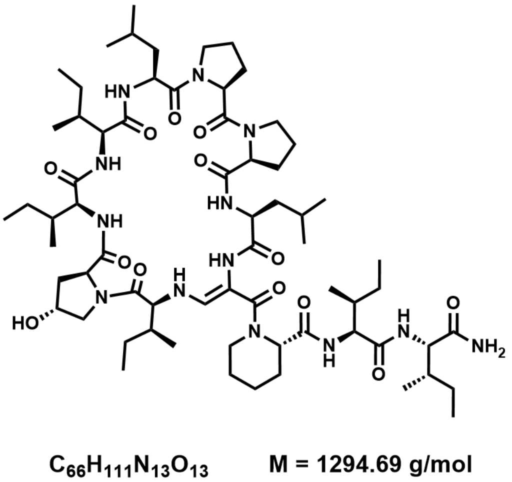

The linear precursor peptide of **53** was prepared via microwave-assisted SPPS using Rink amide PS resin at a scale of 0.1 mmol initial resin capacity. After purification by HPLC (solvent A: $H_2O$, acidified with 0.1% TFA; solvent B: ACN, acidified with 0.1% TFA), 59 mg (45 µmol, 45%) of the peptide precursor was obtained as a white powder. For the oxidation of the serine residue and concomitant cyclization, the peptide was dissolved in 10 mL of ACN and 4 equivalents of DMP were added. The resulting suspension was shaken for 3 hours at RT and immediately purified by HPLC (solvent A: $H_2O$; solvent B: ACN) to give 7.9 mg (6.1 µmol, 13.7%) of **53** in a total yield of 6.1% (calculated in relation to initial resin capacity) as a white powder.

HRMS (ESI): *m/z* calculated for $[M+H]^+$: 1294.8497, found: 1294.8486 (-0.8 ppm); calculated for $[M+2H]^{2+}$: 647.9285, found: 947.9277 (-1.2 ppm); $MS^2$ HRMS (ESI, HCD): *m/z* calculated for ring fragment $[M_{ring}+H]^+$: 940.5866, found: 940.5826 (-4.3 ppm).

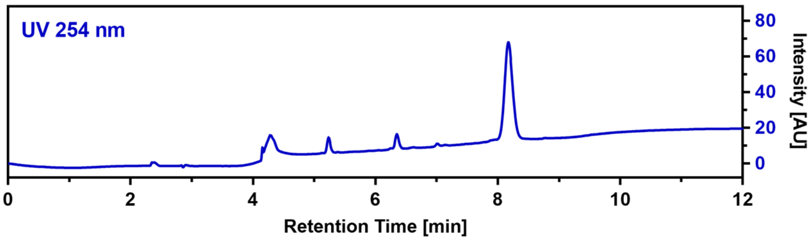

**CalB_C1Nip (54)**

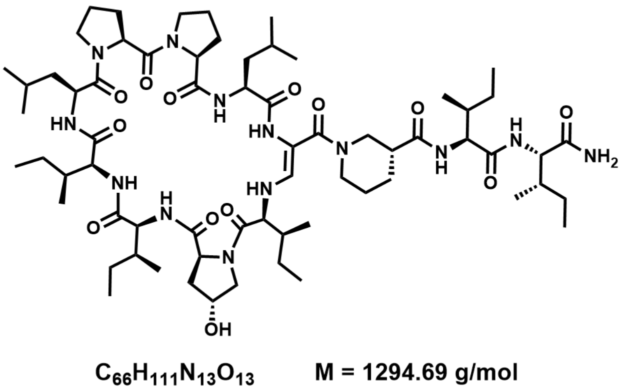

The linear precursor peptide of **54** was prepared via microwave-assisted SPPS using Rink amide PS resin at a scale of 0.1 mmol initial resin capacity. After purification by HPLC (solvent A: $H_2O$, acidified with 0.1% TFA; solvent B: ACN, acidified with 0.1% TFA), 36 mg (27 µmol, 27%) of the peptide precursor was obtained as a white powder. For the oxidation of the serine residue and concomitant cyclization, the peptide was dissolved in 10 mL of ACN and 4 equivalents of DMP were added. The resulting suspension was shaken for 2.5 hours at RT and immediately purified by HPLC (solvent A: $H_2O$; solvent B: ACN) to give 1.4 mg (1.1 µmol, 4.0%) of **54** in a total yield of 1.1% (calculated in relation to initial resin capacity) as a white powder.

HRMS (ESI): *m/z* calculated for $[M+H]^+$: 1294.8497, found: 1294.8499 (0.2 ppm); calculated for $[M+2H]^{2+}$: 647.9285, found: 647.9282 (-0.5 ppm); $MS^2$ HRMS (ESI, HCD): *m/z* calculated for ring fragment $[M_{ring}+H]^+$: 940.5866, found: 940.5836 (-3.2 ppm).

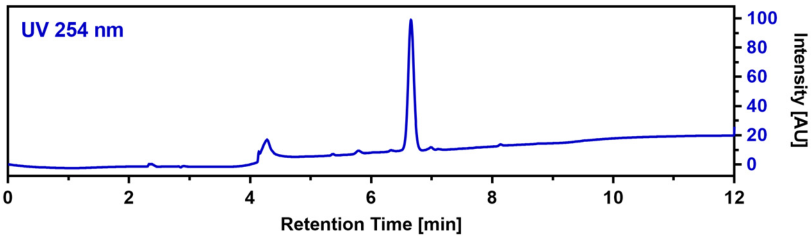

**CalB_C1Oic (55)**

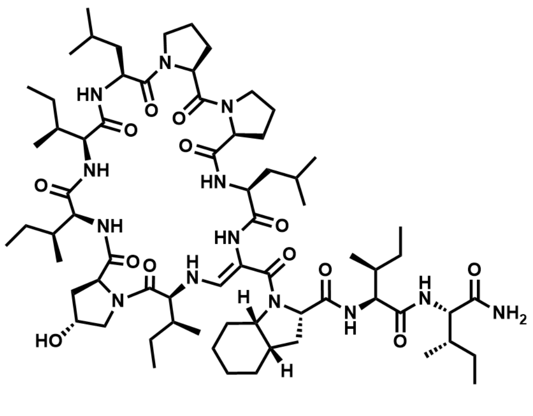

$C_{69}H_{115}N_{13}O_{13}$ M = 1334.76 g/mol

The linear precursor peptide of **55** was prepared via microwave-assisted SPPS using Rink amide PS resin at a scale of 0.1 mmol initial resin capacity. After purification by HPLC (solvent A: $H_2O$, acidified with 0.1% TFA; solvent B: ACN, acidified with 0.1% TFA), 49 mg (36 µmol, 36%) of the peptide precursor was obtained as a white powder. For the oxidation of the serine residue and concomitant cyclization, the peptide was dissolved in 10 mL of ACN and 4 equivalents of DMP were added. The resulting suspension was shaken for 2.5 hours at RT and immediately purified by HPLC (solvent A: $H_2O$; solvent B: ACN) to give 4.1 mg (3.1 µmol, 8.5%) of **55** in a total yield of 3.1% (calculated in relation to initial resin capacity) as a white powder.

HRMS (ESI): *m/z* calculated for $[M+H]^+$: 1334.8810, found: 1334.8815 (0.4 ppm); calculated for $[M+2H]^{2+}$: 667.9442, found: 667.9437 (-0.7 ppm); $MS^2$ HRMS (ESI, HCD): *m/z* calculated for ring fragment $[M_{ring}+H]^+$: 940.5866, found: 940.5851 (-1.6 ppm).

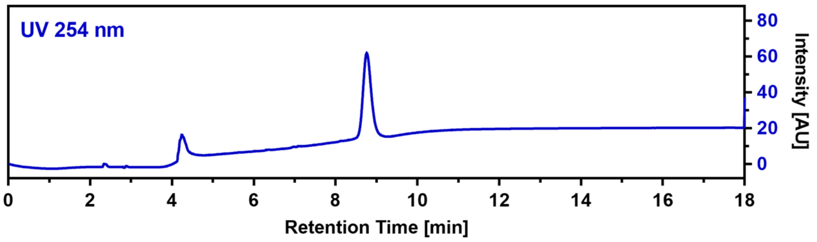

**CalB_C2β-hIle (56)**

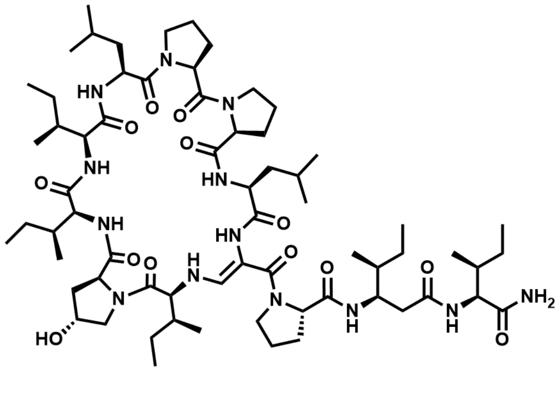

$C_{65}H_{109}N_{13}O_{13}$ M = 1280.67 g/mol

The linear precursor peptide of **56** was prepared via microwave-assisted SPPS using Rink amide PS resin at a scale of 0.1 mmol initial resin capacity. After purification by HPLC (solvent A: $H_2O$, acidified with 0.1% TFA; solvent B: ACN, acidified with 0.1% TFA), 50 mg (38 µmol, 38%) of the peptide precursor was obtained as a white powder. For the oxidation of the serine residue and concomitant cyclization, the peptide was dissolved in 10 mL of ACN and 4 equivalents of DMP were added. The resulting suspension was shaken for 4 hours at RT and immediately purified by HPLC (solvent A: $H_2O$; solvent B: ACN) to give 0.9 mg (0.7 µmol, 1.8%) of **56** in a total yield of 0.7% (calculated in relation to initial resin capacity) as a white powder. HRMS (ESI): *m/z* calculated for $[M+H]^+$:

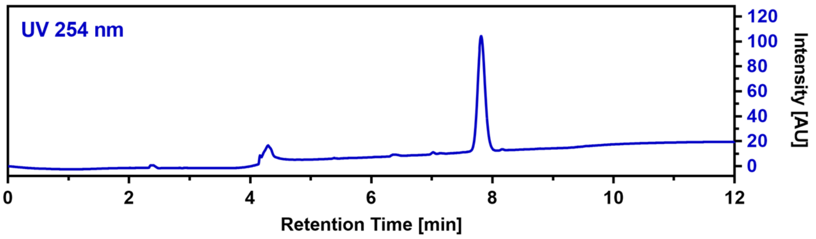

1294.8497, found: 1294.8489 (-0.6 ppm); calculated for $[M+2H]^{2+}$: 647.9285, found: 647.9276 (-1.4 ppm); $MS^2$ HRMS (ESI, HCD): *m/z* calculated for ring fragment $[M_{ring}+H]^+$: 940.5866, found: 940.5830 (-3.8 ppm).

**CalB_C2Aib (57)**

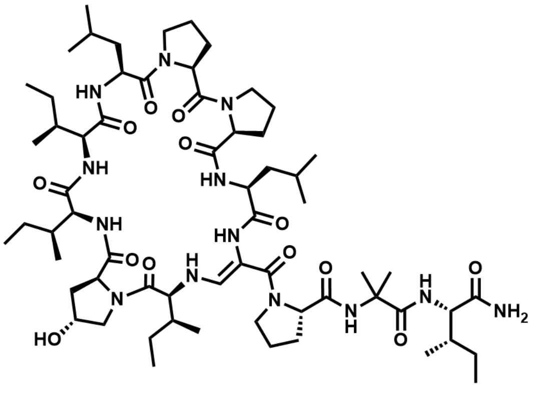

$C_{63}H_{105}N_{13}O_{13}$ M = 1252.61 g/mol

The linear precursor peptide of **57** was prepared via microwave-assisted SPPS using Rink amide PS resin at a scale of 0.1 mmol initial resin capacity. After purification by HPLC (solvent A: $H_2O$, acidified with 0.1% TFA; solvent B: ACN, acidified with 0.1% TFA), 60 mg (47 µmol, 47%) of the peptide precursor was obtained as a white powder. For the oxidation of the serine residue and concomitant cyclization, the peptide was dissolved in 15 mL of ACN and 4 equivalents of DMP were added. The resulting suspension was shaken for 5 hours at RT and immediately purified by HPLC (solvent A: $H_2O$; solvent B: ACN) to give 6.8 mg (5.4 µmol, 11.5%) of **57** in a total yield of 5.4% (calculated in relation to initial resin capacity) as a white powder.

HRMS (ESI): *m/z* calculated for $[M+H]^+$: 1252.8028, found: 1252.8028 (0.0 ppm); calculated for $[M+2H]^{2+}$: 626.9050, found: 626.9048 (-0.3 ppm); $MS^2$ HRMS (ESI, HCD): *m/z* calculated for ring fragment $[M_{ring}+H]^+$: 940.5866, found: 940.5827 (-4.1 ppm).

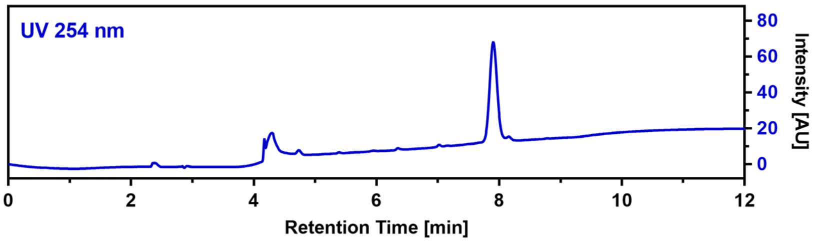

**CalB_C3F (58)**

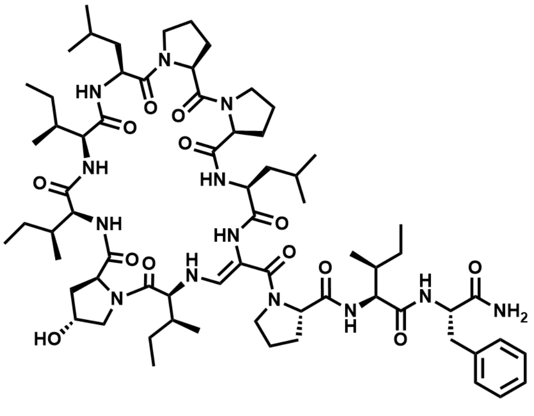

$C_{68}H_{107}N_{13}O_{13}$ M = 1314.68 g/mol

The linear precursor peptide of **58** was prepared via microwave-assisted SPPS using Rink amide PS resin at a scale of 0.1 mmol initial resin capacity. After purification by HPLC (solvent A: $H_2O$, acidified with 0.1% TFA; solvent B: ACN, acidified with 0.1% TFA), 81 mg (61 µmol, 61%) of the peptide precursor was obtained as a white powder. For the oxidation of the serine residue and concomitant cyclization, the peptide was dissolved in 15 mL of ACN and 4 equivalents of DMP were added. The resulting suspension was shaken for 2 hours at RT and immediately purified by HPLC (solvent A: $H_2O$; solvent B: ACN) to give 13.3 mg (10.1 µmol, 16.7%) of **58** in a total yield of 10.1% (calculated in relation to initial resin capacity) as a white powder.

$^1H$ NMR (400 MHz, DMSO) δ 8.98 (d, *J* = 7.0 Hz, 1H), 8.83 (d, *J* = 6.5 Hz, 1H), 8.27 (s, 1H), 7.49-7.42 (m, 2H), 7.39 (d, *J* = 7.0 Hz, 1H), 7.24-7.18 (m, 3H), 7.16 (s, 1H), 7.14-7.06 (m, 4H), 6.73 (d, *J* = 10.0 Hz, 1H), 5.82 (dd, *J* = 13.3, 10.0 Hz, 1H), 5.46-5.28 (m, 1H), 4.67-4.56 (m, 2H), 4.40 (s, 1H), 4.31-4.16 (m, 4H), 4.06 (t, *J* = 8.7 Hz, 1H), 3.91-3.77 (m, 4H), 3.71-3.54 (m, 3H), 3.53-3.44 (m, 2H), 3.25-3.15 (m, 2H), 3.12-3.03 (m, 1H), 2.61-2.54 (m, 1H), 2.35-2.18 (m, 3H), 2.11-2.03 (m, 1H), 2.00-1.85 (m, 8H), 1.81-1.46 (m, 8H), 1.45-1.33 (m, 4H), 1.30-1.19 (m, 1H), 1.17-0.99 (m, 2H), 0.94 (d, *J* = 6.5 Hz, 3H), 0.91-0.72 (m, 30H), 0.67 (t, *J* = 7.4 Hz, 3H), 0.48 (t, *J* = 7.3 Hz, 3H), 0.33 (d, *J* = 6.8 Hz, 3H); HRMS (ESI): *m/z* calculated for $[M+H]^+$: 1314.8184, found: 1314.8175 (-0.7 ppm); calculated for $[M+2H]^{2+}$: 657.9129, found: 657.9122 (-1.1 ppm); $MS^2$ HRMS (ESI, HCD): *m/z* calculated for ring fragment $[M_{ring}+H]^+$: 940.5866, found: 940.5837 (-3.1 ppm).

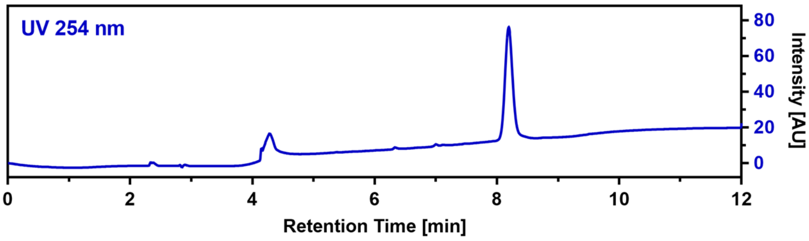

**CalB_C3Aib (59)**

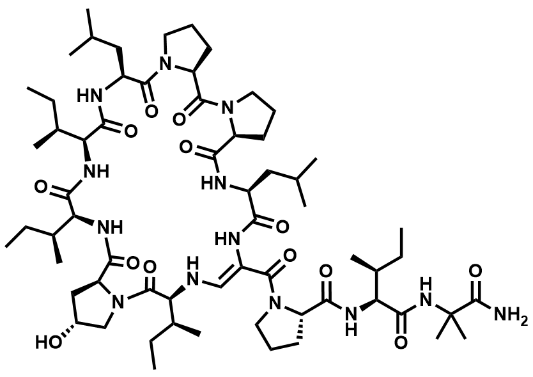

$C_{63}H_{105}N_{13}O_{13}$ M = 1252.61 g/mol

The linear precursor peptide of **59** was prepared via microwave-assisted SPPS using Rink amide PS resin at a scale of 0.1 mmol initial resin capacity. After purification by HPLC (solvent A: $H_2O$, acidified with 0.1% TFA; solvent B: ACN, acidified with 0.1% TFA), 38 mg (30 µmol, 30%) of the peptide precursor was obtained as a white powder. For the oxidation of the serine residue and concomitant cyclization, the peptide was dissolved in 10 mL of ACN and 4 equivalents of DMP were added. The resulting suspension was shaken for 5 hours at RT and immediately purified by HPLC (solvent A: $H_2O$; solvent B: ACN) to give 8.1 mg (6.5 µmol, 21.5%) of **59** in a total yield of 6.5% (calculated in relation to initial resin capacity) as a white powder.

HRMS (ESI): *m/z* calculated for $[M+H]^+$: 1252.8028, found: 1252.8026 (-0.2 ppm); calculated for $[M+2H]^{2+}$: 626.9050, found: 626.9048 (-0.3 ppm); $MS^2$ HRMS (ESI, HCD): *m/z* calculated for ring fragment $[M_{ring}+H]^+$: 940.5866, found: 940.5834 (-3.4 ppm).

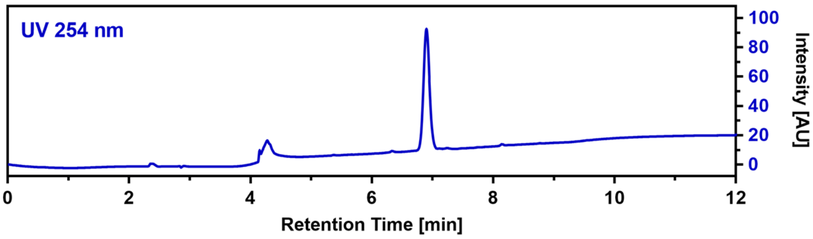

**CalB_C3cPrGly (60)**

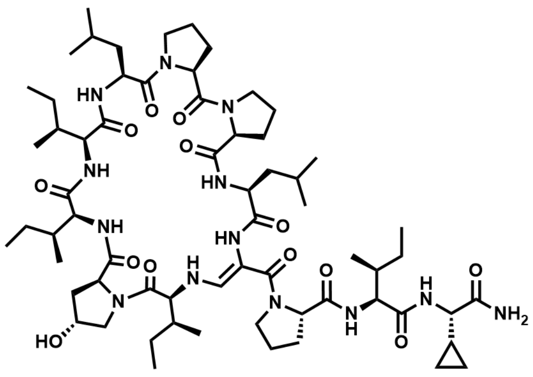

$C_{64}H_{105}N_{13}O_{13}$ M = 1264.62 g/mol

The linear precursor peptide of **60** was prepared via SPPS using Rink amide PS resin at a scale of 0.14 mmol initial resin capacity. After purification by HPLC (solvent A: $H_2O$, acidified with 0.1% TFA; solvent B: ACN, acidified with 0.1% TFA), 69 mg (54 µmol, 39%) of the peptide precursor was obtained as a white powder. For the oxidation of the serine residue and concomitant cyclization, the peptide was dissolved in 15 mL of ACN and 4 equivalents of DMP were added. The resulting suspension was shaken for 4 hours at RT and immediately purified by HPLC (solvent A: $H_2O$; solvent B: ACN) to give 6.2 mg (4.9 µmol, 9.1%) of **60** in a total yield of 3.5% (calculated in relation to initial resin capacity) as a white powder.

$^1$H NMR (700 MHz, DMSO) δ 9.01 (s, 1H), 8.86 (s, 1H), 8.25 (s, 1H), 7.51 (d, *J* = 10.3 Hz, 1H), 7.48 (s, 1H), 7.38 (d, *J* = 6.9 Hz, 1H), 7.22 (s, 1H), 7.07 (d, *J* = 13.3 Hz, 1H), 6.94 (s, 1H), 6.69 (d, *J* = 9.9 Hz, 1H), 5.73 (t, *J* = 10.0 Hz, 1H), 4.63-4.58 (m, 1H), 4.55-4.50 (m, 1H), 4.41 (s, 1H), 4.27-4.21 (m, 3H), 4.14-4.09 (m, 1H), 4.04 (t, *J* = 8.9 Hz, 1H), 3.89 (t, *J* = 10.2 Hz, 1H), 3.83-3.77 (m, 2H), 3.70-3.63 (m, 1H), 3.63-3.56 (m, 2H), 3.51-3.44 (m, 1H), 3.21-3.15 (m, 1H),

2.32-2.19 (m, 3H), 2.11-2.04 (m, 1H), 2.01-1.96 (m, 1H), 1.94-1.82 (m, 8H), 1.81-1.49 (m, 12H), 1.47-1.34 (m, 8H), 1.29-1.21 (m, 2H), 1.19-1.10 (m, 2H), 1.09-0.91 (m, 5H), 0.91-0.70 (m, 36H); HRMS (ESI): *m/z* calculated for $[M+H]^+$: 1264.8028, found: 1264.8015 (-1.0 ppm); calculated for $[M+2H]^{2+}$: 632.9050, found: 632.9043 (-1.1 ppm); $MS^2$ HRMS (ESI, HCD): *m/z* calculated for ring fragment $[M_{ring}+H]^+$: 940.5866, found: 940.5845 (-2.2 ppm).

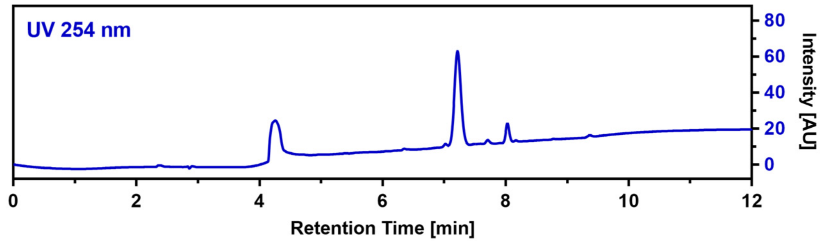

**CalB_C3Tle (61)**

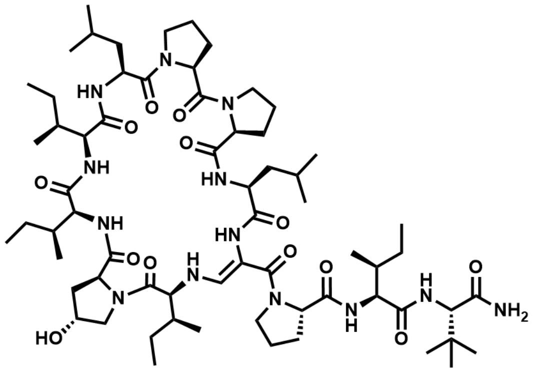

$C_{65}H_{109}N_{13}O_{13}$ M = 1280.67 g/mol

The linear precursor peptide of **61** was prepared via SPPS using Rink amide PS resin at a scale of 0.14 mmol initial resin capacity. After purification by HPLC (solvent A: $H_2O$, acidified with 0.1% TFA; solvent B: ACN, acidified with 0.1% TFA), 51 mg (39 µmol, 28%) of the peptide precursor was obtained as a white powder. For the oxidation of the serine residue and concomitant cyclization, the peptide was dissolved in 15 mL of ACN and 4 equivalents of DMP were added. The resulting suspension was shaken for 4 hours at RT and immediately purified by HPLC (solvent A: $H_2O$; solvent B: ACN) to give 6.4 mg (5.0 µmol, 12.7%) of **61** in a total yield of 3.6% (calculated in relation to initial resin capacity) as a white powder.

$^1$H NMR (400 MHz, DMSO) δ 9.00 (d, *J* = 6.9 Hz, 1H), 8.85 (s, 1H), 8.20 (s, 1H), 7.65 (d, *J* = 9.6 Hz, 1H), 7.37 (d, *J* = 7.1 Hz, 1H), 7.16 (d, *J* = 9.8 Hz, 1H), 6.98 (d, *J* = 14.3 Hz, 2H), 6.89 (s, 1H), 6.66 (d, *J* = 9.8 Hz, 1H), 5.63 (t, *J* = 11.6 Hz, 1H), 5.39 (s, 1H), 4.66-4.56 (m, 1H), 4.55-4.44 (m, 1H), 4.40 (s, 1H), 4.32-4.19 (m, 3H), 4.19-4.10 (m, 1H), 4.08-3.98 (m, 3H), 3.88-3.71 (m, 3H), 3.70-3.55 (m, 4H), 3.33-3.24 (m, 1H), 3.22-3.13 (m, 1H), 3.12-3.04 (m, 1H), 2.35-2.15 (m, 3H), 2.11-2.00 (m, 2H), 1.97-1.81 (m, 8H), 1.77-1.56 (m, 6H), 1.54-1.35 (m, 8H), 1.31-1.20 (m, 1H), 1.19-1.09 (m, 2H), 1.09-0.96 (m, 2H), 0.94-0.71 (m, 45H); HRMS (ESI): *m/z* calculated for $[M+H]^+$: 1280.8341, found: 1280.8334 (-0.5 ppm); calculated for $[M+2H]^{2+}$: 640.9207, found: 640.9203(-0.6 ppm); $MS^2$ HRMS (ESI, HCD): *m/z* calculated for ring fragment $[M_{ring}+H]^+$: 940.5866, found: 940.5834 (-3.4 ppm).

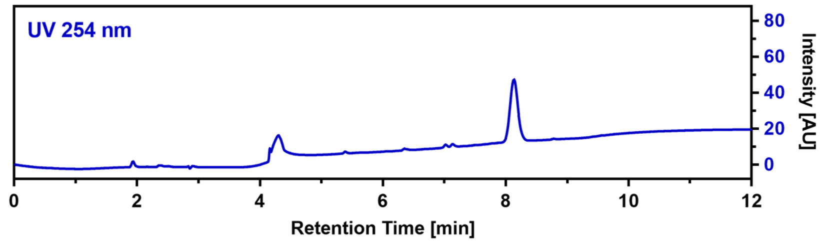

#### CalB_C3Bpa (62)

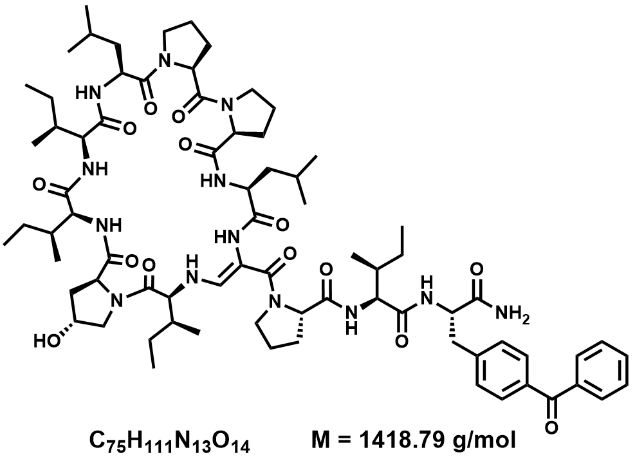

$C_{75}H_{111}N_{13}O_{14}$ M = 1418.79 g/mol

The linear precursor peptide of **62** was prepared via microwave-assisted SPPS using Rink amide PS resin at a scale of 0.14 mmol initial resin capacity. After purification by HPLC (solvent A: $H_2O$, acidified with 0.1% TFA; solvent B: ACN, acidified with 0.1% TFA), 65 mg (45 µmol, 32%) of the peptide precursor was obtained as a white powder. For the oxidation of the serine residue and concomitant cyclization, the peptide was dissolved in 15 mL of ACN and 4 equivalents of DMP were added. The resulting suspension was shaken for 4 hours at RT and immediately purified by HPLC (solvent A: $H_2O$; solvent B: ACN) to give 3.1 mg (2.2 µmol, 4.8%) of **62** in a total yield of 1.6% (calculated in relation to initial resin capacity) as a white powder.

$^1$H NMR (400 MHz, DMSO) δ 8.98 (s, 1H), 8.80 (s, 1H), 8.18 (s, 1H), 7.66-7.59 (m, 4H), 7.57-7.46 (m, 5H), 7.44-7.38 (m, 2H), 7.35 (d, *J* = 7.0 Hz, 1H), 7.31 (s, 1H), 7.20 (d, *J* = 13.3 Hz, 1H), 6.93 (s, 1H), 6.67 (d, *J* = 10.0 Hz, 1H), 5.78 (t, *J* = 11.6 Hz, 1H), 5.36 (s, 1H), 4.63-4.57 (m, 1H), 4.56-4.49 (m, 1H), 4.39 (s, 1H), 4.29-4.16 (m, 5H), 4.04 (t, *J* = 8.7 Hz, 1H), 3.91-3.83 (m, 1H), 3.82 – 3.75 (m, 2H), 3.66 – 3.46 (m, 5H), 3.32 – 3.26 (m, 2H), 3.25 – 3.16 (m, 1H), 3.09 – 3.02 (m, 1H), 2.82 (t, *J* = 12.8 Hz, 1H), 2.35-2.17 (m, 4H), 2.07 (q, *J* = 7.0 Hz, 2H), 1.96-1.79 (m, 7H), 1.75-1.49 (m, 6H), 1.46-1.27 (m, 6H), 1.26-0.98 (m, 7H), 0.85-0.67 (m, 30H), 0.47 (d, *J* = 6.8 Hz, 3H), 0.31 (t, *J* = 7.3 Hz, 3H); HRMS (ESI): *m/z* calculated for $[M+H]^+$: 1418.8446, found: 1418.8438 (-0.6 ppm); calculated for $[M+2H]^{2+}$: 709.9260, found: 709.9253 (-1.0 ppm); MS$^2$ HRMS (ESI, HCD): *m/z* calculated for ring fragment $[M_{ring}+H]^+$: 940.5866, found: 940.5846 (-2.1 ppm).

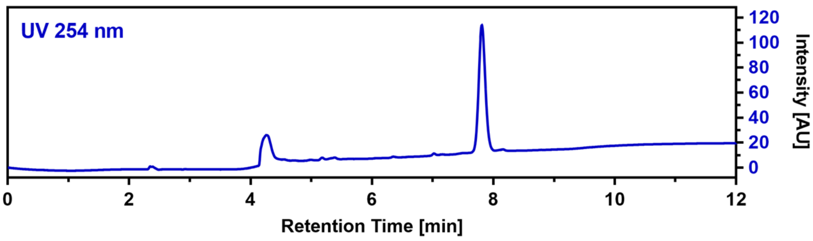

#### CalA_R2P (63)

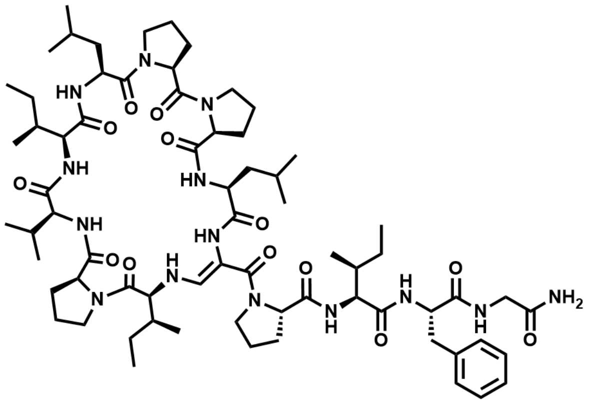

$C_{69}H_{108}N_{14}O_{13}$ M = 1341.71 g/mol

The linear precursor peptide of **63** was prepared via SPPS using Rink amide PS resin at a scale of 0.1 mmol initial resin capacity and purified by HPLC (solvent A: $H_2O$, acidified with 0.1% TFA; solvent B: ACN, acidified with 0.1% TFA). The obtained peptide was then dissolved in 10 mL of ACN and 4 equivalents DMP were added. The resulting suspension was shaken for one hour at RT and immediately purified by HPLC (solvent A: $H_2O$; solvent B: ACN) to give 36.5 mg (27.2 µmol) of **63** in a total yield of 27.2% (calculated in relation to initial resin capacity) as a white powder.

$^1$H NMR (400 MHz, DMSO) δ 8.94 (d, *J* = 7.0 Hz, 1H), 8.69 (d, *J* = 6.8 Hz, 1H), 8.24 (s, 1H), 8.10 (t, *J* = 6.2 Hz, 1H), 7.48 (d, *J* = 8.6 Hz, 1H), 7.44 (d, *J* = 7.0 Hz, 1H), 7.35 (d, *J* = 13.3 Hz, 1H), 7.29 (d, *J* = 8.6 Hz, 1H), 7.23-7.16 (m, 3H), 7.16-7.07 (m, 3H), 7.01 (s, 1H), 6.73 (d, *J* = 9.9 Hz, 1H), 5.84 (dd, *J* = 13.3, 10.1 Hz, 1H), 4.66-4.55 (m, 2H), 4.32-4.19 (m, 3H), 4.14-4.06 (m, 2H), 4.06-3.99 (m, 1H), 3.93-3.82 (m, 2H), 3.82-3.75 (m, 2H), 3.68-3.54 (m, 3H), 3.53-3.42 (m, 3H), 3.26-3.17 (m, 1H), 3.11 (dd, *J* = 13.8, 2.9 Hz, 1H), 3.00 (dd,

$J$ = 10.9, 7.0 Hz, 1H), 2.73-2.58 (m, 2H), 2.35-2.17 (m, 4H), 2.06-1.95 (m, 3H), 1.95-1.81 (m, 7H), 1.80-1.72 (m, 2H), 1.71-1.55 (m, 6H), 1.53-1.44 (m, 2H), 1.44-1.33 (m, 3H), 1.28-1.20 (m, 1H), 1.20-1.09 (m, 2H), 1.08-1.00 (m, 1H), 0.95 (d, $J$ = 6.4 Hz, 3H), 0.90-0.75 (m, 24H), 0.73-0.67 (m, 4H), 0.44-0.36 (m, 6H); HRMS (ESI): *m/z* calculated for $[M+H]^+$: 1341.8293, found: 1341.8319 (1.9 ppm); calculated for $[M+2H]^{2+}$: 671.4183, found: 671.4186 (0.4 ppm); $MS^2$ HRMS (ESI, HCD): *m/z* calculated for ring fragment $[M_{ring}+H]^+$: 910.5761, found:910.5755 (-0.7 ppm).

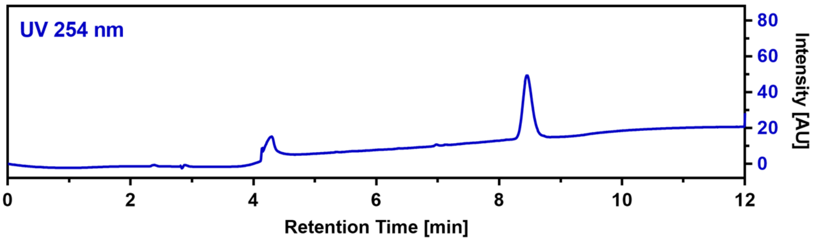

**CalA_R3Abu (64)**

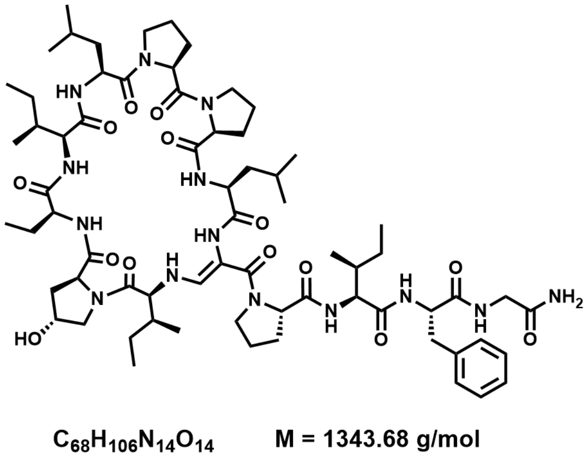

The linear precursor peptide of **64** was prepared via SPPS using Rink amide PS resin at a scale of 0.14 mmol initial resin capacity. After purification by HPLC (solvent A: $H_2O$, acidified with 0.1% TFA; solvent B: ACN, acidified with 0.1% TFA), 90 mg (66 µmol, 47%) of the peptide precursor was obtained as a white powder. For the oxidation of the serine residue and concomitant cyclization, the peptide was dissolved in 20 mL of ACN and 4 equivalents of DMP were added. The resulting suspension was shaken for 5 hours at RT and immediately purified by HPLC (solvent A: $H_2O$; solvent B: ACN) to give 21.1 mg (15.7 µmol, 23.7%) of **64** in a total yield of 11.2% (calculated in relation to initial resin capacity) as a white powder.

$^1H$ NMR (400 MHz, DMSO) δ 8.83 (d, $J$ = 6.8 Hz, 1H), 8.31 (s, 1H), 8.17-8.08 (m, 2H), 7.55 (d, $J$ = 7.3 Hz, 1H), 7.48 (d, $J$ = 8.7 Hz, 1H), 7.35 (d, $J$ = 13.3 Hz, 1H), 7.28-7.21 (m, 2H), 7.21-7.10 (m, 5H), 7.00-6.95 (m, 1H), 6.76 (d, $J$ = 9.9 Hz, 1H), 5.80-5.70 (m, 1H), 5.37 (d, $J$ = 3.0 Hz, 1H), 4.66 (q, $J$ = 6.9 Hz, 1H), 4.58 (td, $J$ = 10.3, 4.2 Hz, 1H), 4.41 (s, 1H), 4.32-4.20 (m, 4H), 4.12 (t, $J$ = 10.2 Hz, 1H), 4.03 (t, $J$ = 8.7 Hz, 1H), 3.98-3.87 (m, 2H), 3.82-3.68 (m, 3H), 3.61-3.39 (m, 6H), 3.27-3.18 (m, 1H), 3.15-3.07 (m, 1H), 2.68-2.58 (m, 1H), 2.35-2.31 (m, 1H), 2.30-2.21 (m, 2H), 2.14-2.01 (m, 3H), 1.98-1.78 (m, 9H), 1.75-1.53 (m, 9H), 1.45-1.28 (m, 5H), 1.20-0.99 (m, 3H), 0.96 (d, $J$ = 6.5 Hz, 3H), 0.90 (d, $J$ = 6.4 Hz, 3H), 0.86-0.75 (m, 18H), 0.71 (t, $J$ = 7.4 Hz, 3H), 0.42 (t, $J$ = 7.3 Hz, 3H), 0.38 (d, $J$ = 6.8 Hz, 3H); HRMS (ESI): *m/z* calculated for $[M+H]^+$: 1343.8086, found: 1343.8087 (0.1 ppm); calculated for $[M+2H]^{2+}$: 672.4079, found: 672.4073 (-0.9 ppm); $MS^2$ HRMS (ESI, HCD): *m/z* calculated for ring fragment $[M_{ring}+H]^+$: 912.5554, found: 912.5527 (-3.0 ppm).

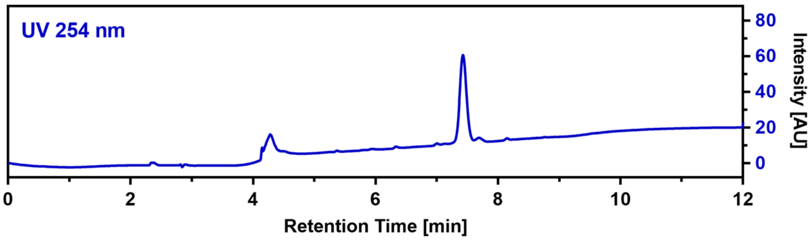

#### CalA_R3Chg (65)

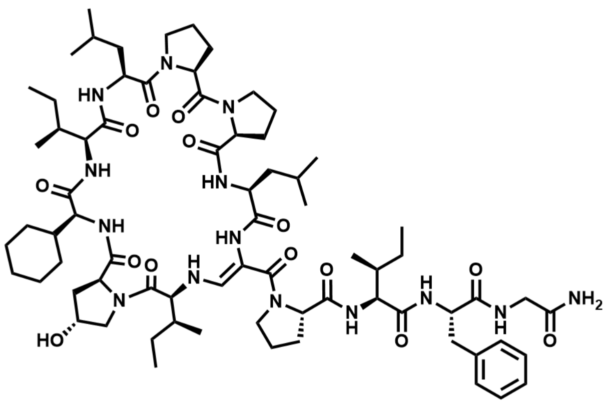

$C_{72}H_{112}N_{14}O_{14}$ M = 1397.77 g/mol

The linear precursor peptide of **65** was prepared via SPPS using Rink amide PS resin at a scale of 0.14 mmol initial resin capacity. After purification by HPLC (solvent A: $H_2O$, acidified with 0.1% TFA; solvent B: ACN, acidified with 0.1% TFA), 40 mg (28 µmol, 20%) of the peptide precursor was obtained as a white powder. For the oxidation of the serine residue and concomitant cyclization, the peptide was dissolved in 10 mL of ACN and 4 equivalents of DMP were added. The resulting suspension was shaken for 5 hours at RT and immediately purified by HPLC (solvent A: $H_2O$; solvent B: ACN) to give 9.4 mg (6.7 µmol, 23.8%) of **65** in a total yield of 4.8% (calculated in relation to initial resin capacity) as a white powder. $^1$H NMR (400 MHz, DMSO) δ 9.01 (d, $J$ = 7.0 Hz, 1H), 8.83 (d, $J$ = 6.6 Hz, 1H), 8.28 (s, 1H), 8.18 (t, $J$ = 6.1 Hz, 1H), 7.46 (d, $J$ = 8.8 Hz, 1H), 7.42 (d, $J$ = 7.0 Hz, 1H), 7.34 (d, $J$ = 13.3 Hz, 1H), 7.29-7.23 (m, 2H), 7.19-7.09 (m, 5H), 6.97 (s, 1H), 6.71 (d, $J$ = 10.0 Hz, 1H), 5.93-5.82 (m, 1H), 5.38 (d, $J$ = 3.0 Hz, 1H), 4.64-4.55 (m, 2H), 4.40 (s, 1H), 4.30-4.20 (m, 4H), 4.10-4.00 (m, 2H), 3.93 (d, $J$ = 7.2 Hz, 1H), 3.89 (d, $J$ = 7.0 Hz, 1H), 3.81-3.74 (m, 3H), 3.70-3.55 (m, 4H), 3.35-3.25 (m, 1H), 3.24-3.15 (m, 1H), 3.12-3.01 (m, 2H), 2.61 (t, $J$ = 12.9 Hz, 1H), 2.55-2.52 (m, 1H), 2.40-2.19 (m, 4H), 2.12-2.03 (m, 1H), 2.01-1.94 (m, 1H), 1.93-1.80 (m, 8H), 1.79-1.71 (m, 1H), 1.71-1.53 (m, 10H), 1.52-1.33 (m, 7H), 1.22 (s, 1H), 1.19-0.98 (m, 7H), 0.94 (d, $J$ = 6.5 Hz, 3H), 0.87-0.74 (m, 18H), 0.69 (t, $J$ = 7.3 Hz, 3H), 0.42 (t, $J$ = 7.3 Hz, 3H), 0.37 (d, $J$ = 6.8 Hz, 3H); HRMS (ESI): *m/z* calculated for $[M+H]^+$: 1397.8555, found: 1394.8557 (0.1 ppm); calculated for $[M+2H]^{2+}$: 699.4314, found: 699.4307 (-1.0 ppm); $MS^2$ HRMS (ESI, HCD): *m/z* calculated for ring fragment $[M_{ring}+H]^+$: 966.6023, found: 966.6001 (-2.3 ppm).

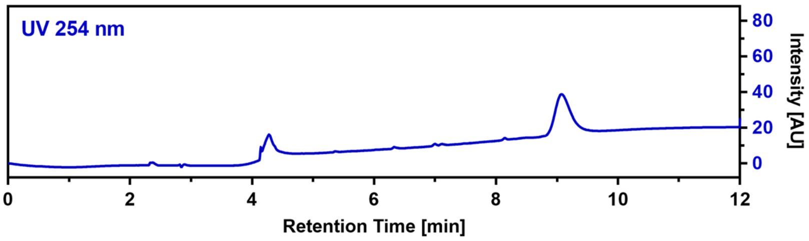

#### CalA_R3Cpg (66)

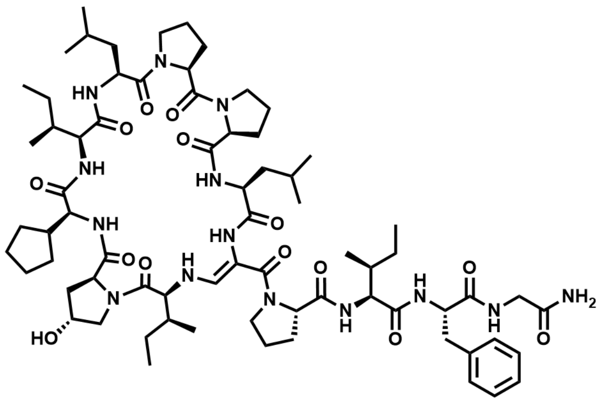

$C_{71}H_{110}N_{14}O_{14}$ M = 1383.75 g/mol

The linear precursor peptide of **66** was prepared via SPPS using Rink amide PS resin at a scale of 0.14 mmol initial resin capacity. After purification by HPLC (solvent A: $H_2O$, acidified with 0.1% TFA; solvent B: ACN, acidified with 0.1% TFA), 19 mg (14 µmol, 10%) of the peptide precursor was obtained as a white powder. For the oxidation of the serine residue and concomitant cyclization, the peptide was dissolved in 10 mL of ACN and 4 equivalents of DMP were added. The resulting suspension was shaken for 6 hours at RT and immediately purified by HPLC (solvent A: $H_2O$; solvent B: ACN) to give 3.5 mg (2.5 µmol, 18.5%) of **66** in a total yield of 1.0% (calculated in relation to initial resin capacity) as a white powder.

$^{1}$H NMR (400 MHz, DMSO) δ 9.08 (d, *J* = 6.9 Hz, 1H), 8.76 (d, *J* = 7.0 Hz, 1H), 8.29 (s, 1H), 8.17 (t, *J* = 6.2 Hz, 1H), 7.47 (d, *J* = 8.7 Hz, 1H), 7.42 (d, *J* = 7.0 Hz, 1H), 7.34 (d, *J* = 13.3 Hz, 1H), 7.28-7.23 (m, 2H), 7.20-7.09 (m, 5H), 6.99-6.95 (m, 1H), 6.72 (d, *J* = 10.0 Hz, 1H), 5.91-5.82 (m, 1H), 5.37 (d, *J* = 3.1 Hz, 1H), 4.65-4.57 (m, 2H), 4.40 (s, 1H), 4.31-4.21 (m, 4H), 4.12-4.01 (m, 2H), 3.96-3.87 (m, 1H), 3.85-3.74 (m, 3H), 3.72-3.66 (m, 1H), 3.63-3.55 (m, 2H), 3.53-3.44 (m, 3H), 3.25-3.16 (m, 1H), 3.13-3.05 (m, 2H), 2.97-2.86 (m, 1H), 2.62 (t, *J* = 12.9 Hz, 1H), 2.55-2.51 (m, 1H), 2.34-2.20 (m, 3H), 2.10-2.02 (m, 1H), 2.02-1.96 (m, 1H), 1.94-1.82 (m, 8H), 1.79-1.70 (m, 1H), 1.69-1.52 (m, 10H), 1.51-1.35 (m, 7H), 1.27-1.20 (m, 3H), 1.19-1.11 (m, 2H), 1.08-1.00 (m, 1H), 0.95 (d, *J* = 6.5 Hz, 3H), 0.89-0.75 (m, 18H), 0.70 (t, *J* = 7.4 Hz, 3H), 0.42 (t, *J* = 7.3 Hz, 3H), 0.38 (d, *J* = 6.8 Hz, 3H); HRMS (ESI): *m/z* calculated for $[M+H]^{+}$: 1383.8399, found: 1383.8401 (0.1 ppm); calculated for $[M+2H]^{2+}$: 692.4236, found: 692.4231 (-0.7 ppm); MS$^2$ HRMS (ESI, HCD): *m/z* calculated for ring fragment $[M_{ring}+H]^{+}$: 952.5867, found: 952.5858 (-0.9 ppm).

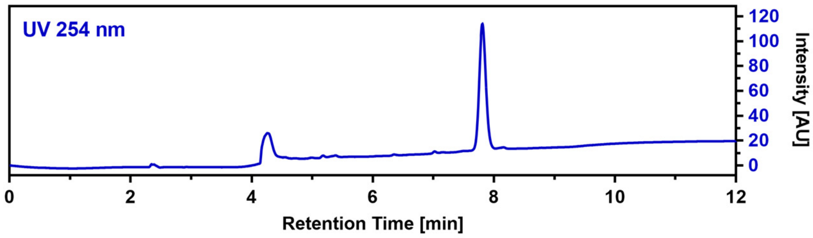

**CalA_R3L (67)**

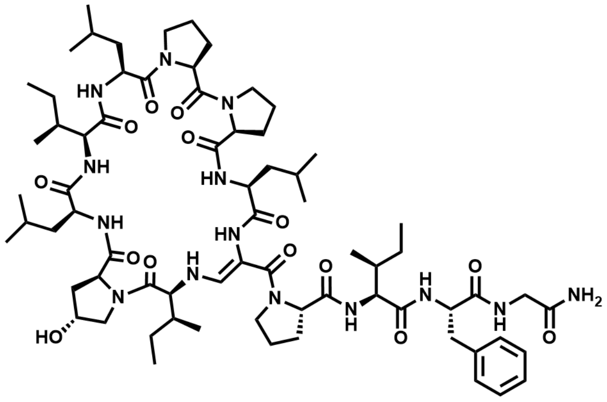

$C_{70}H_{110}N_{14}O_{14}$ M = 1371.73 g/mol

The linear precursor peptide of **67** was prepared via SPPS using Rink amide PS resin at a scale of 0.14 mmol initial resin capacity. After purification by HPLC (solvent A: $H_2O$, acidified with 0.1% TFA; solvent B: ACN, acidified with 0.1% TFA), 26 mg (19 µmol, 13%) of the peptide precursor was obtained as a white powder. For the oxidation of the serine residue and concomitant cyclization, the peptide was dissolved in 10 mL of ACN and 4 equivalents of DMP were added. The resulting suspension was shaken for 5 hours at RT and immediately purified by HPLC (solvent A: $H_2O$; solvent B: ACN) to give 6.4 mg (4.7 µmol, 25.4%) of **67** in a total yield of 3.4% (calculated in relation to initial resin capacity) as a white powder. $^{1}$H NMR (400 MHz, DMSO) δ 8.86 (d, *J* = 6.9 Hz, 1H), 8.31 (s, 1H), 8.19-8.12 (m, 2H), 7.54 (d, *J* = 7.3 Hz, 1H), 7.47 (d, *J* = 8.7 Hz, 1H), 7.34 (d, *J* = 13.3 Hz, 1H), 7.27-7.21 (m, 2H), 7.20-7.12 (m, 5H), 6.97 (s, 1H), 6.75 (d, *J* = 9.9 Hz, 1H), 5.82-5.72 (m, 1H), 5.39 (d, *J* = 3.1 Hz, 1H), 4.65 (q, *J* = 6.9 Hz, 1H), 4.57 (dd, *J* = 10.2, 4.1 Hz, 1H), 4.40 (s, 1H), 4.31-4.18 (m, 4H), 4.10 (t, *J* = 10.2 Hz, 1H), 4.03 (t, *J* = 8.8 Hz, 1H), 3.97-3.87 (m, 2H), 3.80-3.69 (m, 2H), 3.61-3.48 (m, 5H), 3.37-3.28 (m, 2H), 3.26-3.17 (m, 1H), 3.13-3.06 (m, 1H), 2.61 (t, *J* = 12.9 Hz, 1H), 2.54-2.51 (m, 1H), 2.35-2.30 (m, 1H), 2.29-2.20 (m, 2H), 2.13-2.03 (m, 2H), 1.97-1.81 (m, 7H), 1.73-1.36 (m, 15H), 1.34-1.25 (m, 1H), 1.22 (s, 1H), 1.20-1.12 (m, 1H), 1.11-0.99 (m, 2H), 0.95 (d, *J* = 6.5 Hz, 3H), 0.90-0.73 (m, 24H), 0.70 (t, *J* = 7.4 Hz, 3H), 0.42 (t, *J* = 7.3 Hz, 3H), 0.37 (d, *J* = 6.8 Hz, 3H); HRMS (ESI): *m/z* calculated for $[M+H]^{+}$: 1371.8399, found: 1371.8404 (0.4 ppm); calculated for $[M+2H]^{2+}$: 686.4236, found: 686.4233 (-0.4 ppm); MS$^2$ HRMS (ESI, HCD): *m/z* calculated for ring fragment $[M_{ring}+H]^{+}$: 940.5866, found: 940.5856 (-1.1 ppm).

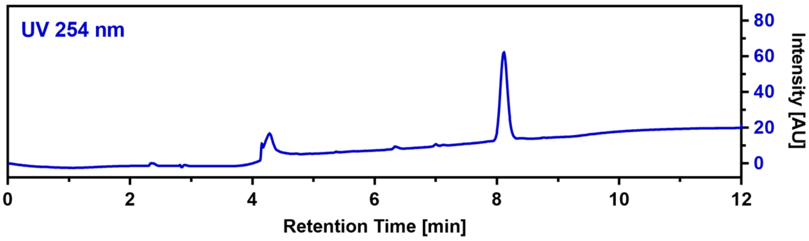

**CalA_R3F (68)**

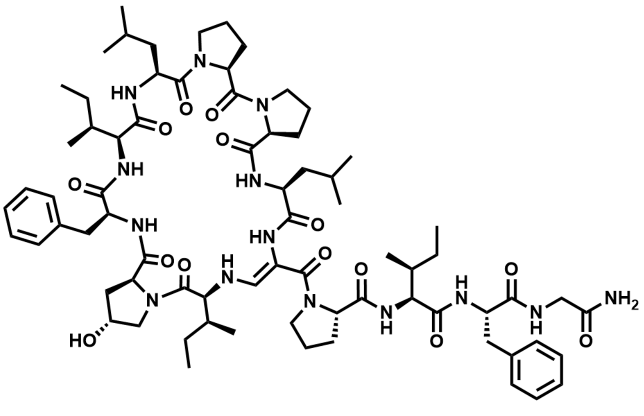

$C_{73}H_{108}N_{14}O_{14}$ M = 1405.75 g/mol

The linear precursor peptide of **68** was prepared via SPPS using Rink amide PS resin at a scale of 0.1 mmol initial resin capacity and purified by HPLC (solvent A: $H_2O$, acidified with 0.1% TFA; solvent B: ACN, acidified with 0.1% TFA). The obtained peptide was then dissolved in 10 mL of ACN and 4 equivalents DMP were added. The resulting suspension was shaken for one hour at RT and immediately purified by HPLC (solvent A: $H_2O$; solvent B: ACN) to give 7.2 mg (5.1 µmol) of **68** in a total yield of 5.1% (calculated in relation to initial resin capacity) as a white powder.

HRMS (ESI): *m/z* calculated for $[M+H]^+$: 1405.8242, found: 1405.8247 (0.4 ppm); calculated for $[M+2H]^{2+}$: 703.4158, found: 703.4160 (0.3 ppm); $MS^2$ HRMS (ESI, HCD): *m/z* calculated for ring fragment $[M_{ring}+H]^+$: 974.5710, found: 974.5698 (-1.2 ppm).

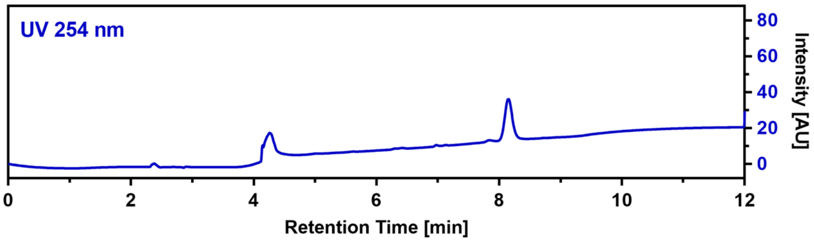

**CalA_R5D (69)**

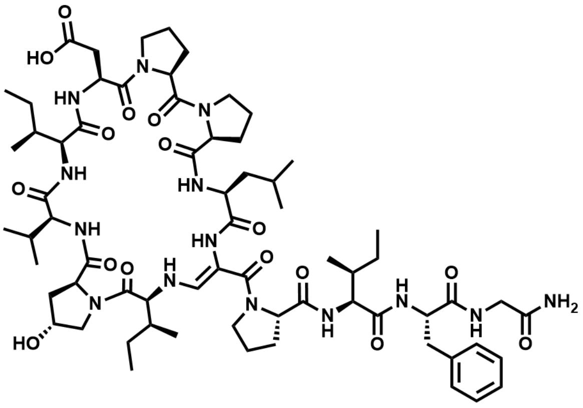

$C_{67}H_{102}N_{14}O_{16}$ M = 1359.64 g/mol

The linear precursor peptide of **69** was prepared via SPPS using Rink amide PS resin at a scale of 0.1 mmol initial resin capacity and purified by HPLC (solvent A: $H_2O$, acidified with 0.1% TFA; solvent B: ACN, acidified with 0.1% TFA). The obtained peptide was then dissolved in 10 mL of ACN and 4 equivalents DMP were added. The resulting suspension was shaken for one hour at RT and immediately purified by HPLC (solvent A: $H_2O$; solvent B: ACN) to give 4.7 mg (3.5 µmol) of **69** in a total yield of 3.5% (calculated in relation to initial resin capacity) as a white powder.

HRMS (ESI): *m/z* calculated for $[M+H]^+$: 1359.7671, found: 1359.7677 (0.4 ppm); calculated for $[M+2H]^{2+}$: 680.3872, found: 680.3870 (-0.3 ppm); $MS^2$ HRMS (ESI, HCD): *m/z* calculated for ring fragment $[M_{ring}+H]^+$: 928.5138, found: 928.5136 (-0.2 ppm).

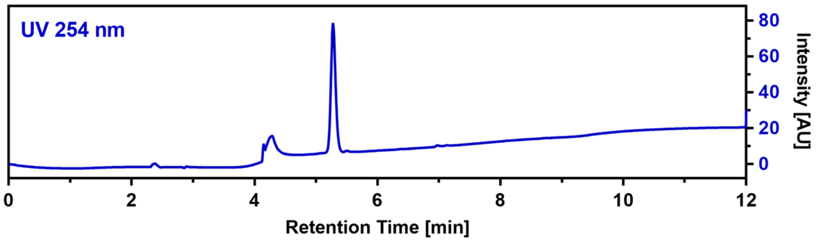

#### CalA_R8W (70)

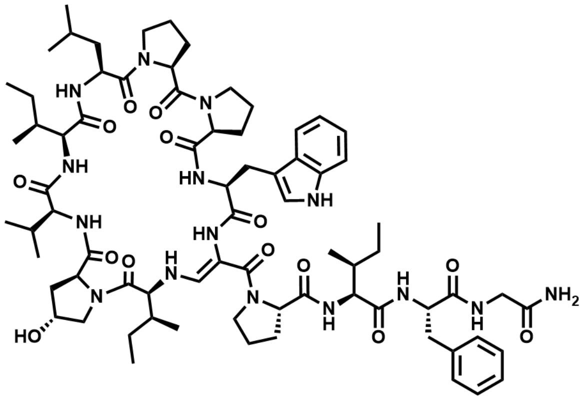

$C_{74}H_{107}N_{15}O_{14}$ M = 1430.76 g/mol

The linear precursor peptide of **70** was prepared via SPPS using Rink amide PS resin at a scale of 0.1 mmol initial resin capacity and purified by HPLC (solvent A: $H_2O$, acidified with 0.1% TFA; solvent B: ACN, acidified with 0.1% TFA). The obtained peptide was then dissolved in 10 mL of ACN and 4 equivalents DMP were added. The resulting suspension was shaken for one hour at RT and immediately purified by HPLC (solvent A: $H_2O$; solvent B: ACN) to give 6.2 mg (4.3 µmol) of **70** in a total yield of 4.3% (calculated in relation to initial resin capacity) as a white powder.

HRMS (ESI): *m/z* calculated for $[M+H]^+$: 1430.8195, found: 1430.8207 (0.8 ppm); calculated for $[M+2H]^{2+}$: 715.9134, found: 715.9127 (-1.0 ppm); $MS^2$ HRMS (ESI, HCD): *m/z* calculated for ring fragment $[M_{ring}+H]^+$: 999.5662, found: 999.5626 (-3.6 ppm).

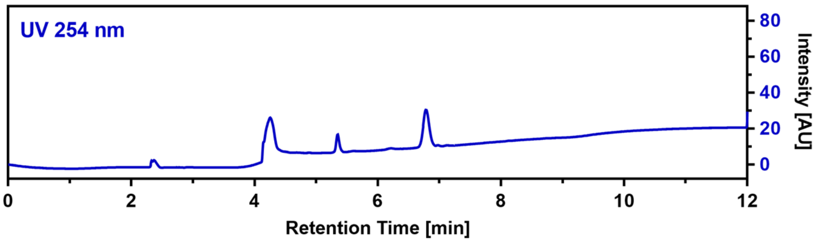

#### CalA_R8V (71)

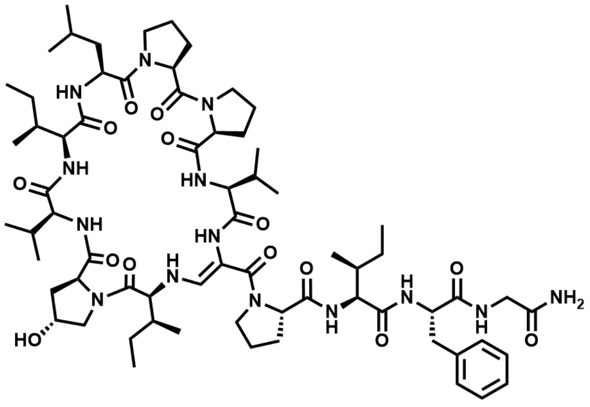

$C_{68}H_{106}N_{14}O_{14}$ M = 1343.68 g/mol

The linear precursor peptide of **71** was prepared via SPPS using Rink amide PS resin at a scale of 0.1 mmol initial resin capacity and purified by HPLC (solvent A: $H_2O$, acidified with 0.1% TFA; solvent B: ACN, acidified with 0.1% TFA). The obtained peptide was then dissolved in 10 mL of ACN and 4 equivalents DMP were added. The resulting suspension was shaken for one hour at RT and immediately purified by HPLC (solvent A: $H_2O$; solvent B: ACN) to give 9.2 mg (6.8 µmol) of **71** in a total yield of 6.8% (calculated in relation to initial resin capacity) as a white powder.

$^1H$ NMR (400 MHz, DMSO) δ 8.96 (s, 1H), 8.80 (s, 1H), 8.36 (s, 1H), 8.10 (t, $J$ = 6.1 Hz, 1H), 7.53 (d, $J$ = 8.3 Hz, 1H), 7.37 (d, $J$ = 6.5 Hz, 1H), 7.31 (d, $J$ = 13.3 Hz, 1H), 7.28-7.13 (m, 7H), 6.95 (s, 1H), 6.69 (d, $J$ = 9.9 Hz, 1H), 5.73 (dd, $J$ = 14.1, 9.9 Hz, 1H), 5.37 (d, $J$ = 3.1 Hz, 1H), 4.55 (td, $J$ = 10.3, 5.0 Hz, 2H), 4.40 (s, 1H), 4.30-4.20 (m, 4H), 4.07-3.97 (m, 2H), 3.90 (dd, $J$ = 16.5, 7.2 Hz, 1H), 3.79-3.71 (m, 4H), 3.66-3.52 (m, 3H), 3.48 (dd, $J$ = 16.7, 5.1 Hz, 1H), 3.26-3.16 (m, 2H), 3.12 (dd, $J$ = 13.7, 3.1 Hz, 1H), 3.03-2.95 (m, 1H), 2.69-2.57 (m, 2H), 2.47-2.38 (m, 1H), 2.32-2.21 (m, 3H), 2.12-2.04 (m, 2H), 2.01-1.96 (m, 1H), 1.96-1.86 (m, 5H), 1.86-1.76 (m, 2H), 1.76-1.60 (m, 3H), 1.55 (td, $J$ = 13.5, 6.6 Hz, 2H), 1.49-1.32 (m, 4H), 1.27-1.17 (m, 2H), 1.17-1.02 (m, 2H), 0.99 (d, $J$ = 7.2 Hz, 3H), 0.97 (d, $J$ = 6.9 Hz, 3H), 0.89 (d, $J$ = 6.6 Hz, 3H), 0.87 (d, $J$ = 6.5 Hz, 3H), 0.84-0.74 (m, 15H), 0.74-0.69 (m, 4H), 0.41 (t, $J$ = 7.3 Hz, 3H),

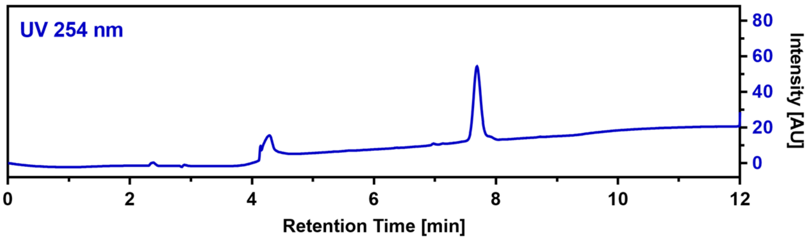

0.38 (d, $J$ = 6.8 Hz, 3H); HRMS (ESI): $m/z$ calculated for $[M+H]^+$: 1343.8086, found: 1343.8092 (0.4 ppm); calculated for $[M+2H]^{2+}$: 672.9096, found: 672.9101 (0.7 ppm); $MS^2$ HRMS (ESI, HCD): $m/z$ calculated for ring fragment $[M_{ring}+H]^+$: 912.5553, found: 912.5564 (1.2 ppm).

**CalA_C3L (72)**

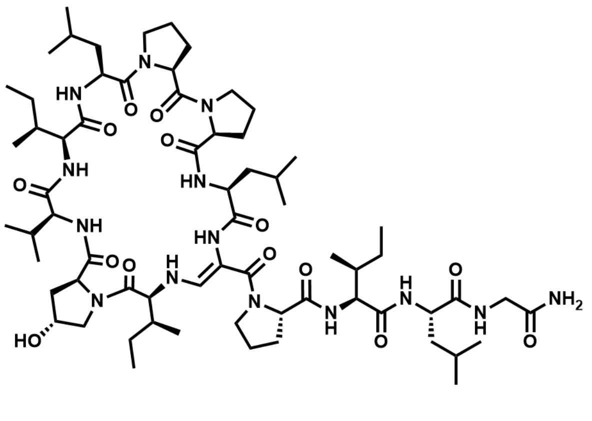

$C_{66}H_{110}N_{14}O_{14}$ M = 1323.69 g/mol

The linear precursor peptide of **72** was prepared via SPPS using Rink amide PS resin at a scale of 0.1 mmol initial resin capacity and purified by HPLC (solvent A: $H_2O$, acidified with 0.1% TFA; solvent B: ACN, acidified with 0.1% TFA). The obtained peptide was then dissolved in 10 mL of ACN and 4 equivalents DMP were added. The resulting suspension was shaken for one hour at RT and immediately purified by HPLC (solvent A: $H_2O$; solvent B: ACN) to give 6.3 mg (4.8 µmol) of **72** in a total yield of 4.8% (calculated in relation to initial resin capacity) as a white powder.

$^1$H NMR (400 MHz, DMSO) δ 9.04 (d, $J$ = 6.8 Hz, 1H), 8.83 (d, $J$ = 5.8 Hz, 1H), 8.29 (d, $J$ = 6.4 Hz, 1H), 8.28 (s, 1H), 7.46 (d, $J$ = 6.9 Hz, 1H), 7.43 (d, $J$ = 8.1 Hz, 1H), 7.35 (d, $J$ = 6.5 Hz, 1H), 7.32 (d, $J$ = 3.1 Hz, 1H), 7.19 (s, 1H), 6.91 (s, 1H), 6.63 (d, $J$ = 10.1 Hz, 1H), 5.81 (dd, $J$ = 12.9, 9.8 Hz, 1H), 5.36 (d, $J$ = 3.0 Hz, 1H), 4.66-4.54 (m, 2H), 4.40 (s, 1H), 4.29-4.16 (m, 4H), 4.15-4.07 (m, 2H), 4.07-4.01 (m, 1H), 3.92 (dd, $J$ = 16.2, 7.1 Hz, 1H), 3.79 (dd, $J$ = 8.5, 6.6 Hz, 1H), 3.76-3.66 (m, 2H), 3.65-3.55 (m, 2H), 3.54-3.45 (m, 2H), 3.40 (dd, $J$ = 16.2, 5.1 Hz, 1H), 3.22-3.12 (m, 1H), 3.01-2.94 (m, 1H), 2.71-2.62 (m, 1H), 2.35-2.29 (m, 1H), 2.28-2.19 (m, 2H), 2.12-2.04 (m, 2H), 2.03-1.96 (m, 1H), 1.95-1.80 (m, 7H), 1.76-1.66 (m, 3H), 1.66-1.55 (m, 3H), 1.54-1.37 (m, 8H), 1.30-1.22 (m, 2H), 1.21-1.09 (m, 1H), 1.02-0.93 (m, 1H), 0.89 (d, $J$ = 5.1 Hz, 3H), 0.88-0.79 (m, 30H), 0.78 (d, $J$ = 3.8 Hz, 3H), 0.76 (d, $J$ = 3.8 Hz, 3H), 0.75-0.70 (m, 3H); HRMS (ESI): $m/z$ calculated for $[M+H]^+$: 1323.8399, found: 1323.8419 (1.5 ppm); calculated for $[M+2H]^{2+}$: 662.4236, found: 662.4237 (0.2 ppm); $MS^2$ HRMS (ESI, HCD): $m/z$ calculated for ring fragment $[M_{ring}+H]^+$: 926.5710, found: 926.5701 (-1.0 ppm).

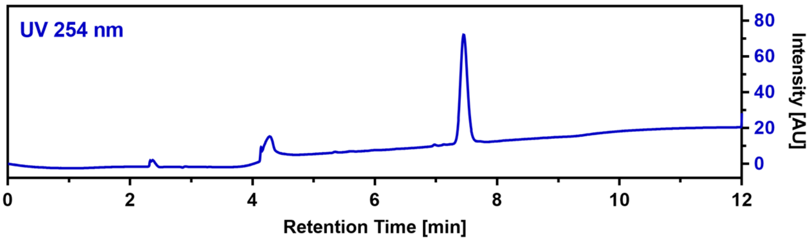

**CalA_C3I (73)**

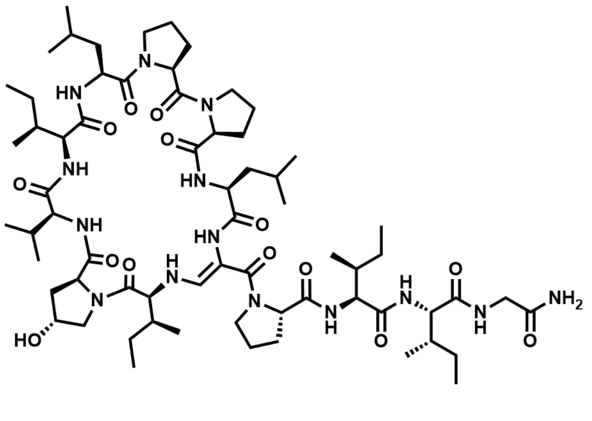

$C_{66}H_{110}N_{14}O_{14}$ M = 1323.69 g/mol

The linear precursor peptide of **73** was prepared via SPPS using Rink amide PS resin at a scale of 0.1 mmol initial resin capacity and purified by HPLC (solvent A: $H_2O$, acidified with 0.1% TFA; solvent B: ACN, acidified with 0.1% TFA). The obtained peptide was then dissolved in 10 mL of ACN and 4 equivalents DMP were added. The resulting suspension was shaken for one hour at RT and immediately purified by HPLC (solvent A: $H_2O$; solvent B: ACN) to give 2.3 mg (1.7 µmol) of **73** in a total yield of 1.7% (calculated in relation to initial resin capacity) as a white powder.

HRMS (ESI): $m/z$ calculated for $[M+H]^+$: 1323.8399, found: 1323.8405 (0.5 ppm); calculated for $[M+2H]^{2+}$: 662.4236, found: 662.4233

(-0.5 ppm); MS[2] HRMS (ESI, HCD): *m/z* calculated for ring fragment $[M_{ring}+H]^+$: 926.5710, found: 926.5721 (1.2 ppm).

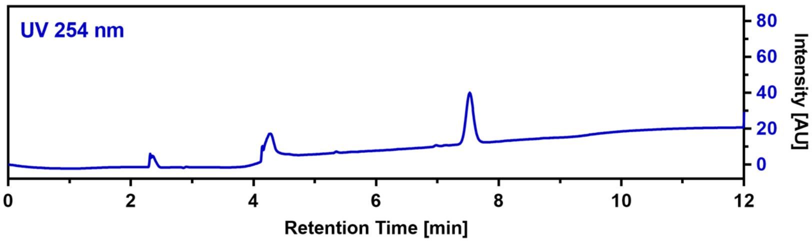

**CalA_C3X (74)**

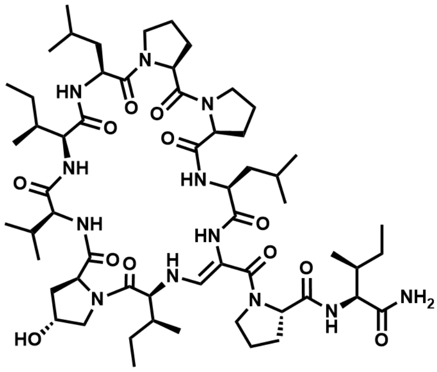

$C_{58}H_{96}N_{12}O_{12}$ M = 1153.48 g/mol

The linear precursor peptide of **74** was prepared via SPPS using Rink amide PS resin at a scale of 0.1 mmol initial resin capacity and purified by HPLC (solvent A: $H_2O$, acidified with 0.1% TFA; solvent B: ACN, acidified with 0.1% TFA). The obtained peptide was then dissolved in 10 mL of ACN and 4 equivalents DMP were added. The resulting suspension was shaken for one hour at RT and immediately purified by HPLC (solvent A: $H_2O$; solvent B: ACN) to give 8.8 mg (7.6 µmol) of **74** in a total yield of 7.6% (calculated in relation to initial resin capacity) as a white powder.

$^1H$ NMR (400 MHz, DMSO) δ 8.95 (s, 1H), 8.77 (s, 1H), 8.06 (s, 1H), 7.47 (d, *J* = 9.1 Hz, 1H), 7.42 (d, *J* = 6.8 Hz, 1H), 7.15 (d, *J* = 12.9 Hz, 1H), 7.05 (s, 1H), 6.81 (s, 1H), 6.74 (d, *J* = 9.9 Hz, 1H), 5.56 (dd, *J* = 12.9, 10.2 Hz, 1H), 5.36 (d, *J* = 3.1 Hz, 1H), 4.61 (q, *J* = 6.7 Hz, 1H), 4.51-4.43 (m, 1H), 4.39 (s, 1H), 4.33 (dd, *J* = 9.4, 7.5 Hz, 1H), 4.27-4.21 (m, 1H), 4.04 (t, *J* = 9.0 Hz, 1H), 3.99 (dd, *J* = 9.1, 6.3 Hz, 1H), 3.94 (t, *J* = 9.7 Hz, 1H), 3.83 (t, *J* = 7.5 Hz, 1H), 3.78 (d, *J* = 11.4 Hz, 1H), 3.67 (dd, *J* = 10.9, 3.5 Hz, 1H), 3.64-3.54 (m, 2H), 3.54-3.43 (m, 2H), 3.29-3.23 (m, 1H), 3.21-3.13 (m, 1H), 3.07-2.97 (m, 1H), 2.69-2.58 (m, 1H), 2.32-2.15 (m, 3H), 2.12-2.04 (m, 1H), 2.02-1.98 (m, 1H), 1.97-1.84 (m, 7H), 1.80-1.69 (m, 4H), 1.66-1.58 (m, 3H), 1.56-1.37 (m, 7H), 1.36-1.27 (m, 1H), 1.22-1.10 (m, 1H), 1.08-0.94 (m, 2H), 0.92-0.89 (m, 9H), 0.86-0.79 (m, 27H); HRMS (ESI): *m/z* calculated for $[M+H]^+$: 1153.7343, found: 1153.7338 (-0.4 ppm); calculated for $[M+2H]^{2+}$: 577.3708, found: 577.3702 (-1.0 ppm); MS[2] HRMS (ESI, HCD): *m/z* calculated for ring fragment $[M_{ring}+H]^+$: 926.5710, found: 926.5721 (1.2 ppm).

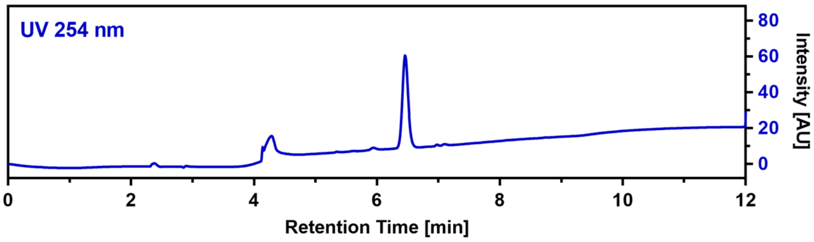

**CalA_C4Aha (75)**

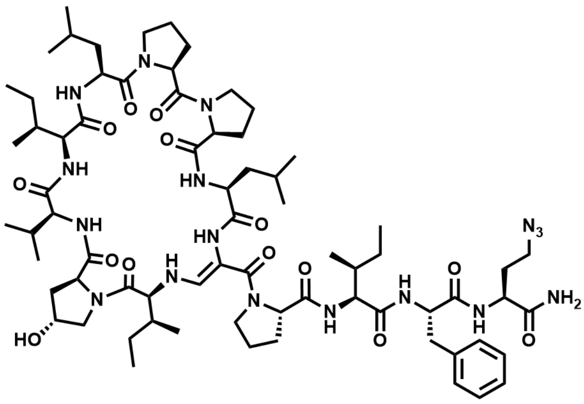

$C_{71}H_{111}N_{17}O_{14}$ M = 1426.77 g/mol

The linear precursor peptide of **75** was prepared via SPPS using Rink amide PS resin at a scale of 0.1 mmol initial resin capacity and purified by HPLC (solvent A: $H_2O$, acidified with 0.1% TFA; solvent B: ACN, acidified with 0.1% TFA). The obtained peptide was then dissolved in 10 mL of ACN and 4 equivalents DMP were added. The resulting suspension was shaken for one hour at RT and immediately purified by HPLC (solvent A: $H_2O$; solvent B: ACN) to give 8.2 mg (5.7 µmol) of **75** in a total yield of 5.7% (calculated in relation to initial resin capacity) as a white powder.

$^1H$ NMR (400 MHz, DMSO) δ 9.00 (s, 1H), 8.77 (s, 1H), 8.15 (s, 1H), 7.68 (d, *J* = 7.2 Hz, 1H), 7.45 (d, *J* = 8.1 Hz, 1H), 7.40 (d, *J* = 8.4 Hz, 1H), 7.35 (d,

*J* = 7.2 Hz, 1H), 7.26-7.18 (m, 3H), 7.14-7.02 (m, 4H), 6.77 (s, 1H), 6.72 (d, *J* = 9.7 Hz, 1H), 5.77 (dd, *J* = 12.9, 10.0 Hz, 1H), 5.37 (d, *J* = 3.0 Hz, 1H), 4.64 (q, *J* = 6.9 Hz, 1H), 4.56 (dd, *J* = 9.9, 5.0 Hz, 1H), 4.39 (s, 1H), 4.31-4.24 (m, 2H), 4.22-4.12 (m, 3H), 4.10-4.01 (m, 1H), 3.85-3.70 (m, 4H), 3.64 (dd, *J* = 11.1, 3.5 Hz, 1H), 3.61-3.53 (m, 2H), 3.52-3.44 (m, 4H), 3.24 (td, *J* = 11.0, 6.0 Hz, 1H), 3.15 (dd, *J* = 13.3, 3.0 Hz, 1H), 2.99 (dd, *J* = 10.9, 6.8 Hz, 1H), 2.84-2.73 (m, 1H), 2.71-2.55 (m, 1H), 2.34-2.19 (m, 3H), 2.12-2.05 (m, 1H), 2.04-1.97 (m, 2H), 1.96-1.81 (m, 9H), 1.79-1.68 (m, 4H), 1.66-1.54 (m, 3H), 1.51-1.41 (m, 1H), 1.39-1.32 (m, 1H), 1.30-1.21 (m, 2H), 1.18-1.05 (m, 5H), 0.96 (d, *J* = 6.8 Hz, 3H), 0.91 (d, *J* = 6.5 Hz, 3H), 0.88 (d, *J* = 6.6 Hz, 6H), 0.85-0.74 (m, 15H), 0.70 (t, *J* = 7.4 Hz, 3H), 0.59-0.52 (m, 1H), 0.50 (d, *J* = 6.8 Hz, 3H), 0.26 (t, *J* = 7.4 Hz, 3H); HRMS (ESI): *m/z* calculated for $[M+H]^+$: 1426.8569, found: 1426.8575 (0.4 ppm); calculated for $[M+2H]^{2+}$: 713.9321, found: 713.9322 (0.1 ppm); $MS^2$ HRMS (ESI, HCD): *m/z* calculated for ring fragment $[M_{ring}+H]^+$: 926.5710, found: 926.5727 (1.8 ppm).

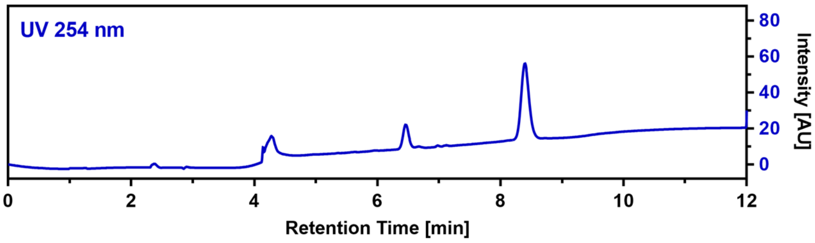

**CalA_C4X (76)**

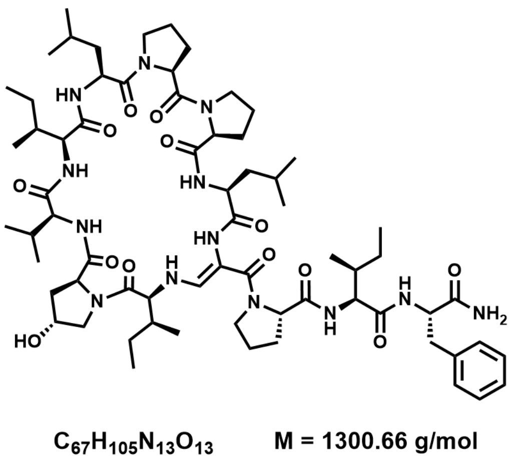

The linear precursor peptide of **76** was prepared via SPPS using Rink amide PS resin at a scale of 0.1 mmol initial resin capacity and purified by HPLC (solvent A: $H_2O$, acidified with 0.1% TFA; solvent B: ACN, acidified with 0.1% TFA). The obtained peptide was then dissolved in 10 mL of ACN and 4 equivalents DMP were added. The resulting suspension was shaken for one hour at RT and immediately purified by HPLC (solvent A: $H_2O$; solvent B: ACN) to give 1.9 mg (1.5 µmol) of **76** in a total yield of 1.5% (calculated in relation to initial resin capacity) as a white powder.

$^1$H NMR (400 MHz, DMSO) δ 9.02 (s, 1H), 8.82 (s, 1H), 8.27 (s, 1H), 7.45 (d, *J* = 9.0 Hz, 2H), 7.39 (d, *J* = 7.4 Hz, 1H), 7.22-7.18 (m, 3H), 7.15 (d, *J* = 7.8 Hz, 1H), 7.14-7.08 (m, 4H), 6.73 (d, *J* = 9.6 Hz, 1H), 5.86-5.77 (m, 1H), 5.38 (d, *J* = 3.0 Hz, 1H), 4.66-4.55 (m, 2H), 4.41 (s, 1H), 4.30-4.17 (m, 4H), 4.06 (t, *J* = 8.2 Hz, 1H), 3.88 (t, *J* = 10.5 Hz, 1H), 3.84-3.77 (m, 3H), 3.68-3.63 (m, 1H), 3.62-3.55 (m, 2H), 3.53-3.44 (m, 2H), 3.24-3.16 (m, 2H), 3.02-2.94 (m, 1H), 2.70-2.61 (m, 1H), 2.56 (t, *J* = 12.7 Hz, 1H), 2.33-2.20 (m, 3H), 2.13-2.05 (m, 1H), 2.02-1.97 (m, 1H), 1.96-1.84 (m, 7H), 1.82-1.68 (m, 3H), 1.66-1.54 (m, 4H), 1.54-1.46 (m, 2H), 1.44-1.32 (m, 3H), 1.30-1.21 (m, 3H), 1.14-1.01 (m, 2H), 0.94 (d, *J* = 6.6 Hz, 3H), 0.90-0.77 (m, 24H), 0.74-0.70 (m, 1H), 0.67 (t, *J* = 7.4 Hz, 3H), 0.48 (t, *J* = 7.2 Hz, 3H), 0.33 (d, *J* = 6.8 Hz, 3H); HRMS (ESI): *m/z* calculated for $[M+H]^+$: 1300.8028, found: 1300.8048 (1.5 ppm); calculated for $[M+2H]^{2+}$: 650.9050, found: 950.9052 (0.3 ppm); $MS^2$ HRMS (ESI, HCD): *m/z* calculated for ring fragment $[M_{ring}+H]^+$: 926.5710, found: 926.5722 (1.3 ppm).

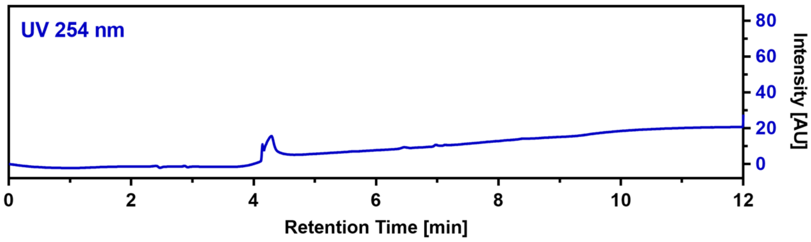

**CalB_R2Oic_R3Chg (77)**

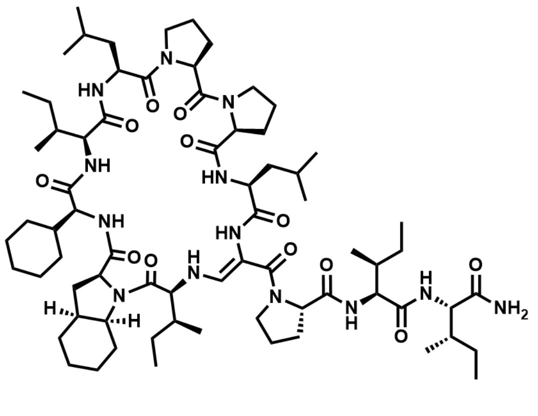

$C_{71}H_{117}N_{13}O_{12}$ M = 1344.80 g/mol

The linear precursor peptide of **77** was prepared via microwave-assisted SPPS using Rink amide PS resin at a scale of 0.1 mmol initial resin capacity. After purification by HPLC (solvent A: $H_2O$, acidified with 0.1% TFA; solvent B: ACN, acidified with 0.1% TFA), 72 mg (52 µmol, 52%) of the peptide precursor was obtained as a white powder. For the oxidation of the serine residue and concomitant cyclization, the peptide was dissolved in 15 mL of ACN and 3 equivalents of DMP were added. The resulting suspension was shaken for 3.5 hours at RT and immediately purified by HPLC (solvent A: $H_2O$; solvent B: ACN) to give 4.1 mg (3.0 µmol, 5.8%) of **77** in a total yield of 3.0% (calculated in relation to initial resin capacity) as a white powder.

$^1$H NMR (700 MHz, DMSO) δ 8.85 (d, *J* = 7.0 Hz, 1H), 8.58 (d, *J* = 6.6 Hz, 1H), 8.26 (s, 1H), 7.62 – 7.55 (m, 2H), 7.30 (d, *J* = 9.6 Hz, 1H), 7.12 (d, *J* = 13.4 Hz, 1H), 7.05 (s, 1H), 7.03 (s, 1H), 6.66 (d, *J* = 10.0 Hz, 1H), 5.58 (dd, *J* = 13.4, 10.0 Hz, 1H), 4.59 (q, *J* = 6.7 Hz, 1H), 4.53 (td, *J* = 10.3, 3.8 Hz, 1H), 4.27 (dd, *J* = 10.0, 7.7 Hz, 1H), 4.23 (t, *J* = 7.9 Hz, 1H), 4.17 (dd, *J* = 9.6, 5.1 Hz, 1H), 4.13-4.06 (m, 2H), 4.04 (dt, *J* = 12.4, 6.5 Hz, 1H), 3.95 (t, *J* = 8.8 Hz, 1H), 3.81 (t, *J* = 10.2 Hz, 1H), 3.73 (dd, *J* = 9.5, 6.6 Hz, 1H), 3.60-3.51 (m, 2H), 3.51-3.45 (m, 1H), 3.45-3.41 (m, 1H), 3.28 (dt, *J* = 10.2, 7.0 Hz, 1H), 3.18 (td, *J* = 10.9, 5.8 Hz, 1H), 3.07 (dd, *J* = 11.1, 7.0 Hz, 1H), 2.49-2.44 (m, 2H), 2.34-2.30 (m, 1H), 2.29-2.25 (m, 1H), 2.25-2.19 (m, 1H), 2.03-1.96 (m, 2H), 1.93-1.83 (m, 7H), 1.82-1.78 (m, 1H), 1.77-1.72 (m, 2H), 1.71-1.64 (m, 7H), 1.63-1.55 (m, 5H), 1.54-1.40 (m, 9H), 1.31-1.18 (m, 7H), 1.17-1.10 (m, 3H), 1.06-0.99 (m, 2H), 0.94-0.90 (m, 1H), 0.90-0.86 (m, 12H), 0.86-0.85 (m, 1H), 0.84-0.77 (m, 18H), 0.75 (t, *J* = 6.4 Hz, 3H), 0.73 (t, *J* = 6.0 Hz, 3H); $^{13}$C NMR (176 MHz, DMSO) δ 173.0, 172.8, 172.8, 172.4, 171.9, 171.5, 171.4, 171.4, 171.3, 171.2, 171.2, 167.2, 142.3, 98.1, 64.8, 63.9, 63.8, 62.5, 61.2, 61.1, 59.8, 58.1, 57.4, 57.3, 50.0, 49.2, 48.7, 47.1, 46.1, 41.1, 40.4, 37.5, 36.8, 36.3, 36.2, 35.8, 29.7, 29.5, 29.0, 28.5, 26.3, 25.8, 25.6, 25.2, 25.1, 25.1, 25.0, 24.9, 24.8, 24.6, 24.0, 23.6, 23.5, 23.1, 22.7, 22.2, 20.9, 19.4, 15.8, 15.7, 15.1, 11.5, 11.0, 10.8, 9.5; HRMS (ESI): *m/z* calculated for $[M+H]^+$: 1344.9017, found: 1344.9020 (0.2 ppm); calculated for $[M+2H]^{2+}$: 672.9545, found: 972.9536 (-1.3 ppm); $MS^2$ HRMS (ESI, HCD): *m/z* calculated for ring fragment $[M_{ring}+H]^+$: 1004.6543, found: 1004.6513 (-3.0 ppm).

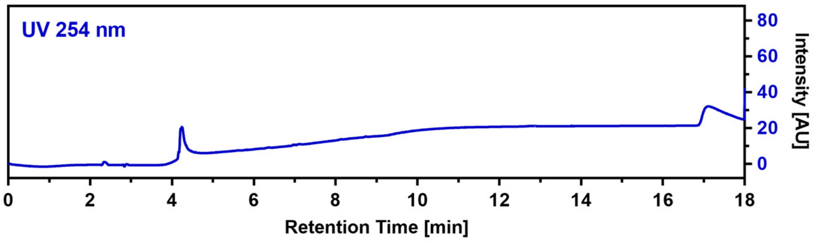

**CalB_R3Pra (78)**

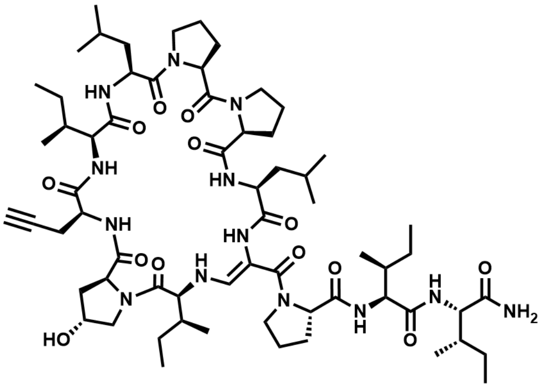

$C_{64}H_{103}N_{13}O_{13}$ M = 1262.61 g/mol

The linear precursor peptide of **78** was prepared via microwave-assisted SPPS using Rink amide PS resin at a scale of 0.1 mmol initial resin capacity. After purification by HPLC (solvent A: $H_2O$, acidified with 0.1% TFA; solvent B: ACN, acidified with 0.1% TFA), 42 mg (33 µmol, 33%) of the peptide precursor was obtained as a white powder. For the oxidation of the serine residue and concomitant cyclization, the peptide was dissolved in 8 mL of ACN and 3 equivalents of DMP were added. The resulting suspension was shaken for 3 hours at RT and immediately purified by HPLC (solvent A: $H_2O$; solvent B: ACN) to give 9.2 mg (7.3 µmol, 22.3%) of **78** in a total yield of 7.3% (calculated in relation to initial resin capacity) as a white powder.

$^1$H NMR (400 MHz, DMSO) δ 9.13 (d, *J* = 6.8 Hz, 1H), 8.30 (s, 1H), 7.80 (d, *J* = 9.0 Hz, 1H), 7.60 (d, *J* = 10.0 Hz, 1H), 7.49 (d, *J* = 7.6 Hz, 1H), 7.23 (d, *J* = 9.7 Hz, 1H), 7.14 (s, 1H), 7.07 (d, *J* = 13.2 Hz, 1H), 6.97 (s, 1H), 6.68 (d, *J* = 9.9 Hz, 1H), 5.62-5.50 (m, 1H), 5.40 (d, *J* = 3.1 Hz, 1H), 4.75-4.64 (m, 1H), 4.56-4.46 (m, 1H), 4.42 (s, 1H), 4.33-4.21 (m, 3H), 4.19-4.13 (m, 1H), 4.12-4.06 (m, 1H), 4.05-3.95 (m, 3H), 3.77-3.69 (m, 2H), 3.62-3.44 (m, 4H), 3.39-3.33 (m, 1H), 3.23-3.13 (m, 1H), 3.09-2.99 (m, 1H), 2.97-2.93 (m, 1H), 2.78-2.69 (m, 1H), 2.41-2.32 (m, 1H), 2.31-2.17 (m, 2H), 2.12-2.01 (m, 2H), 2.01-1.75 (m, 8H), 1.74-1.55 (m, 6H), 1.54-1.33 (m, 8H), 1.28-1.21 (m, 1H), 1.19-0.96 (m, 5H), 0.95-0.69 (m, 36H); HRMS (ESI): *m/z* calculated for $[M+H]^+$: 1262.7871, found: 1262.7857 (-1.1 ppm); calculated for $[M+2H]^{2+}$: 631.8972, found: 631.8963 (-1.4 ppm); $MS^2$ HRMS (ESI, HCD): *m/z* calculated for ring fragment $[M_{ring}+H]^+$: 922.5397, found: 922.5377 (-2.2 ppm).

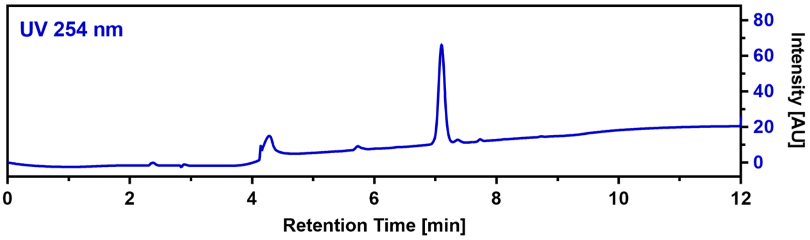

**CalB_R4Pra (79)**

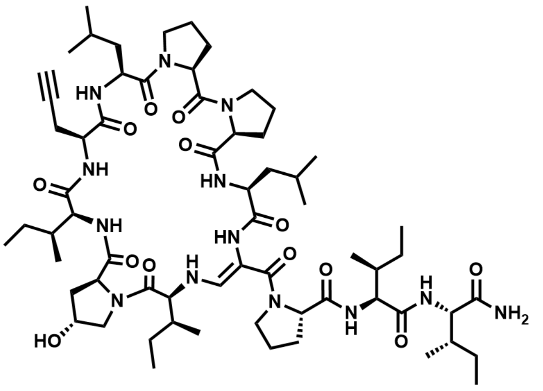

$C_{64}H_{103}N_{13}O_{13}$ M = 1262.61 g/mol

The linear precursor peptide of **79** was prepared via microwave-assisted SPPS using Rink amide PS resin at a scale of 0.1 mmol initial resin capacity. After purification by HPLC (solvent A: $H_2O$, acidified with 0.1% TFA; solvent B: ACN, acidified with 0.1% TFA), 29 mg (23 µmol, 23%) of the peptide precursor was obtained as a white powder. For the oxidation of the serine residue and concomitant cyclization, the peptide was dissolved in 5 mL of ACN and 3 equivalents of DMP were added. The resulting suspension was shaken for 2.5 hours at RT and immediately purified by HPLC (solvent A: $H_2O$; solvent B: ACN) to give 2.6 mg (2.1 µmol, 9.1%) of **79** in a total yield of 2.1% (calculated in relation to initial resin capacity) as a white powder.

$^1$H NMR (400 MHz, DMSO) δ 8.81 (s, 1H), 8.73 (s, 1H), 8.30 (s, 1H), 7.59 (d, *J* = 9.9 Hz, 1H), 7.55 (d, *J* = 7.0 Hz, 1H), 7.23 (d, *J* = 9.6 Hz, 1H), 7.14 (s, 1H), 7.07 (d, *J* = 13.3 Hz, 1H), 6.96 (d, *J* = 2.8 Hz, 1H), 6.67 (d, *J* = 9.7 Hz, 1H), 5.70-5.60 (m, 1H), 5.45-5.33 (m, 1H), 4.63-4.56 (m, 1H), 4.56-4.48 (m, 1H), 4.42 (s, 1H), 4.28-4.18 (m, 3H), 4.17-4.10 (m, 1H), 4.10-4.01 (m, 3H),

3.98-3.89 (m, 1H), 3.78-3.67 (m, 2H), 3.61-3.48 (m, 4H), 3.34-3.28 (m, 1H), 3.23-3.13 (m, 2H), 2.98-2.94 (m, 1H), 2.60-2.53 (m, 1H), 2.33-2.20 (m, 3H), 2.11-1.97 (m, 2H), 1.95-1.84 (m, 8H), 1.73-1.56 (m, 6H), 1.51-1.39 (m, 8H), 1.24 (s, 1H), 1.17-0.95 (m, 5H), 0.90-0.76 (m, 36H); HRMS (ESI): *m/z* calculated for $[M+H]^+$: 1262.7871, found: 1262.7861 (-0.8 ppm); $MS^2$ HRMS (ESI, HCD): *m/z* calculated for ring fragment $[M_{ring}+H]^+$: 922.5397, found: 922.5407 (1.1 ppm).

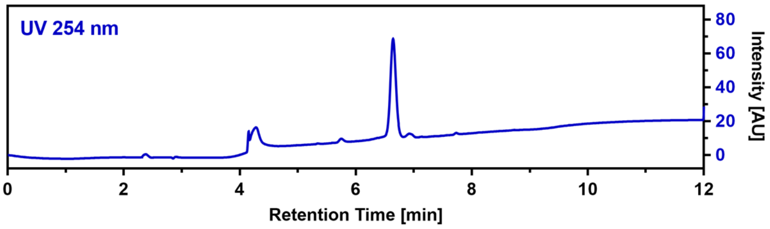

**CalB_R7Pra (80)**

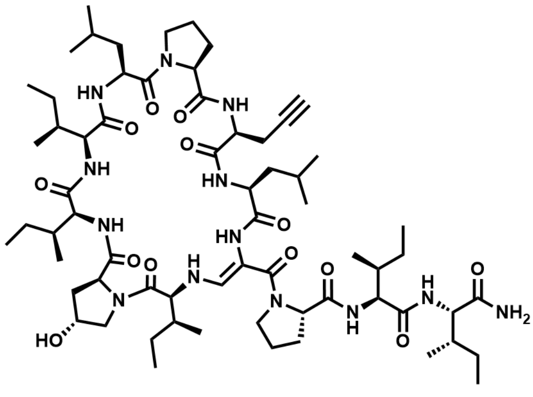

The linear precursor peptide of **80** was prepared via microwave-assisted SPPS using Rink amide PS resin at a scale of 0.1 mmol initial resin capacity. After purification by HPLC (solvent A: $H_2O$, acidified with 0.1% TFA; solvent B: ACN, acidified with 0.1% TFA), 40 mg (31 µmol, 22%) of the peptide precursor was obtained as a white powder. For the oxidation of the serine residue and concomitant cyclization, the peptide was dissolved in 10 mL of ACN and 4 equivalents of DMP were added. The resulting suspension was shaken for 5 hours at RT and immediately purified by HPLC (solvent A: $H_2O$; solvent B: ACN) to give 1.7 mg (1.3 µmol, 4.3%) of **80** in a total yield of 0.9% (calculated in relation to initial resin capacity) as a white powder. HRMS (ESI): *m/z* calculated for $[M+H]^+$: 1278.8184, found: 1278.8171 (-1.0 ppm); $MS^2$ HRMS (ESI, HCD): *m/z* calculated for ring fragment $[M_{ring}+H]^+$: 938.5710, found: 938.5712 (0.2 ppm).

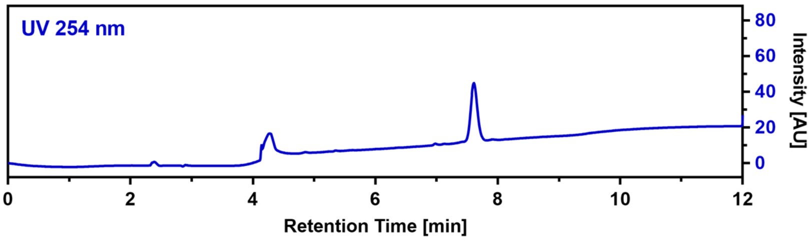

**CalB_C3Pra (81)**

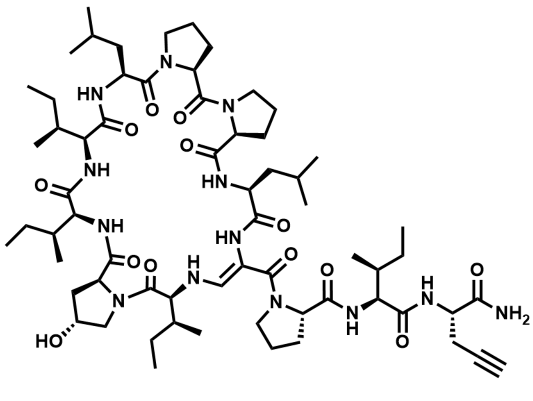

The linear precursor peptide of **81** was prepared via microwave-assisted SPPS using Rink amide PS resin at a scale of 0.1 mmol initial resin capacity. After purification by HPLC (solvent A: $H_2O$, acidified with 0.1% TFA; solvent B: ACN, acidified with 0.1% TFA), 43 mg (33 µmol, 33%) of the peptide precursor was obtained as a white powder. For the oxidation of the serine residue and concomitant cyclization, the peptide was dissolved in 5 mL of ACN and 4 equivalents of DMP were added. The resulting suspension was shaken for 2.5 hours at RT and immediately purified by HPLC (solvent A: $H_2O$; solvent B: ACN) to give 1.9 mg (1.5 µmol, 4.5%) of **81** in a total yield of 1.5% (calculated in relation to initial resin capacity) as a white powder.

$^1$H NMR (400 MHz, DMSO) δ 8.99 (d, *J* = 7.0 Hz, 1H), 8.85 (d, *J* = 6.5 Hz, 1H), 8.35 (s, 1H), 7.78 (d, *J* = 8.7 Hz, 1H), 7.44-7.35 (m, 3H), 7.16 (s, 1H), 7.08 (d, *J* = 13.4 Hz, 1H), 6.70 (d, *J* = 9.8 Hz, 1H), 5.83-5.70 (m, 1H), 5.40 (d, *J* = 3.1 Hz, 1H), 4.64-4.49 (m, 2H), 4.41 (s, 1H), 4.28-4.19 (m, 5H), 4.18-4.12 (m, 1H), 4.06-4.00 (m, 1H), 3.91-3.84 (m, 1H), 3.82-3.75 (m, 2H), 3.70-3.56 (m, 4H), 3.52-3.45 (m, 1H), 3.23-3.13 (m, 1H), 3.12-3.03 (m, 1H), 2.73-2.68 (m, 1H), 2.68-2.59 (m, 1H), 2.31-2.18 (m, 3H), 2.14-2.04 (m, 2H), 1.99-1.77 (m, 8H), 1.71-1.48 (m, 6H), 1.47-1.32 (m, 8H), 1.31-1.21 (m, 1H), 1.19-0.99 (m, 5H), 0.92-.72 (m, 36H); HRMS (ESI): *m/z* calculated for $[M+H]^+$: 1262.7871, found: 1262.7865 (-0.5 ppm); calculated for $[M+2H]^{2+}$: 631.8972, found: 631.8967 (-0.8 ppm); MS$^2$ HRMS (ESI, HCD): *m/z* calculated for ring fragment $[M_{ring}+H]^+$: 940.5866, found: 940.5854 (-1.3 ppm).

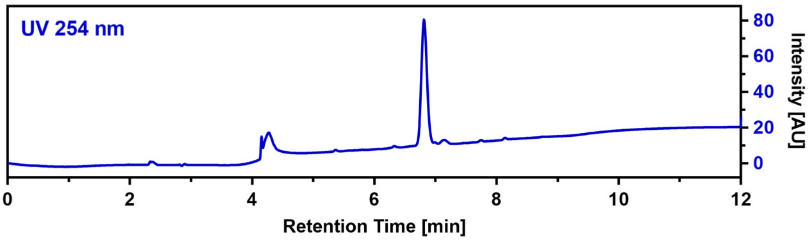

**CalB_C4Pra (82)**

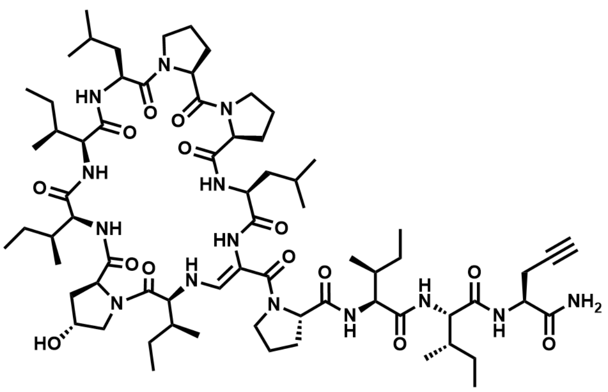

$C_{70}H_{114}N_{14}O_{14}$ M = 1375.77 g/mol

The linear precursor peptide of **82** was prepared via microwave-assisted SPPS using Rink amide PS resin at a scale of 0.2 mmol initial resin capacity. After purification by HPLC (solvent A: $H_2O$, acidified with 0.1% TFA; solvent B: ACN, acidified with 0.1% TFA), 130 mg (93 µmol, 46%) of the peptide precursor was obtained as a white powder. For the oxidation of the serine residue and concomitant cyclization, the peptide was dissolved in 20 mL of ACN and 4 equivalents of DMP were added. The resulting suspension was shaken for 4 hours at RT and immediately purified by HPLC (solvent A: $H_2O$; solvent B: ACN) to give 25.3 mg (18.4 µmol, 19.7%) of **82** in a total yield of 9.0% (calculated in relation to initial resin capacity) as a white powder.

$^1$H NMR (700 MHz, DMSO) δ 9.01 (s, 1H), 8.85 (s, 1H), 8.17 (s, 1H), 8.10 (d, *J* = 8.3 Hz, 1H), 7.47 (d, *J* = 8.7 Hz, 1H), 7.36 (d, *J* = 7.0 Hz, 1H), 7.26 (d, *J* = 7.7 Hz, 1H), 7.22 (s, 1H), 6.98 (d, *J* = 13.1 Hz, 1H), 6.70 (s, 1H), 6.63 (d, *J* = 9.9 Hz, 1H), 5.71 (dd, *J* = 13.3, 10.1 Hz, 1H), 4.62 (q, *J* = 6.8 Hz, 1H), 4.52 (td, *J* = 9.9, 4.9 Hz, 1H), 4.41 (s, 1H), 4.29-4.22 (m, 4H), 4.04 (t, *J* = 9.0 Hz, 1H), 3.99 (dd, *J* = 8.7, 7.3 Hz, 1H), 3.89 (t, *J* = 8.4 Hz, 1H), 3.80 (t, *J* = 7.5 Hz, 1H), 3.68-3.66 (m, 2H), 3.66-3.56 (m, 4H), 3.32-3.28 (m, 1H), 3.23-3.18 (m, 1H), 3.09-3.05 (m, 1H), 2.85 (t, *J* = 2.6 Hz, 1H), 2.71-2.67 (m, 1H), 2.56 (ddd, *J* = 16.9, 10.0, 2.6 Hz, 1H), 2.32-2.27 (m, 1H), 2.25-2.19 (m, 2H), 2.09-2.05 (m, 1H), 2.02-1.94 (m, 2H), 1.93-1.83 (m, 6H), 1.81-1.68 (m, 4H), 1.65-1.57 (m, 4H), 1.57-1.50 (m, 3H), 1.49-1.36 (m, 6H), 1.30-1.25 (m, 1H), 1.23 (s, 1H), 1.22-1.17 (m, 1H), 1.16-1.09 (m, 2H), 1.07-1.00 (m, 1H), 0.94-0.90 (m, 1H), 0.90-0.88 (m, 9H), 0.86 (d, *J* = 6.6 Hz, 3H), 0.85-0.80 (m, 24H), 0.78 (d, *J* = 6.7 Hz, 3H), 0.71 (t, *J* = 7.4 Hz, 3H); $^{13}$C NMR (176 MHz, DMSO) δ 173.4, 172.7, 172.2, 172.2, 172.2, 171.9, 171.8, 171.5, 171.4, 171.2, 171.1, 167.3, 161.7, 141.4, 99.3, 81.7, 72.2, 68.8, 65.4, 64.0, 62.6, 61.7, 60.0, 59.4, 58.8, 58.2, 56.7, 52.4, 50.2, 49.2, 48.7, 47.0, 46.2, 41.2, 40.9, 38.1, 37.5, 36.2, 35.7, 35.4, 32.5, 29.6, 28.5, 26.3, 25.7, 25.6, 25.1, 24.9, 24.7, 24.6, 24.0, 23.2, 22.5, 22.1, 21.4, 21.3, 15.6, 15.4, 15.3, 15.3, 14.5, 14.0, 11.3, 11.2, 10.5, 10.3, 9.9;

HRMS (ESI): *m/z* calculated for $[M+H]^+$: 1375.8712, found: 1375.8708 (-0.3 ppm); calculated for $[M+2H]^{2+}$: 688.4392, found: 688.4384 (-1.2 ppm); $MS^2$ HRMS (ESI, HCD): *m/z* calculated for ring fragment $[M_{ring}+H]^+$: 940.5866, found: 940.5852 (-1.5 ppm).

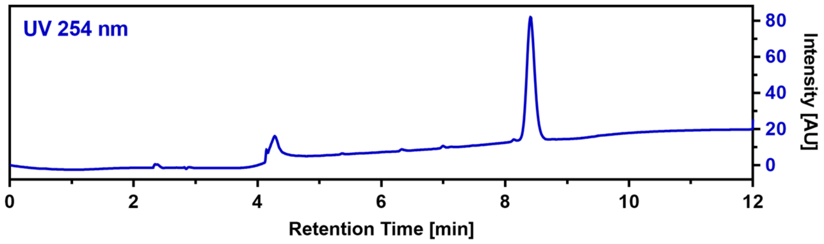

**CalB_C5Pra (83)**

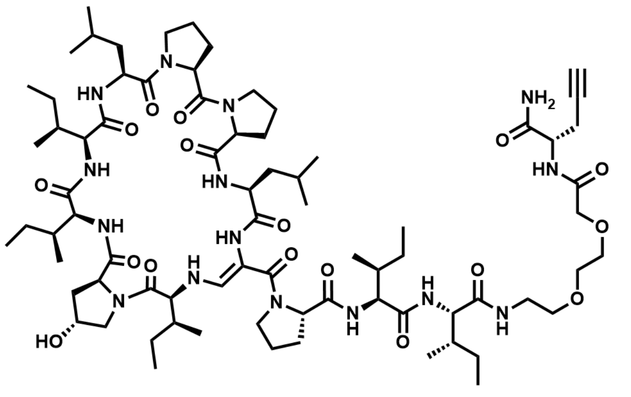

$C_{76}H_{125}N_{15}O_{17}$ M = 1520.92 g/mol

The linear precursor peptide of **83** was prepared via microwave-assisted SPPS using Rink amide PS resin at a scale of 0.1 mmol initial resin capacity. After purification by HPLC (solvent A: $H_2O$, acidified with 0.1% TFA; solvent B: ACN, acidified with 0.1% TFA), 48 mg (31 µmol, 31%) of the peptide precursor was obtained as a white powder. For the oxidation of the serine residue and concomitant cyclization, the peptide was dissolved in 10 mL of ACN and 3 equivalents of DMP were added. The resulting suspension was shaken for 2.5 hours at RT and immediately purified by HPLC (solvent A: $H_2O$; solvent B: ACN) to give 6.6 mg (4.3 µmol, 13.9%) of **83** in a total yield of 4.3% (calculated in relation to initial resin capacity) as a white powder.

HRMS (ESI): *m/z* calculated for $[M+H]^+$: 1520.9451, found: 1520.9440 (-0.7 ppm); calculated for $[M+2H]^{2+}$: 760.9762, found: 760.9751 (-1.4 ppm); $MS^2$ HRMS (ESI, HCD): *m/z* calculated for ring fragment $[M_{ring}+H]^+$: 940.5866, found: 940.5825 (-4.4 ppm).

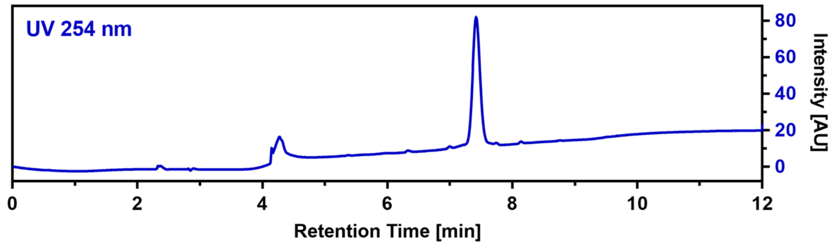

**CalB_R3phLeu (84)**

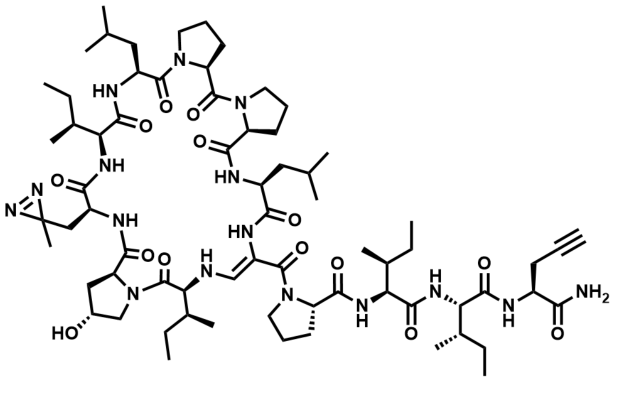

$C_{69}H_{110}N_{16}O_{14}$ M = 1387.74 g/mol

The linear precursor peptide of **84** was prepared via microwave-assisted SPPS up to the R2 position using Rink amide PS resin at a scale of 0.1 mmol initial resin capacity. The loading of the resin was quantified through a spectrometric Fmoc-Test and a manual coupling of photo-leucine was performed using 4 equivalents of amino acid combined with 2-(1*H*-benzotriazol-1-yl)-1,1,3,3-tetramethyluronium hexafluorophosphate (HBTU) and hydroxybenzotriazole (HOBt) (each 4 equivalents) as well as 5 equivalents of DIPEA in 10 mL DMF by shaking the suspension over night at room temperature. The resin was then thoroughly washed with DMF and a second round of microwave-assisted SPPS was performed for completion of the peptide sequence. After purification by HPLC (solvent A: $H_2O$, acidified with 0.1% TFA; solvent B: ACN, acidified with 0.1% TFA), 38 mg

(27 µmol, 27%) of the peptide precursor was obtained as a white powder. For the oxidation of the serine residue and concomitant cyclization, 36 mg of peptide was dissolved in 5 mL of ACN and 4 equivalents of DMP were added. The resulting suspension was shaken for 4.5 hours at RT and immediately purified by HPLC (solvent A: $H_2O$; solvent B: ACN) to give 11.8 mg (8.5 µmol, 33.6%) of **84** in a total yield of 8.5% (calculated in relation to initial resin capacity) as a white powder.

$^1$H NMR (400 MHz, DMSO) δ 9.13 (d, $J$ = 6.4 Hz, 1H), 8.19 (s, 1H), 8.06 (d, $J$ = 8.2 Hz, 1H), 7.88 (d, $J$ = 8.6 Hz, 1H), 7.51 (d, $J$ = 8.8 Hz, 1H), 7.49 (d, $J$ = 7.4 Hz, 1H), 7.26 (d, $J$ = 7.9 Hz, 1H), 7.21 (s, 1H), 7.00 (d, $J$ = 13.1 Hz, 1H), 6.74 (s, 1H), 6.67 (d, $J$ = 10.2 Hz, 1H), 5.60 (dd, $J$ = 13.1, 10.2 Hz, 1H), 5.43 (d, $J$ = 3.0 Hz, 1H), 4.67 (q, $J$ = 7.0 Hz, 1H), 4.52-4.46 (m, 1H), 4.45 (s, 1H), 4.36-4.31 (m, 1H), 4.31-4.21 (m, 3H), 4.07-3.95 (m, 3H), 3.92 (t, $J$ = 8.3 Hz, 1H), 3.79-3.73 (m, 1H), 3.72-3.64 (m, 2H), 3.63-3.51 (m, 5H), 3.50-3.44 (m, 1H), 3.27-3.18 (m, 1H), 2.85 (t, $J$ = 2.6 Hz, 1H), 2.73-2.65 (m, 1H), 2.60-2.52 (m, 1H), 2.42-2.31 (m, 1H), 2.27-2.17 (m, 3H), 2.11-2.02 (m, 3H), 2.01-1.93 (m, 4H), 1.92-1.83 (m, 2H), 1.81-1.76 (m, 1H), 1.75-1.70 (m, 1H), 1.70-1.54 (m, 7H), 1.52-1.40 (m, 5H), 1.39-1.33 (m, 2H), 1.25-1.13 (m, 2H), 1.10-1.04 (m, 1H), 1.02 (s, 3H), 1.00-0.95 (m, 1H), 0.92 (d, $J$ = 6.3 Hz, 3H), 0.90 (d, $J$ = 6.3 Hz, 3H), 0.88 (d, $J$ = 6.6 Hz, 3H), 0.86-0.79 (m, 21H), 0.79-0.77 (m, 1H), 0.76-0.70 (m, 6H); HRMS (ESI): $m/z$ calculated for $[M+H]^+$: 1387.8460, found: 1387.8449 (-0.8 ppm); calculated for $[M+2H]^{2+}$: 694.4267, found: 694.4256 (-1.6 ppm); $MS^2$ HRMS (ESI, HCD): $m/z$ calculated for ring fragment $[M_{ring}+H]^+$: 952.5615, found: 952.5581 (-3.6 ppm).

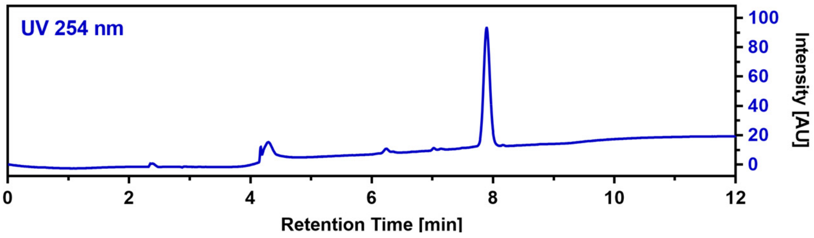

**CalB_R5phLeu (85)**

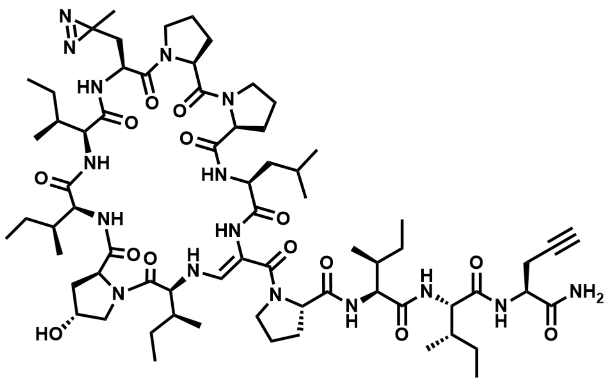

$C_{69}H_{110}N_{16}O_{14}$ M = 1387.74 g/mol

The linear precursor peptide of **85** was prepared via microwave-assisted SPPS using Rink amide PS resin at a scale of 0.1 mmol initial resin capacity. After purification by HPLC (solvent A: $H_2O$, acidified with 0.1% TFA; solvent B: ACN, acidified with 0.1% TFA), 42 mg (30 µmol, 30%) of the peptide precursor was obtained as a white powder. For the oxidation of the serine residue and concomitant cyclization, the peptide was dissolved in 5 mL of ACN and 4 equivalents of DMP were added. The resulting suspension was shaken for 2 hours at RT and immediately purified by HPLC (solvent A: $H_2O$; solvent B: ACN) to give 11.2 mg (8.1 µmol, 26.8%) of **85** in a total yield of 8.1% (calculated in relation to initial resin capacity) as a white powder.

HRMS (ESI): $m/z$ calculated for $[M+H]^+$: 1387.8460, found: 1387.8451 (-0.6 ppm); calculated for $[M+2H]^{2+}$: 694.4267, found: 694.4267 (-2.7 ppm); $MS^2$ HRMS (ESI, HCD): $m/z$ calculated for ring fragment $[M_{ring}+H]^+$: 952.5615, found: 952.5582 (-3.5 ppm).

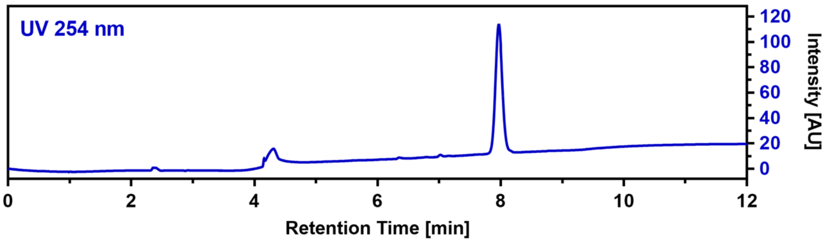

**CalB_R8phLeu (86)**

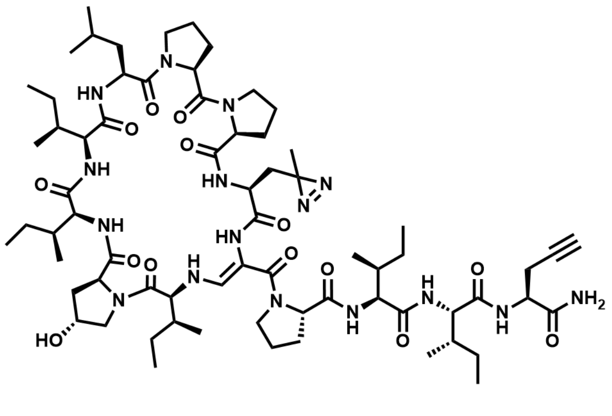

$C_{69}H_{110}N_{16}O_{14}$ M = 1387.74 g/mol

The linear precursor peptide of **86** was prepared via microwave-assisted SPPS up to the R2 position using Rink amide PS resin at a scale of 0.1 mmol initial resin capacity. The loading of the resin was quantified through a spectrometric Fmoc-Test and a manual coupling of photo-leucine was performed using 4 equivalents of amino acid combined with HBTU and HOBt (each 4 equivalents) as well as 5 equivalents of DIPEA in 10 mL DMF by shaking the suspension over night at room temperature. The resin was then thoroughly washed with DMF and a second round of microwave-assisted SPPS was performed for completion of the peptide sequence. After purification by HPLC (solvent A: $H_2O$, acidified with 0.1% TFA; solvent B: ACN, acidified with 0.1% TFA), 19 mg (13 µmol, 13%) of the peptide precursor was obtained as a white powder. For the oxidation of the serine residue and concomitant cyclization, the peptide was dissolved in 5 mL of ACN and 4 equivalents of DMP were added. The resulting suspension was shaken for 6.5 hours at RT and immediately purified by HPLC (solvent A: $H_2O$; solvent B: ACN) to give 1.4 mg (1.0 µmol, 7.6%) of **86** in a total yield of 1.0% (calculated in relation to initial resin capacity) as a white powder.

HRMS (ESI): *m/z* calculated for $[M+H]^+$: 1387.8460, found: 1387.8451 (-0.6 ppm); calculated for $[M+2H]^{2+}$: 694.4267, found: 694.4256 (-1.6 ppm); $MS^2$ HRMS (ESI, HCD): *m/z* calculated for ring fragment $[M_{ring}+H]^+$: 952.5615, found: 952.5597 (-1.9 ppm).

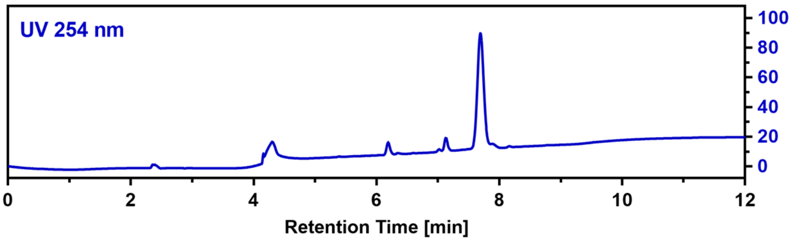

**CalB_C4Pra(Cy3) (87)**

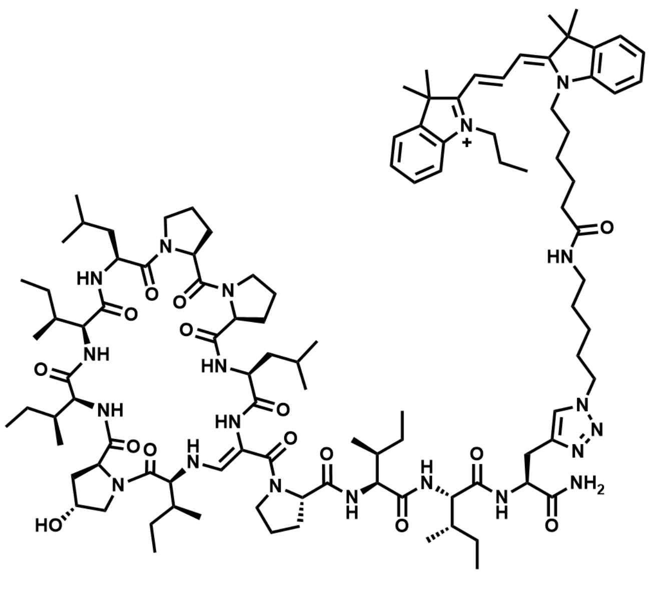

$C_{107}H_{165}N_{20}O_{15}^+$ M = 1971.62 g/mol

For the copper-catalyzed azide-alkyne cycloaddition (CuAAC), 12.2 mg of **82** were dissolved in 1 mL of ACN and 1.8 equivalents of Cy3-azide, 0.4 eq. Tris[(1-benzyl-1*H*-1,2,3-triazol-4-yl)methyl]amin) (TBTA) as well as 4 eq. of DIPEA were added. To start the reaction, 0.6 eq. of CuI were added. Immediately, the micro reaction vessel was flooded with argon and the mixture was shaken for 26 hours at room temperature. Another 0.1 equivalents of CuI were added and shaking was continued overnight. The crude product was purified by HPLC (solvent A: $H_2O$, acidified with 0.1% TFA; solvent B: ACN, acidified with 0.1% TFA) to give 13.3 mg (6.4 µmol) of **87** in a total yield of 72% as a deep purple powder.

HRMS (ESI): *m/z* calculated for $M^+$: 1971.2792, found: 1971.2798 (0.3 ppm); calculated for $[M^+ + H]^{2+}$: 986.1432, found: 986.1426 (-0.6 ppm); $MS^2$ HRMS (ESI, HCD): *m/z* calculated for ring fragment $[M^+_{ring} + H]^+$: 940.5866, found: 940.5839 (-2.9 ppm).

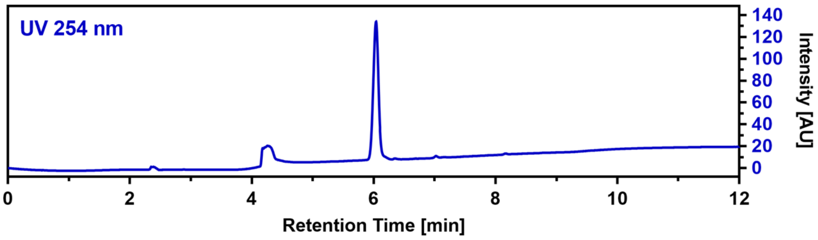

**CalB_C4Pra(Rhodamine) (88)**

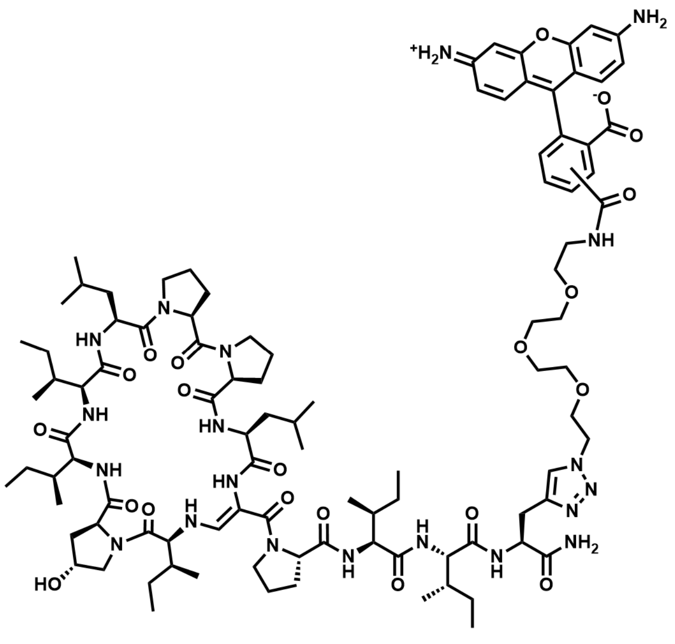

$C_{99}H_{144}N_{20}O_{21}$ M = 1950.36 g/mol

For CuAAC, 1.0 mg of 5/6-Carboxy-rhodamine-azide (CLK-AZ105, Jena Bioscience) was dissolved in 300 µL ACN and 1.6 eq. of **82**, 0.4 eq. TBTA as well as 4 eq. of DIPEA were added. To start the reaction, 0.6 eq. of CuI were added. Immediately, the micro reaction vessel was flooded with argon and the mixture was shaken for 24 hours at room temperature. The crude product was purified by HPLC (solvent A: $H_2O$, acidified with 0.1% TFA; solvent B: ACN, acidified with 0.1% TFA) to give 1.4 mg (0.7 µmol) of **88** in a total yield of 25% as a brown powder.

HRMS (ESI): *m/z* calculated for $M^+$: 1951.0922, found: 1951.0919 (-0.2 ppm); calculated for $[M+H]^{2+}$: 976.0497, found: 976.0497 (-0.5 ppm); $MS^2$ HRMS (ESI, HCD): *m/z* calculated for ring fragment $[M_{ring}+H]^+$: 940.5866, found: 940.5829 (-3.9 ppm).

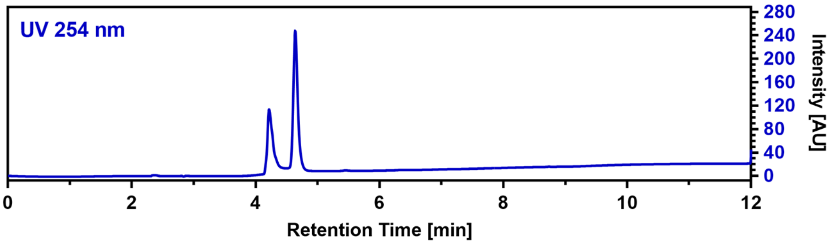

#### Cy3-Control (89)

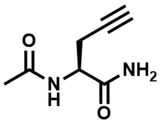

$C_7H_{10}N_2O_2$
M = 154.17 g/mol

*N*-Fmoc-L-propargylglycin (CC05047, Carbolution) was amidated and acetylated via an automated microwave-assisted protocol in a Liberty Blue peptide synthesizer (CEM) using Rink-Amide PS resin and the obtained amino acid building block **89a** was purified by HPLC (solvent A: $H_2O$, acidified with 0.1% TFA; solvent B: ACN, acidified with 0.1% TFA). $^1H$ NMR (400 MHz, DMSO) δ 8.04 (d, *J* = 8.2 Hz, 1H), 7.42 (s, 1H), 7.12 (s, 1H), 4.32 (td, *J* = 8.0, 5.6 Hz, 1H), 2.82 (t, *J* = 2.7 Hz, 1H), 2.58-2.52 (m, 1H), 2.46-2.36 (m, 1H), 1.86 (s, 3H); $^{13}C$ NMR (101 MHz, DMSO) δ 171.82, 169.20, 80.88, 72.69, 51.25, 22.53, 21.76; HRMS (ESI): *m/z* calculated for $[M+H]^+$: 155.0815, found: 155.0813 (-1.3 ppm);

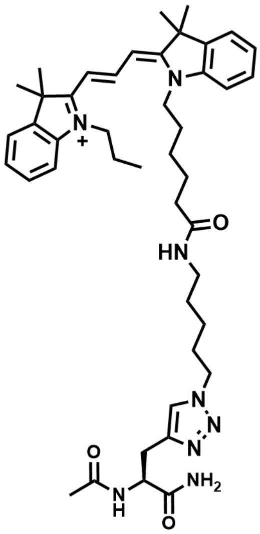

$C_{44}H_{61}N_8O_3^+$
M = 750.02 g/mol

For CuAAC, 5.0 mg of Cy3-Azide (dissolved in DMSO) was mixed with 1.5 eq. of **89a** and 0.2 eq. TBTA as well as 4 eq. of DIPEA were added. To start the reaction, 0.6 eq. of CuI were added. Immediately, the micro reaction vessel was flooded with argon and the mixture was shaken for 24 hours at room temperature. The crude product was purified by HPLC (solvent A: $H_2O$, acidified with 0.1% TFA; solvent B: ACN, acidified with 0.1% TFA) to give 6.1 mg (7.1 µmol) of **90** in a total yield of 84% as a deep purple powder.

$^1H$ NMR (400 MHz, DMSO) δ 8.36 (t, *J* = 13.4 Hz, 1H), 8.01 (d, *J* = 8.4 Hz, 1H), 7.76-7.70 (m, 2H), 7.64 (dd, *J* = 7.5, 4.0 Hz, 2H), 7.52-7.40 (m, 4H), 7.38 (s, 1H), 7.33-7.26 (m, 2H), 7.04 (s, 1H), 6.52 (dd, *J* = 13.4, 8.7 Hz, 2H), 4.43 (td, *J* = 8.7, 5.1 Hz, 1H), 4.27 (t, *J* = 7.0 Hz, 2H), 4.11 (t, *J* = 7.3 Hz, 4H), 3.04 (dd, *J* = 14.7, 5.1 Hz, 1H), 3.00-2.92 (m, 2H), 2.82 (dd, *J* = 14.7, 9.0 Hz, 1H), 2.05 (t, *J* = 7.3 Hz, 2H), 1.81-1.79 (m, 3H), 1.78-1.72 (m, 4H), 1.70 (s, 6H), 1.69 (s, 6H), 1.55 (p, *J* = 7.4 Hz, 2H), 1.41-1.29 (m, 4H), 1.19-1.10 (m, 2H), 0.98 (t, *J* = 7.4 Hz, 3H); HRMS (ESI): *m/z* calculated for $M^+$: 749.4861, found: 749.4857 (-0.5 ppm); calculated for $[M+H]^{2+}$: 375.2467, found: 375.2463 (-1.1 ppm).

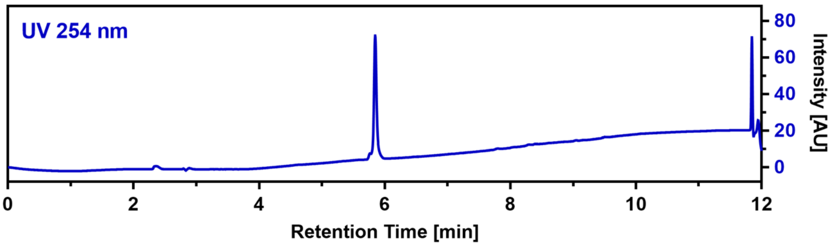

**CalB_R3Pra(Cy3) (90)**

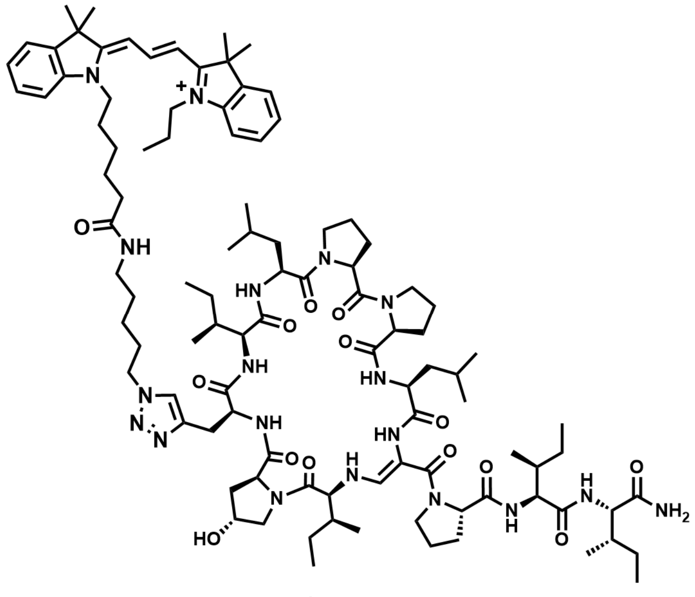

$C_{101}H_{154}N_{19}O_{14}^{+}$ M = 1858.46 g/mol

For CuAAC, 4.3 mg of **78** were dissolved in 100 µL of ACN and 1.2 equivalents of Cy3-azide (500 mM DMSO stock), 0.3 eq. TBTA (5 mM stock) as well as 4 eq. of DIPEA were added. To start the reaction, 0.5 eq. of CuI (10 mM ACN stock) were added. Immediately, the micro reaction vessel was flooded with argon and the mixture was shaken for 4 hours at room temperature. Another 0.1 eq. of CuI were added and shaking was continued overnight. The crude product was purified by HPLC (solvent A: $H_2O$, acidified with 0.1% TFA; solvent B: ACN, acidified with 0.1% TFA) to give 0.6 mg (0.3 µmol) of **90** in a total yield of 9% as a deep purple powder.

HRMS (ESI): *m/z* calculated for $M^+$: 1858.1951, found: 1858.1942 (-0.5 ppm); calculated for $[M^++H]^{2+}$: 929.6012, found: 929.6005 (-0.8 ppm); $MS^2$ HRMS (ESI, HCD): *m/z* calculated for ring fragment $[M^+_{ring}+H]^{2+}$: 758.9758, found 758.9727 (-4.1 ppm).

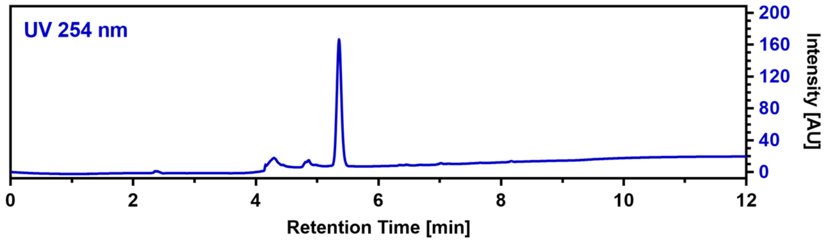

**CalB_R4Pra(Cy3) (91)**

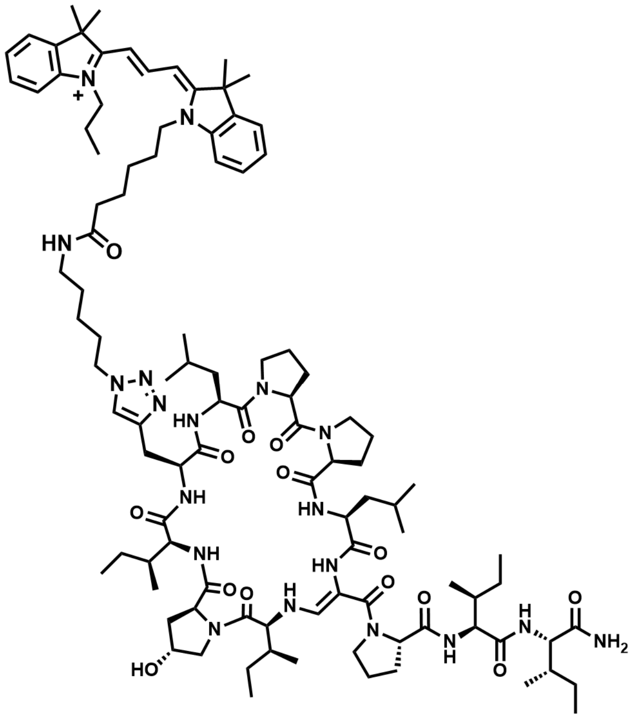

$C_{101}H_{154}N_{19}O_{14}^{+}$ M = 1858.46 g/mol

For CuAAC, 1.2 mg of **79** were dissolved in 100 µL of ACN and 1.3 equivalents of Cy3-azide (500 mM DMSO stock), 0.5 eq. TBTA (5 mM stock) as well as 4 eq. of DIPEA were added. To start the reaction, 0.5 eq. of CuI (10 mM ACN stock) were added. Immediately, the micro reaction vessel was flooded with argon and the mixture was shaken for 4 hours at room temperature. Another 0.25 eq. of Cy3-azide as well as 0.2 eq. of CuI were added and shaking was continued overnight. The crude product was purified by HPLC (solvent A: $H_2O$, acidified with 0.1% TFA; solvent B: ACN, acidified with 0.1% TFA) to give 0.9 mg (0.5 µmol) of **91** in a total yield of 49% as a deep purple powder.
HRMS (ESI): *m/z* calculated for $M^+$: 1858.1951, found: 1858.1937 (-0.8 ppm); calculated for $[M^++H]^{2+}$: 929.6012, found: 929.5993 (-2.0 ppm); $MS^2$ HRMS (ESI, HCD): *m/z* calculated for ring fragment $[M^+_{ring}+H]^{2+}$: 758.9758, found 758.9746 (-1.6 ppm).

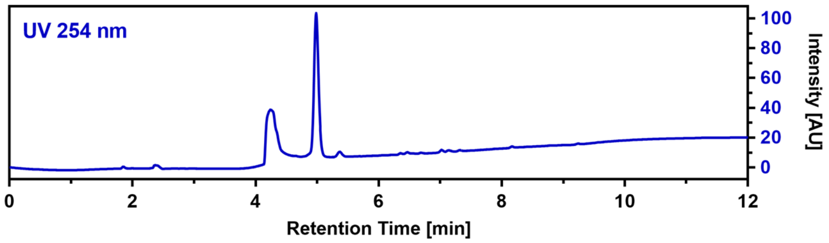

### NMR Spectra

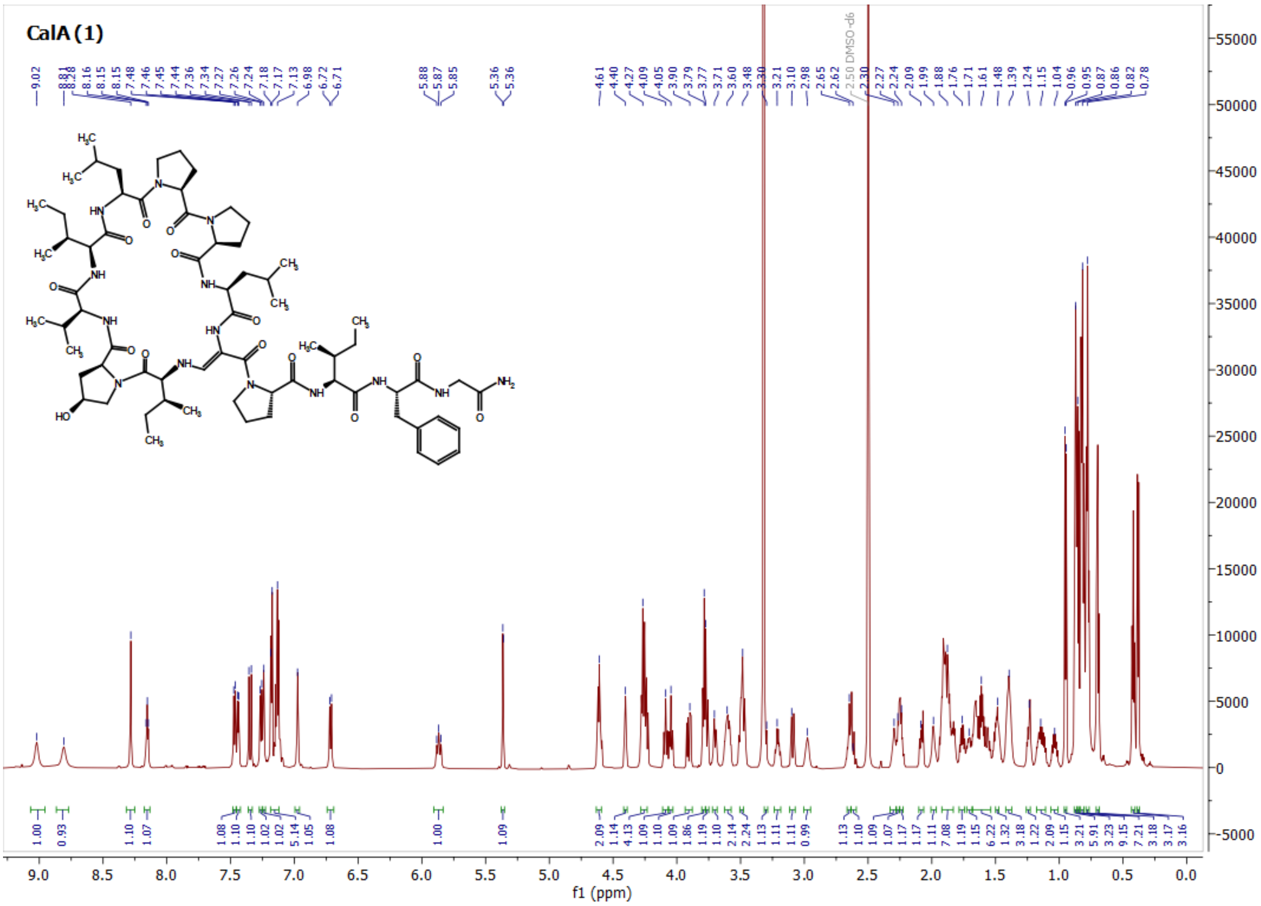

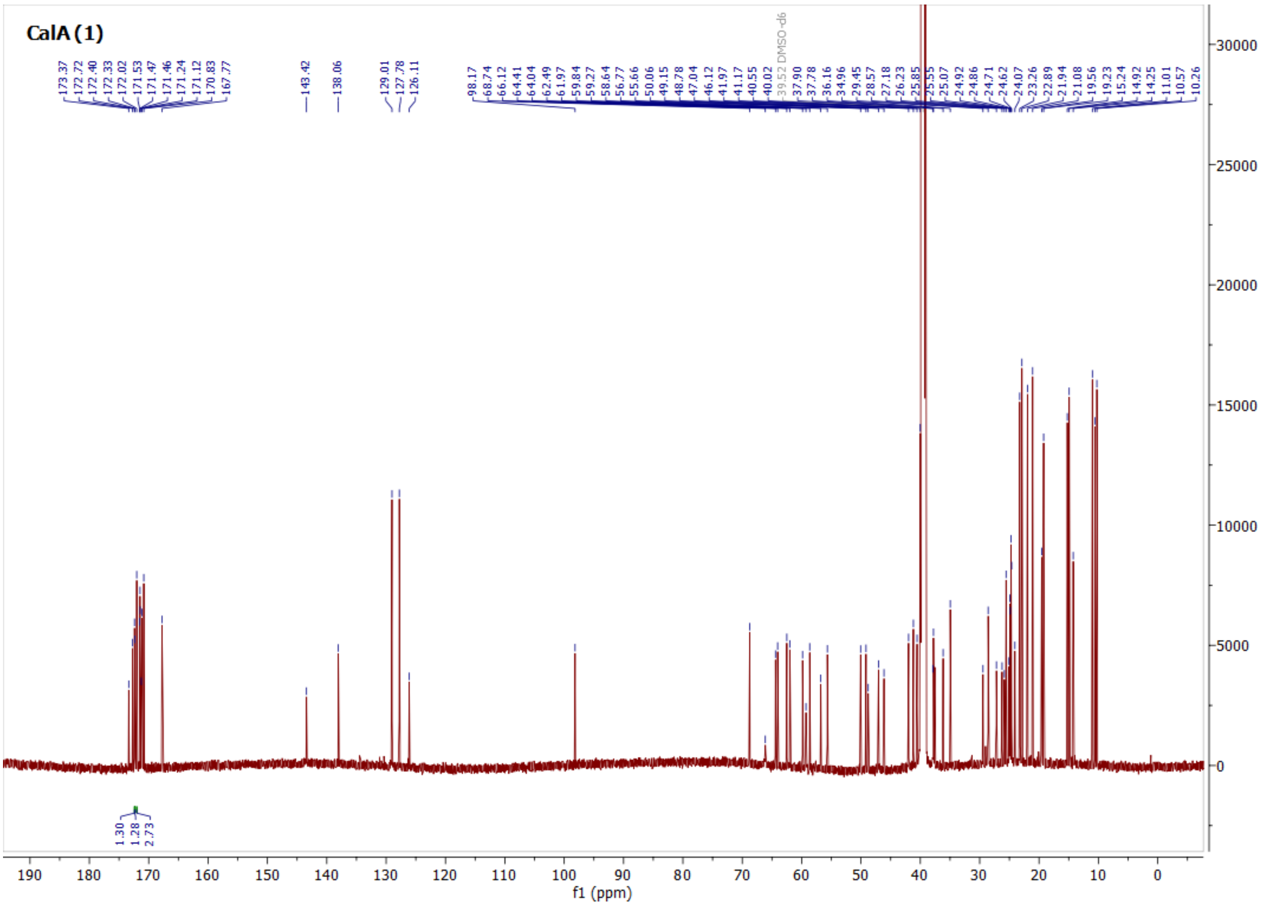

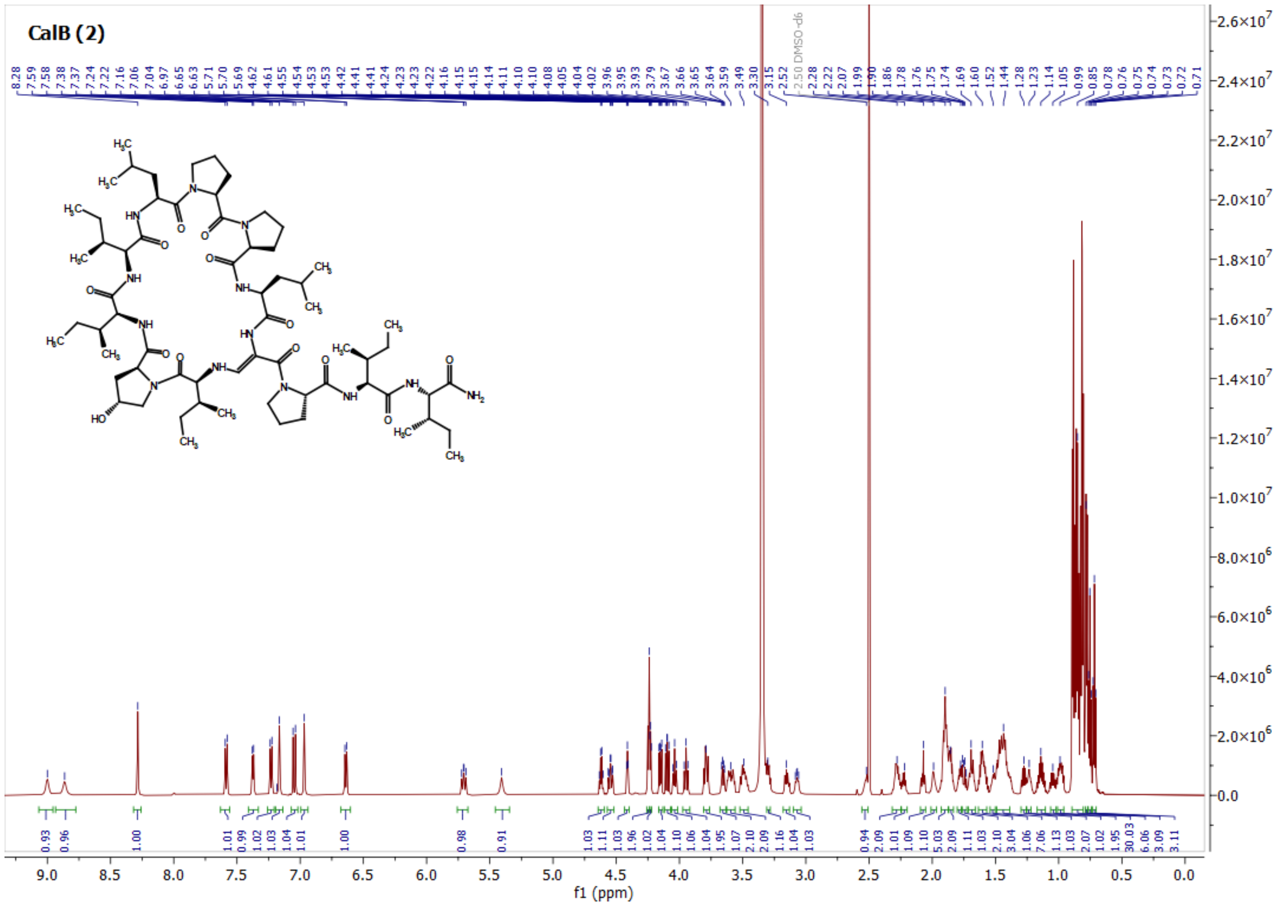
CalB (2)
f1 (ppm)

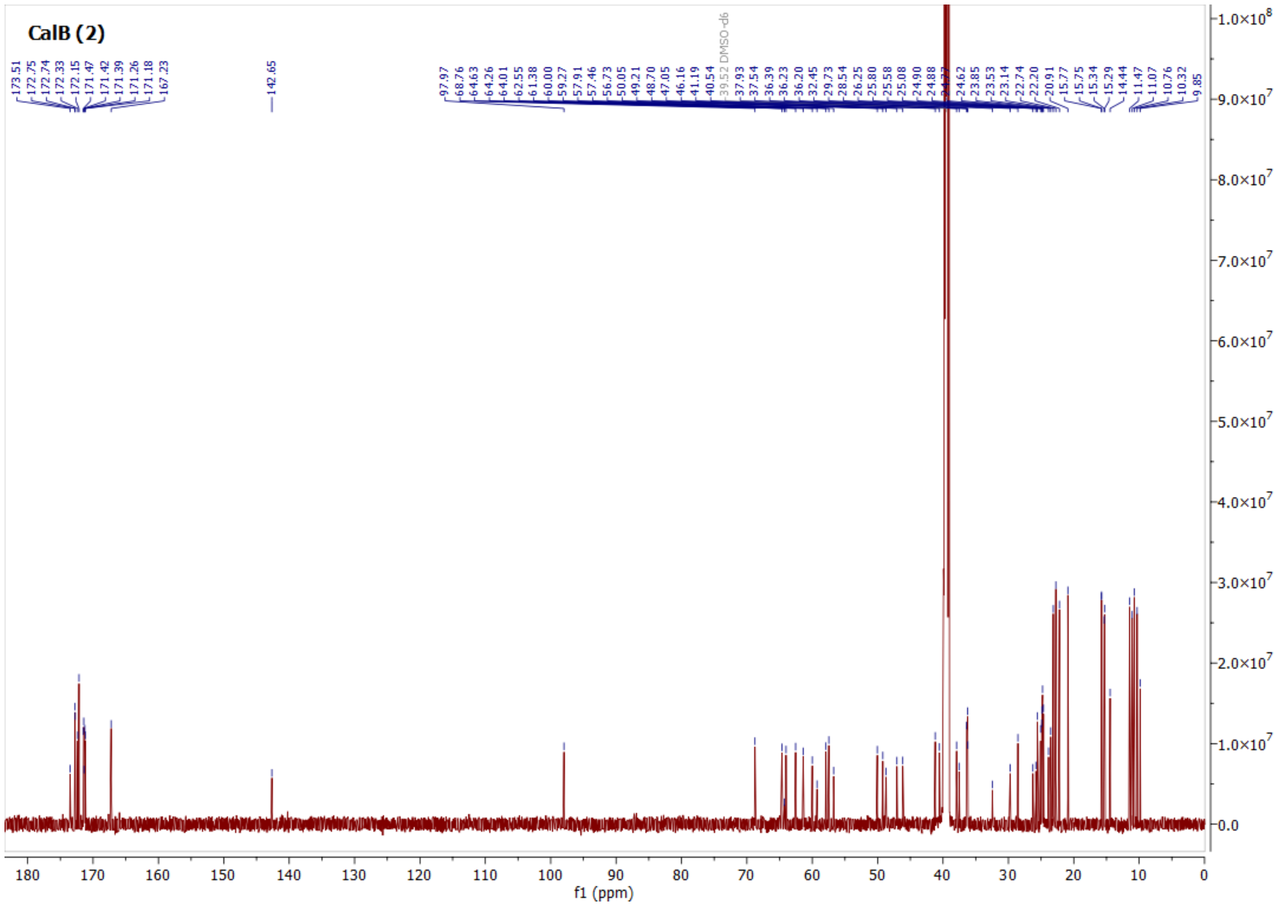
CalB (2)
f1 (ppm)

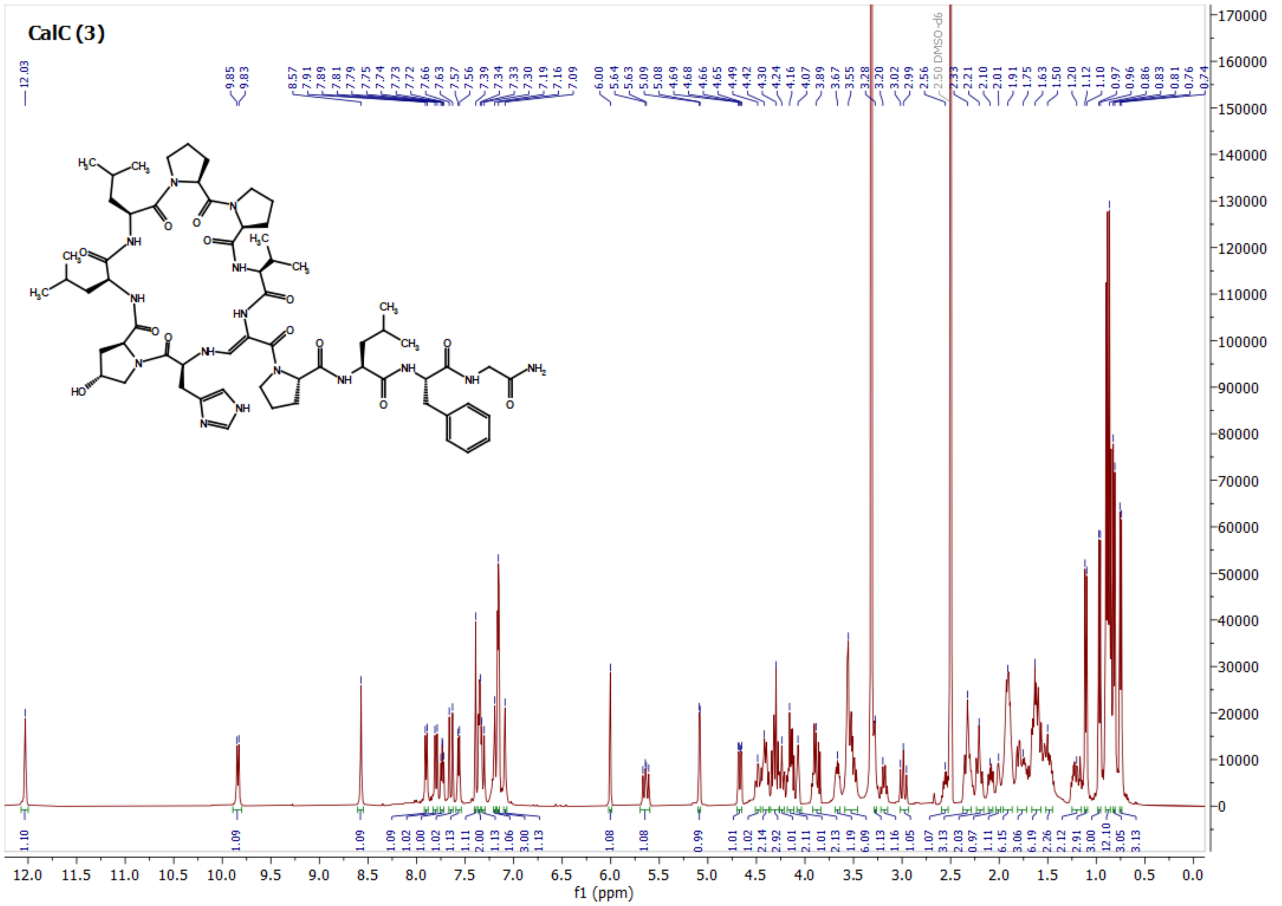
CalC (3)
f1 (ppm)
2.50 DMSO-d6

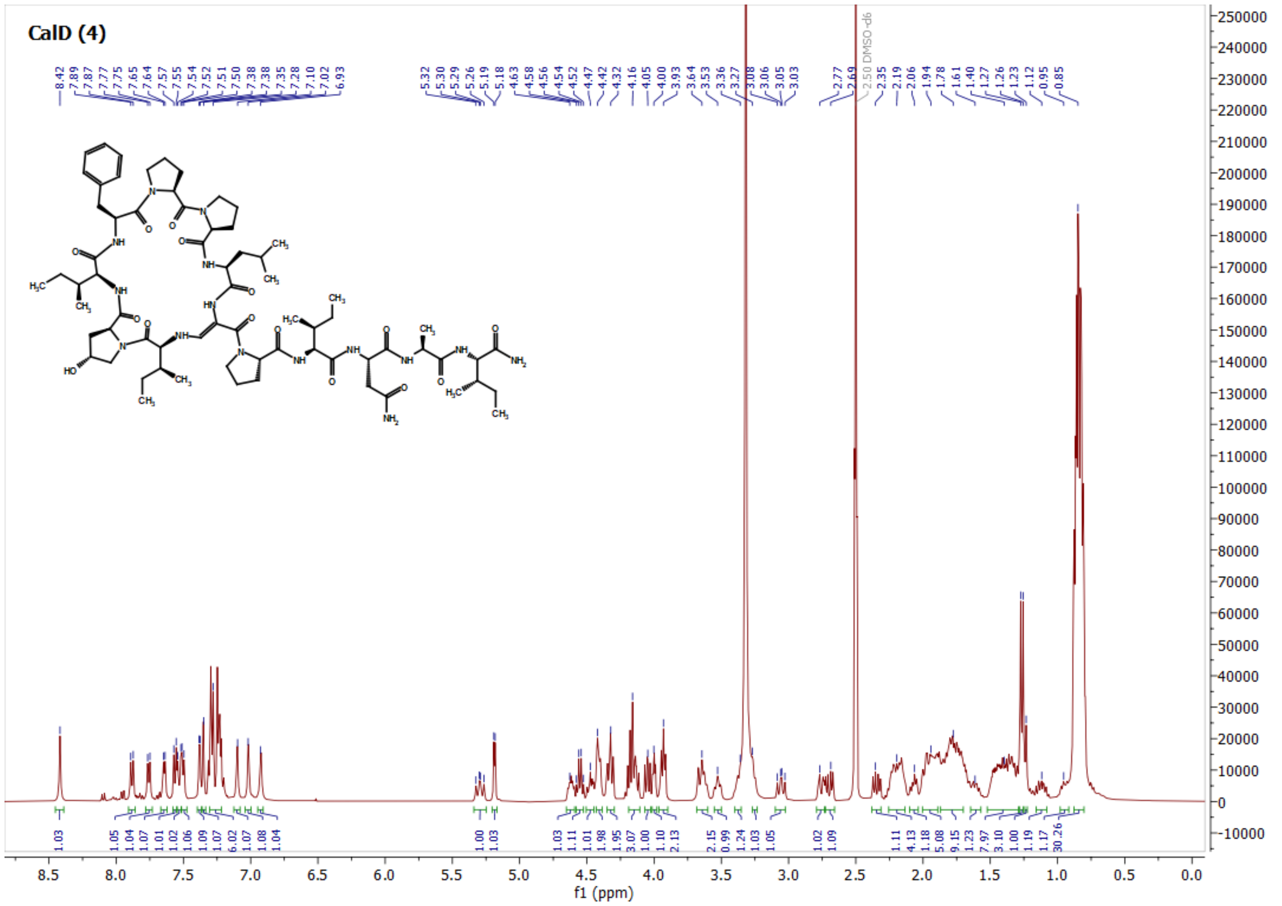
CalD (4)
f1 (ppm)
2.50 DMSO-d6

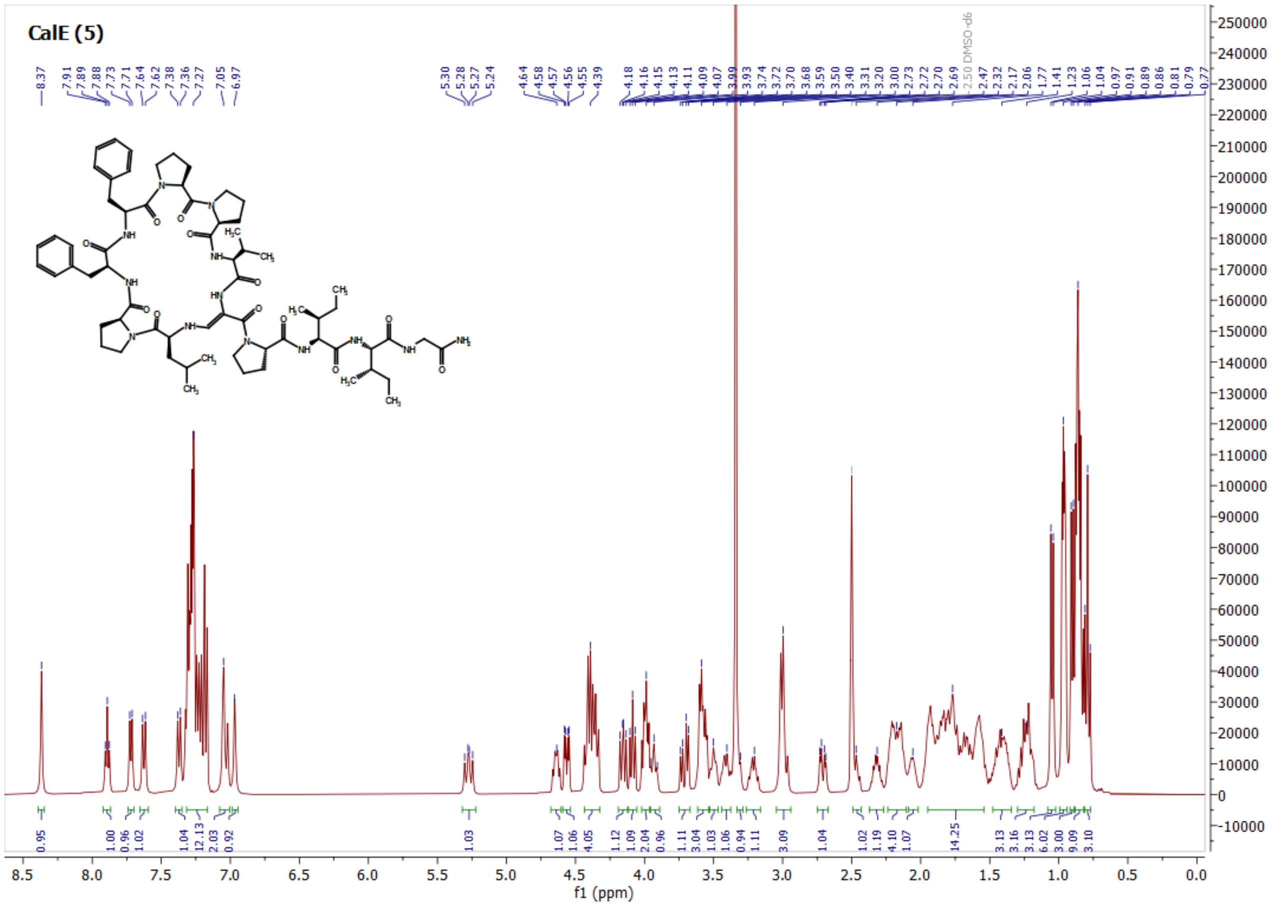
CalE (5)
f1 (ppm)

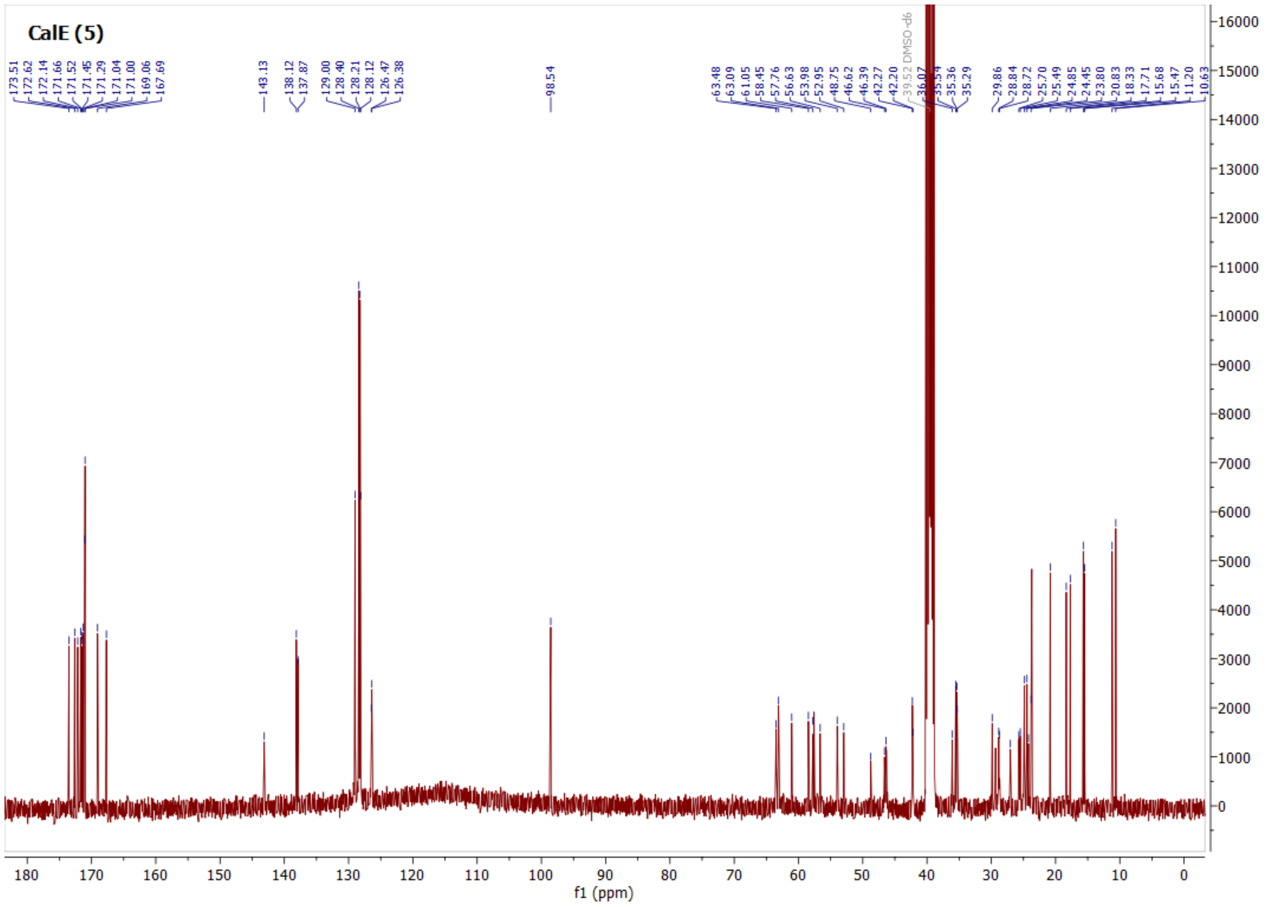
CalE (5)
f1 (ppm)

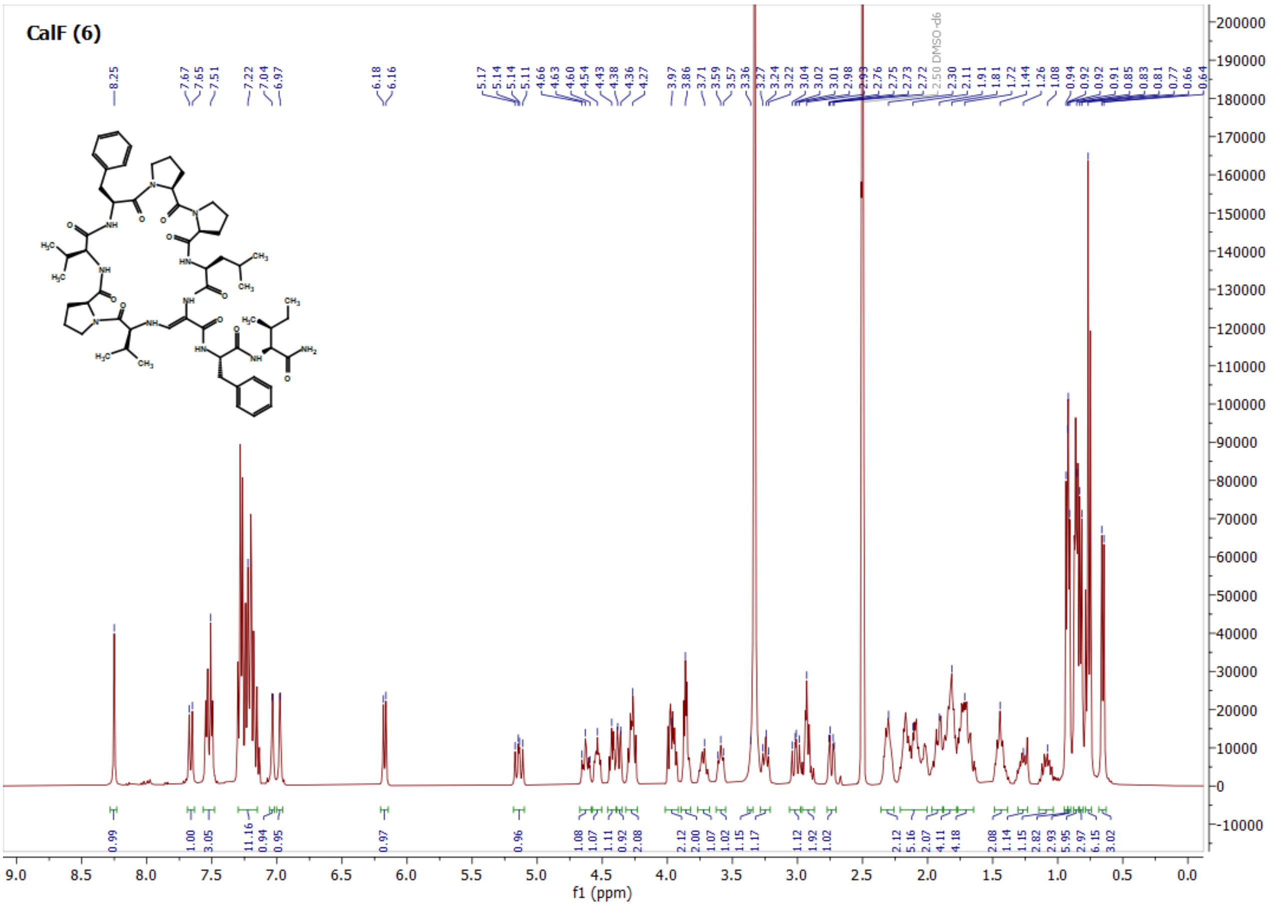
CalF (6)

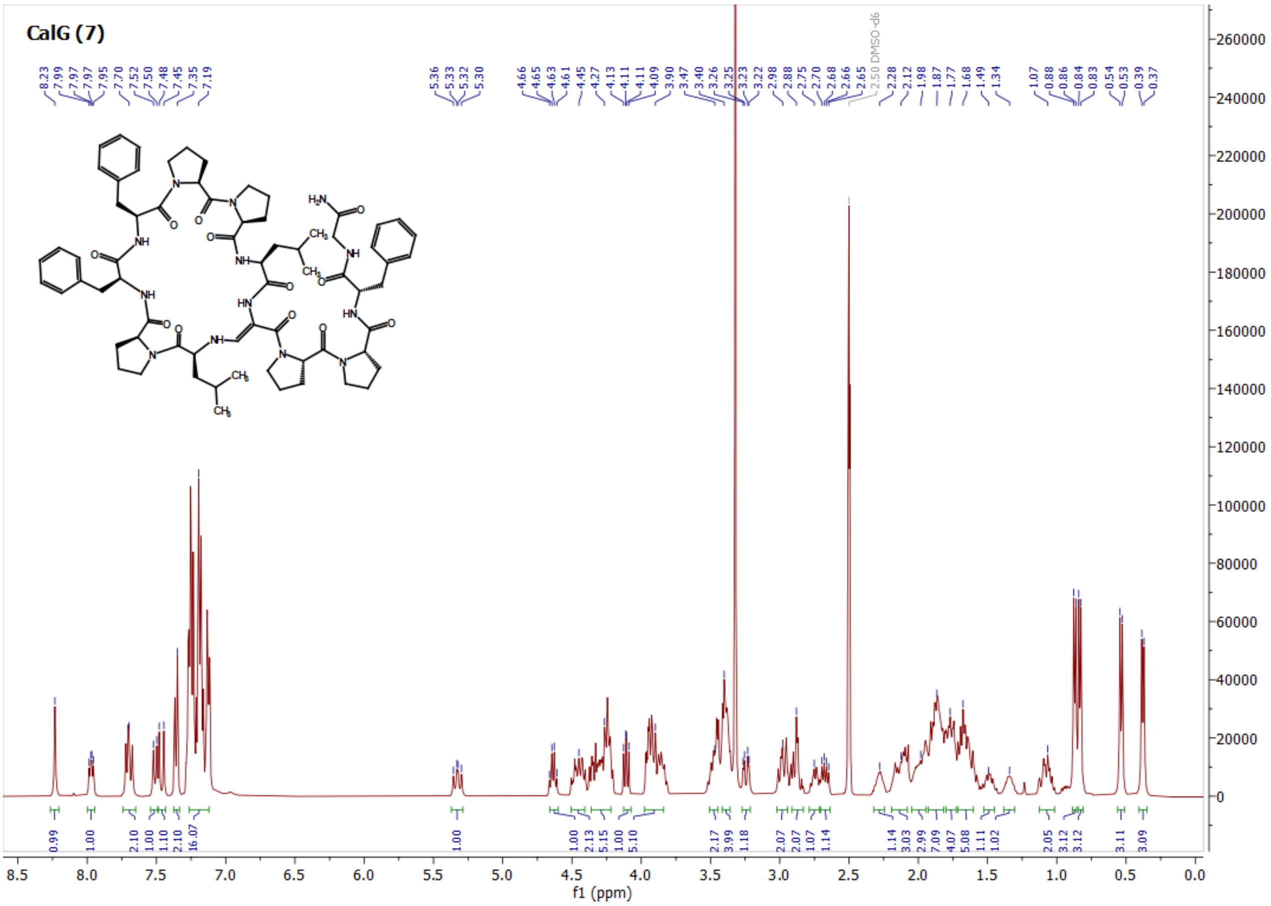
CalG (7)

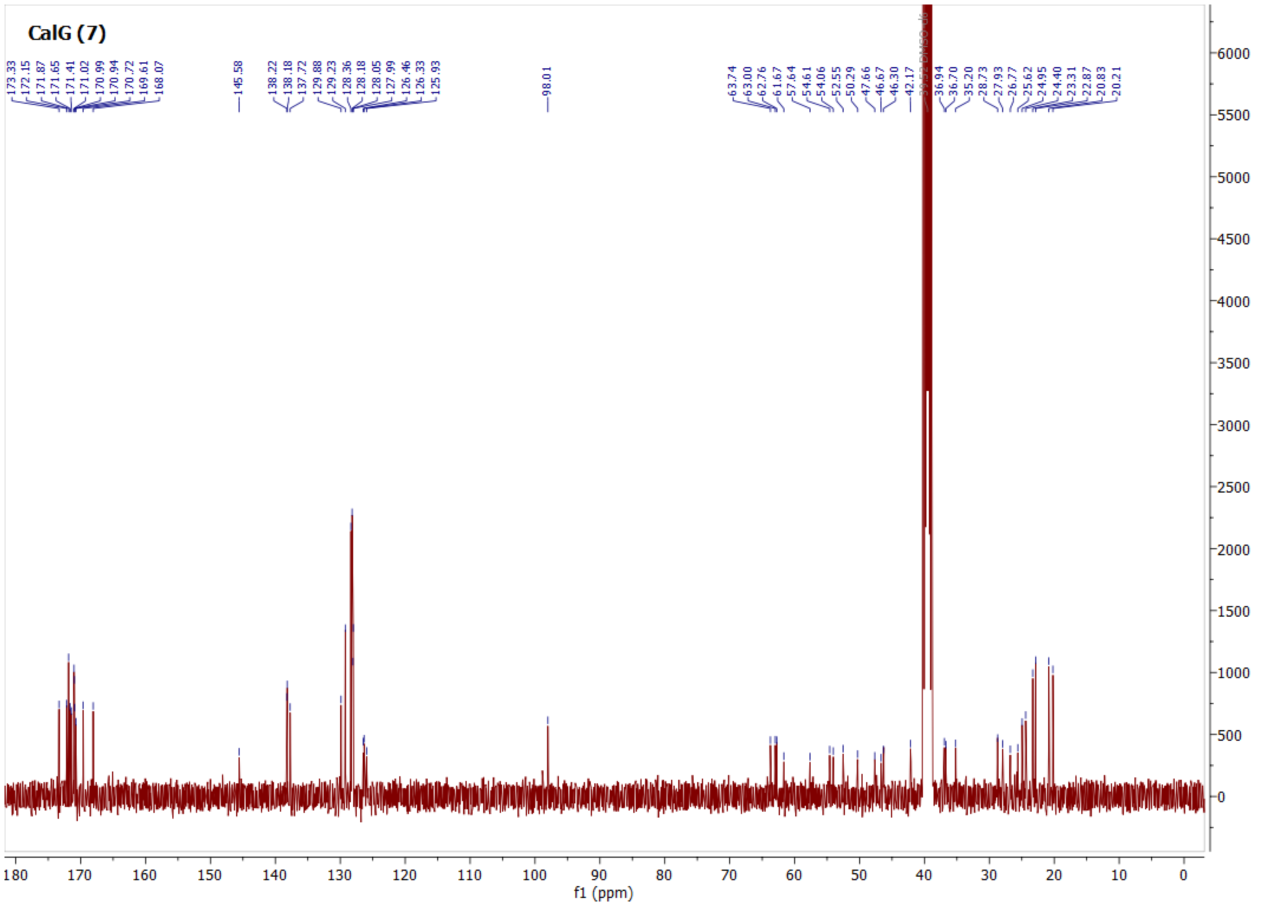
CalG (7)
173.33 172.15 171.87 171.65 171.41 171.02 170.99 170.94 170.72 169.61 168.07
145.58
138.22 138.18 137.72 129.88 129.23 128.36 128.18 128.05 127.99 126.46 126.33 125.93
98.01
63.74 63.00 62.76 61.67 57.64 54.61 54.06 52.55 50.29 47.66 46.67 46.30 42.17
36.94 36.70 35.20 28.73 27.93 26.77 25.62 24.95 24.40 23.31 22.87 20.83 20.21
f1 (ppm)

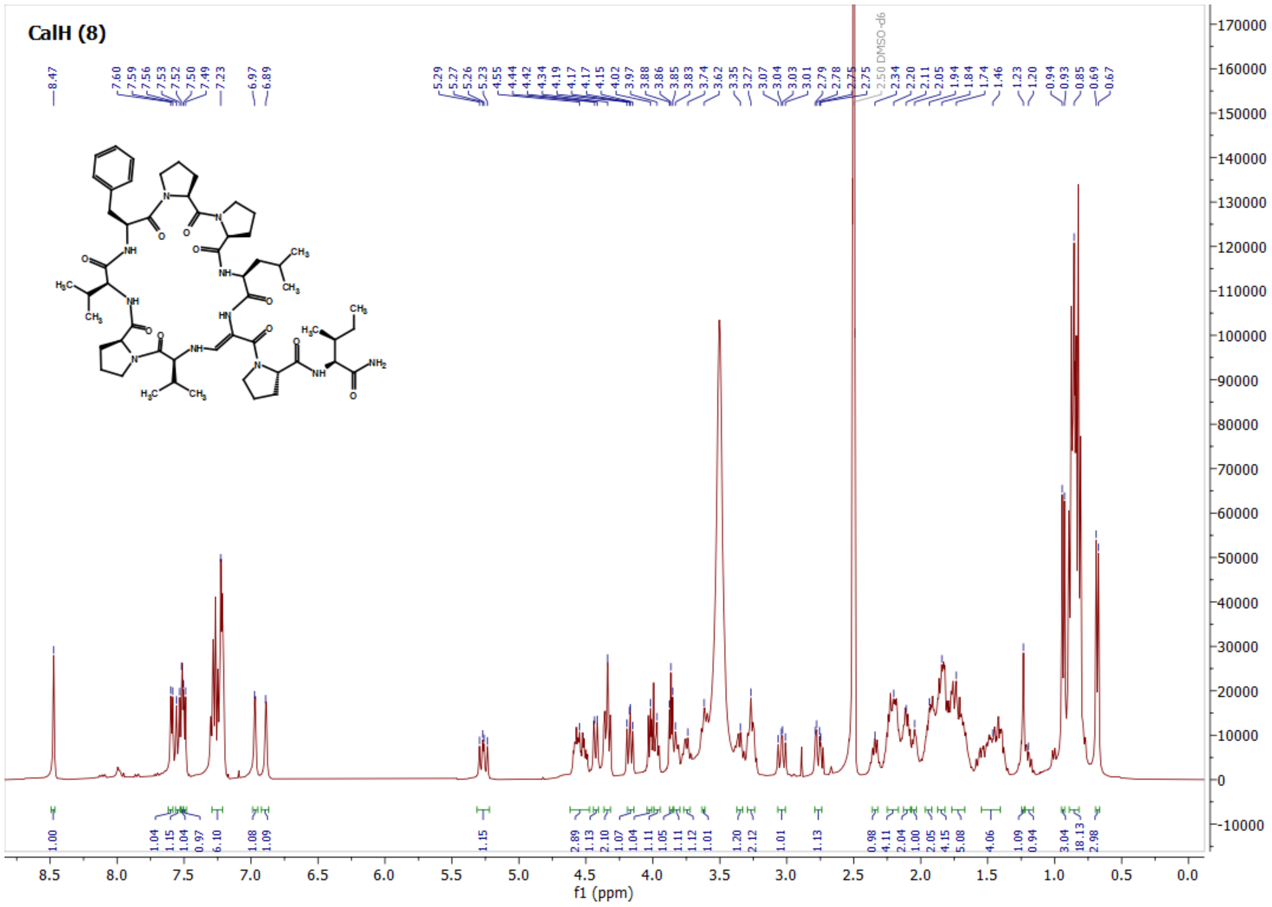
CalH (8)
8.47
7.60 7.59 7.56 7.53 7.52 7.50 7.49 7.23
6.97 6.89
5.29 5.27 5.26 5.23 4.55 4.44 4.42 4.34 4.19 4.17 4.17 4.15 4.02 3.97 3.88 3.86 3.85 3.83 3.74 3.62 3.35 3.27 3.07 3.04 3.03 3.01 2.79 2.78 2.75
2.50 DMSO-d6
2.34 2.20 2.11 2.05 1.94 1.84 1.74 1.46 1.23 1.20 0.94 0.93 0.85 0.69 0.67
f1 (ppm)

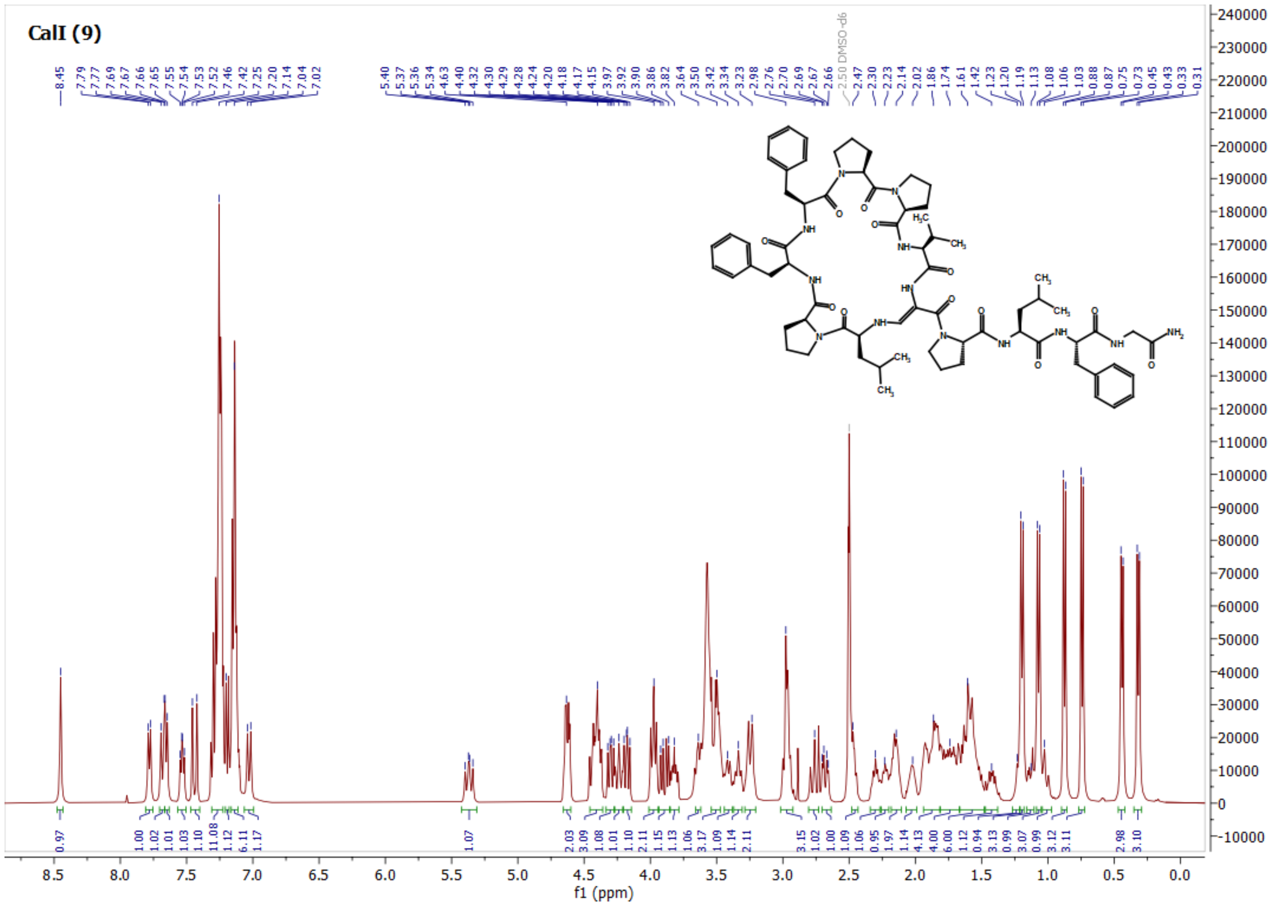
CalI (9)
f1 (ppm)

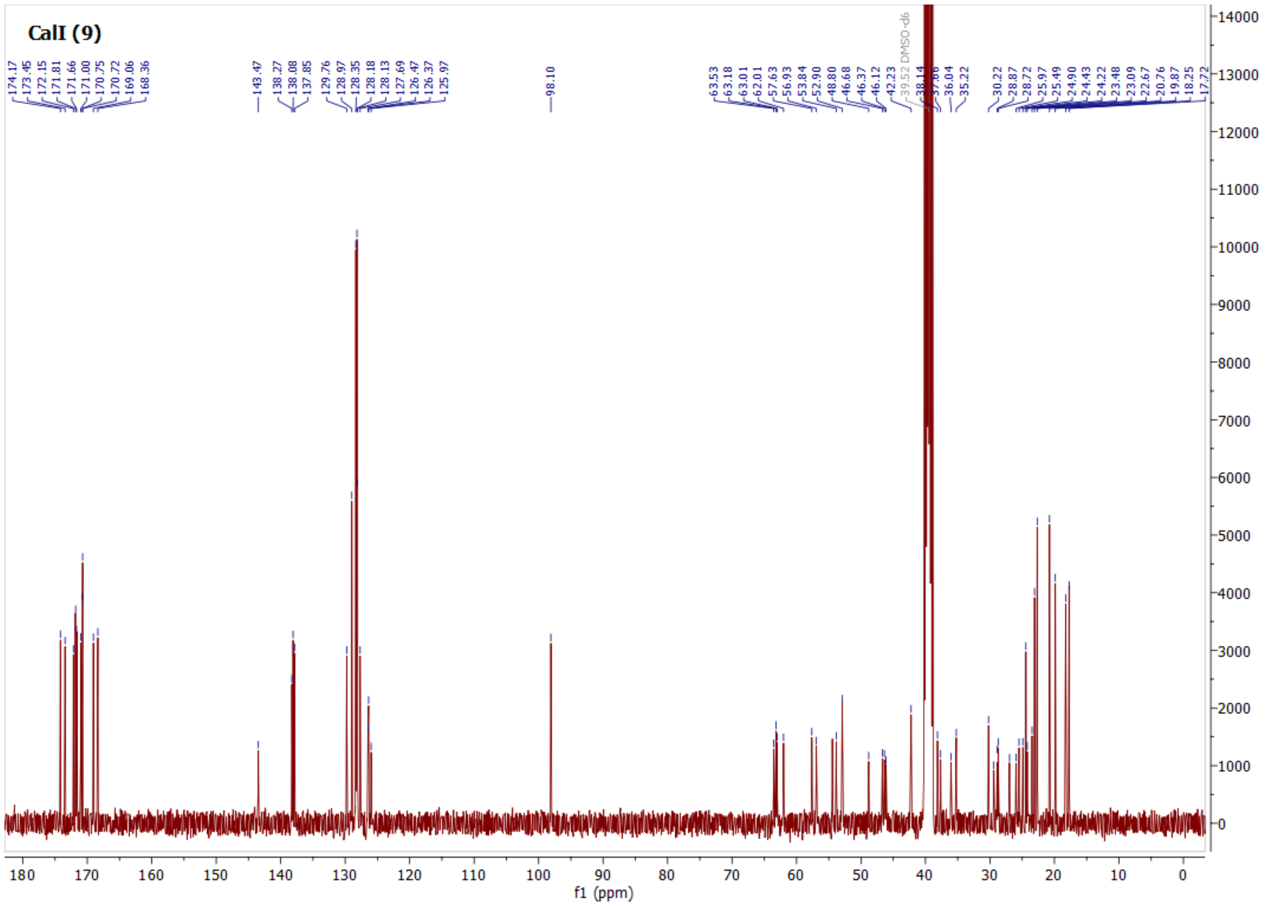
CalI (9)
f1 (ppm)

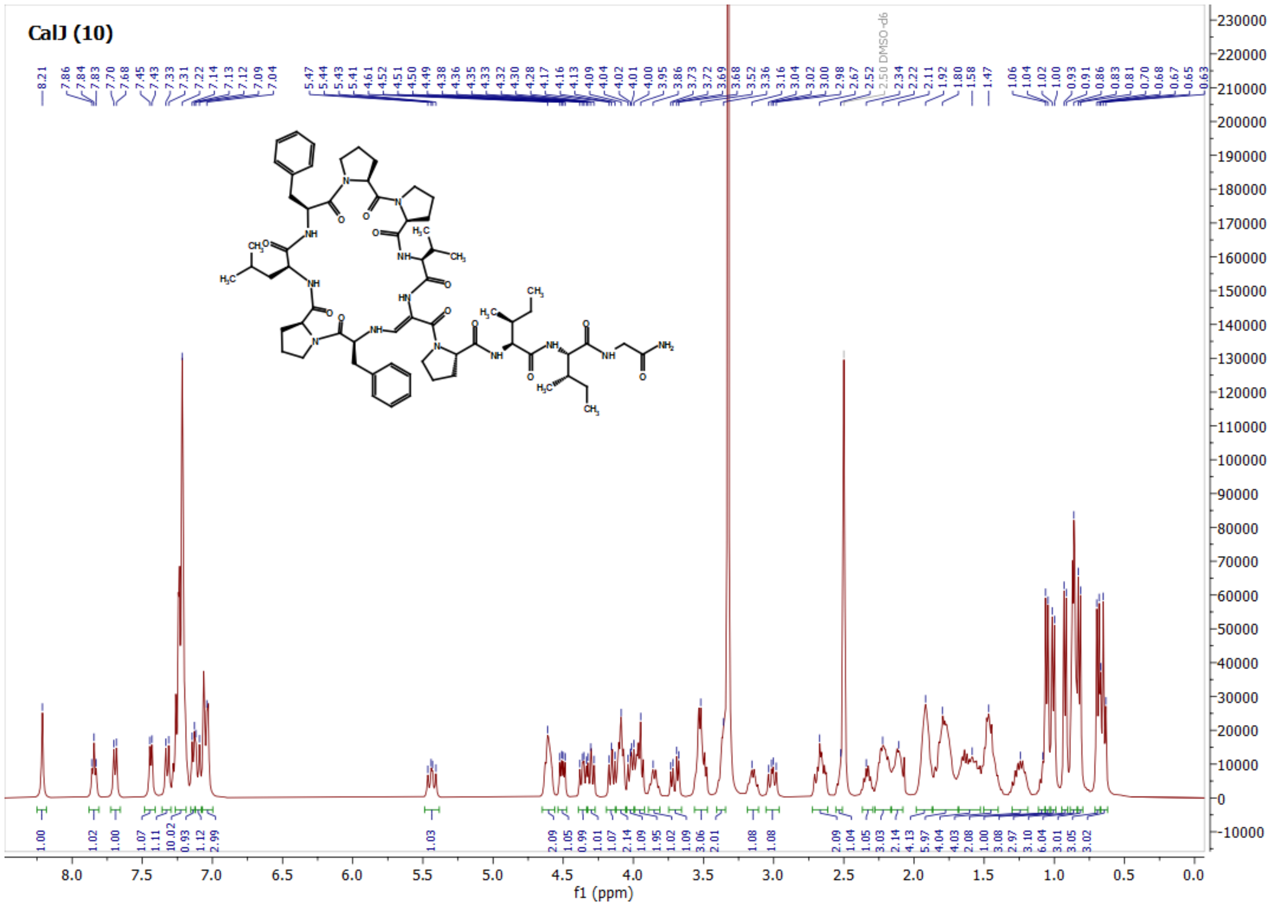
CalJ (10)
f1 (ppm)

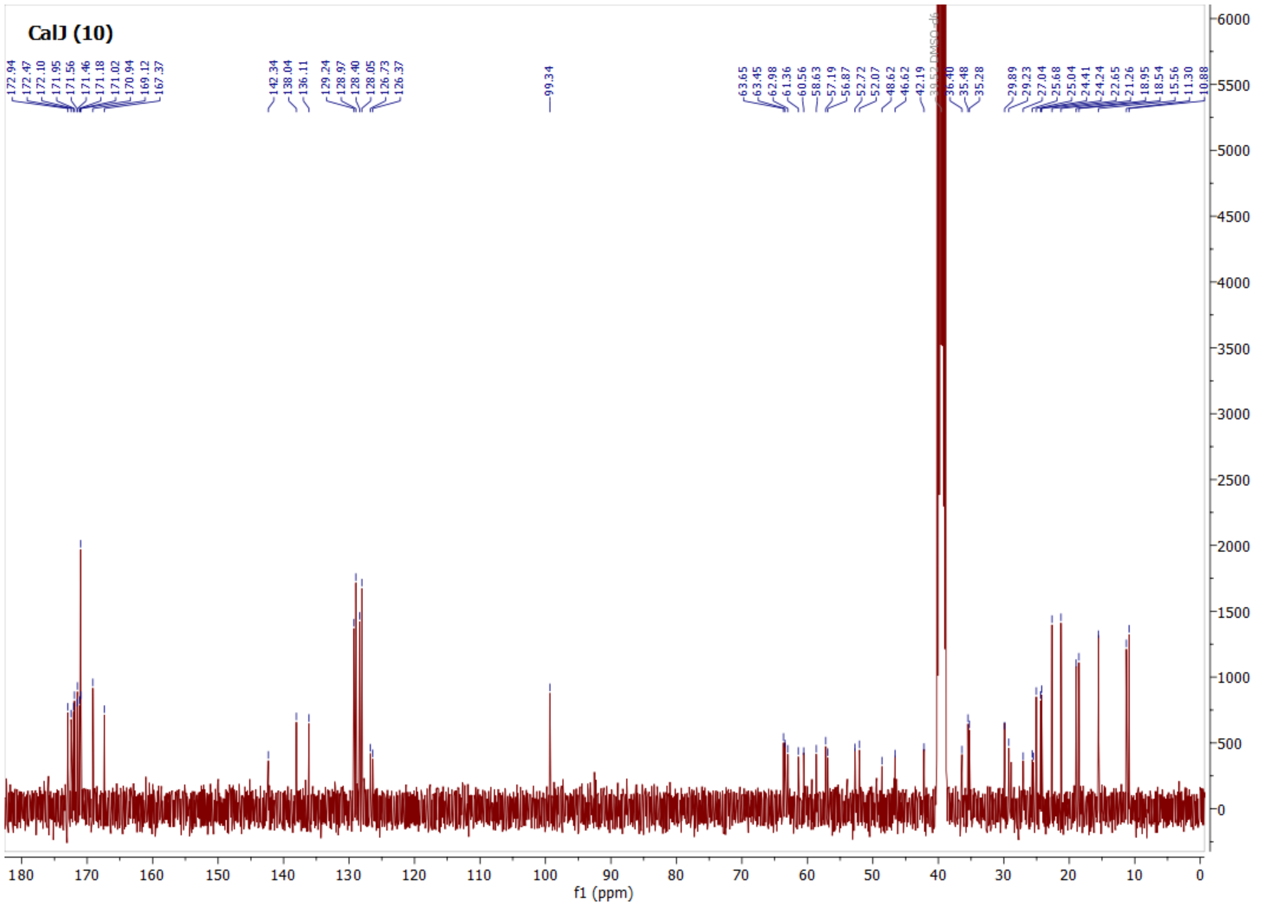
CalJ (10)
f1 (ppm)

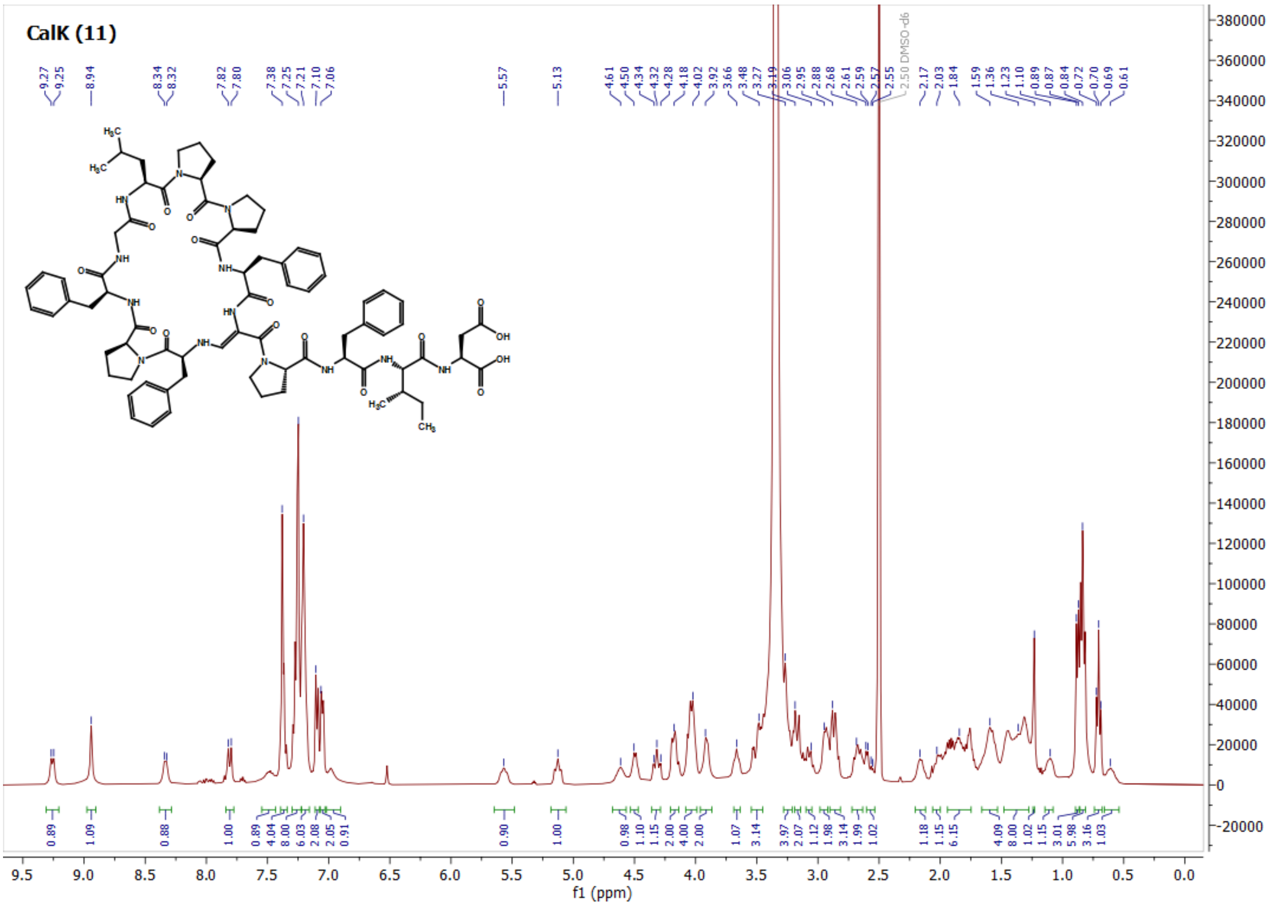
CalK (11)
f1 (ppm)

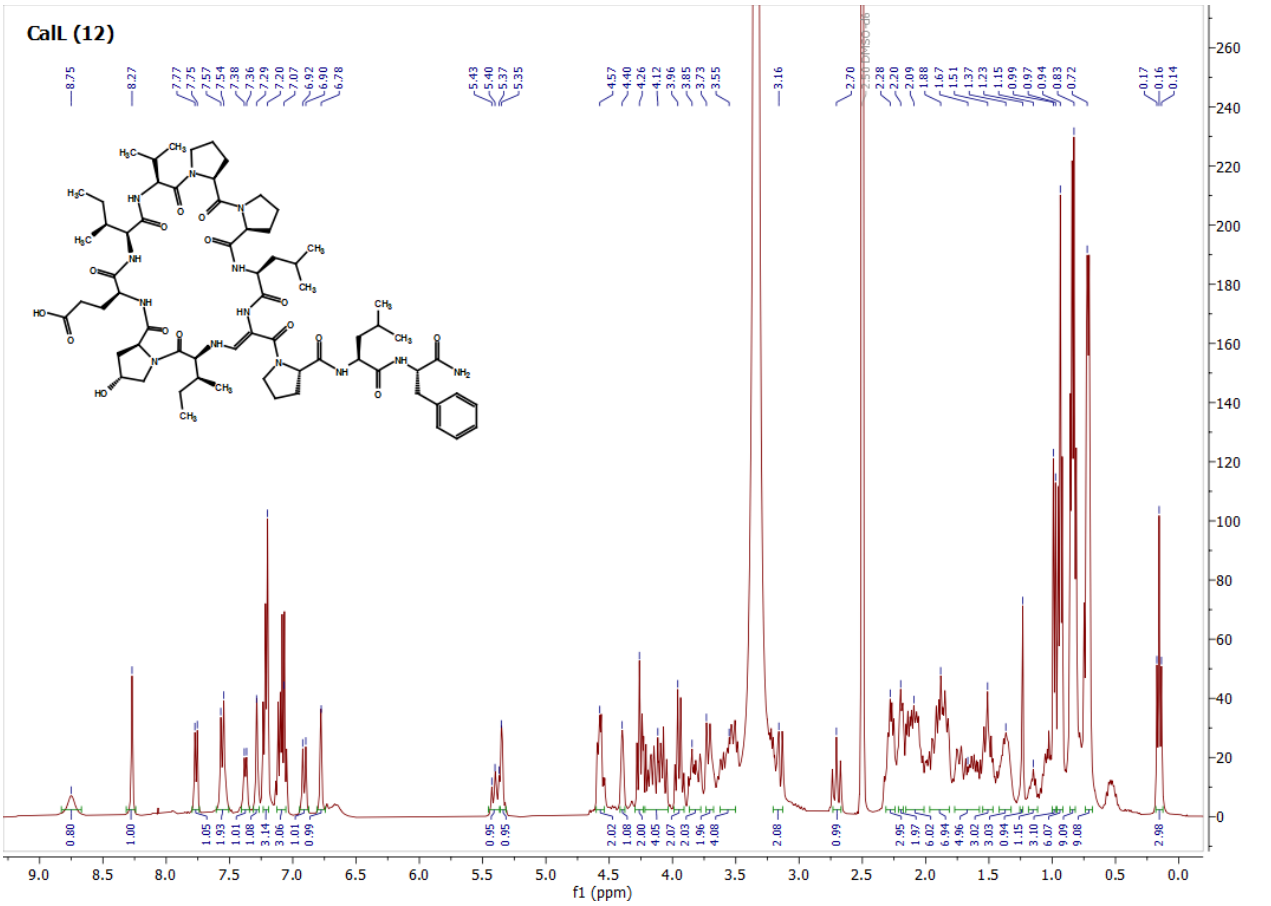
CalL (12)
f1 (ppm)

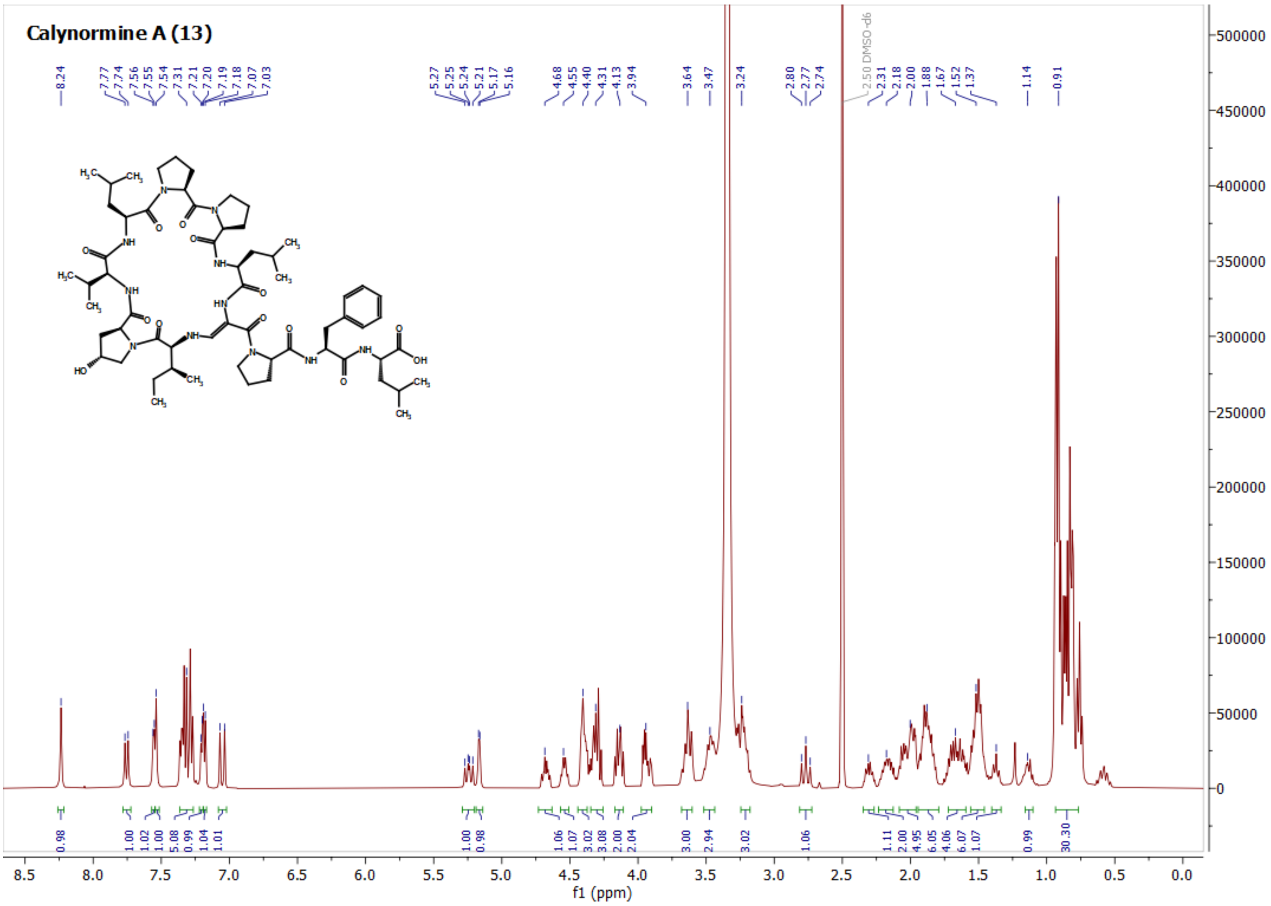
Calynormine A (13)
f1 (ppm)

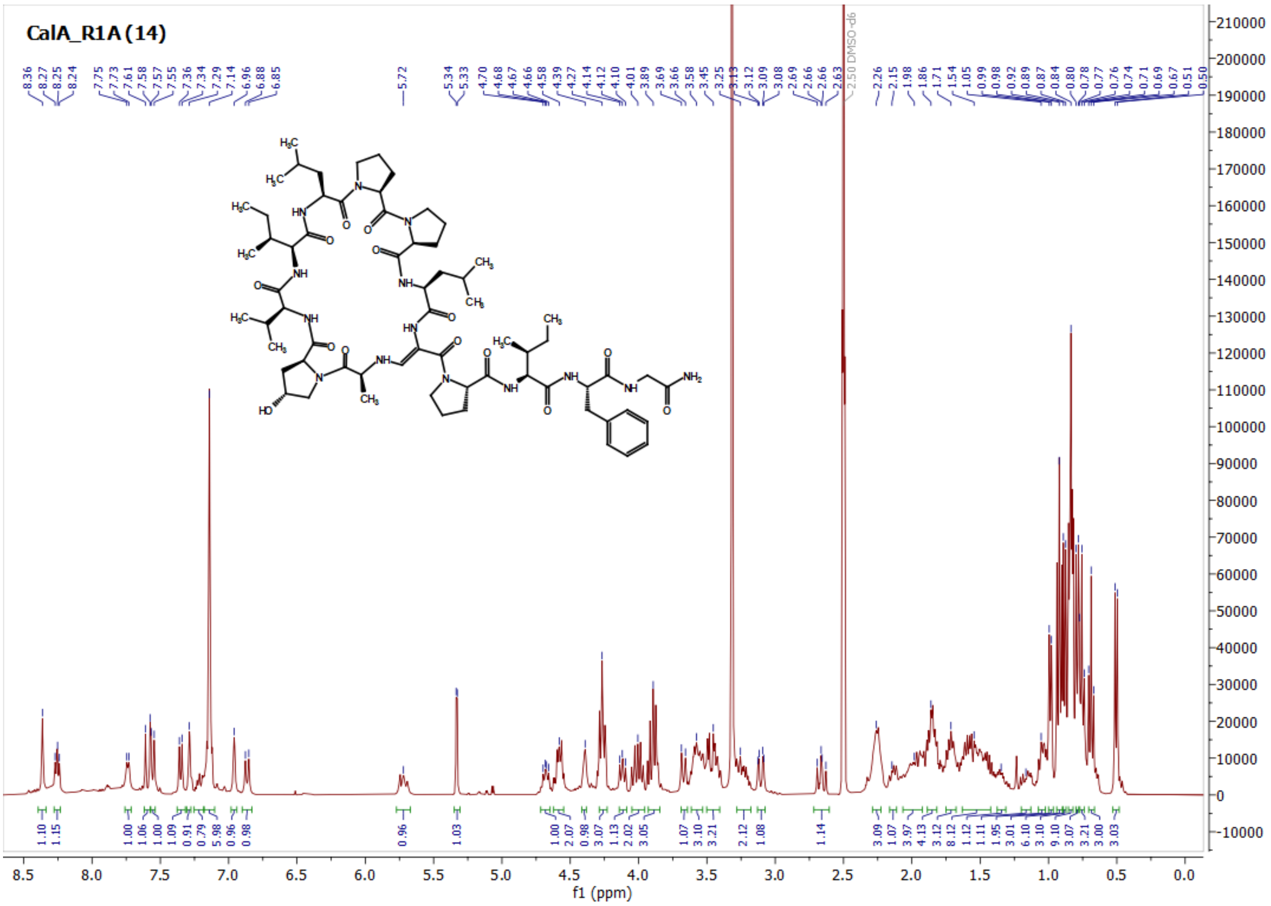
CalA_R1A (14)
f1 (ppm)

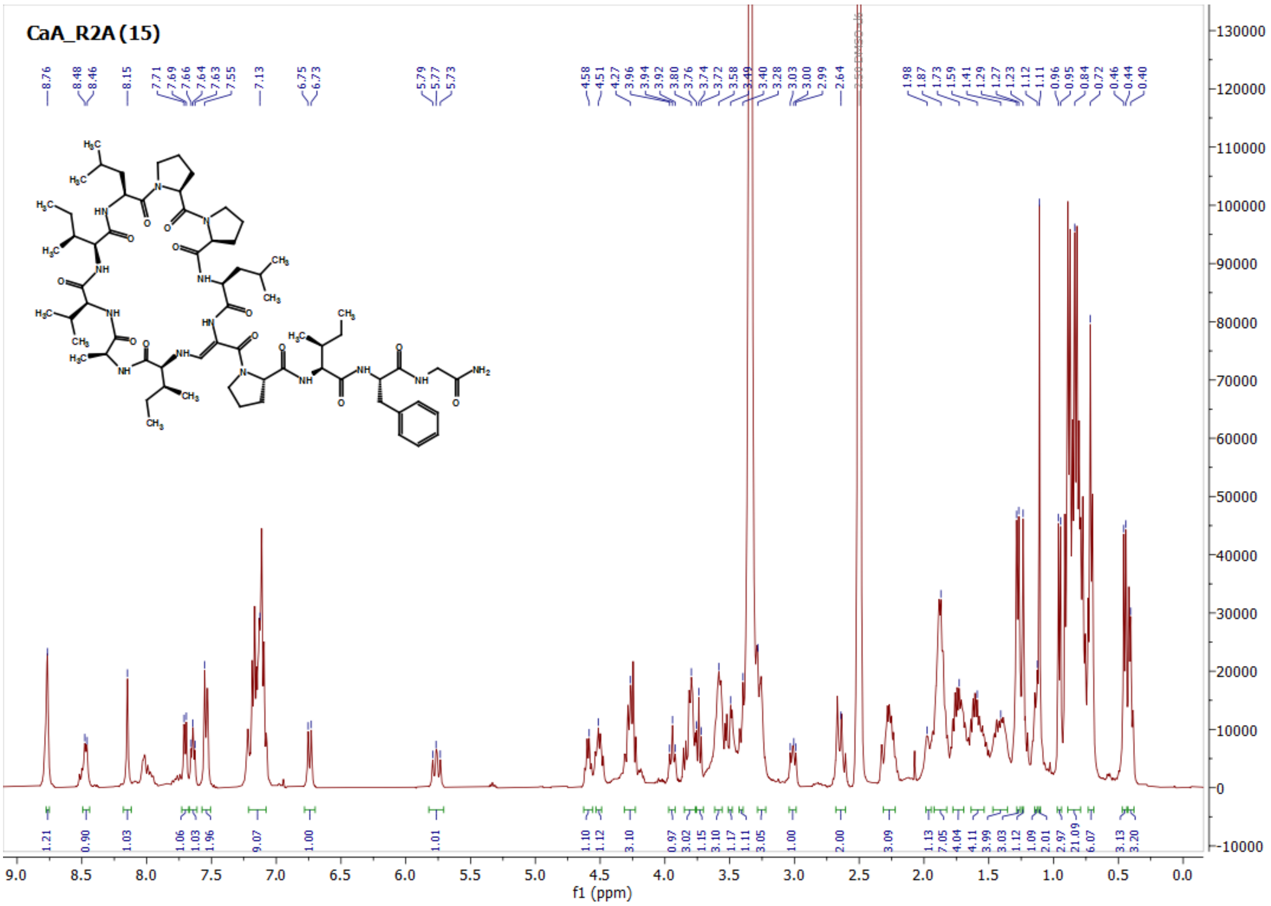
CaA_R2A (15)
f1 (ppm)

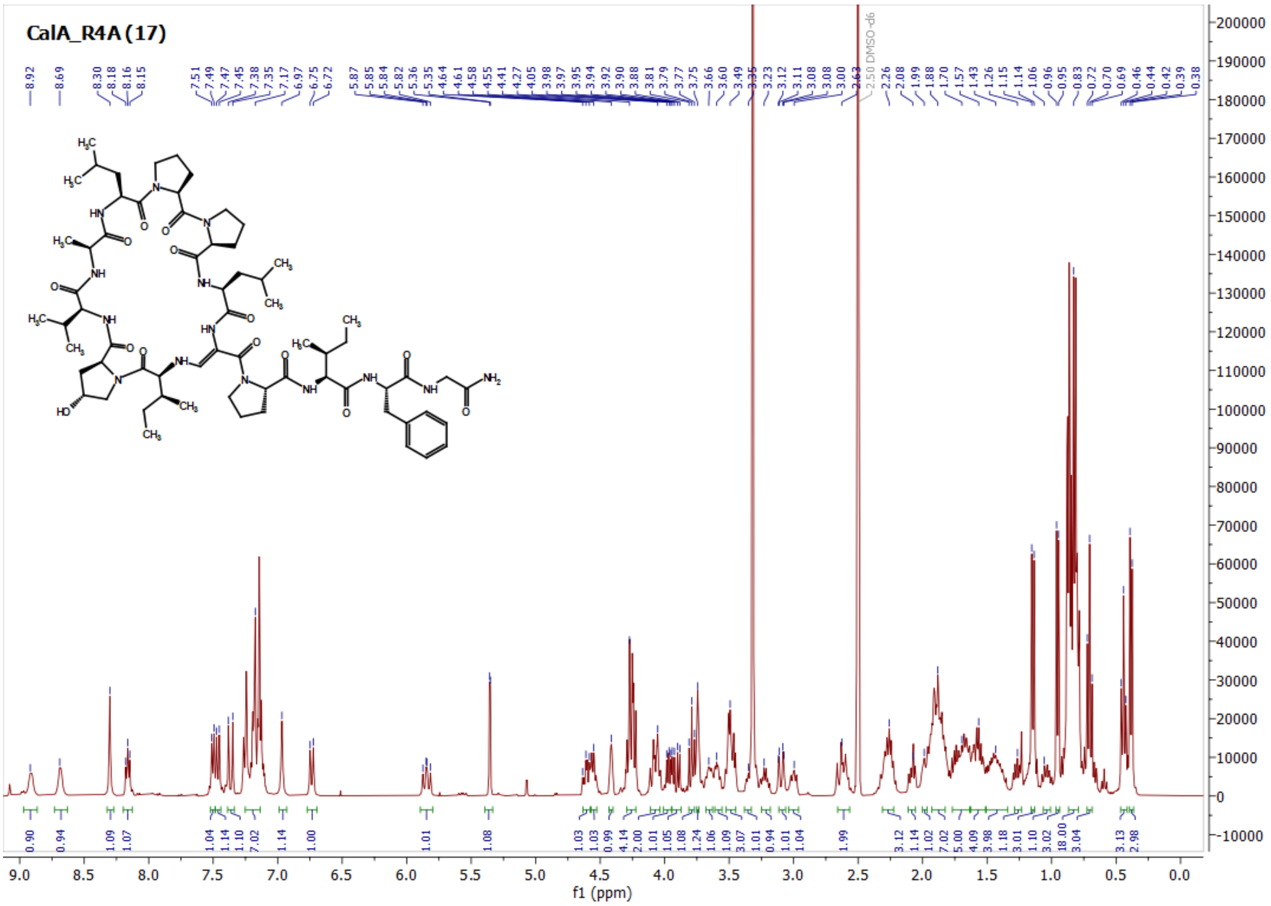
CalA_R4A (17)
f1 (ppm)

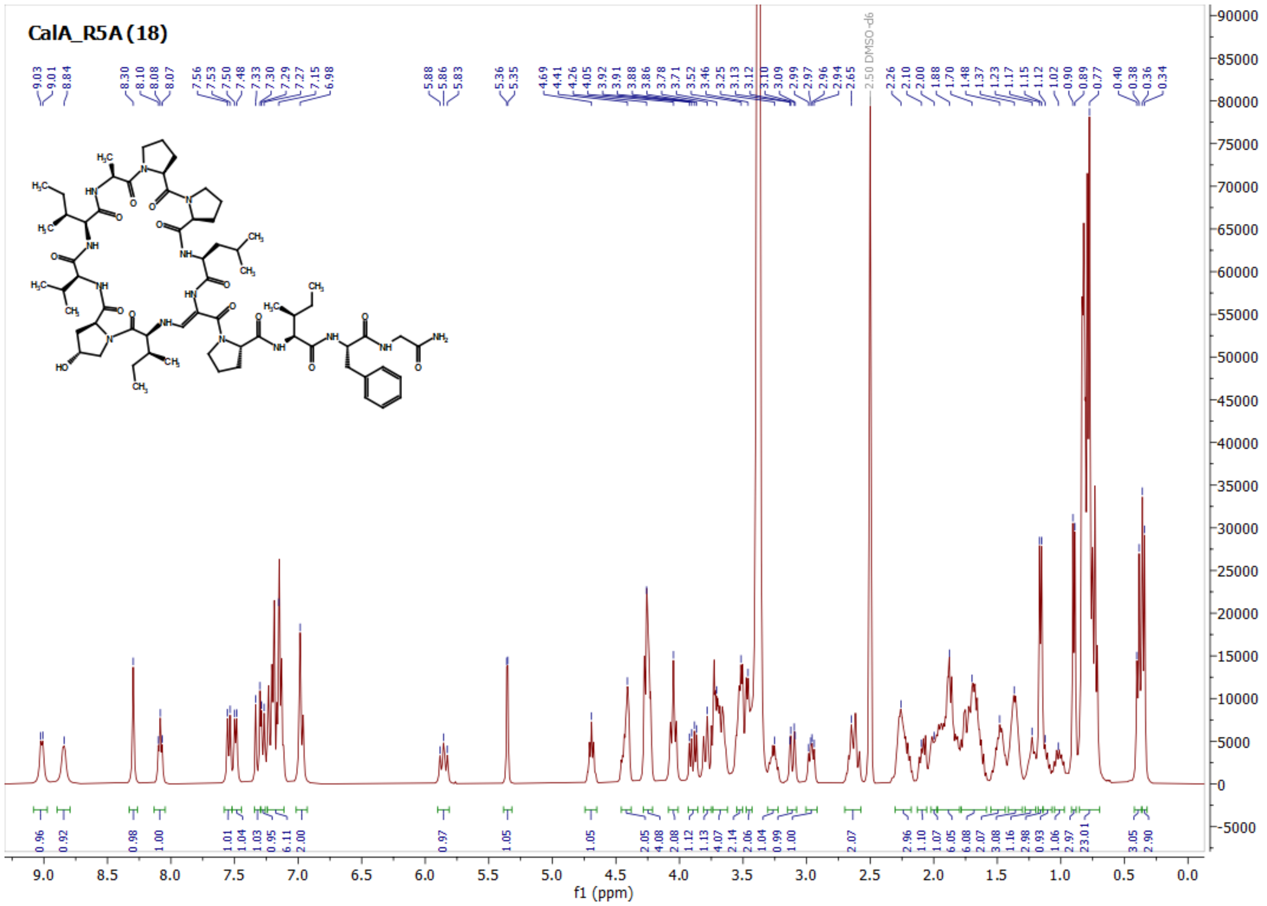
CalA_R5A (18)
f1 (ppm)

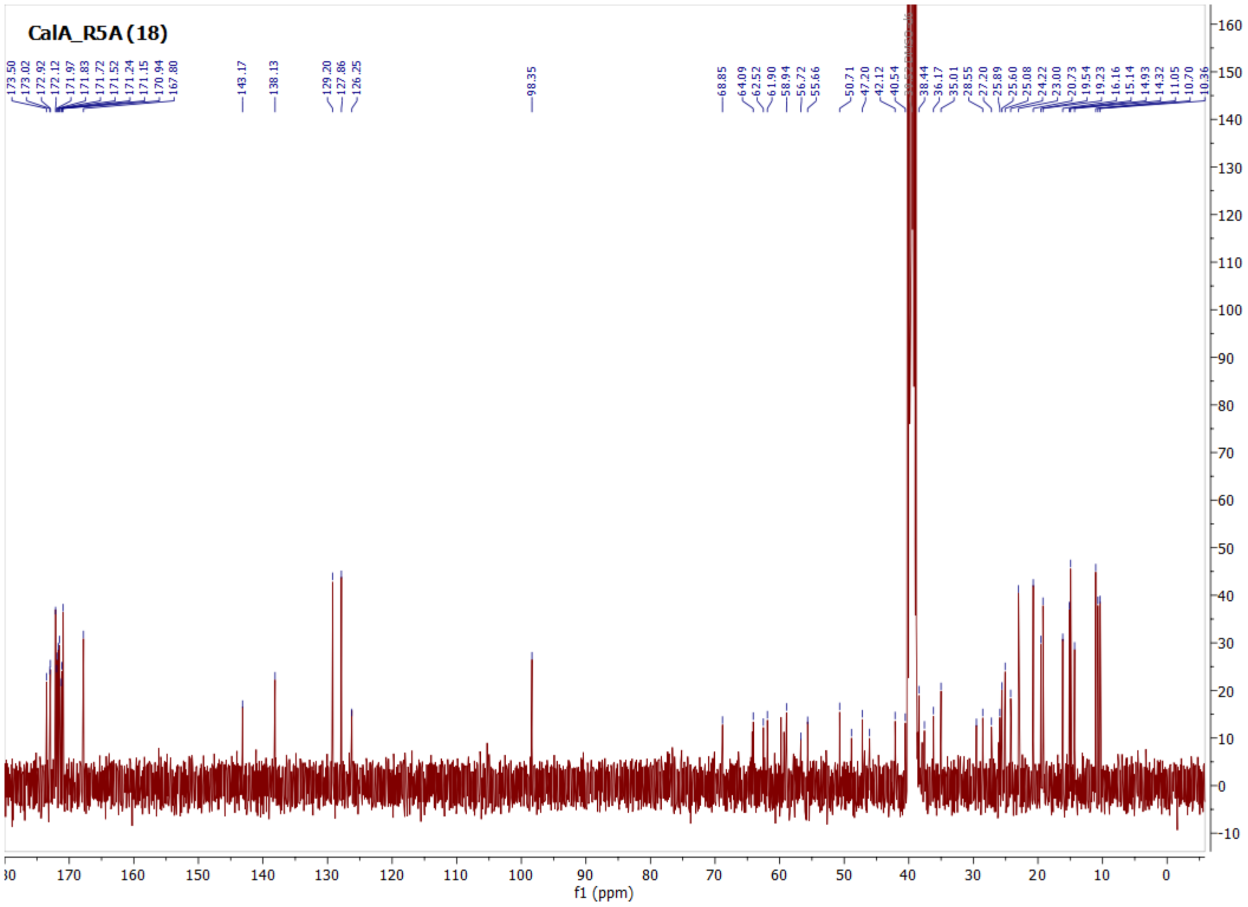
CalA_R5A (18)
f1 (ppm)

CaA_R8A (21)

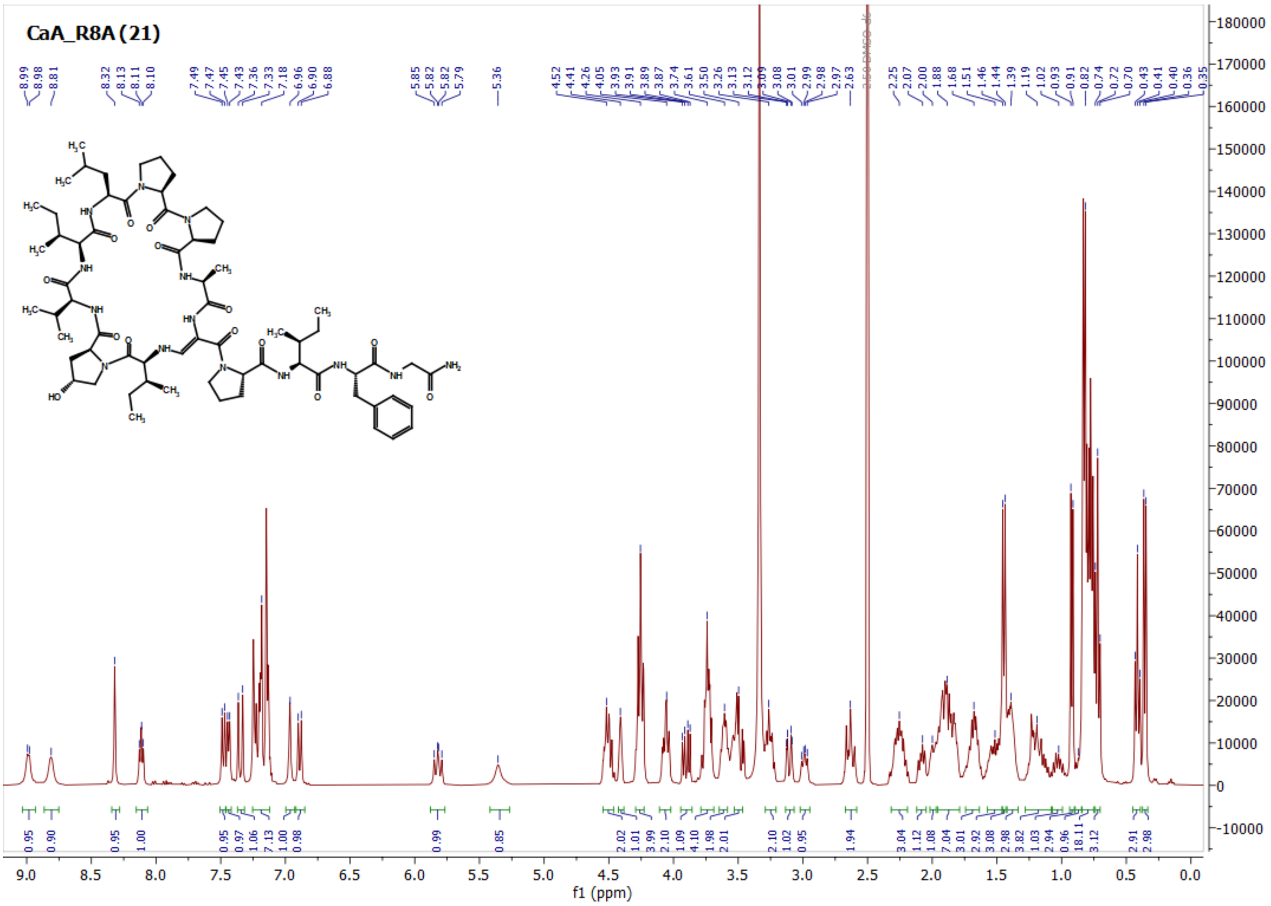

CalA_C2A (23)

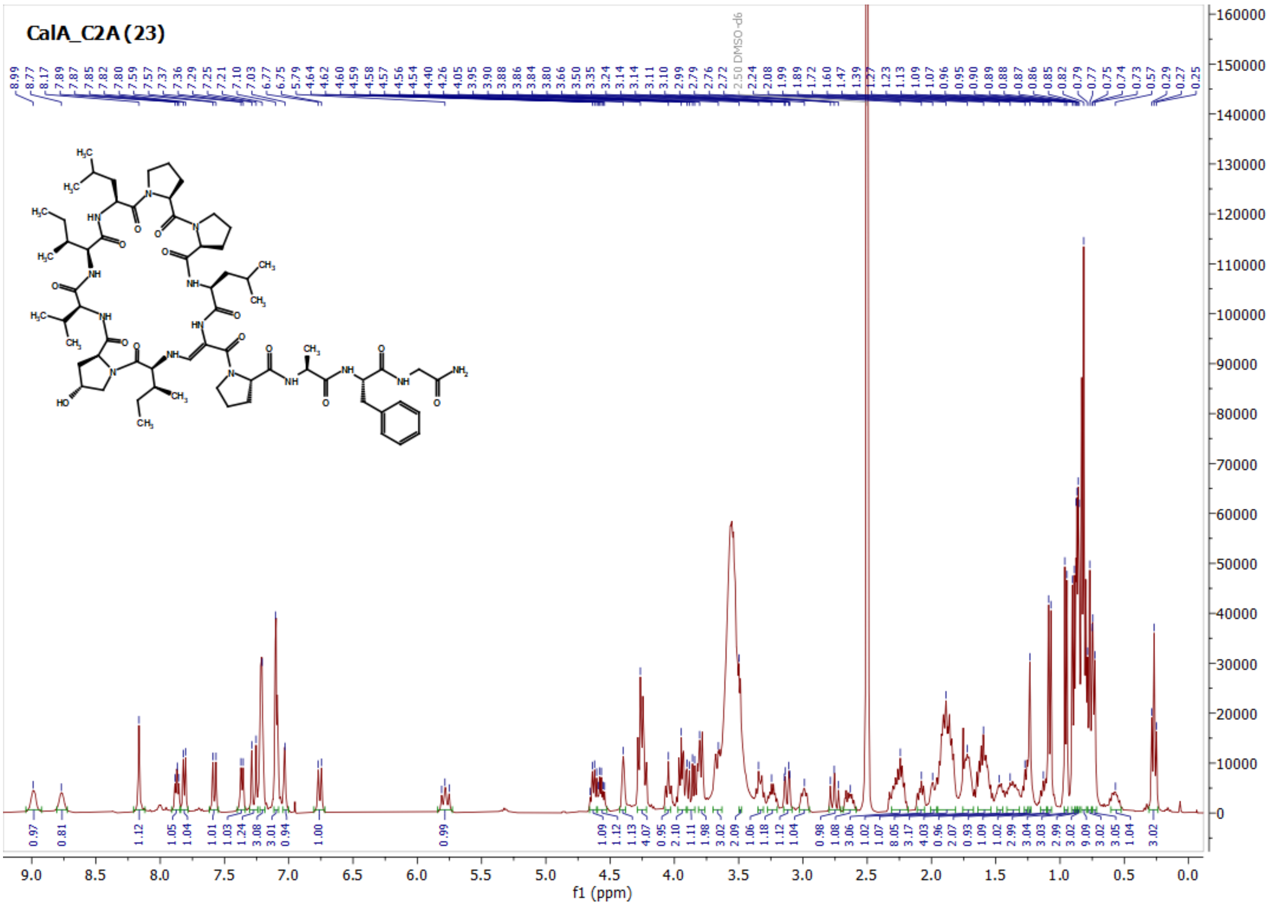

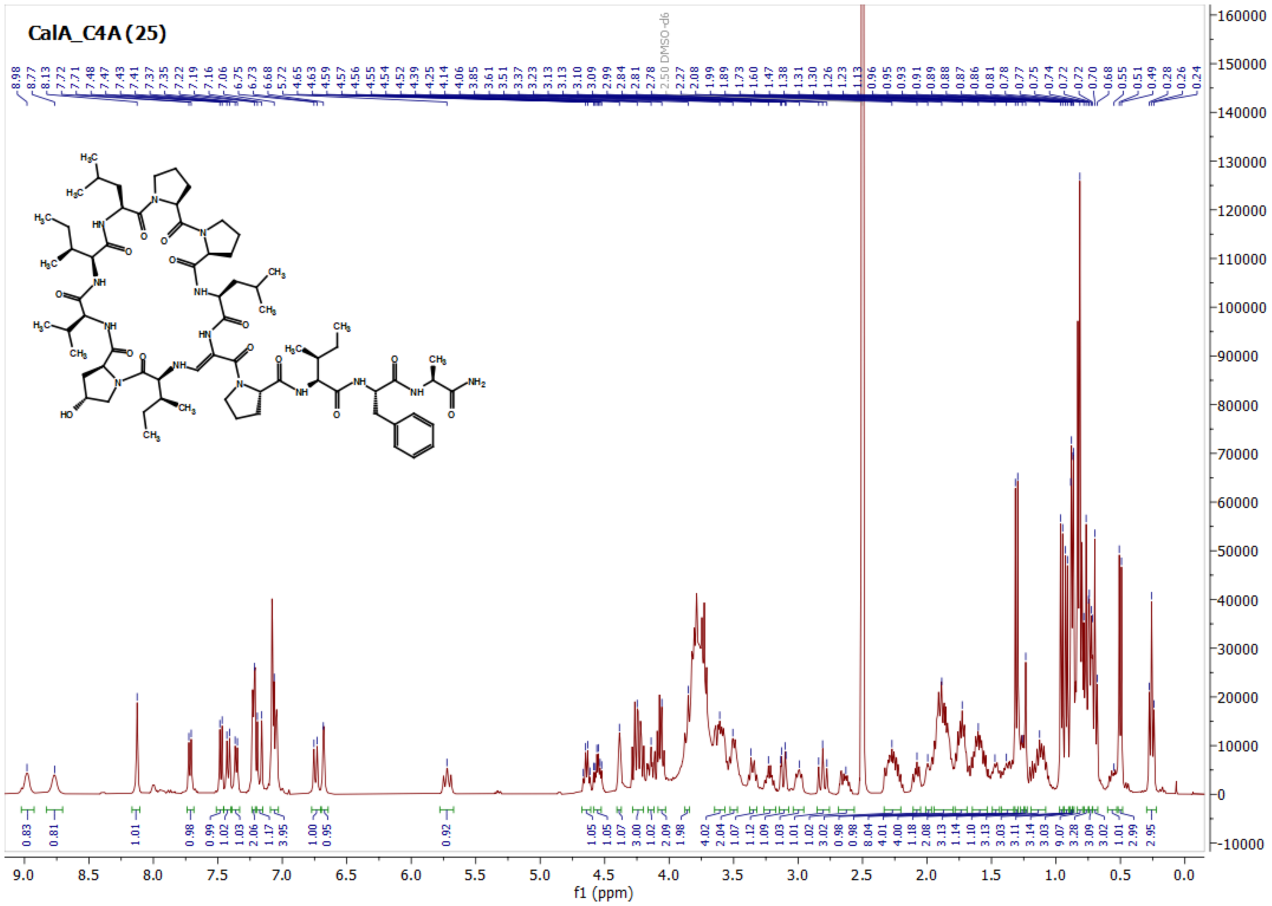
CalA_C4A (25)
f1 (ppm)

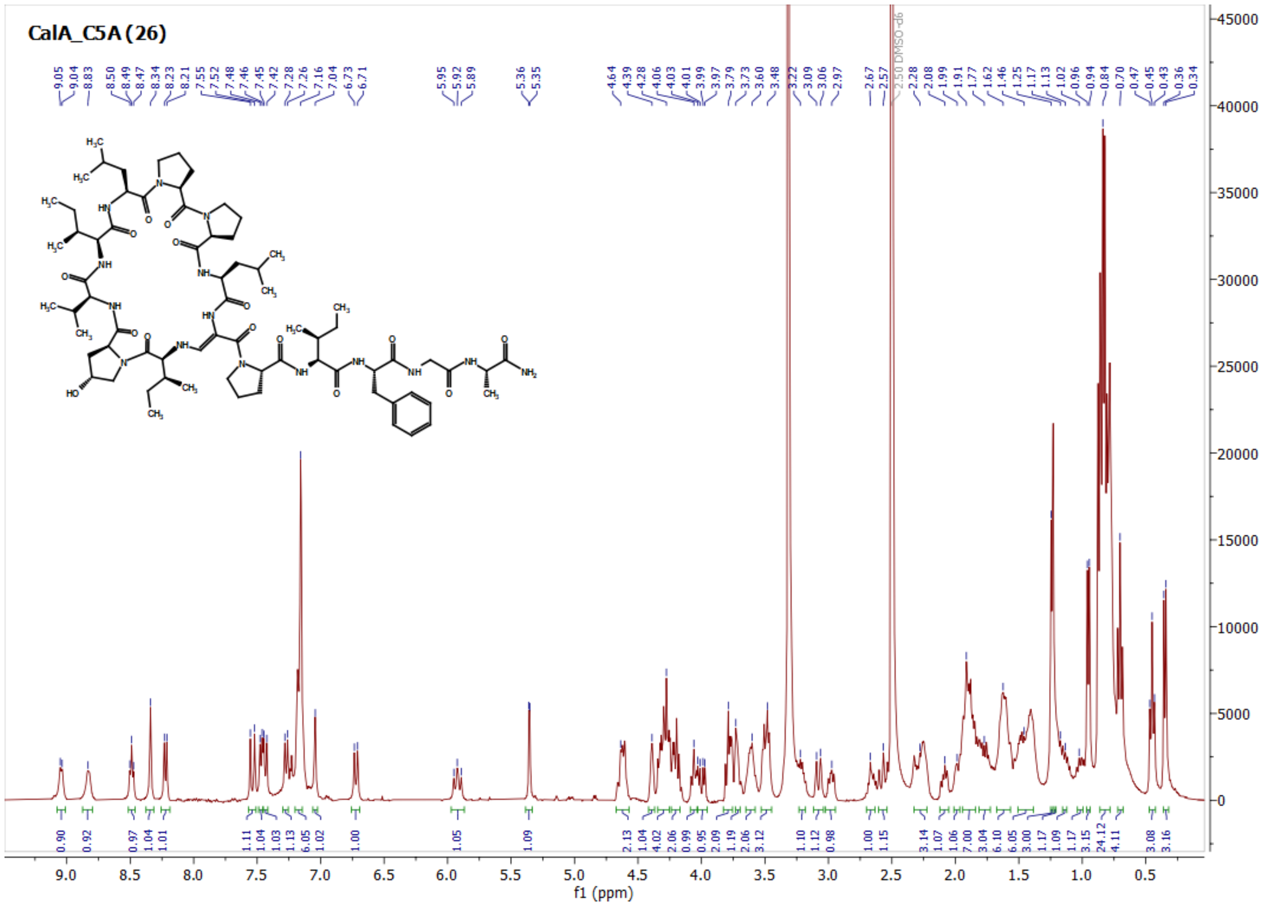
CalA_C5A (26)
f1 (ppm)

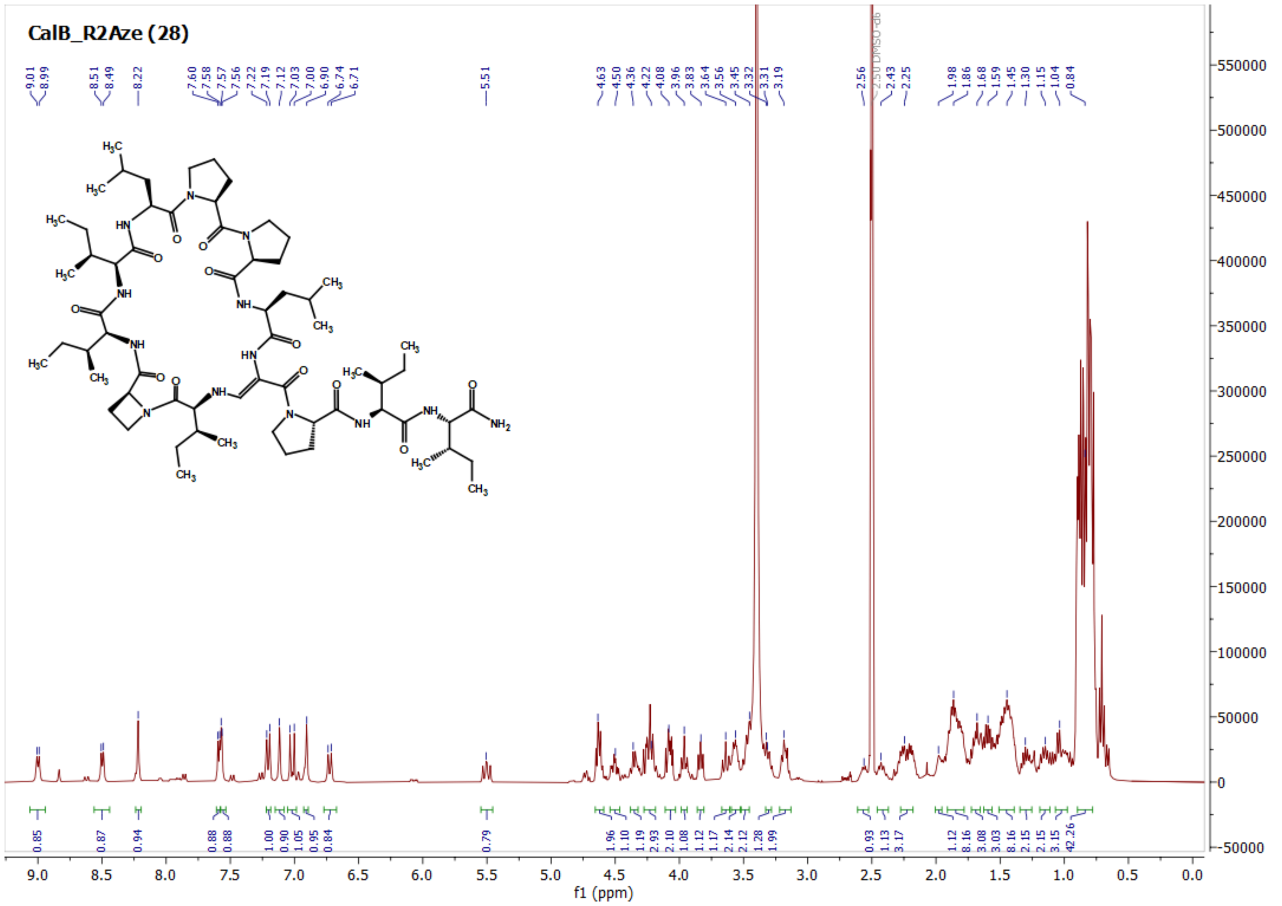

CalB_R2Aze (28)

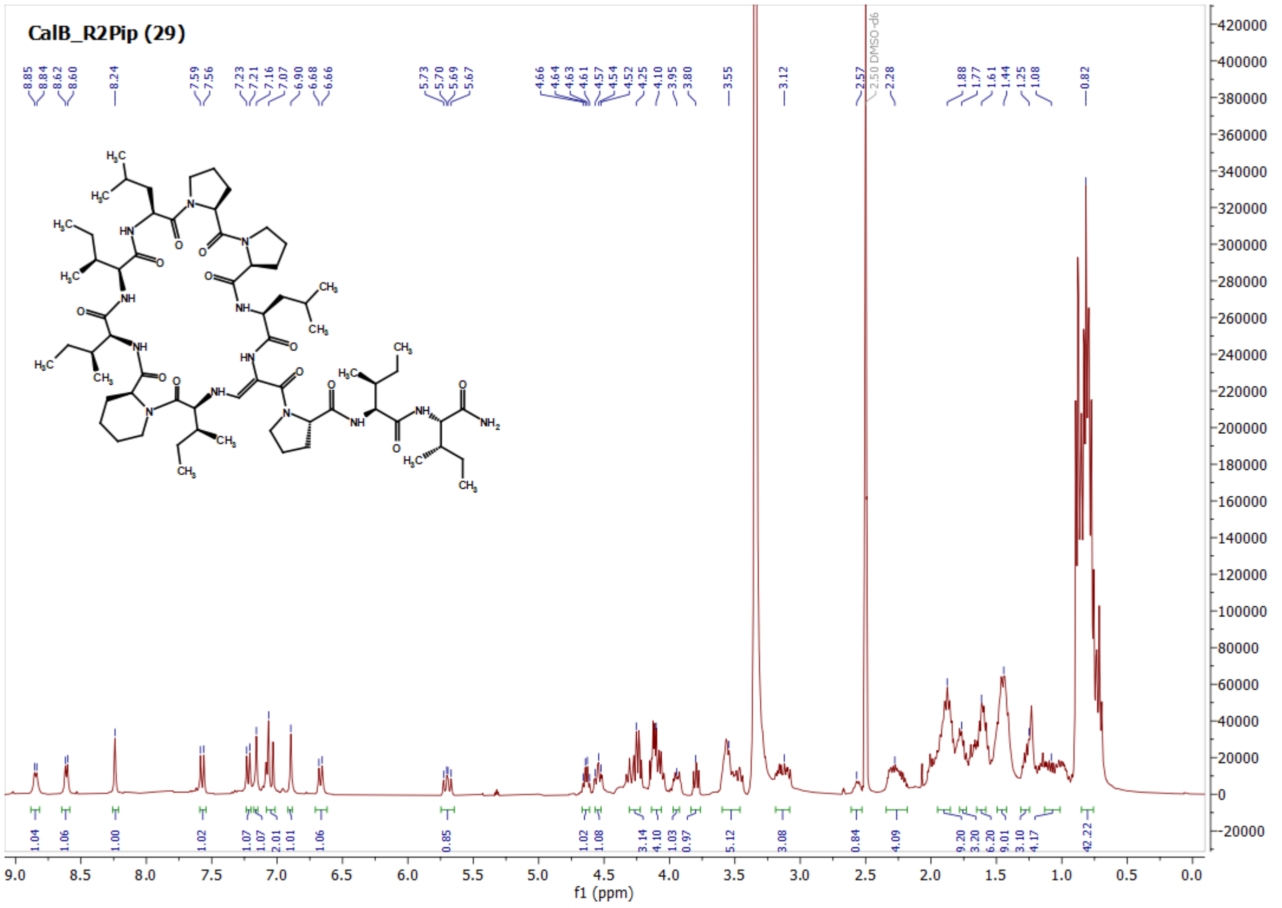

CalB_R2Pip (29)

CalB_R2Oic (31)

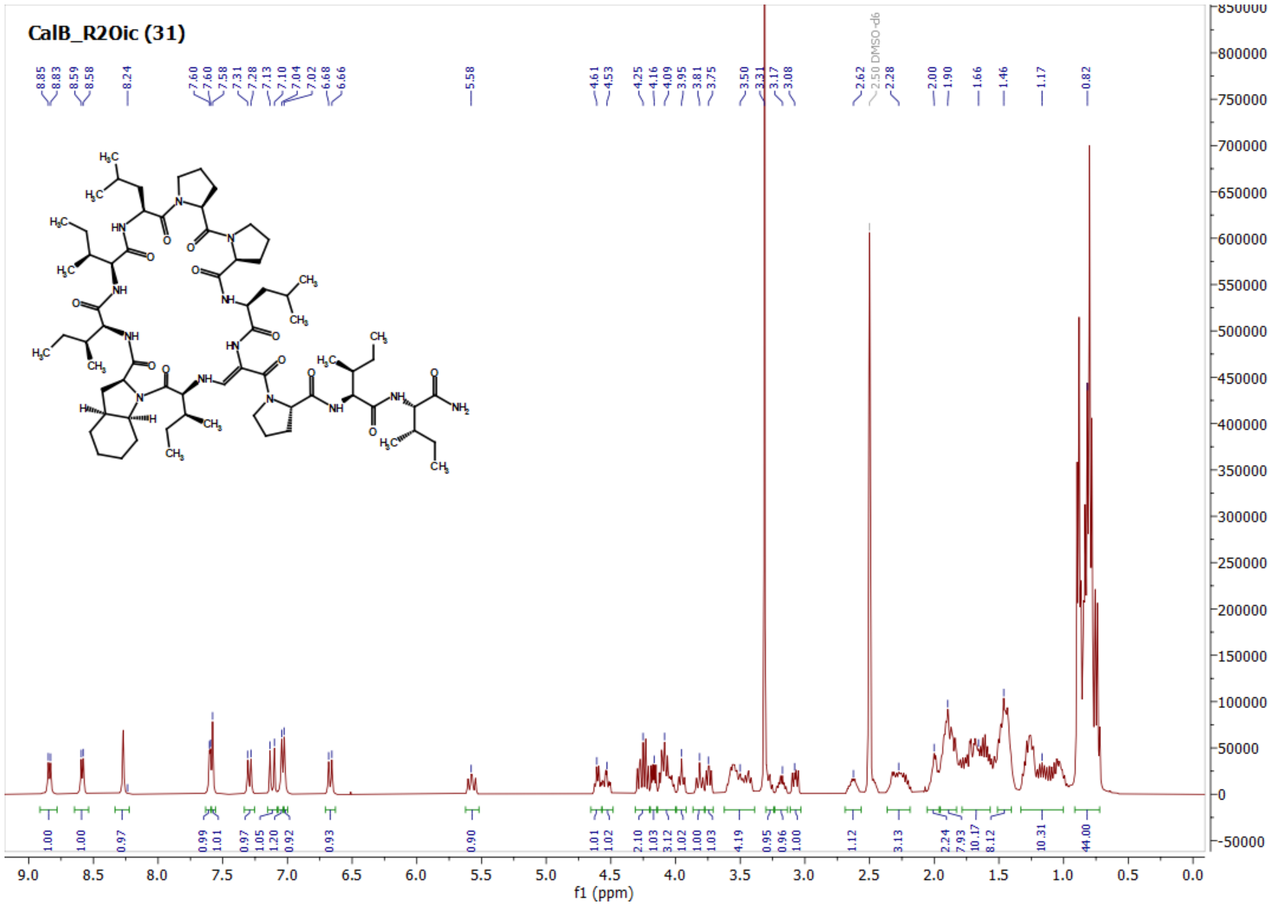

CalB_R3hLeu (32)

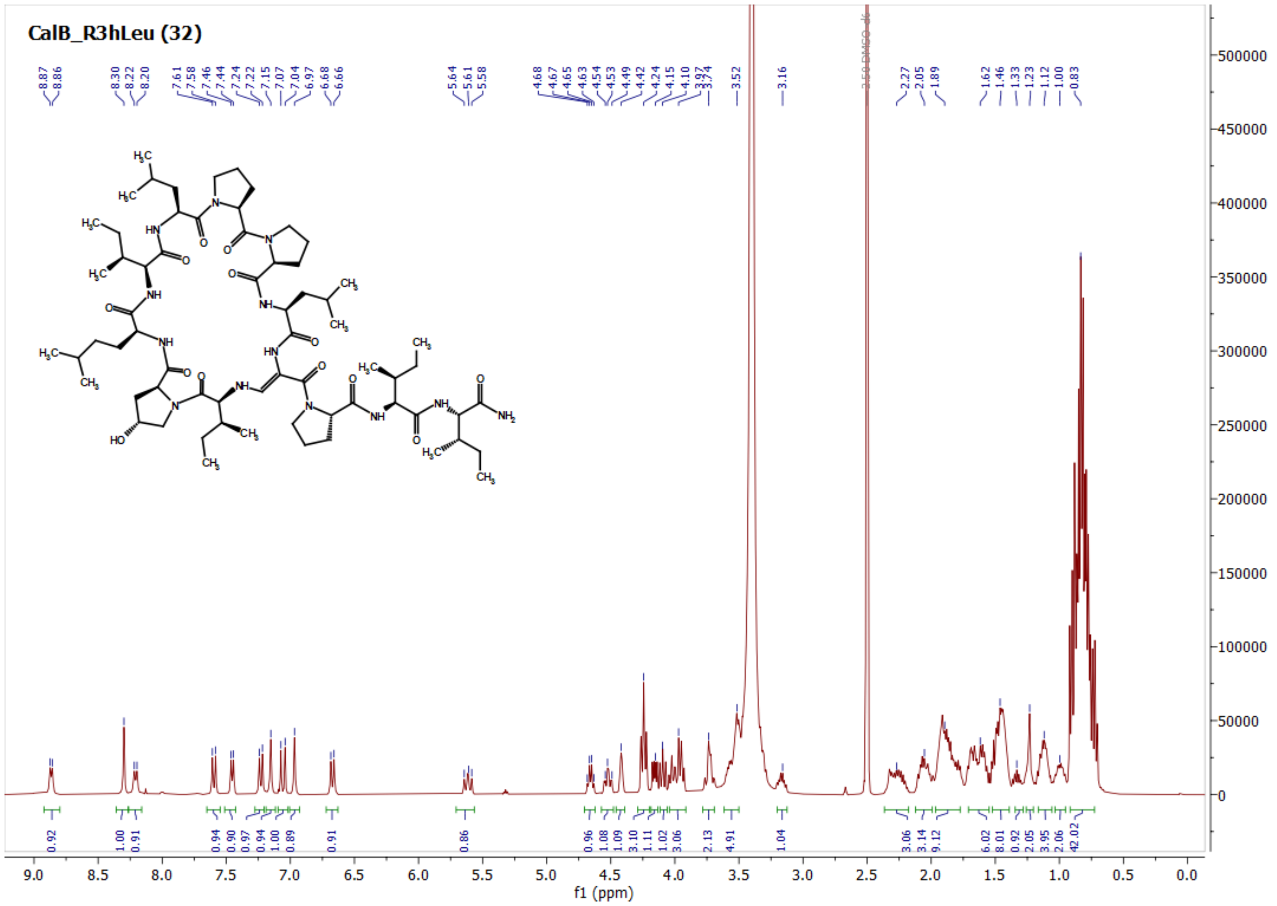

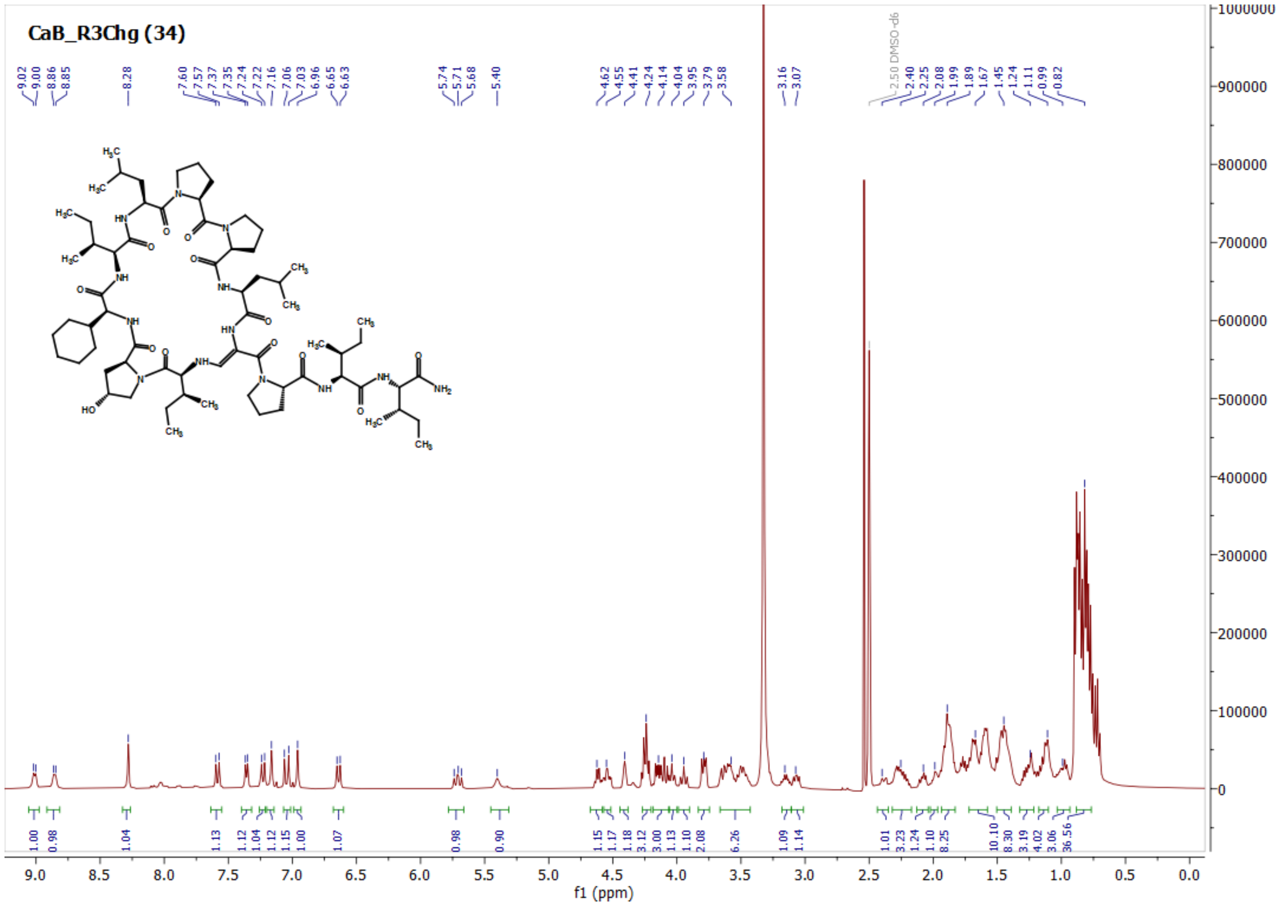
CaB_R3Chg (34)
f1 (ppm)

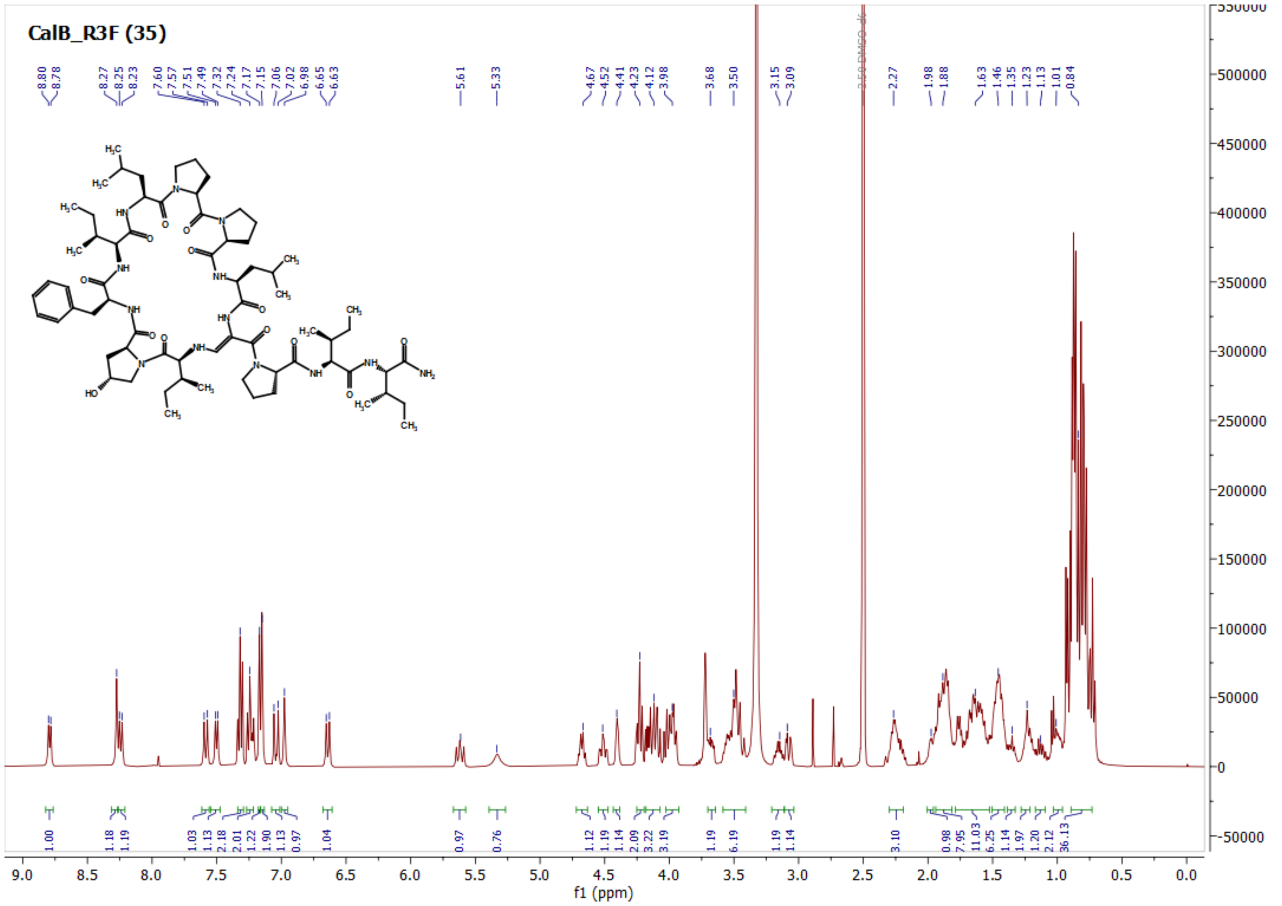
CalB_R3F (35)
f1 (ppm)

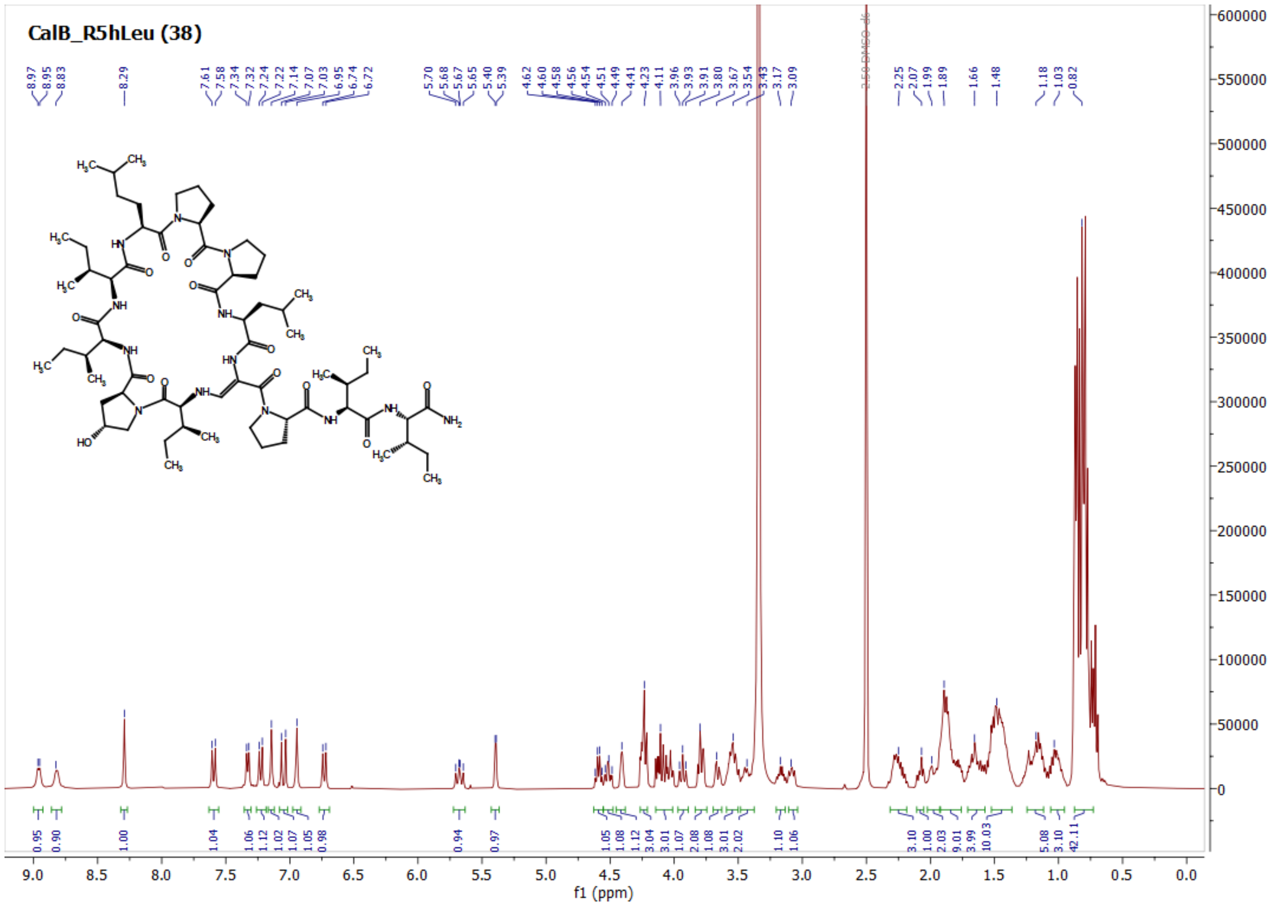
CalB_R5hLeu (38)
f1 (ppm)

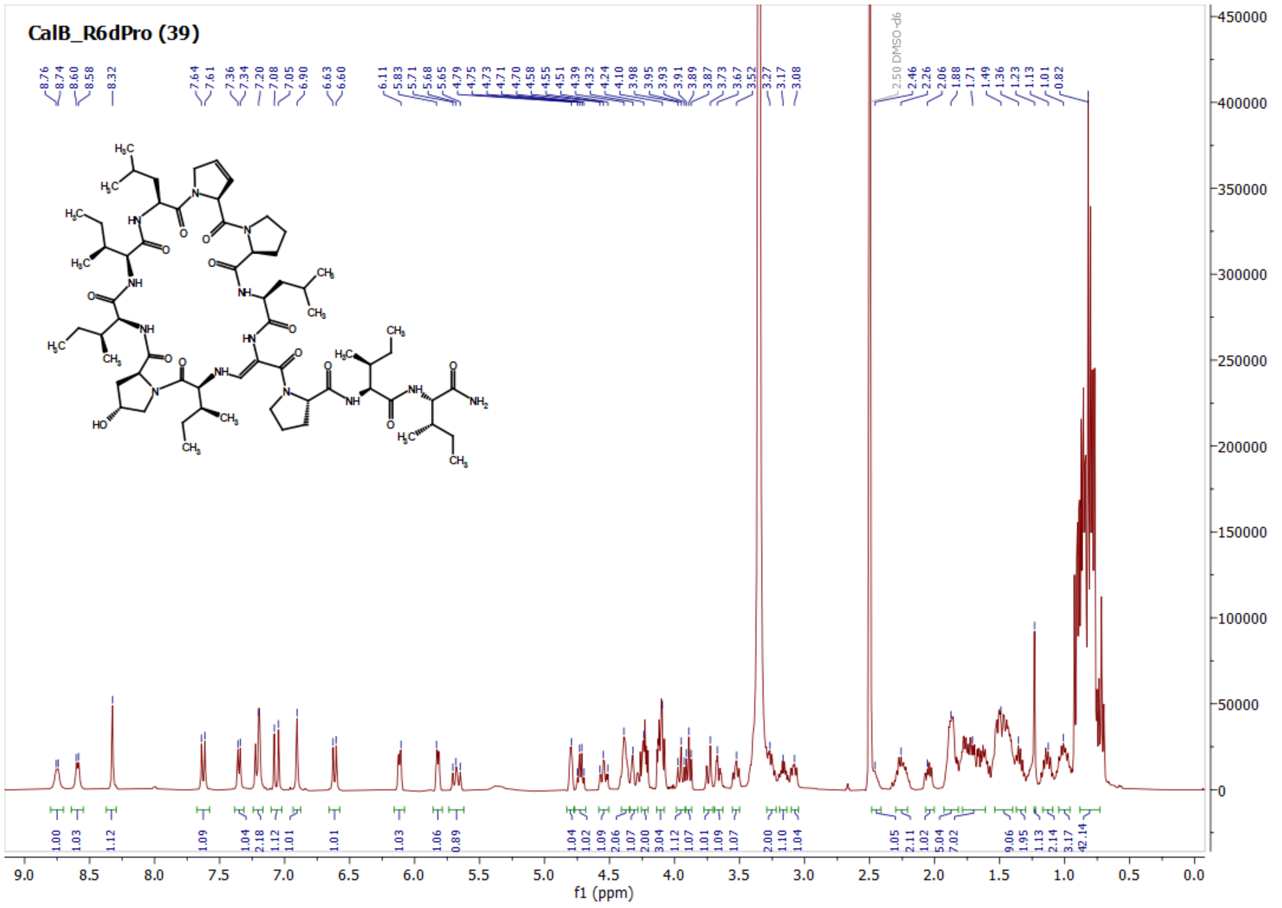
CalB_R6dPro (39)
f1 (ppm)

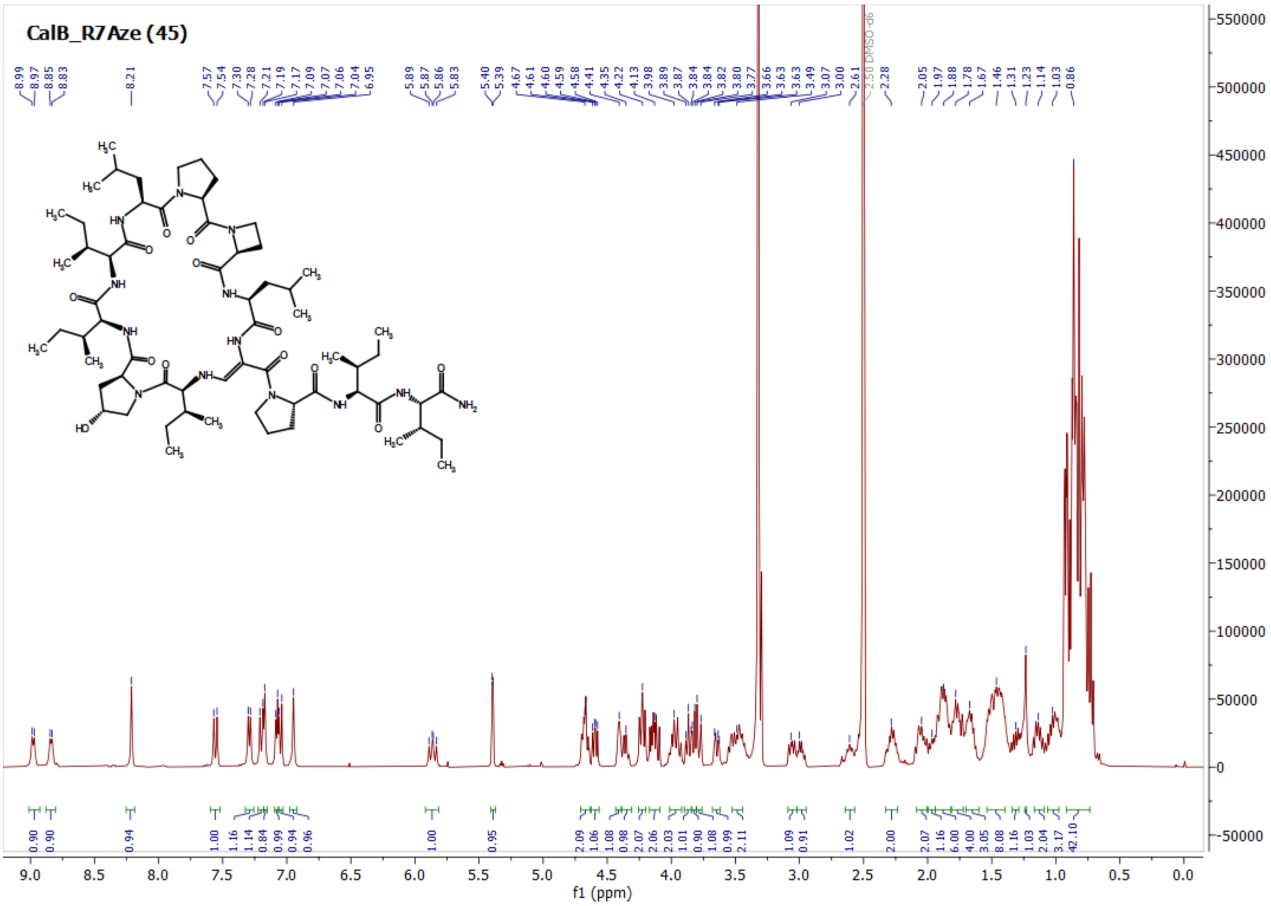
CalB_R7Aze (45)
f1 (ppm)

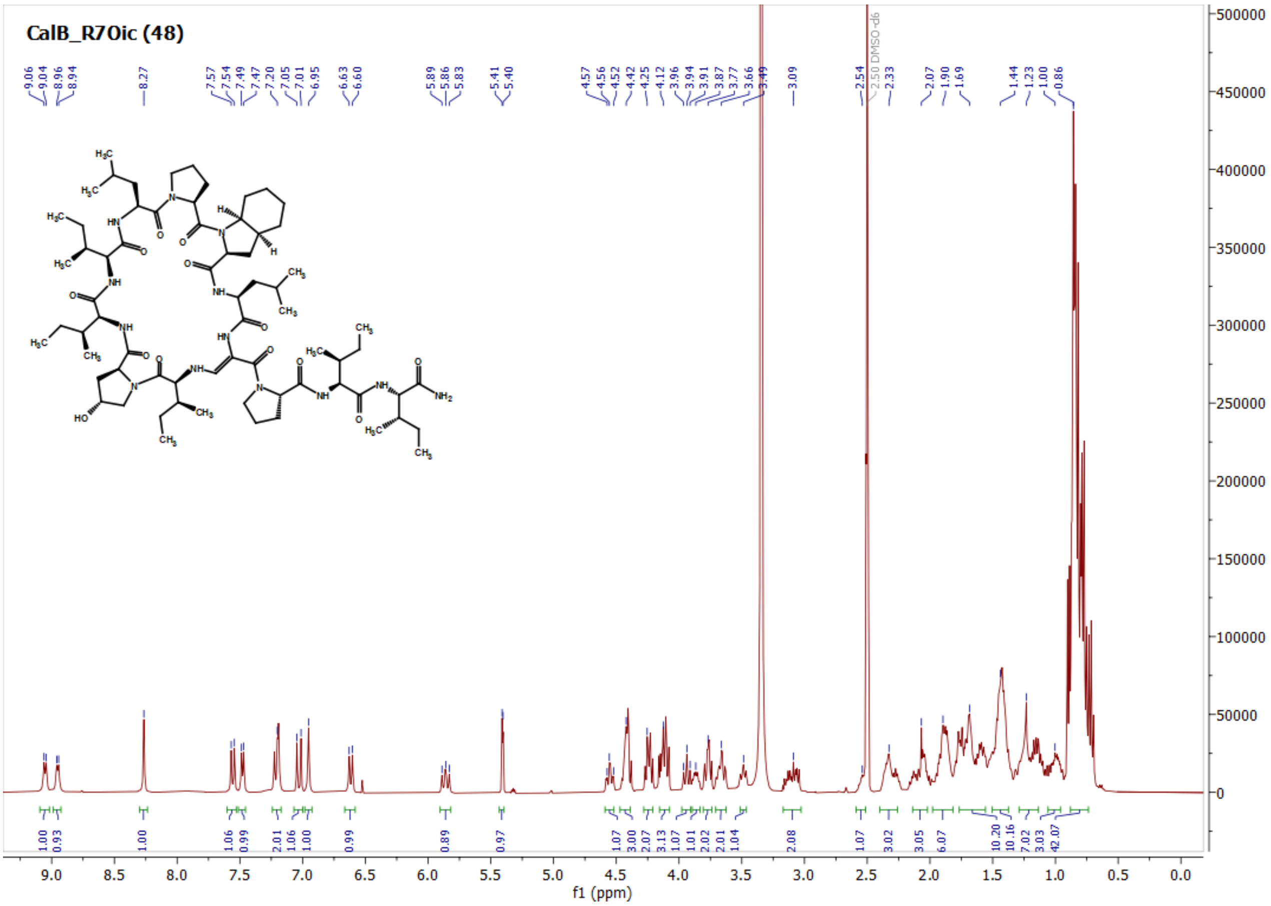
CalB_R7Oic (48)
f1 (ppm)

CalB_R8hLeu (49)

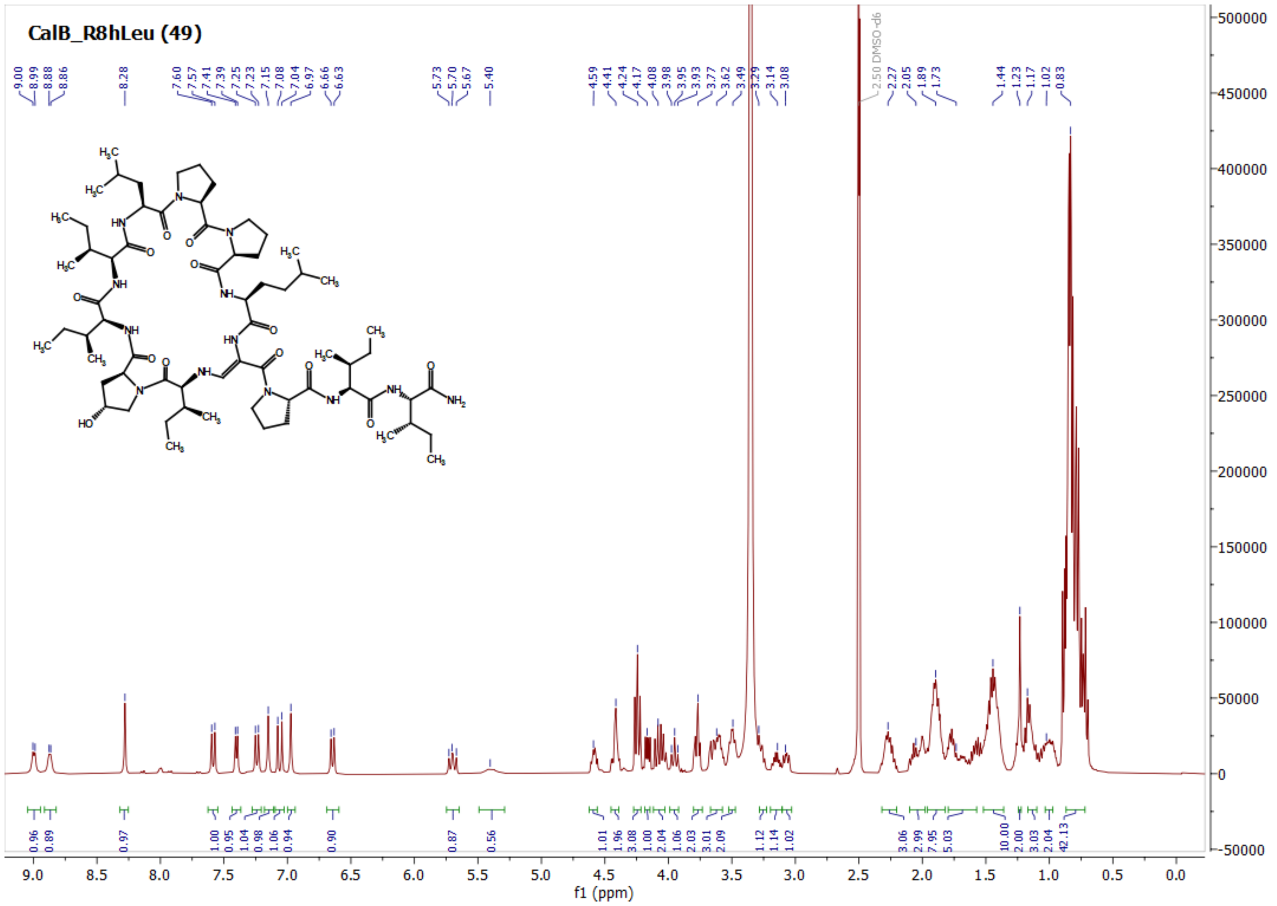

CalB_C1Aze (52)

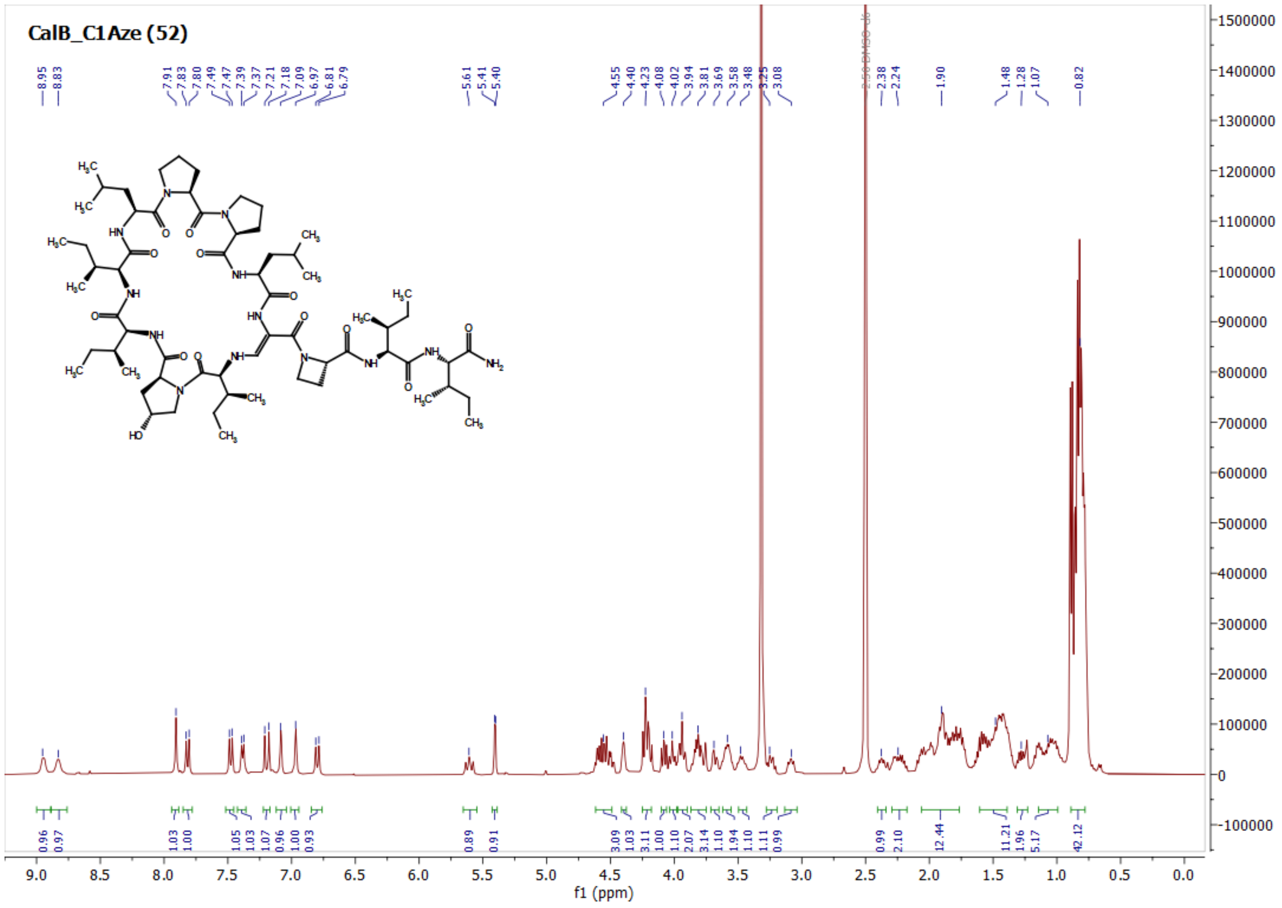

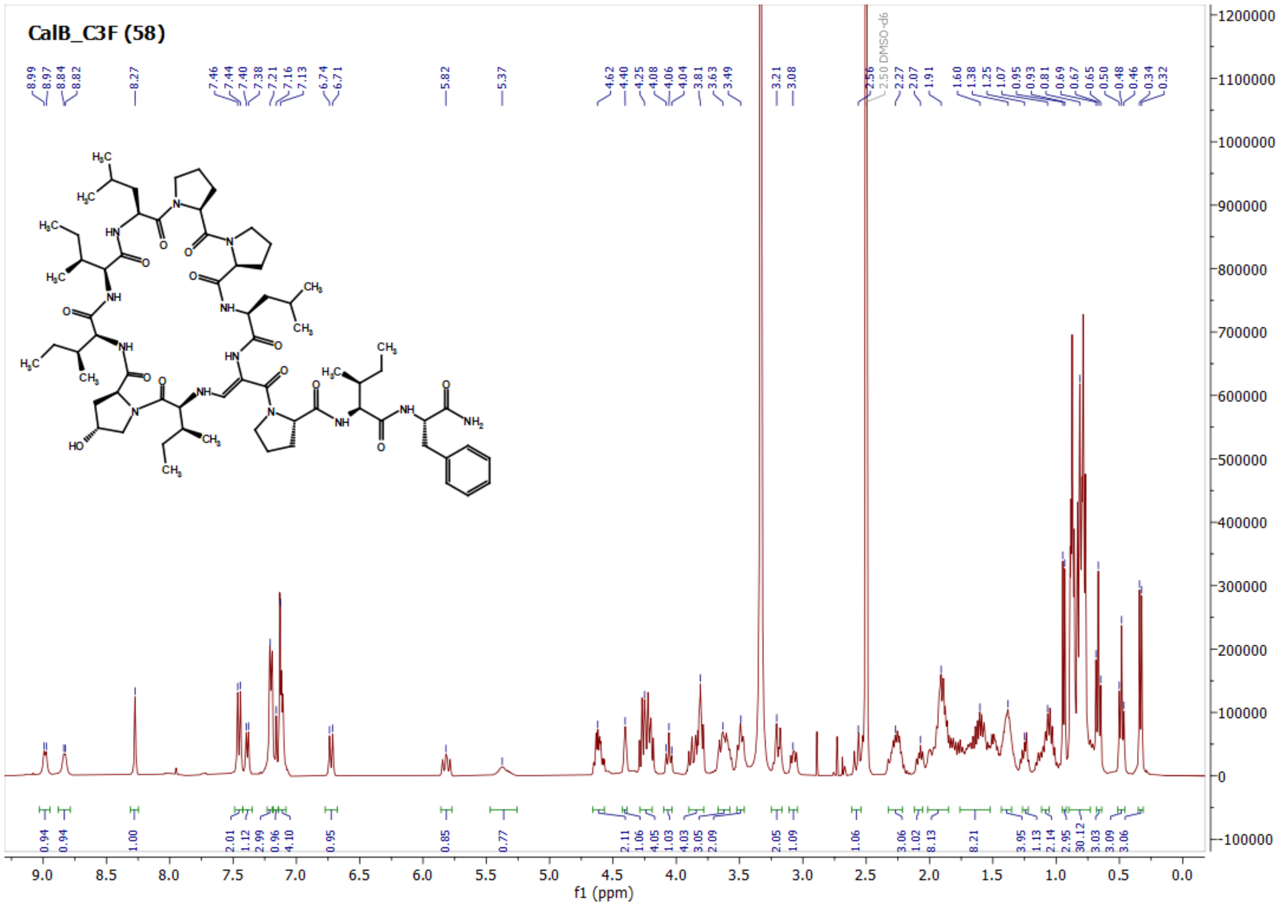
CalB_C3F (58)
f1 (ppm)

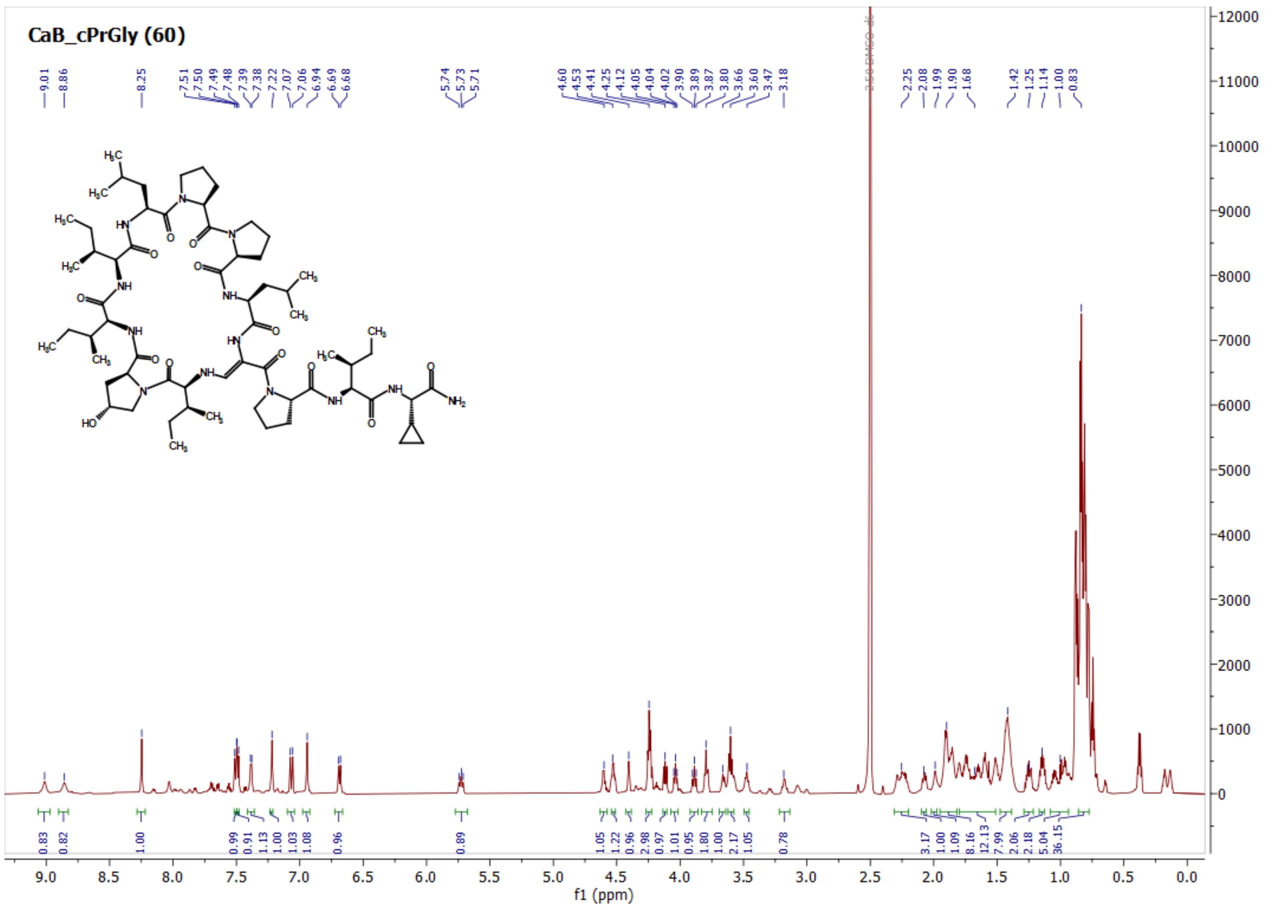
CaB_cPrGly (60)
f1 (ppm)

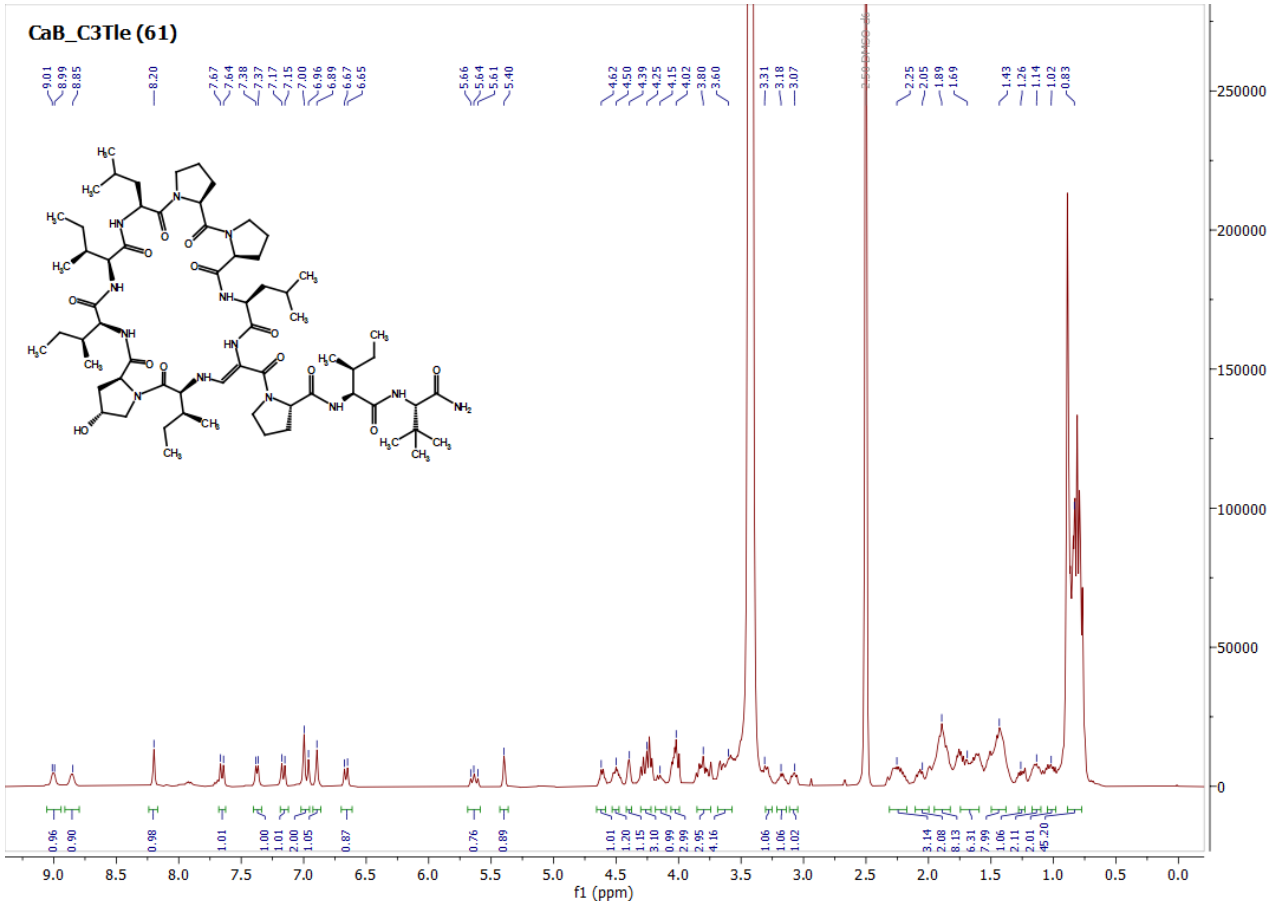
CaB_C3Tle (61)
f1 (ppm)

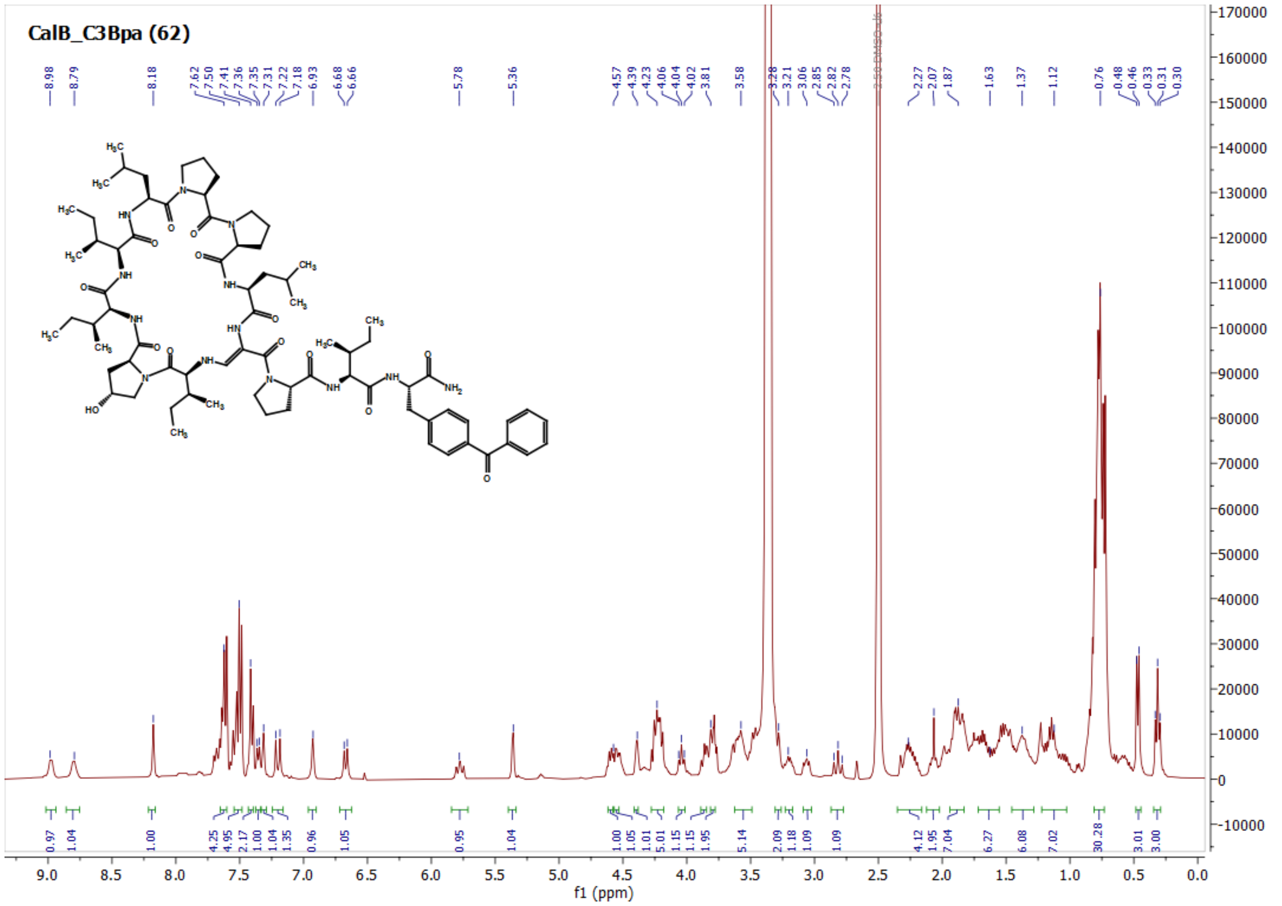
CalB_C3Bpa (62)
f1 (ppm)

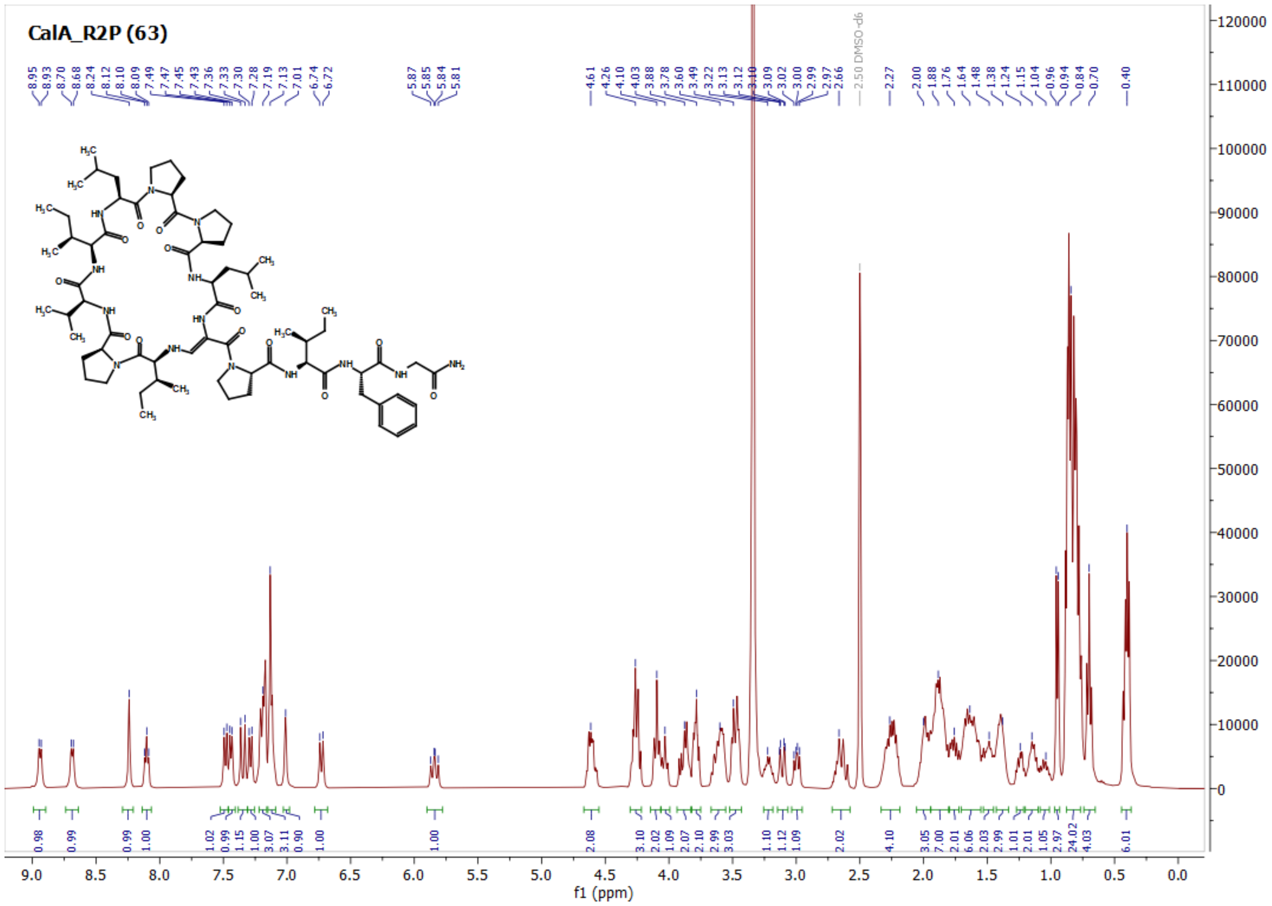
CalA_R2P (63)
f1 (ppm)

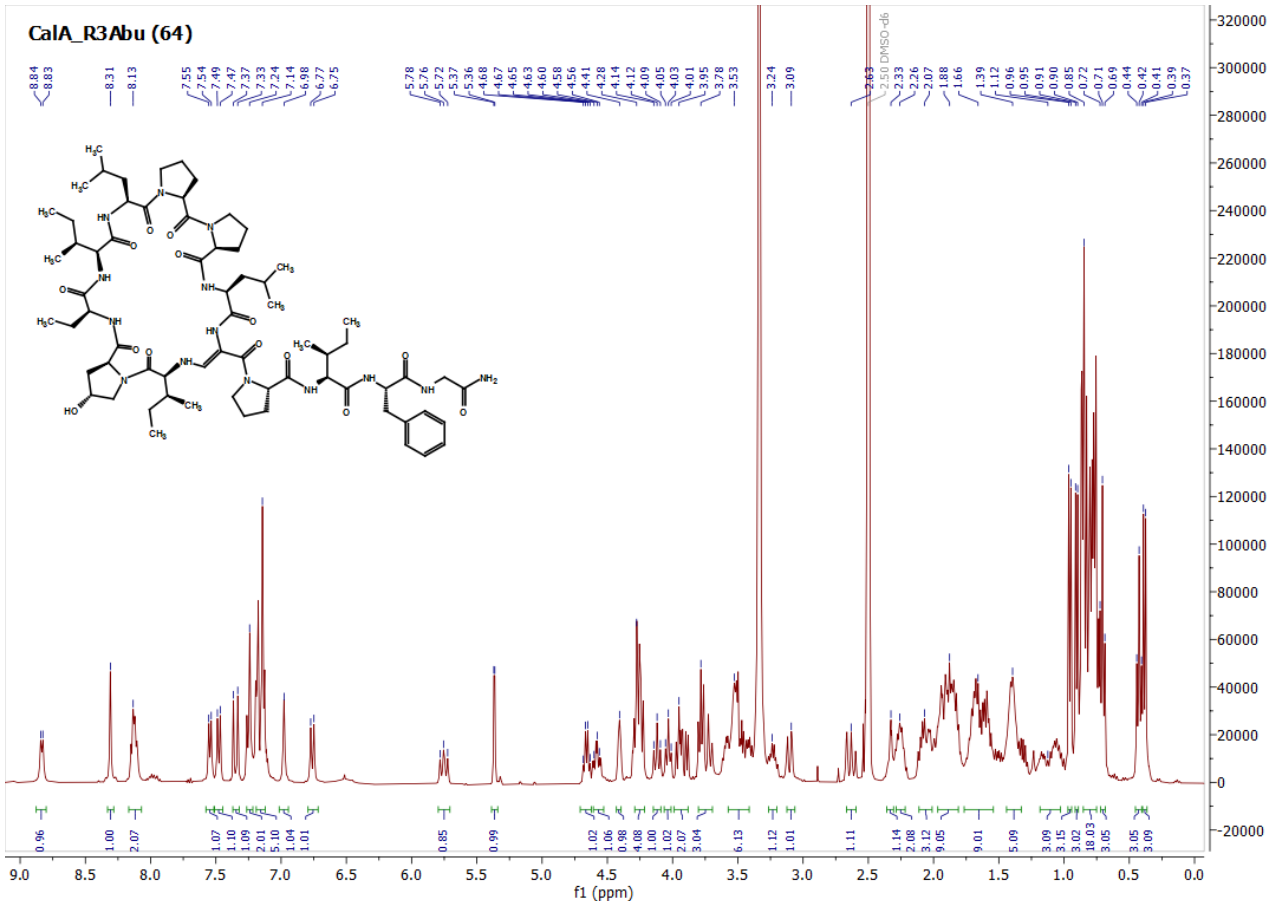
CalA_R3Abu (64)
f1 (ppm)

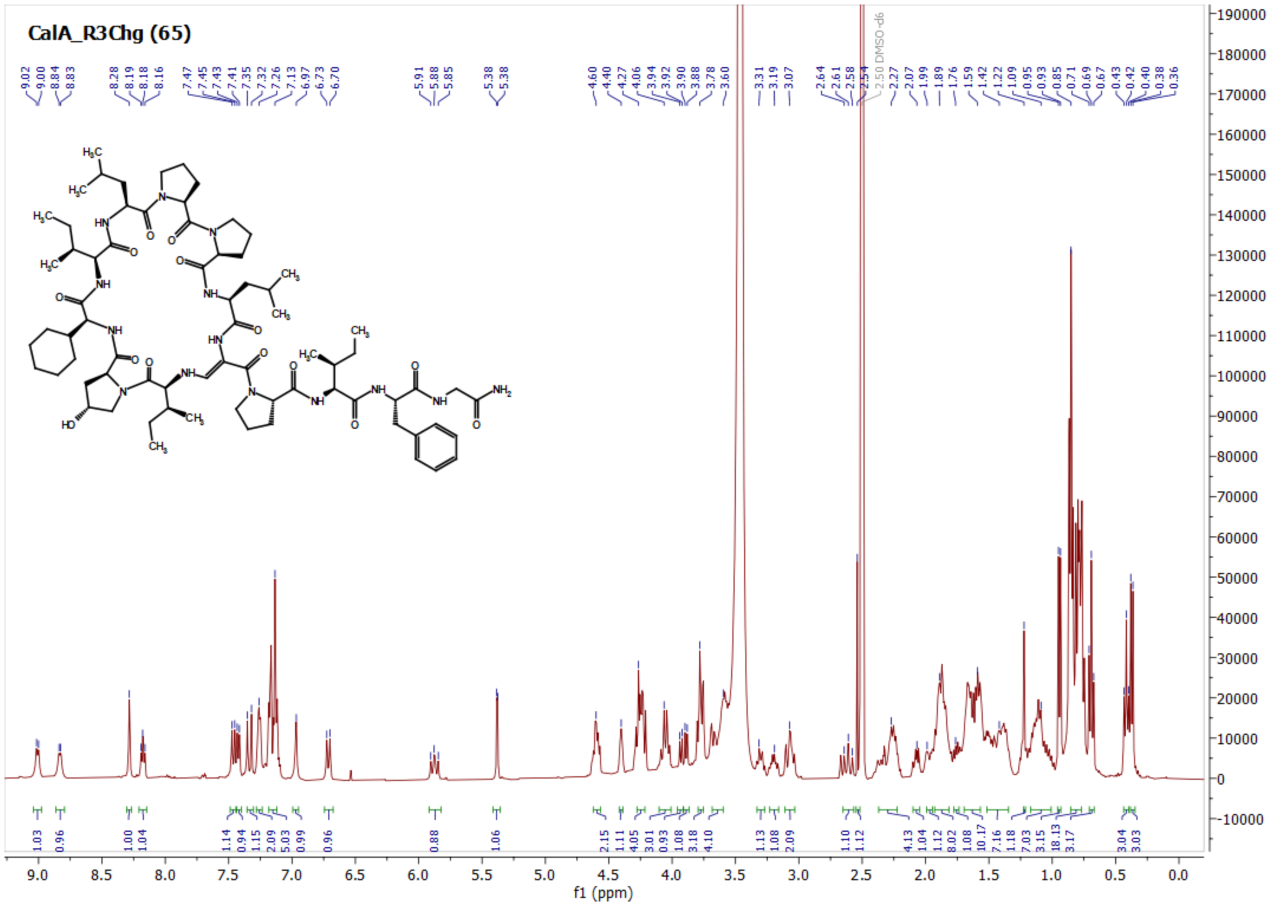
CalA_R3Chg (65)
2.50 DMSO-d6
f1 (ppm)

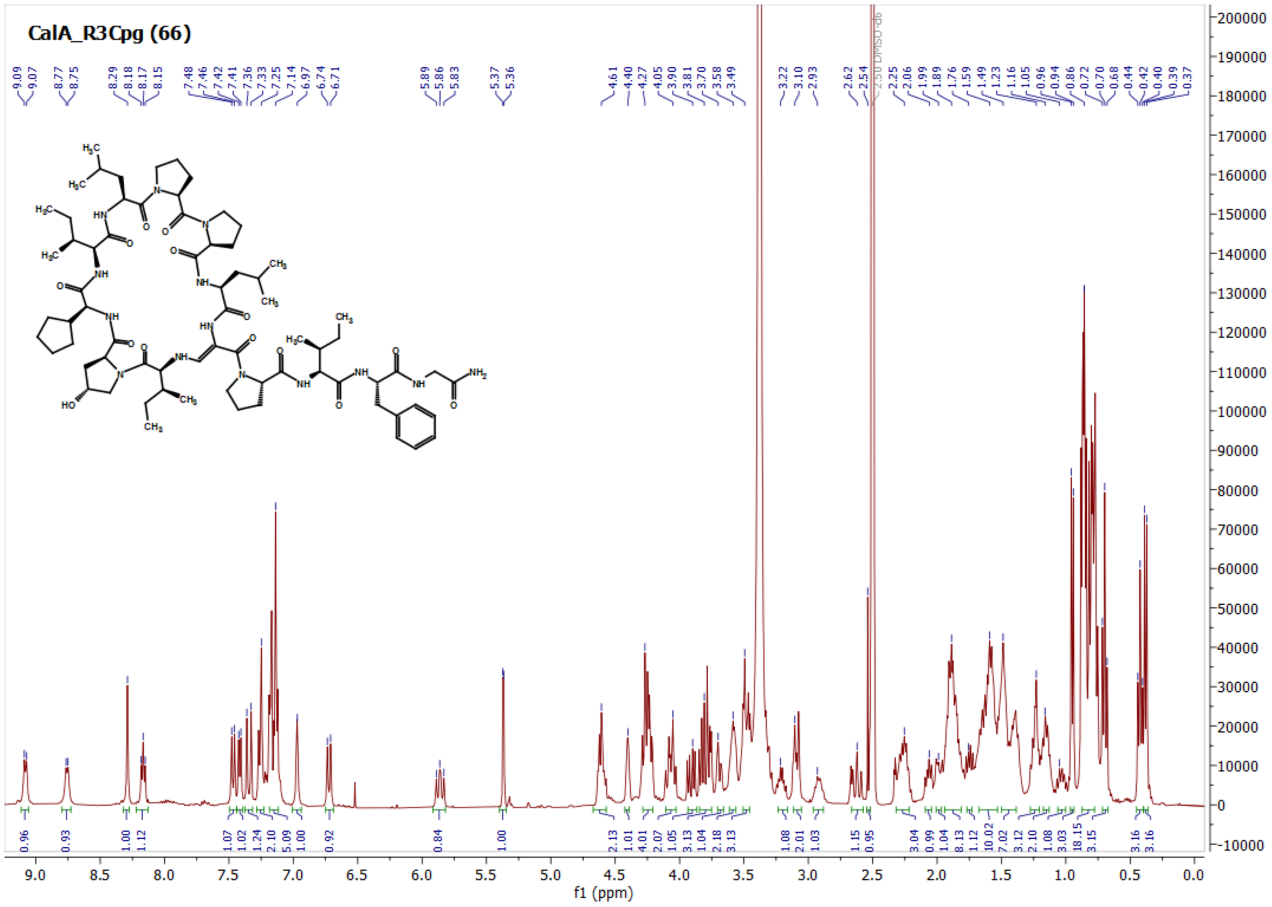
CalA_R3Cpg (66)
2.50 DMSO-d6
f1 (ppm)

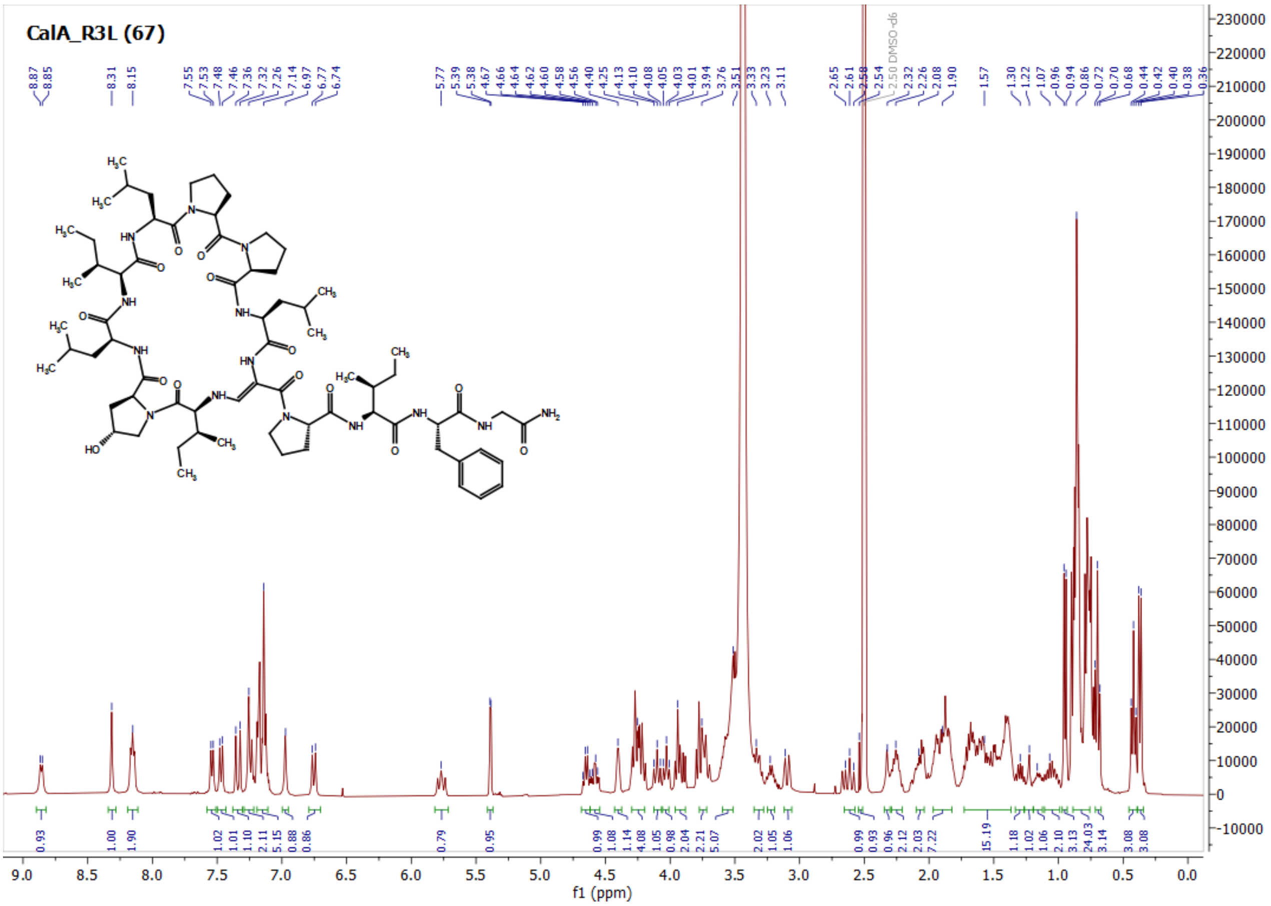
CalA_R3L (67)
f1 (ppm)

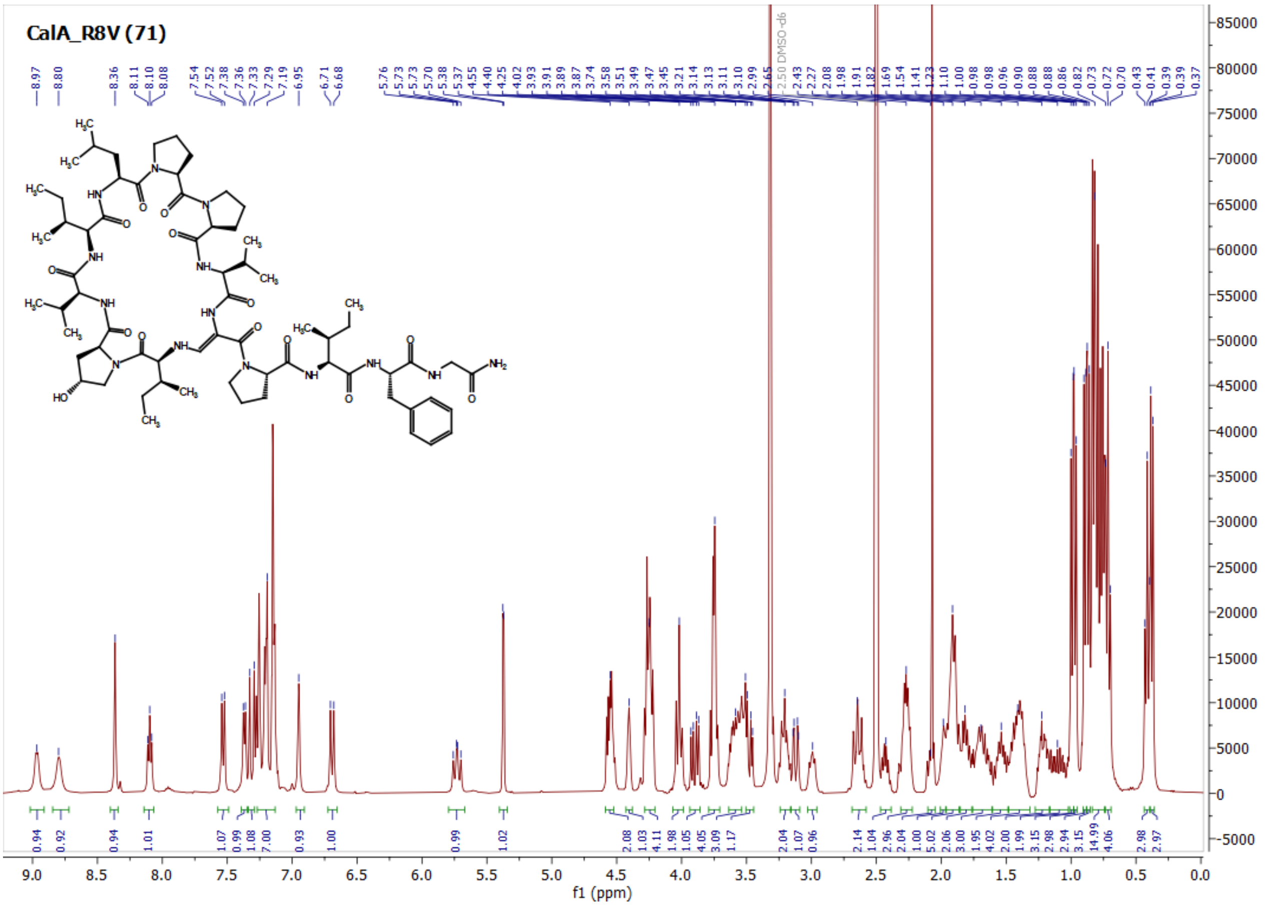
CalA_R8V (71)
f1 (ppm)

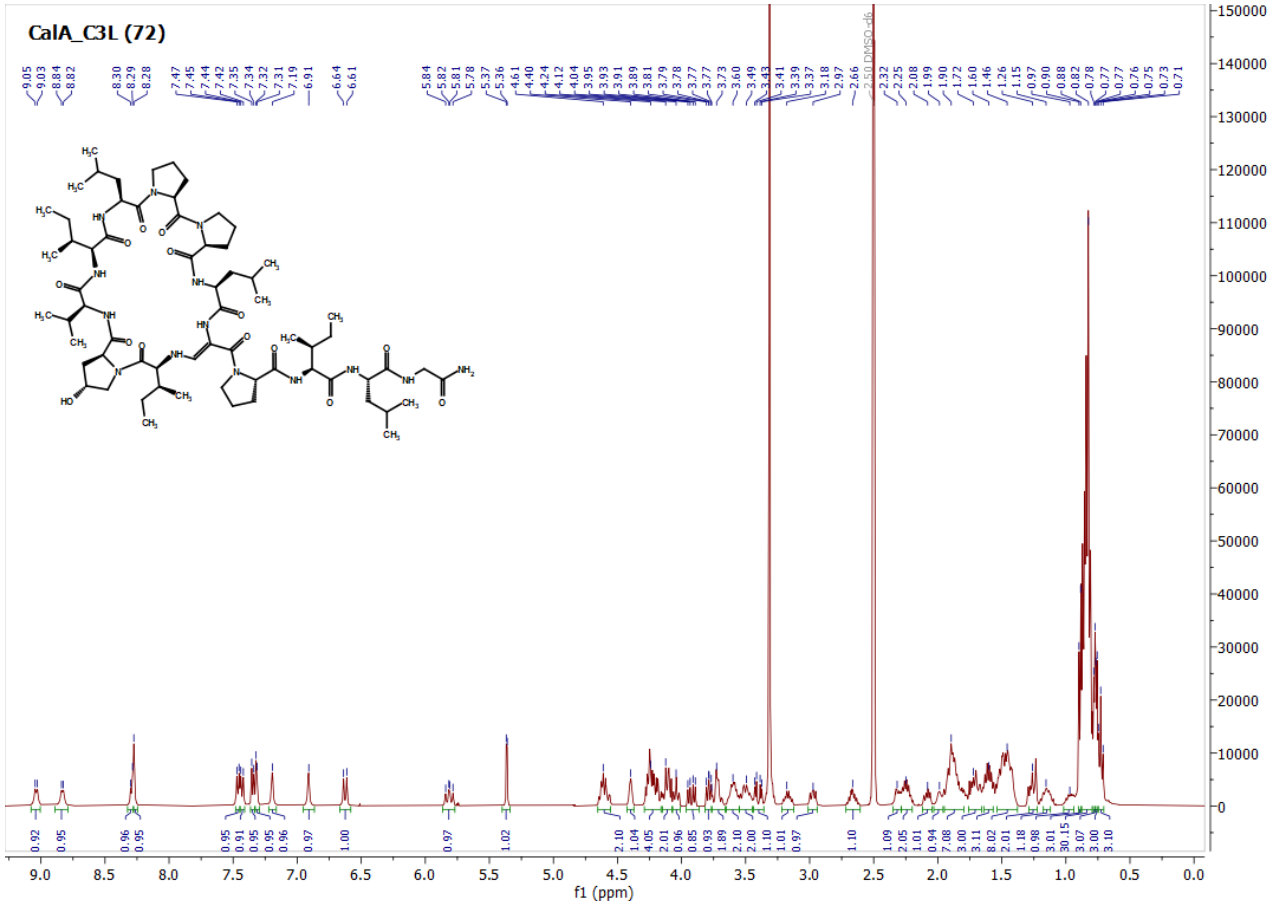
CalA_C3L (72)
f1 (ppm)

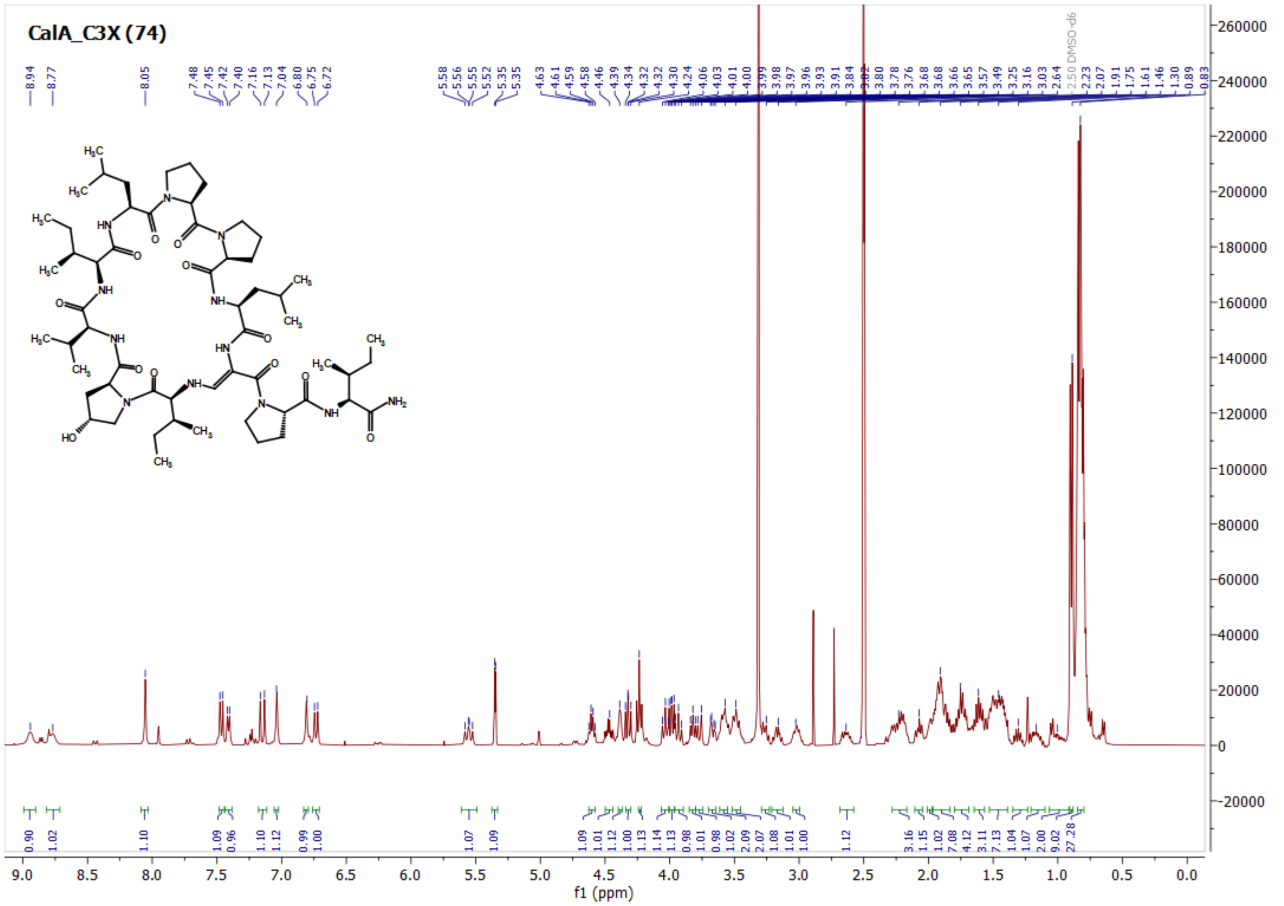
CalA_C3X (74)
f1 (ppm)

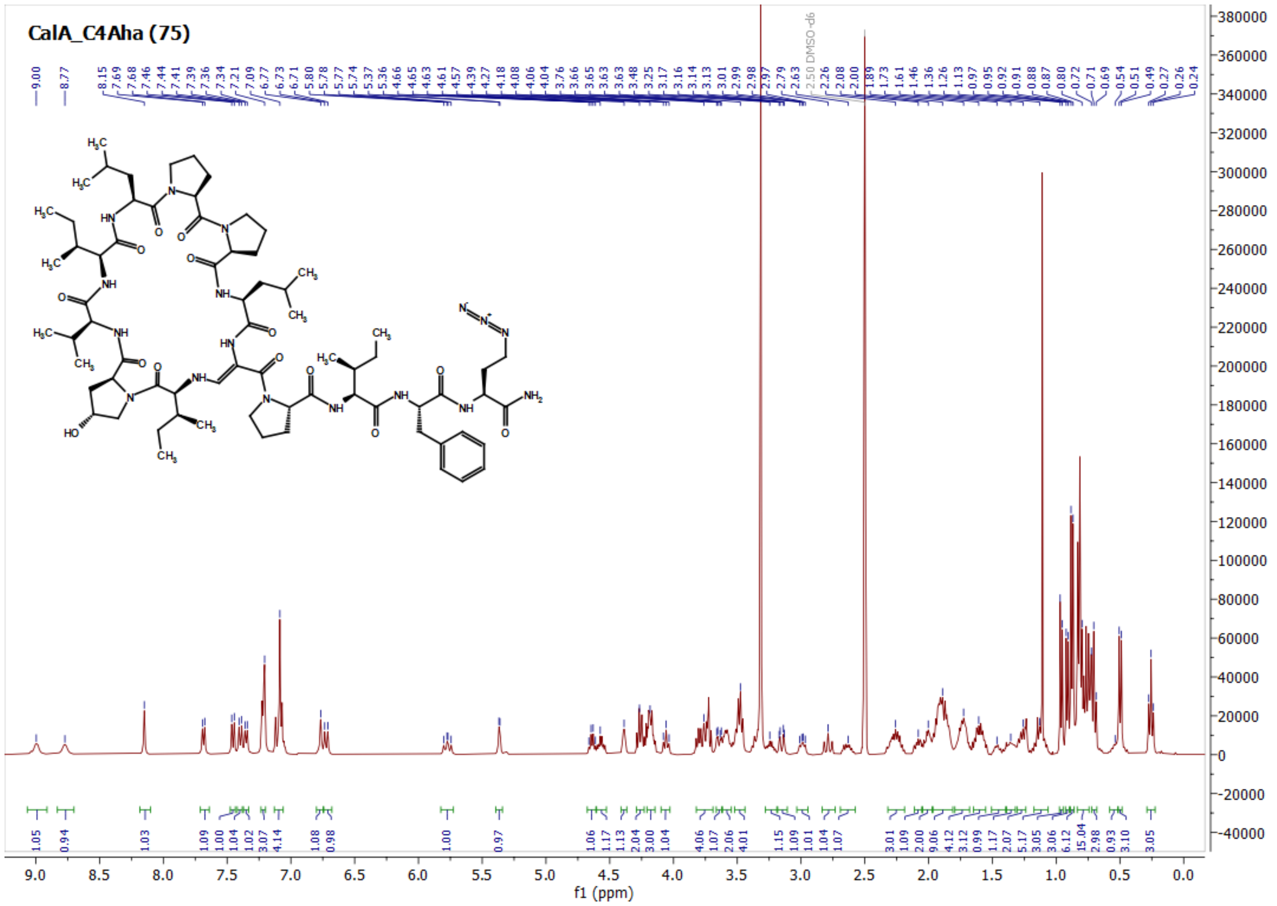
CalA_C4Aha (75)
2.50 DMSO-d6
f1 (ppm)

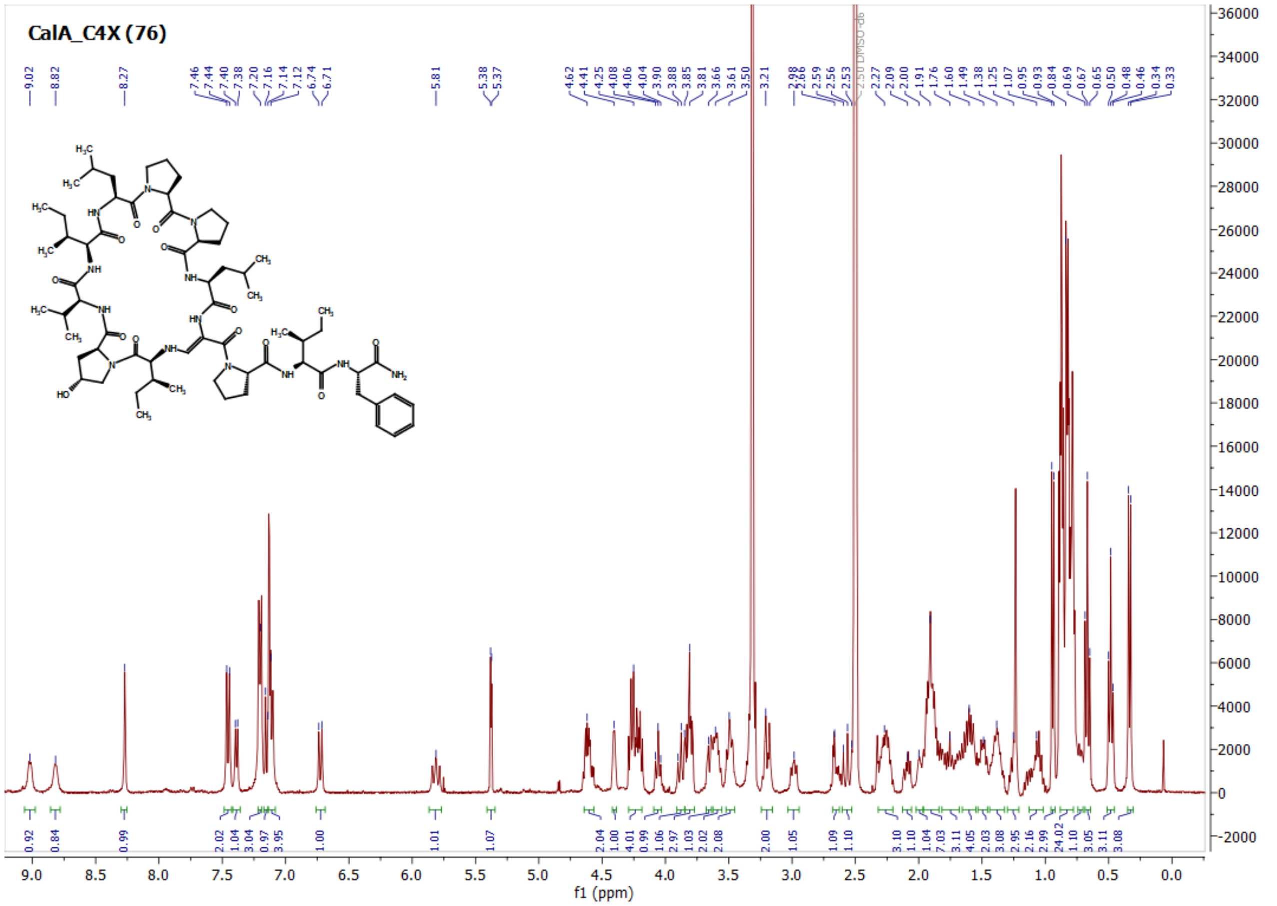
CalA_C4X (76)
2.50 DMSO-d6
f1 (ppm)

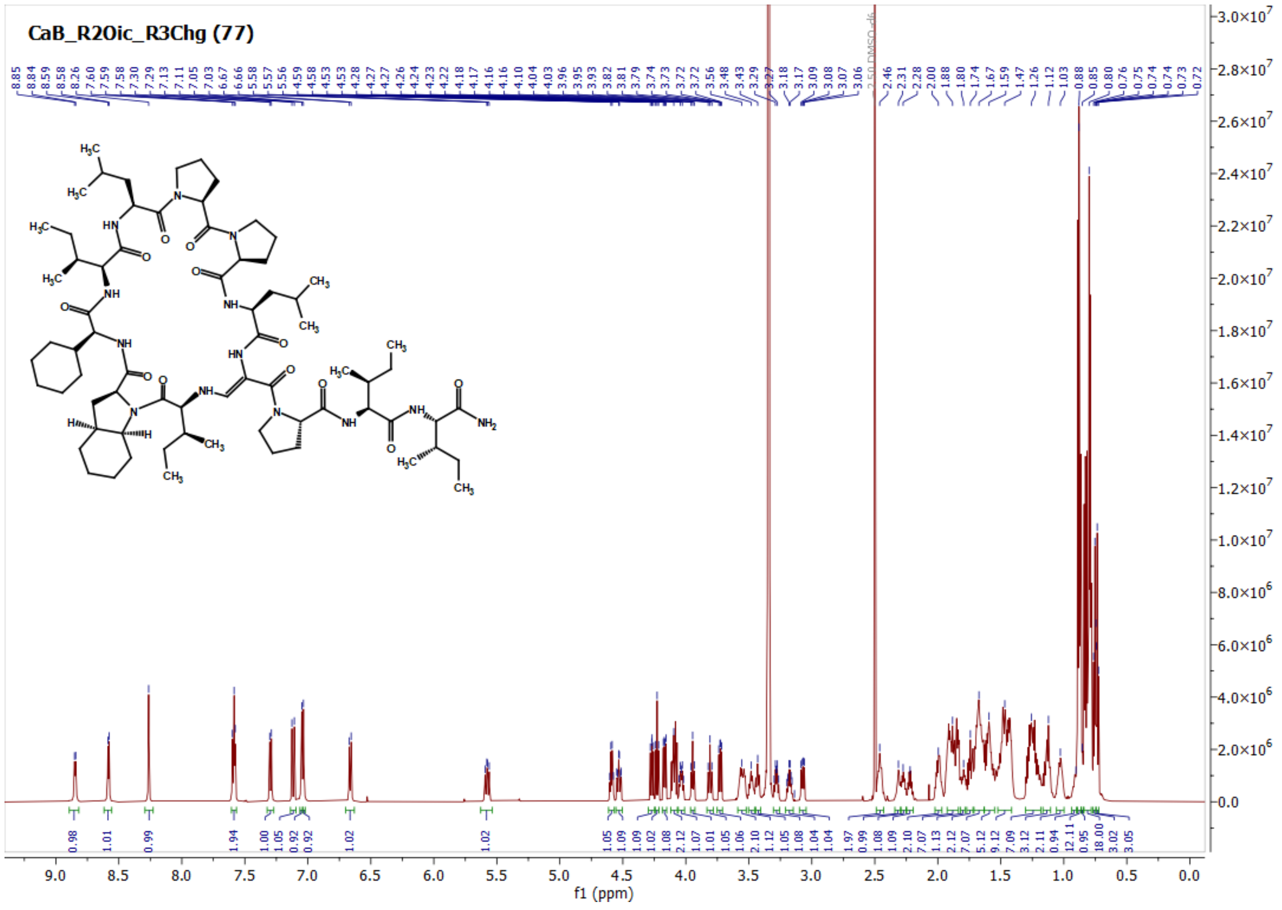
CaB_R2Oic_R3Chg (77)
f1 (ppm)

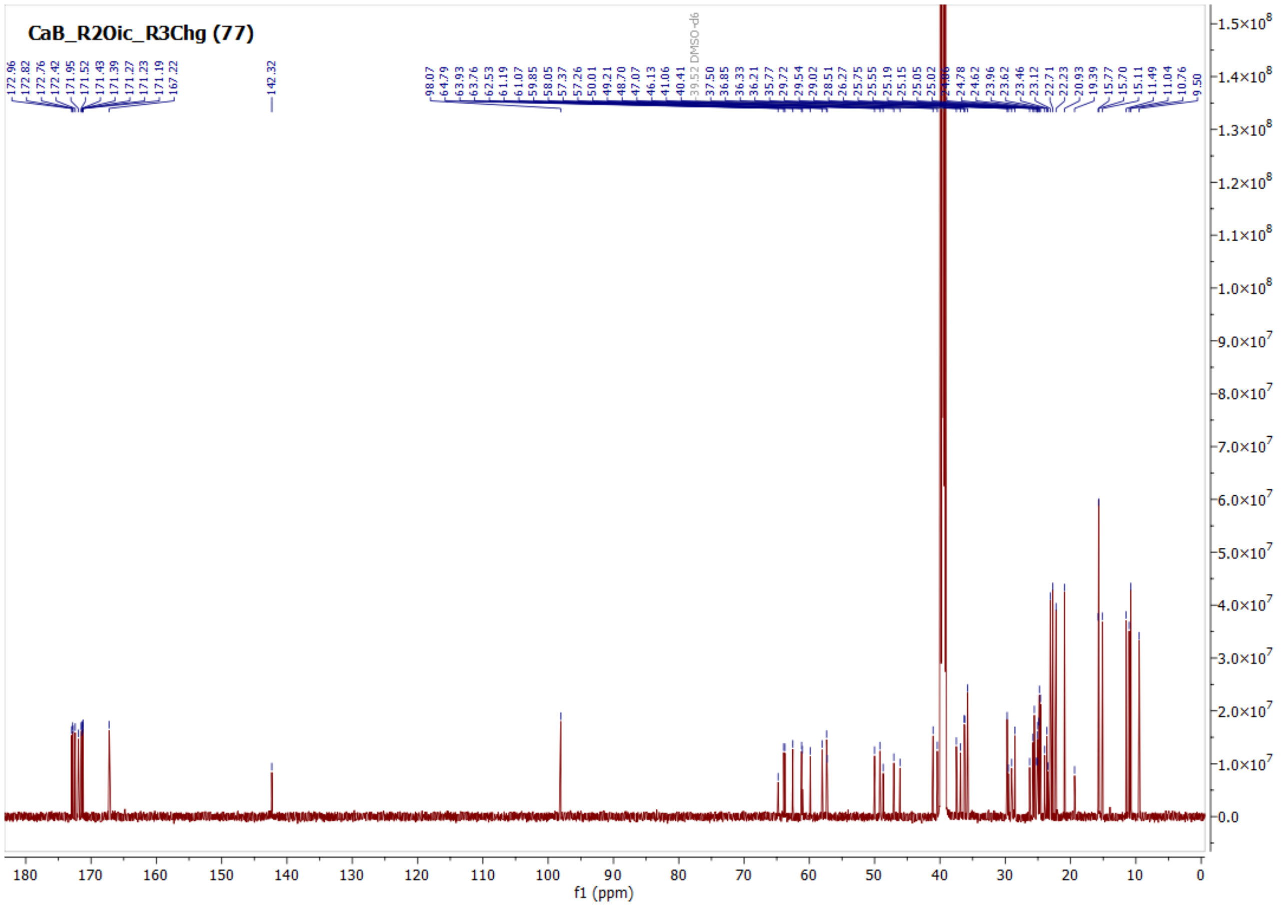
CaB_R2Oic_R3Chg (77)
f1 (ppm)

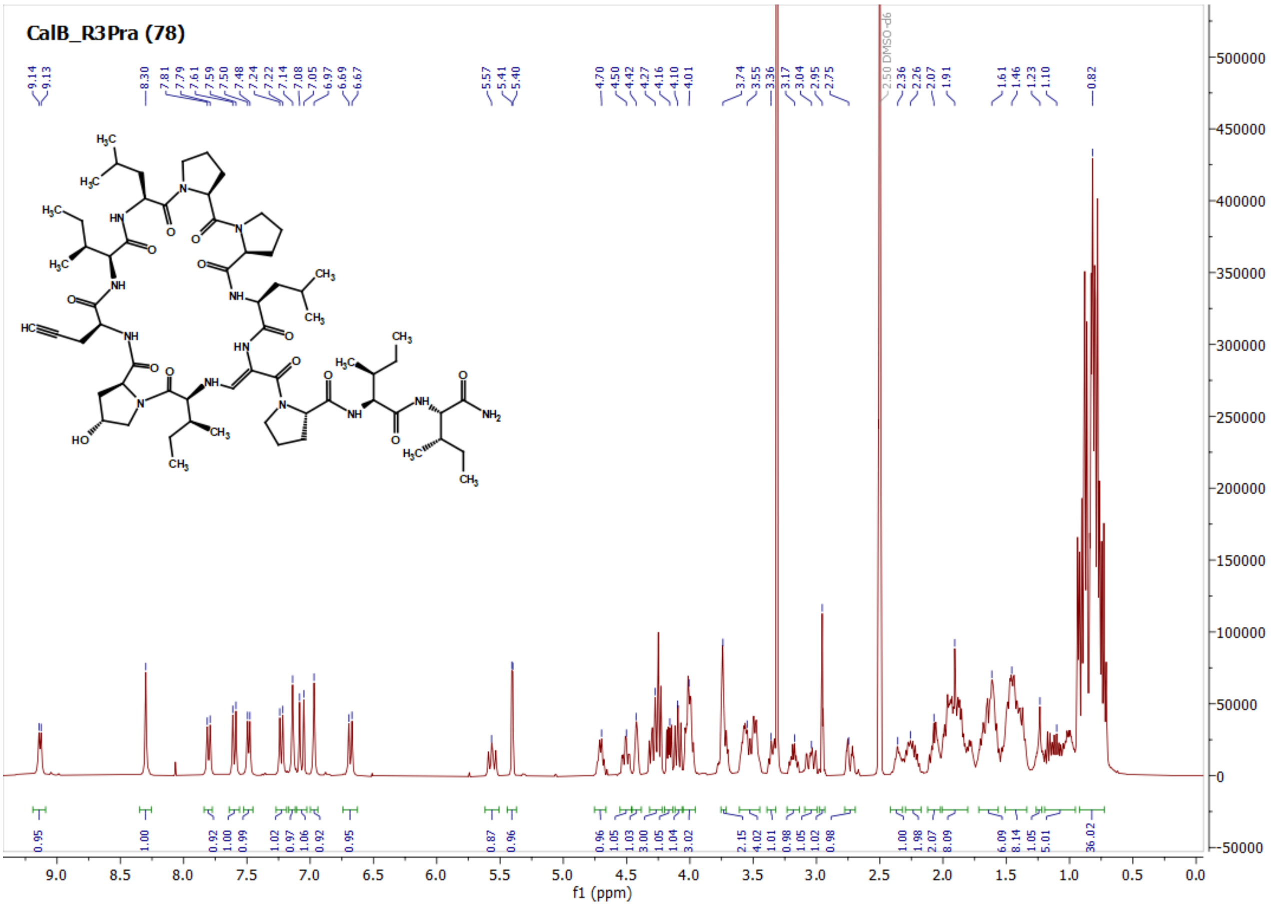
CalB_R3Pra (78)
2.50 DMSO-d6
f1 (ppm)

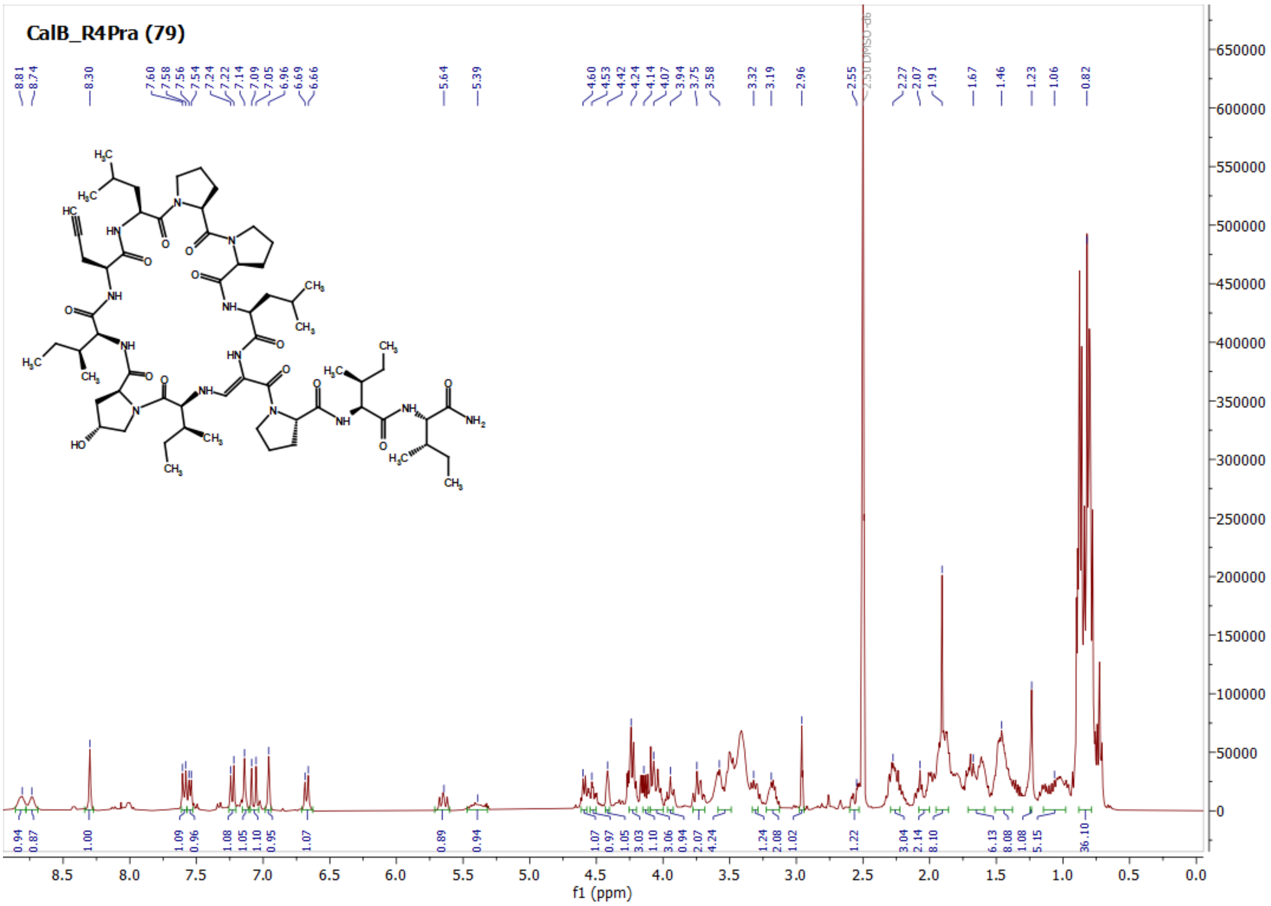
CalB_R4Pra (79)
2.50 DMSO-d6
f1 (ppm)

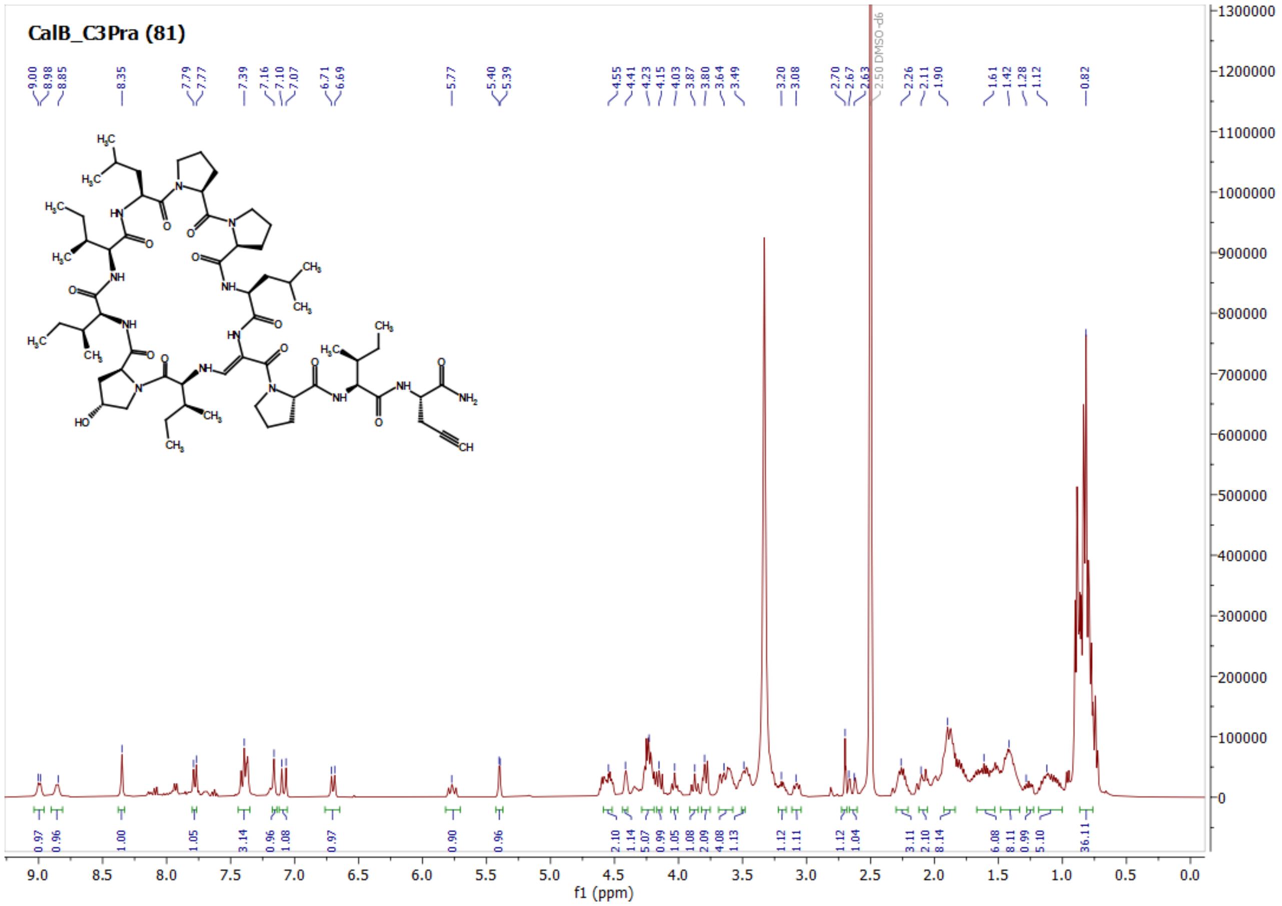
CalB_C3Pra (81)
f1 (ppm)

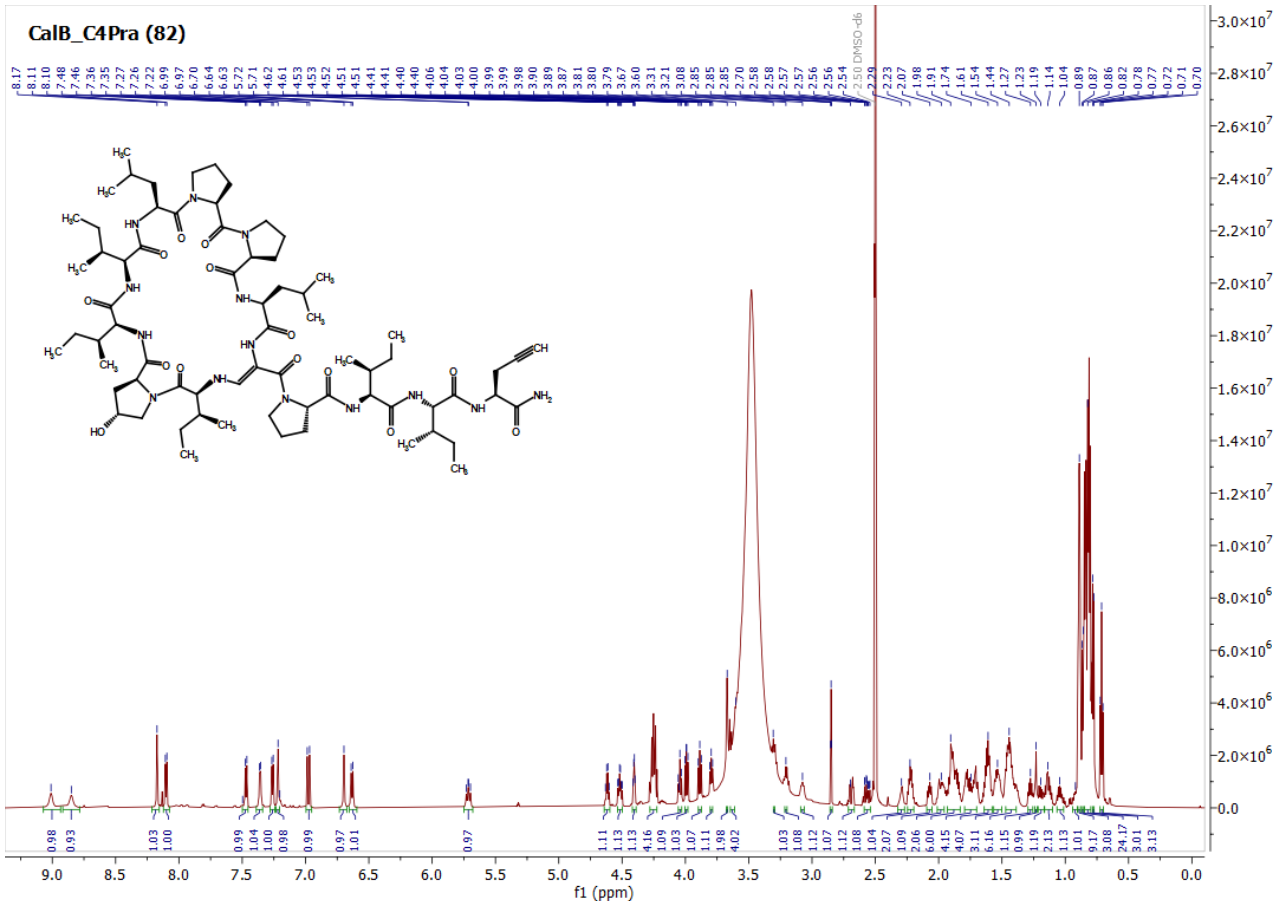
CalB_C4Pra (82)
f1 (ppm)

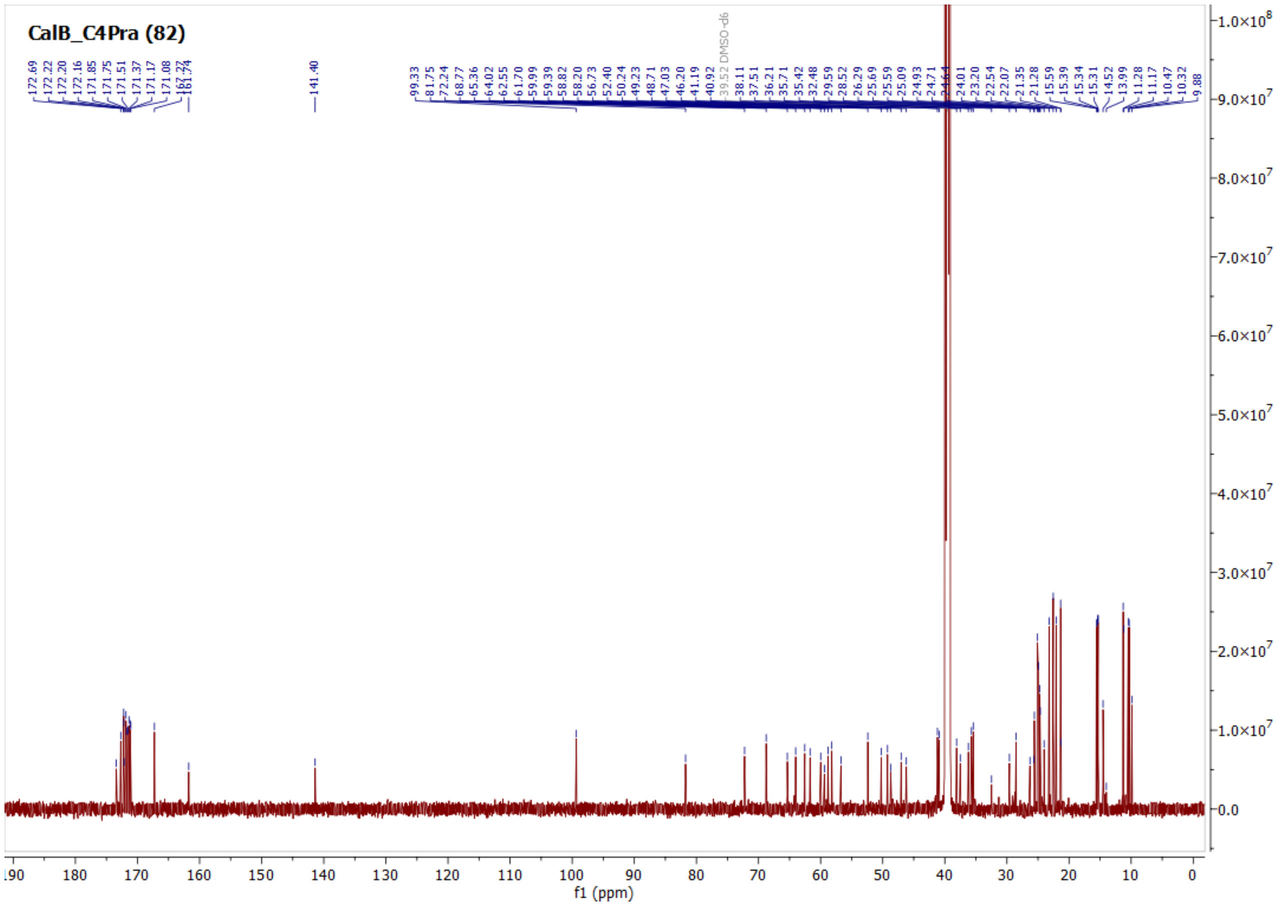
CalB_C4Pra (82)
39.52 DMSO-d6
f1 (ppm)

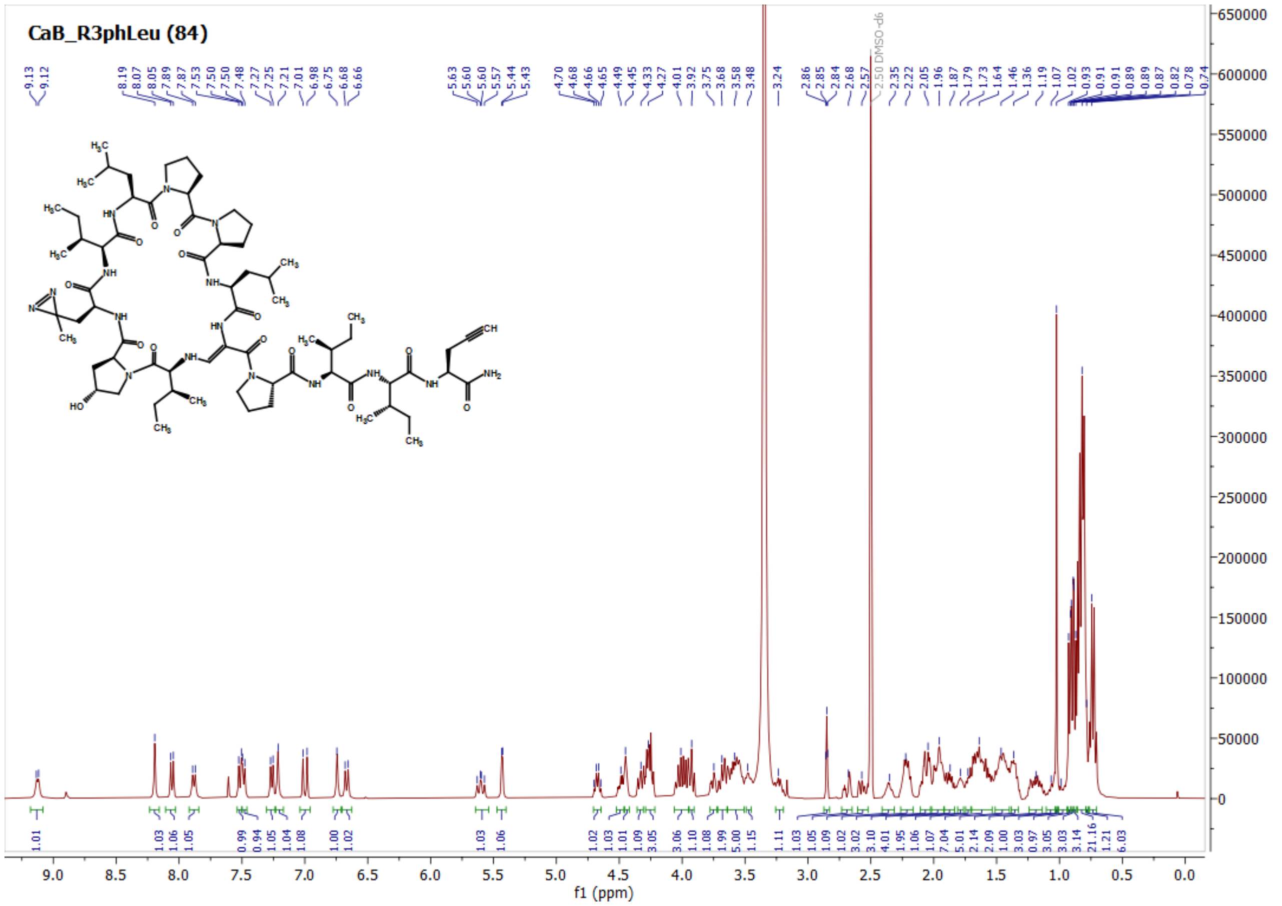
CaB_R3phLeu (84)
2.50 DMSO-d6
f1 (ppm)

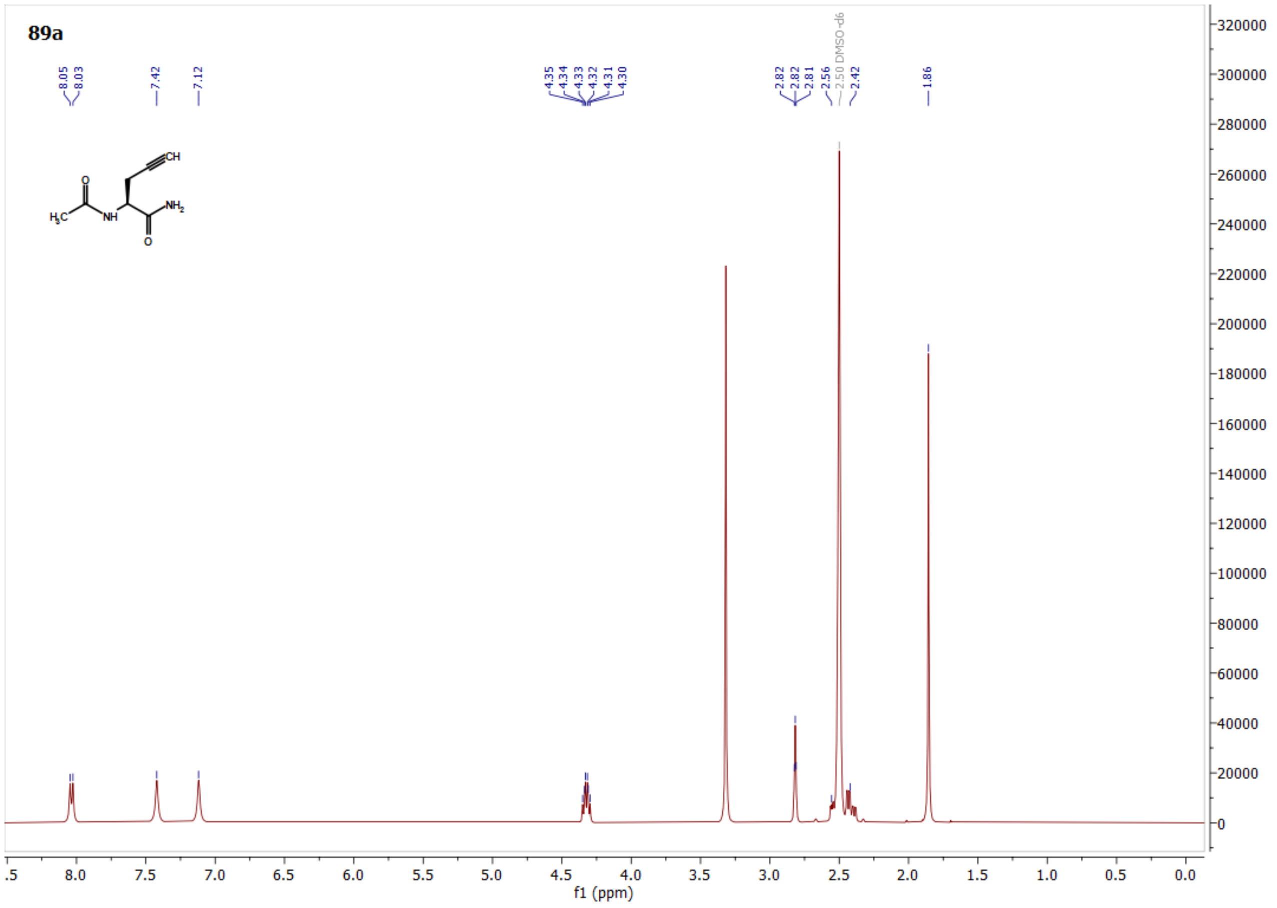
89a
8.05
8.03
7.42
7.12
4.35
4.34
4.33
4.32
4.31
4.30
2.82
2.82
2.81
2.56
2.50 DMSO-d6
2.42
1.86
CH
O
H3C
NH
NH2
O
f1 (ppm)

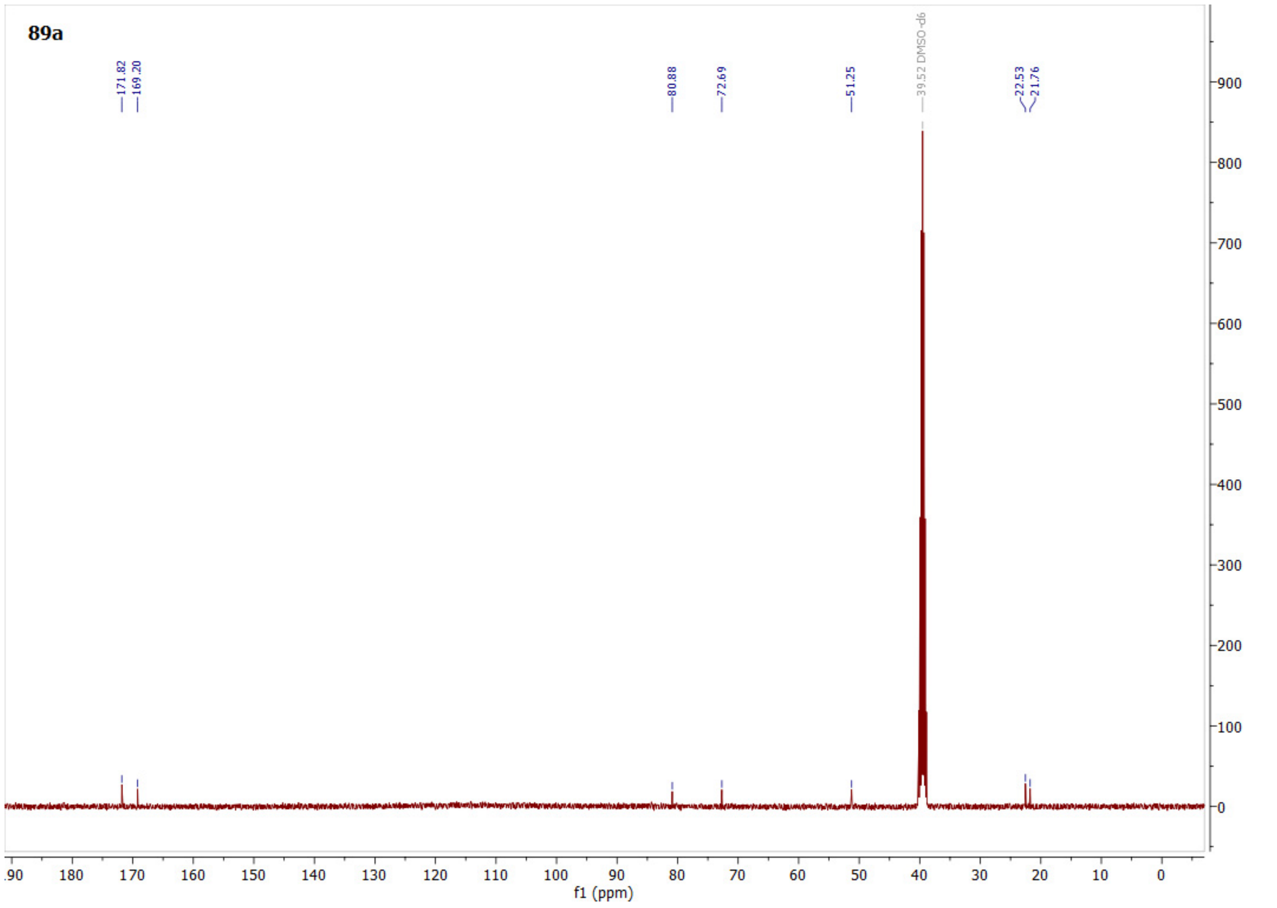
89a
171.82
169.20
80.88
72.69
51.25
39.52 DMSO-d6
22.53
21.76
f1 (ppm)

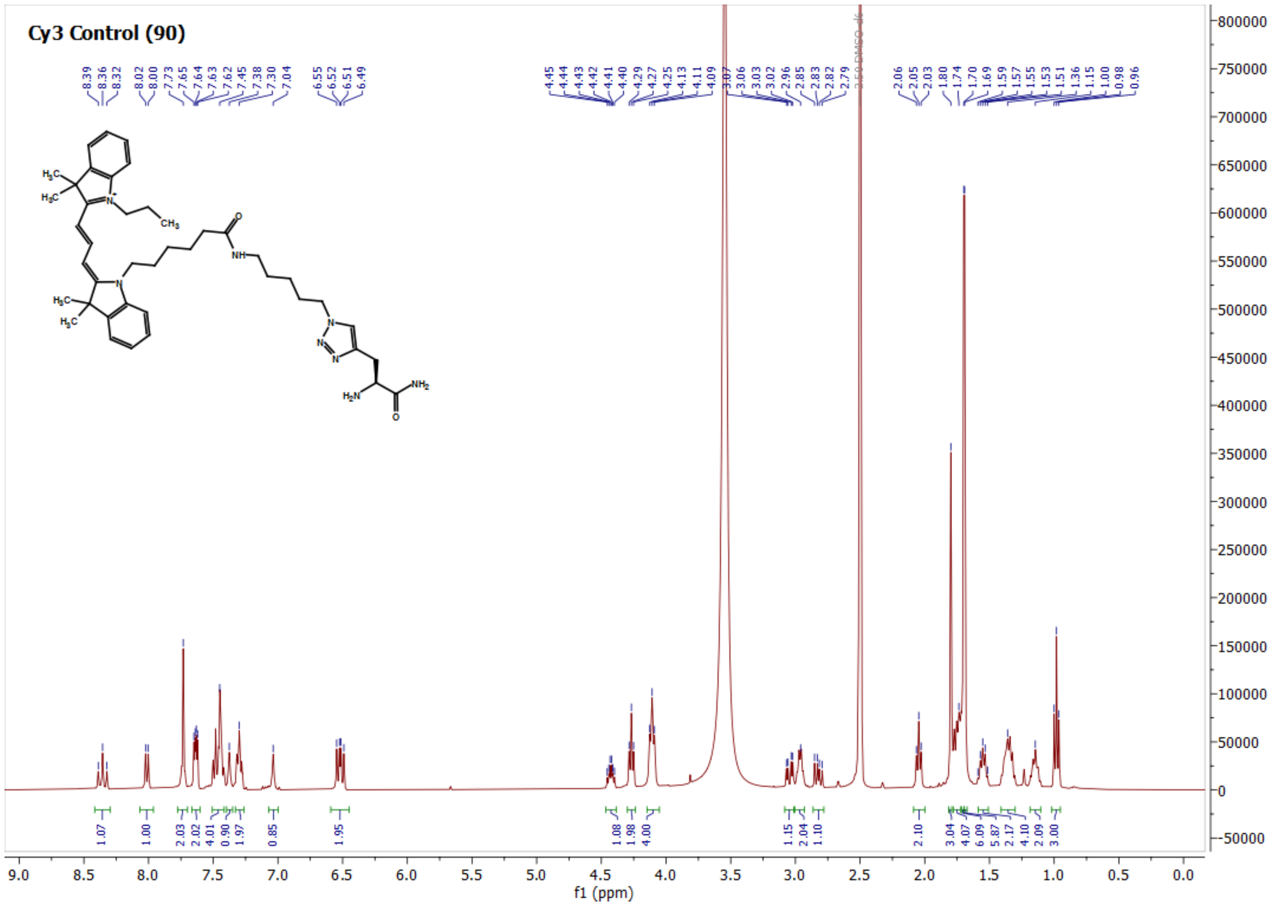
Cy3 Control (90)
f1 (ppm)
